# Phylogenomics of the mega genus *Bulbophyllum* (Orchidaceae) with implications for infrageneric classification in the Asian clade

**DOI:** 10.64898/2026.03.30.715161

**Authors:** Consolata Nanjala, Lalita Simpson, Ai-Qun Hu, Vidushi Patel, James A. Nicholls, Stephen J. Bent, Stephan W. Gale, Gunter A. Fischer, Stephanie Goedderz, André Schuiteman, Darren M. Crayn, Mark A. Clements, Katharina Nargar

**Author notes:** Correspondence: Consolata Nanjala, Katharina Nargar.

## Abstract

Understanding evolutionary relationships in hyperdiverse plant groups remains a major challenge in systematics. The orchid genus *Bulbophyllum*, the second largest genus of flowering plants, represents an exceptional example of phylogenetic and morphological complexity. Relationships, particularly within the species-rich Asian clade, have remained poorly resolved due to extensive morphological variation and limited resolution in previous phylogenetic studies.

Here, we reconstructed phylogenetic relationships using 63 plastid genes from 355 specimens representing 322 species and 65 of the 97 recognised sections of *Bulbophyllum*. Our analyses confirmed that the genus comprises five major evolutionary lineages comprised of species predominantly from Australasia, Madagascar, Continental Africa, Neotropics, and Asia. We provide the first robust phylogenetic evidence for a dichotomous split within the Asian clade into two well-supported lineages: the Asian-Malesian clade and the Malesian-Papuasian clade, with the latter containing a strongly supported Papuasian subclade. Additionally, this study supports the monophyly of several currently recognised sections while clarifying relationships in previously problematic groups.

This study provides the most comprehensive plastid-based phylogenomic framework for *Bulbophyllum* to date and establishes a foundation for future taxonomic revision and integrative analyses of diversification and trait evolution within this hyperdiverse genus.

## INTRODUCTION

Orchidaceae is one of the most rapidly radiating groups (Serna-Sánchez et al., 2021; Pérez-Escobar et al., 2024) comprising ca. 31,482 species and around 758 genera (WFO, 2025). Orchids exhibit a wide range of morphological, physiological, and ecological adaptations including epiphytism and CAM photosynthesis (Zhang et al., 2023), specialisation with mycorrhizal fungi (Jacquemyn and Merckx, 2019), and diverse pollination strategies including deceit pollination (Johnson et al., 2013), all of which are presumed to be among the drivers of their extraordinary diversity (Givnish et al., 2015). However, the relative contributions of these factors to orchid rarity and diversification remain poorly understood (Gravendeel et al., 2004; Tremblay et al., 2005; Crain and Tremblay, 2014; Ackerman et al., 2023), primarily due to the complex evolutionary and biogeographical history of the family and unresolved phylogenetic relationships among species (Neubig et al., 2009; Serna-Sánchez et al., 2021). Addressing these gaps requires comprehensive systematic research leveraging robust, well-sampled phylogenies to accurately trace the evolutionary history of the family particularly at the genus and species levels (Zhang et al., 2023; Pérez-Escobar et al., 2024). However, phylogenetic resolution at lower taxonomic levels in Orchidaceae has remained challenging due to the limited genetic variation in commonly used markers (e.g., nuclear ITS and plastid *rbcL*, *matK*, and intergenic spacers) (Cameron et al., 1999; Chase et al., 2003; Li et al., 2016; Hu et al., 2020). The advent of high-throughput sequencing is now enabling rapid advancements in our understanding of orchid relationships and diversification (Givnish et al., 2015; Xiang et al., 2016; Serna-Sánchez et al., 2021; Pérez-Escobar et al., 2024).

The genus *Bulbophyllum* Thouars (Orchidaceae) is the second-largest genus of flowering plants and the largest orchid genus, comprising approximately 2,190 species (WFO, 2025). The genus is a prime example of phylogenetic complexity, exhibiting remarkable morphological and ecological diversity (Gamisch and Comes, 2019; Vermeulen et al., 2014; Vermeulen et al., 2015; Vermeulen et al., 2018). This has resulted in various classifications of the group throughout its taxonomical history, which have been largely reliant on morphological traits (Bentham, 1881, 1883; Blume, 1825; Reichenbach, 1857; Dressler, 1974; Seidenfaden, 1979; Seidenfaden and Wood, 1992; Dressler, 1993; Vermeulen, 2014; Vermeulen et al., 2014; Vermeulen et al., 2015; Chase et al., 2015). However, assessment of competing systematic classifications of the mega genus has been hampered by a lack of robust molecular phylogenetic evidence.

*Bulbophyllum* has a pantropical distribution spanning continental Africa, Madagascar, the Americas, the Asia-Pacific and the Australasian (Australia and New Zealand) regions, although species richness is unevenly distributed within the genus (Gravendeel, 2014). The highest diversity of *Bulbophyllum* occurs in Southeast Asia which contains ca. 1600 species (Gravendeel, 2014), with New Guinea having the highest species richness in the region, harbouring approximately 660 species (Cribb and Hermans, 2009; Cámara-Leret et al., 2020). *Bulbophyllum* species can be found in a variety of habitats, ranging from seasonally dry forests in the tropical and subtropical lowlands to wet montane cloud forests (Vermeulen, 2014; Vermeulen et al., 2015). Species in this genus are generally epiphytic, sometimes lithophytic, or rarely terrestrial herbs having pseudobulbs of a single internode and leaves which are mainly persistent and sometimes deciduous (Gravendeel and Vermeulen, 2014). Typically, one or two (rarely more) non-sheathing leaves are present at the top of the pseudobulb, ranging from a few mm to more than 1.5 m in length, while the unbranched inflorescence, originating from the base of the pseudobulb or the rhizome, bears one-to-many flowers (Fig. 1; Gravendeel and Vermeulen, 2014; Vermeulen, 2014). In addition to their varied growth habits, several *Bulbophyllum* species have been shown to exhibit fly pollination syndrome (Borba et al., 1999; Tan and Nishida, 2000; Ong, 2025).

**Figure 1.**
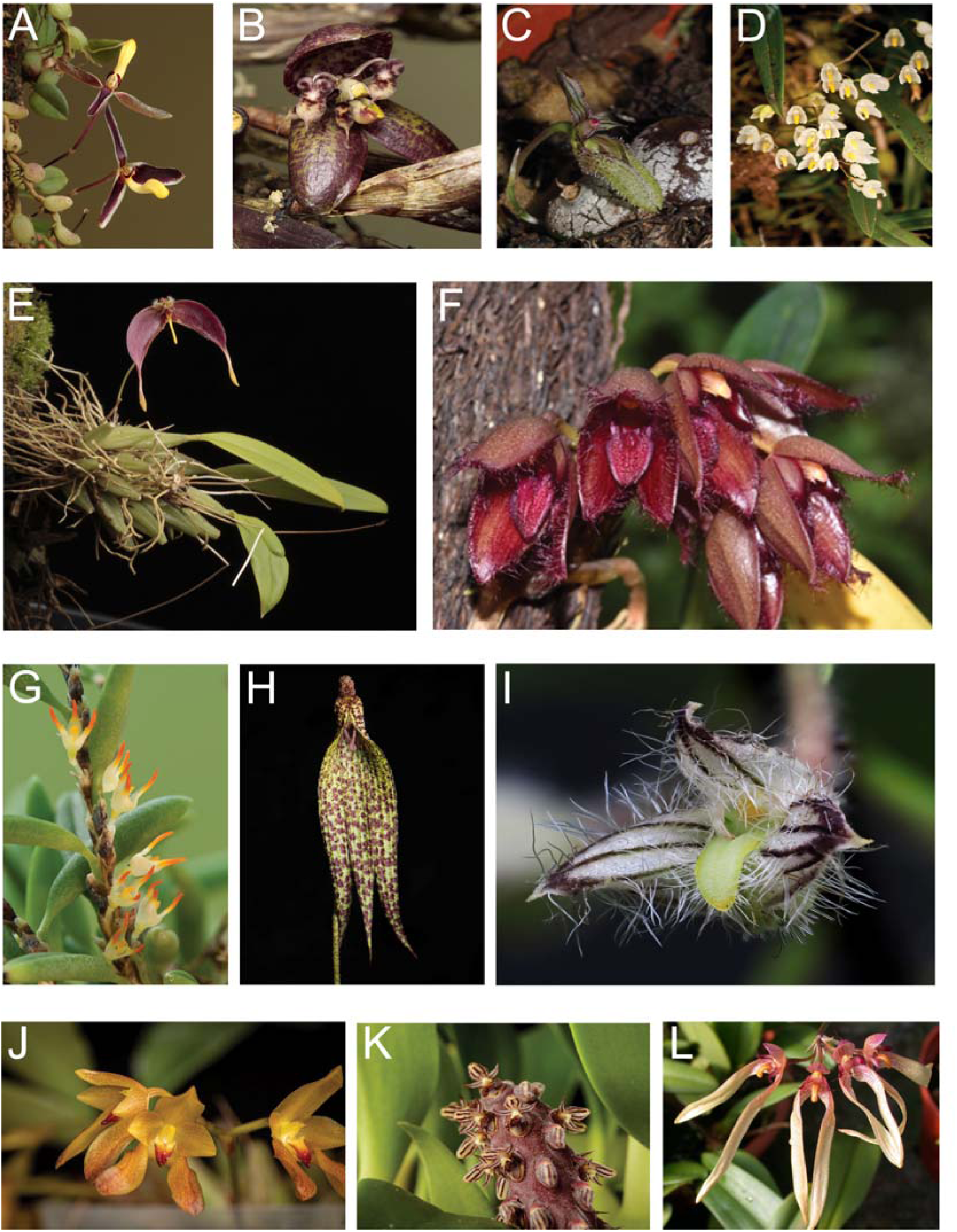
Morphological diversity within the genus *Bulbophyllum*. (A) *Bulbophyllum alkmaarense* J.J.Sm. (sect. *Codonosiphon* Schltr.); (B) *Bulbophyllum macrorhopalon* Schltr. (sect. *Epicrianthes* (Blume) Hook.f.); (C) *Bulbophyllum polliculosum* Seidenf. (sect. *Drymoda* (Lindl.) J.J.Verm.); (D) *Bulbophyllum lageniforme* F.M.Bailey (sect. *Adelopetalum* (Fitzg.) J.J. Verm.); (E) *Bulbophyllum nasica* Schltr. (sect. *Polymeres* (Blume) J.J.Verm. & O’Byrne); (F) *Bulbophyllum dayanum* Rchb.f. (sect. *Achrochaene* (Lindl.) J.J.Verm.); (G) *Bulbophyllum gadgarrense* Rupp (sect. *Oxysepala* Benth. & Hook.f.); (H) *Bulbophyllum muricatum* J.J.Sm. (sect. *Monosepalum* (Schltr.) J.J.Sm.); (I) *Bulbophyllum lindleyanum* Griff. (sect. *Hirtula* Ridl.); (J) *Bulbophyllum cootesii* M.A.Clem. (sect. *Lepidorhiza* Schltr.); (K) *Bulbophyllum saurocephalum* Rchb.f. (sect. *Saurocephalum* J.J.Verm.); (L) *Bulbophyllum longiflorum* Thouars (sect. *Cirrhopetalum* (Lindl.) Rchb.f). Photo credits: Andre Schuiteman (A, B, E, H, I); Knitta Brandorff (C); T. Linderhaus (D, G); Mark Clements (F, L); Eva Lanyi (J); Hannele Lahti (K). **Alt text.** A 12-panel photographic plate showing floral and vegetative morphological diversity across the genus *Bulbophyllum*. Each panel (A-L) displays a close-up of a different *Bulbophyllum* species, highlighting variation in flower shape, size, colour, and inflorescence structure in 12 species representing various sectional groups within the genus.

Initially described by Du Petit-Thouars (1822), early infrageneric classification for *Bulbophyllum* consisted of four unnamed sections based on inflorescence characteristics (Du Petit-Thouars, 1822). Blume (1825) distinguished several genera and sections in what is now considered a single genus *Bulbophyllum* (Gravendeel et al., 2014; Chase et al., 2015). The taxonomic study by Reichenbach (1861) based on characters of the inflorescence divided *Bulbophyllum* into three subgenera (*Uniflora*, *Spicata seu Racemosa*, and *Umbellata*) which were further subdivided into ten sections and additional unnamed groupings including sections *Cirrhopetalum* (Lindl.) Rchb.f., *Brachyantha* Rchb.f., and “Rhachis foliiformis”. Subsequent taxonomic investigations, of which Schlechter’s (1911-1914) was the most influential, proposed various infrageneric classifications. These taxonomic treatments were primarily informed by morphological traits, including inflorescence characteristics, pseudobulb morphology, and floral characteristics such as sepal and petal morphology, to delineate sectional divisions within the genus, resulting in more than 70 sections, most of which were somewhat loosely defined (Holttum, 1957; Schlechter, 1911-1914; Dockrill, 1969; Seidenfaden, 1979; Vermeulen, 2014; Vermeulen et al., 2014).

*Bulbophyllum* has been traditionally placed within subtribe Bulbophyllinae Schltr. within tribe Dendrobieae Endl., alongside several smaller satellite genera such as *Cirrhopetalum*, *Drymoda*, *Pedilochilus* Schltr., *Sunipia* Buch.-Ham. ex Sm., and *Trias* Lindl. (Vermeulen, 1991; Dressler, 1993; Vermeulen, 1993; Cameron et al., 1999; de Witte and Vermeulen, 2010). However, early molecular phylogenetic studies revealed that these satellite genera were nested within *Bulbophyllum*, providing evidence of the non-monophyly of the genus as traditionally circumscribed (Gravendeel et al., 2004; van den Berg et al., 2005; Fischer et al., 2007; Gravendeel et al., 2014; Hosseini et al., 2016). To render the genus *Bulbophyllum* monophyletic, two competing taxonomic approaches were proposed. One favoured maintaining the recognition of the smaller satellite genera alongside a more narrowly defined genus, *Bulbophyllum s.s.,* necessitating the resurrection of genera formerly synonymised with *Bulbophyllum* and the establishment of additional genera, e.g. through elevation of previously established sections to genus level (Garay et al., 1994), an approach advocated mainly by Jones and Clements (2001), Clements and Jones (2002), and Jones (2021). The alternative and subsequently more broadly adopted approach consisted of sinking the previously recognised satellite genera into *Bulbophyllum* to arrive at a more broadly circumscribed generic concept, that of *Bulbophyllum s.l.* (Gravendeel et al., 2014; Vermeulen, 2014; Chase et al., 2015).

The revised taxonomic circumscription of *Bulbophyllum* led to the synonymy of 64 generic names under the genus and the recognition of 97 sections (Pridgeon et al., 2014), some of which have been further validated or renamed in subsequent taxonomic studies (Vermeulen et al., 2014, 2020). Most recently, several sections recognised in earlier classifications but synonymised or merged with others by taxonomists in Pridgeon et al. (2014), have been reinstated such as *Tripudianthes* Seidenf., *Ephippium* Schltr., and *Oxysepala* (Vermeulen et al., 2015; Nguyen et al., 2023), while a new section, *Physometra* J.J.Verm., Suksathan & Watthana, has been erected within *Bulbophyllum* (Vermeulen et al., 2017; Nowak et al., 2023). Informed by phylogenetic studies within Orchidaceae more broadly, the most recent higher-level orchid classification proposed the sinking of subtribe Bulbophyllinae into subtribe Dendrobiinae, together with its sister genus *Dendrobium* Sw., as well as sinking tribe Dendrobieae into Malaxideae (Chase et al., 2015), which is being increasingly adopted.

Recent molecular phylogenetic studies in *Bulbophyllum* support the monophyly of the more broadly defined genus and initially identified four main lineages which revealed a broad-scale biogeographical signal within the genus, termed the Asian, Continental African, Madagascan, and Neotropical clades (Gravendeel et al., 2004; Fischer et al., 2007; Smidt et al., 2011; Gamisch and Comes, 2019). However, the term “Asian” in the aforementioned studies has been used in a broad sense, encompassing taxa distributed across the neighbouring floristic region of Australasia (Australia, New Zealand and the Pacific islands). Gravendeel et al. (2004) conducted a phylogenetic study based on plastid data (*matK*), analysing species of *Bulbophyllum* from all continents. Their findings suggested that non-Asian taxa formed four distinct lineages, with the continental African and Neotropical species constituting a monophyletic group each. Together, these two lineages formed a sister group to the Madagascan clade, and this entire group was sister to the Asian clade. Using nuclear ITS and two to four plastid regions, Fischer et al. (2007) (ca. 150 species, 5 Neotropical) and Smidt et al. (2011) (ca. 50 species, 42 Neotropical) found strongly supported evidence for the monophyly of the Neotropical group, a sister group relationship with the African group, as well as the existence of six Neotropical lineages supported by morphological traits (Smidt et al., 2011). Gamisch and Comes (2019) conducted the first large-scale phylogenetic analysis and divergence-time estimation of *Bulbophyllum* based on ITS sequence data, analysing 320 samples globally. The study identified the Asia-Pacific region as the ancestral area of the genus. While previous studies clarified the monophyly of continental groups particularly the phylogenetic relationships in the Neotropical and Madagascan regions of *Bulbophyllum*, they only offered limited insights into the Asian region which accounts for ca. 70% of *Bulbophyllum* species (Vermeulen, 2014; Gamisch and Comes, 2019).

Within the Asian clade of *Bulbophyllum*, few molecular phylogenetic studies have investigated infrageneric relationships in more detail, with most research focusing on smaller subgeneric groups or specific sectional alliances within this group (Hu et al., 2020; Simpson et al., 2024; Wonnapinij and Sriboonlert, 2015). Wonnapinij and Sriboonlert (2015) using ITS sequences showed that all species of *Bulbophyllum* section *Trias* were embedded within *Bulbophyllum*, supporting the sinking of *Trias* into *Bulbophyllum* (Vermeulen, 2008; Gravendeel et al., 2014). Hu et al. (2020) reconstructed phylogenetic relationships within the Asian *Cirrhopetalum* alliance comprising 15 sections (*Biflorae* Garay, Hamer & Siegerist, *Blepharistes* J.J.Verm., *Brachyantha*, *Cirrhopetaloides* Garay, Hamer & Siegerist, *Cirrhopetalum*, *Desmosanthes* (Blume) J.J.Sm., *Emarginatae* Garay, Hamer & Siegerist, *Ephippium*, *Eublepharon* J.J.Verm., *Lemniscata* Pfitz., *Macrostylidia* Garay, Hamer & Siegerist, *Medusa* Pfitz., *Plumata* J.J.Verm., *Reptantia* J.J.Verm., and *Rhytionanthos* (Garay, Hamer & Siegerist) J.J.Verm.) as well as a broad sampling of eight additional Asian sections of *Bulbophyllum*, using four DNA regions (*nr*ITS, *XDH*, *mat*K, *psb*A-*trn*H) from 95 taxa. Their findings supported the monophyly of a revised *Cirrhopetalum* alliance that also includes section *Desmosanthes*. However, Hu et al. (2020) also showed that many of the previously proposed sections within this alliance were non-monophyletic and lacked clear boundaries, as they had been based largely on overlapping or loosely defined morphological traits. A similar study based on 202 Asian *Bulbophyllum* species and three molecular markers (ITS, *mat*K, and *psb*A-*trn*H) also found most sections within the *Cirrhopetalum* alliance to be non-monophyletic (Thawara et al., 2024).

Zavala-Páez et al. (2020) presented the first comprehensive study of plastid genomes in the genus, using Neotropical species to identify structural characteristics and hotspot regions. Simpson et al. (2024) conducted the first phylogenomic analysis in the genus, with focus on the Asian clade of *Bulbophyllum*. Their study included 70 plastid and nuclear loci across 136 samples, representing 114 species from 41 of the 67 recognised Asian sections accounting for approximately 60% of the Asian sections as recognised by Vermeulen (2014). The study resolved five major lineages, identifying an Australasian lineage as the earliest diverging clade, sister to the remaining four previously recognised lineages the Neotropical, African, Madagascan, and Asian clades. Additionally, the study also revealed the non-monophyly of several taxonomic sections, including *Beccariana* Pfitz., *Brachyantha*, and *Brachypus* Schltr. (Simpson et al., 2024).

Despite considerable efforts to resolve the phylogenetic relationships within *Bulbophyllum*, substantial geographic and taxonomic sampling gaps remain. Half of the sections, for instance *Macrobulbon* Schltr., *Monosepalum*, *Piestobulbon* Schltr., *Uncifera* Schltr., and *Lepanthanthe* Schltr., have never been included in molecular phylogenetic studies. Additionally, most previous studies relied on a limited number of nuclear and plastid markers, often resulting in only low to moderate support in the resulting phylogenetic frameworks (Fischer et al., 2007; Smidt et al., 2011; Hosseini et al., 2012, 2016; Gamisch and Comes, 2019; Thawara et al., 2024). While Simpson et al. (2024) employed multiple plastid protein coding genes to reconstruct phylogenetic relationships within *Bulbophyllum*, ca. 40% of the Asian sections remained unsampled, precluding insights into their phylogenetic placement and monophyly.

Using an expanded dataset that includes the first densely sampled representation of the Asian clade and a multi-locus plastid matrix, this study aims to:

1. generate the most robust phylogenetic framework for *Bulbophyllum* to date,
2. elucidate evolutionary relationships within the genus, with emphasis on evaluating monophyly at the sectional level, and
3. re-assess infrageneric concepts within its largest and most diverse evolutionary lineage, the Asian clade.

## MATERIALS AND METHODS

### Taxon sampling

A total of 355 *Bulbophyllum* specimens were sampled for this study, representing 65 of the 97, or ca. 67% of the sections of *Bulbophyllum* recognised in the most recent classification of the genus (Vermeulen, 2014). Of the 355 specimens, 299 belonged to 51 sections of the Asian clade, representing approximately 76% of the 67 sections recognised for this clade in Vermeulen (2014). In addition, 25 samples representing four Asian sections not recognised in Vermeulen (2014) (*Tripudianthes*, *Macrobulbon*, *Oxysepala*, and *Fruticicola* Schltr.) were also included in the study (Supporting Information, Table S1). The remaining 31 samples comprised species from Madagascar, Africa, Neotropics, and Australasia. Details of sampling coverage of species within sections are provided in Supporting Information, Table S1.

The following 16 Asian sections recognised in Vermeulen (2014) were not included in this study, either because of lack of material or because the material used did not yield DNA of sufficient quality: *Aeschynanthoides* Carr., *Antennata* Pfitz., *Balaenoidea* Pfitz., *Biflorae*, *Biseta* J.J.Verm. ex N.Pearce, *Eublepharon*, *Henosis* (Hook.f.) Ormerod, *Hemisterrantha* J.J.Verm., *Macrocaulia* (Blume) Aver., *Monomeria* (Lindl.) J.J.Verm., *Phreatiopsis* J.J.Verm. & O’Byrne, *Pseudopelma* J.J.Verm. & O’Byrne, *Rhinanthera* J.J.Verm., *Saurocephalum*, *Schistopetalum* Schltr., and *Tapeinoglossum* Schltr. Note that section *Sunipia* sensu Vermeulen (2014) is now called section *Ione* (Lindl.) J.J.Verm. (Vermeulen et al., 2014), section *Recurvae* Garay, Hamer & Siegerist (Vermeulen, 2014) is now section *Ephippium* (Vermeulen et al., 2015), and section *Hemisterrantha* has not been validly published.

Additional DNA sequence data for 19 *Bulbophyllum* species and 94 outgroup taxa were included based on previous phylogenetic studies on orchids (Givnish et al., 2015; Kim et al., 2020; Zavala-Páez et al., 2020; Serna-Sánchez et al., 2021; Li et al., 2023; Pérez-Escobar et al., 2024; Goedderz et al., 2024; Simpson et al., 2024; Wu et al., 2024), obtained from GenBank (https://www.ncbi.nlm.nih.gov/genbank/). Taxonomic concepts applied were based on the accepted taxonomy in International Plant Names Index (https://www.ipni.org/) and Plants of the World Online (https://powo.science.kew.org/), while sectional taxonomy followed IOSPE (Internet Orchid Species Photo Encyclopedia [http://www.orchidspecies.com/]), which provides a baseline classification distilled from recently published works. Details on plant material and voucher information for each sample are provided in Supporting Information, Table S2.

### DNA extraction, amplification, and sequencing

Genomic DNA was extracted from approximately 5 to 20 mg of silica-dried or fresh leaf material using commercial DNA extraction kits (Qiagen DNeasy plant kit, Venlo, Netherlands) and ChargeSwitch gDNA plant kit (Invitrogen, Carlsbad, USA), following the manufacturers’ instructions, or using the CTAB protocol (Doyle, 1991).

Library preparation was either performed at the Australian Genome Research Facility (AGRF) or the National Research Collections Australia (NRCA), CSIRO. At the AGRF, libraries for high-throughput sequencing were constructed using between 5 to 100 ng total DNA using the NEB Next Ultra II DNA library preparation kit (New England Biolabs, Ipswich, MA, USA), with an insert size of 350 base pairs (bp) following the manufacturer’s protocol. Libraries were indexed with uniquely dual-indexed primers and amplified in pools of 16 libraries using 12-15 PCR cycles. Fragment size selection was carried out using AMPure XP beads (Beckman Coulter, Beverly, USA) following the manufacturer’s instructions. DNA concentration of each library was quantified using Qubit fluorometer HD dsDNA kit (Invitrogen, California, USA) following the manufacturer’s protocol and fragment lengths were determined using the Fragment Analyser NGS kit (Agilent, California, USA). Sequencing libraries were then multiplexed 96 times and DNA sequencing with 125 or 150 bp paired-end reads performed on Illumina high-throughput sequencing NovaSeq 600 SP platform at the Australian Genomic Research Facility (Melbourne, Australia).

At NRCA (CSIRO), DNA was sonicated to an average target length of approximately 200 bp using a Covaris LE220 ultra-sonicator (Woburn, Massachusetts, USA) and libraries built using the QIAseq UltraLow Input Library kit (Qiagen, Clayton, Australia) using custom dual-indexed adapters. Final libraries were size-assessed on a Fragment Analyser using the high-sensitivity NGS Fragment Kit, quantified using a fluorescence method and pooled equimolarly. Samples were sequenced on a NovaSeq S1 flowcell (Illumina, California, USA) with paired-end 2 x 150 cycle chemistry at the Biomolecular Resource Facility within the John Curtin School of Medical Research, Australian National University (Canberra, Australia). The raw reads were deposited in the European Nucleotide Archive (ENA) database.

### Sequence assembly, annotation, and alignment

Sequences were assembled and edited using Geneious Prime v.2023.2 (https://www.geneious.com). Illumina sequences were assembled to a reference set of plastid coding regions extracted from the reference genome of *Bulbophyllum inconspicuum* Maxim. (Genbank accession number NC_046811) downloaded from GenBank. Assembly was conducted with the highest quality threshold and a minimum coverage of ten reads. The assemblies were checked for quality and edited manually where necessary. Coding regions of the respective genes from 374 *Bulbophyllum* and 94 outgroup samples were then extracted using the ’extract’ function in Geneious. Sequences affected by mutations leading to frame shifts with internal stop codons were excluded from phylogenetic analyses (Supporting Information, Table S3). Each extracted coding region, totalling 63 plastid protein coding loci from 468 samples were aligned using the Geneious Prime plugin MAFFT v.1.5.0 (Katoh et al., 2002; Katoh and Standley, 2013) with default settings. The resulting gene alignments were manually inspected and then concatenated into a plastid dataset comprising 53,578 bp for phylogenetic analysis. Variable sites were checked in Geneious Prime v.2023.2 and included 2% of the concatenated dataset (53,578 bp). Missing data comprised ca. 5.6% of the concatenated matrix (Supporting Information, Table S3).

### Phylogenetic analyses

Phylogenetic analyses were conducted on the concatenated plastid dataset (63 plastid protein coding genes: 468 samples) using the maximum likelihood approach within the IQ-TREE2 v.2 package (Minh et al., 2020) where the concatenated matrix was treated as a single partition and the optimal model was selected automatically by the program. The best fit model was identified as GTR+F+R6 using IQ-TREE2’s ModelFinder (Kalyaanamoorthy et al., 2017), based on the Akaike Information Criterion (AIC) (Akaike, 1974). Nodal support was evaluated through running 1000 replicates to obtain the Shimodaira-Hasegawa-like approximate Likelihood Ratio Test (SH-aLRT) and 1000 ultrafast bootstraps using the ‘-bb’ option (Guindon et al., 2010; Minh et al., 2013; Hoang et al., 2018; Minh et al., 2020). The combination of SH-aLRT alongside ultrafast bootstrap support provides more reliable assessment of nodal support than either measure alone (Minh et al., 2013). Minh et al. (2013) and Hoang et al. (2018) demonstrated that SH-aLRT values ≥80 generally correspond well to bootstrap values ≥95, and the use of both measures in combination reduces the risk of both false positives and false negatives in support assessment. Nodes were only considered strongly supported when both SH-aLRT and UFBoot values met their respective thresholds, providing a conservative and robust criterion for evaluating phylogenetic support across the tree. In the present study, nodal support values were categorized as follows: nodes with both SH-aLRT and UFBS values ≥95 were considered strongly/well supported, ≥ 90-94 were moderately supported, ≥ 80-89 were weakly supported, while values <80 were treated as unsupported. In cases where only one measure (either SH-aLRT or UFBS) was high while the other indicated lack of support, the node was classified as unsupported. All resulting trees were rooted to relevant outgroups and visually displayed using Figtree v.1.4.4 (http://tree.bio.ed.ac.uk/software/figtree/).

## RESULTS

### Phylogenetic relationships within *Bulbophyllum*

The maximum likelihood phylogenetic tree inferred from the 63 plastid loci concatenated alignment provides robust support for the monophyly of *Bulbophyllum* and its sister group relationship with *Dendrobium* (SH-aLRT/UFboot 99.9/100) (Fig. 2; Supporting Information, Fig. S1). Within *Bulbophyllum*, five major lineages corresponding to the broader biogeographic regions Australasia, Madagascar, Africa, Neotropics and Asia, are retrieved with moderate to strong bootstrap support (Fig. 2). The complete phylogram (outgroups pruned except for sister group *Dendrobium*) is provided in Supporting Information, Fig. S2. The first diverging lineage, the Australasian clade (Clade A: 92.5/97), harbours section *Adelopetalum* and Australasian species of section *Minutissima* Pfitz. Neither of these sections is retrieved as monophyletic (Fig. 3).

**Figure 2.**
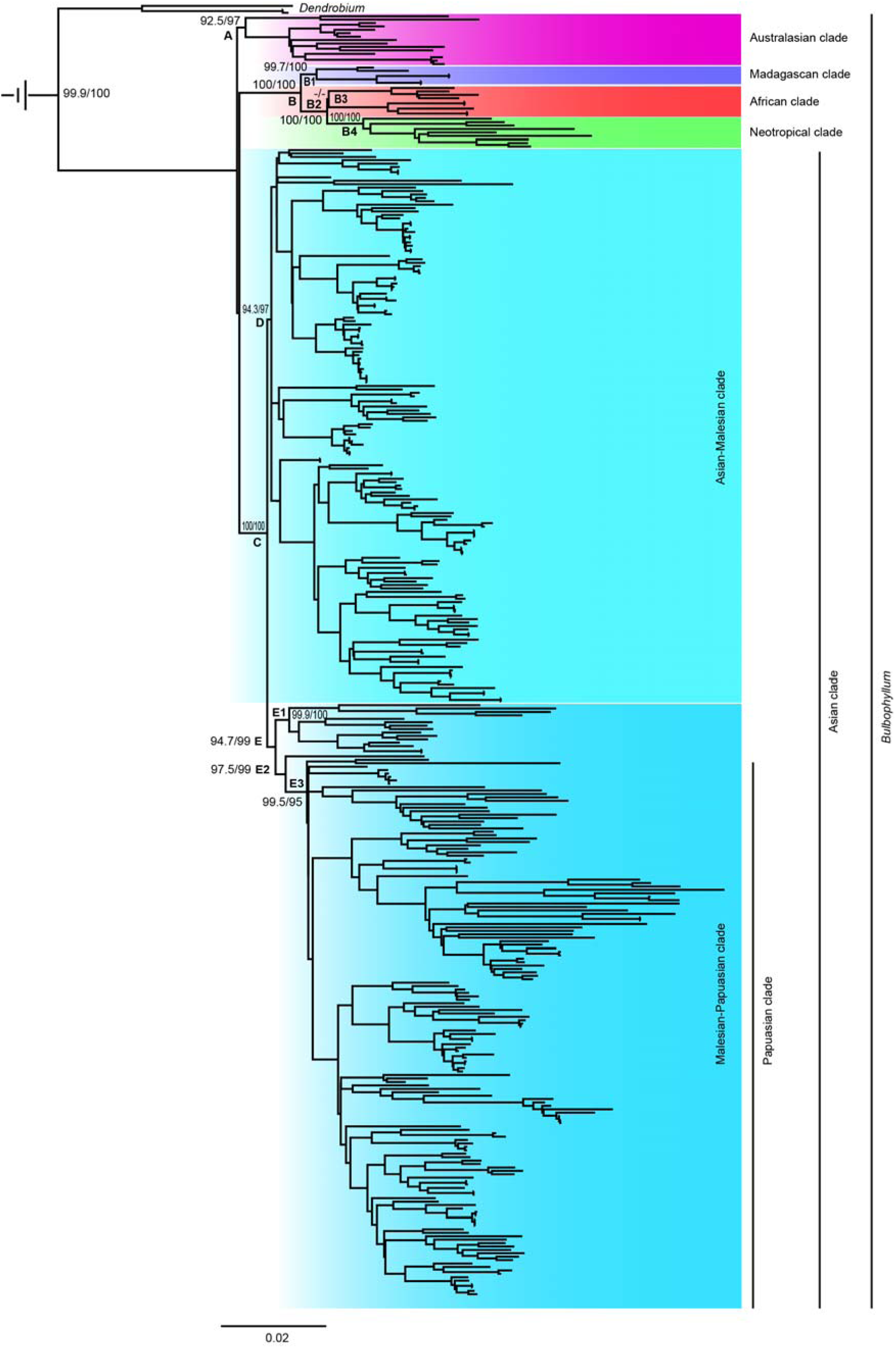
Maximum likelihood phylogenetic tree showing the relationships between *Bulbophyllum* and its sister group *Dendrobium* and the relationships among major biogeographic lineages within *Bulbophyllum*. The tree is based on 468 taxa (other outgroups not depicted) and was constructed using 63 plastid loci. Support values are displayed above each branch, with SH-aLRT followed by UFBoot values. Values <80 are indicated by “-/-”. The scale bar represents branch length, corresponding to 0.02 expected substitutions per site. The log-likelihood value of the best-scoring maximum likelihood tree was -480636.010. Outgroups outside Dendrobieae not depicted. **Alt text.** A maximum-likelihood phylogenetic tree showing the placement of *Bulbophyllum* relative to its sister group *Dendrobium*, with outgroups outside Dendrobieae excluded from view. The tree is colour coded to depict the major biogeographic lineages within *Bulbophyllum*, namely the Australasian, Madagascan, African, Neotropical, and Asian clades. The Papuasian clade is nested within the Malesian-Papuasian lineage of the broader Asian clade.

**Figure 3:**
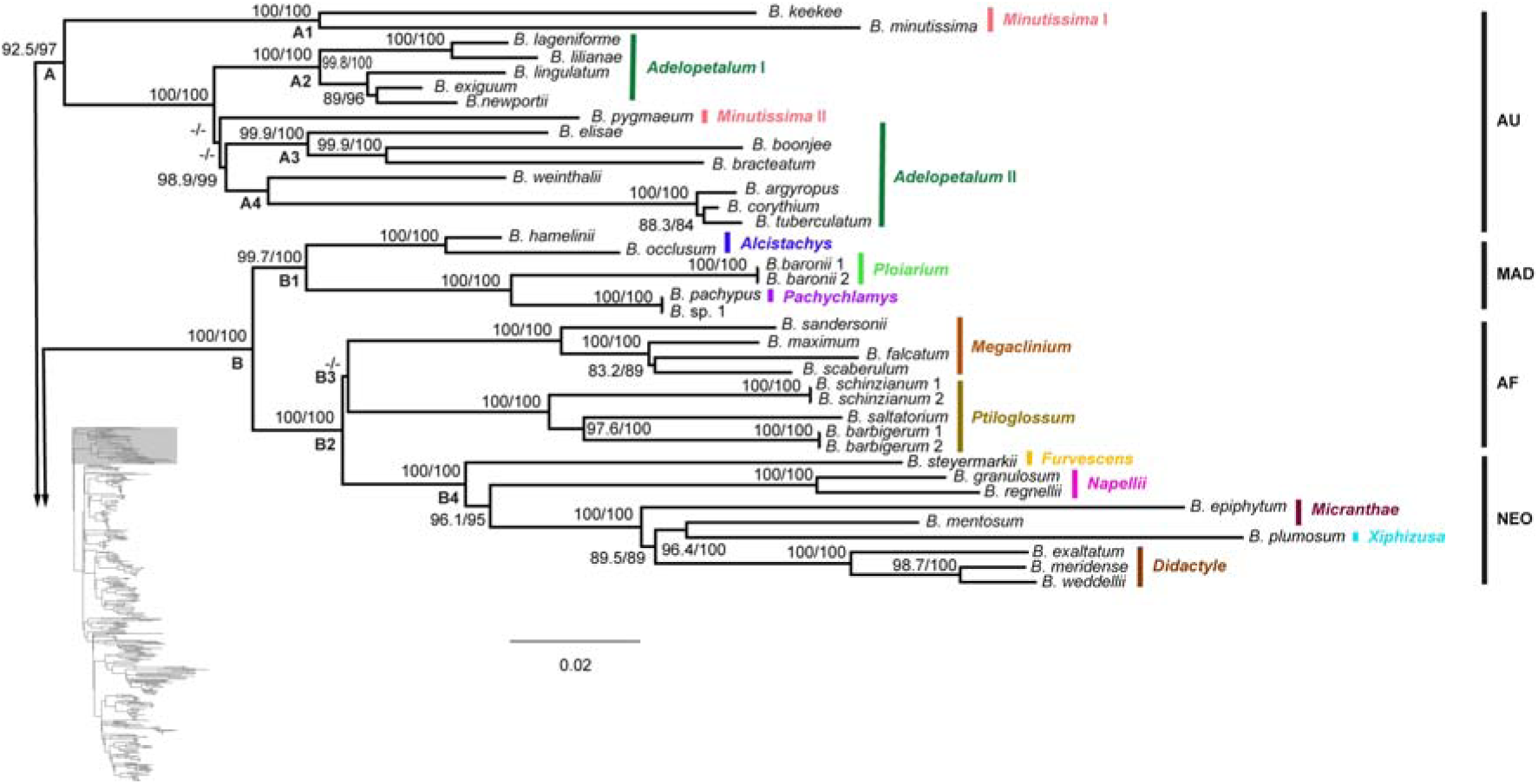
Phylogenetic relationships within the Australasian (A: AU), Madagascan (B1: MAD), African (B3: AF) and Neotropical (B4: NEO) lineages of *Bulbophyllum*. The maximum likelihood tree is based on 63 plastid loci from 468 taxa (outgroups not shown). Support values are displayed above each branch, with SH-aLRT followed by UFBoot values. Values <80 are indicated by “-/-”. The scale bar represents branch length, corresponding to 0.02 expected substitutions per site. Sections are indicated in bold after the species names. **Alt text.** A maximum-likelihood phylogenetic tree showing relationships within four biogeographic clades of *Bulbophyllum*: Australasian (AU), Madagascan (MAD), African (AF), and Neotropical (NEO). Species names are followed by their corresponding section in bold.

Next diverging is Clade B (100/100), in which the Madagascan clade (B1) diverges first, followed by the African (B3) and the Neotropical clade (B4) (Fig. 2 and Fig. 3). In sister group position to Clade B, the Asian Clade (C) is retrieved with maximum support (Fig. 2: 100/100).

The Asian clade is divided into two major groups: the Asian-Malesian clade, comprising species primarily from the Asian and Malesian regions (subclade D: 94.3/97), and the Malesian-Papuasian clade, comprising species mainly from Malesia and the Papuasian region (subclade E: 94.7/99) (Fig. 2). It should be noted that species distributions do not align perfectly with these biogeographic boundaries. Not all species in the Malesian-Papuasian clade occur in Malesia and Papuasia, and similarly, many species of the Asian-Malesian clade are found in Papuasia (Vermeulen, 2014; Vermeulen et al., 2015).

Within the Asian-Malesian lineage (D), two larger main clades are retrieved, labelled F and G (Fig. 4 and Fig. 5 respectively; Figs S1 and S2). Among the 12 sections placed within clade F (*Altisceptrum* J.J.Sm., *Beccariana*, *Blepharistes*, *Drymoda*, *Hymenobractea* Schltr., *Intervallatae* Ridl., *Lemniscata*, *Lepidorhiza*, *Reptantia*, *Sestochilus* Benth. & Hook.f., *Stenochilus* J.J.Sm., *Trias*), evidence of non-monophyly is found for (1) section *Lepidorhiza*, with section *Hymenobractea* and section *Intervallatae* nested within, and (2) section *Lemniscata*, whose members are found within clade F and G. The monophyly of section *Sestochilus* and section *Stenochilus* is well supported, as well as their sister group relationship. Other well-supported sections within clade F are section *Beccariana* and section *Trias*.

**Figure 4.**
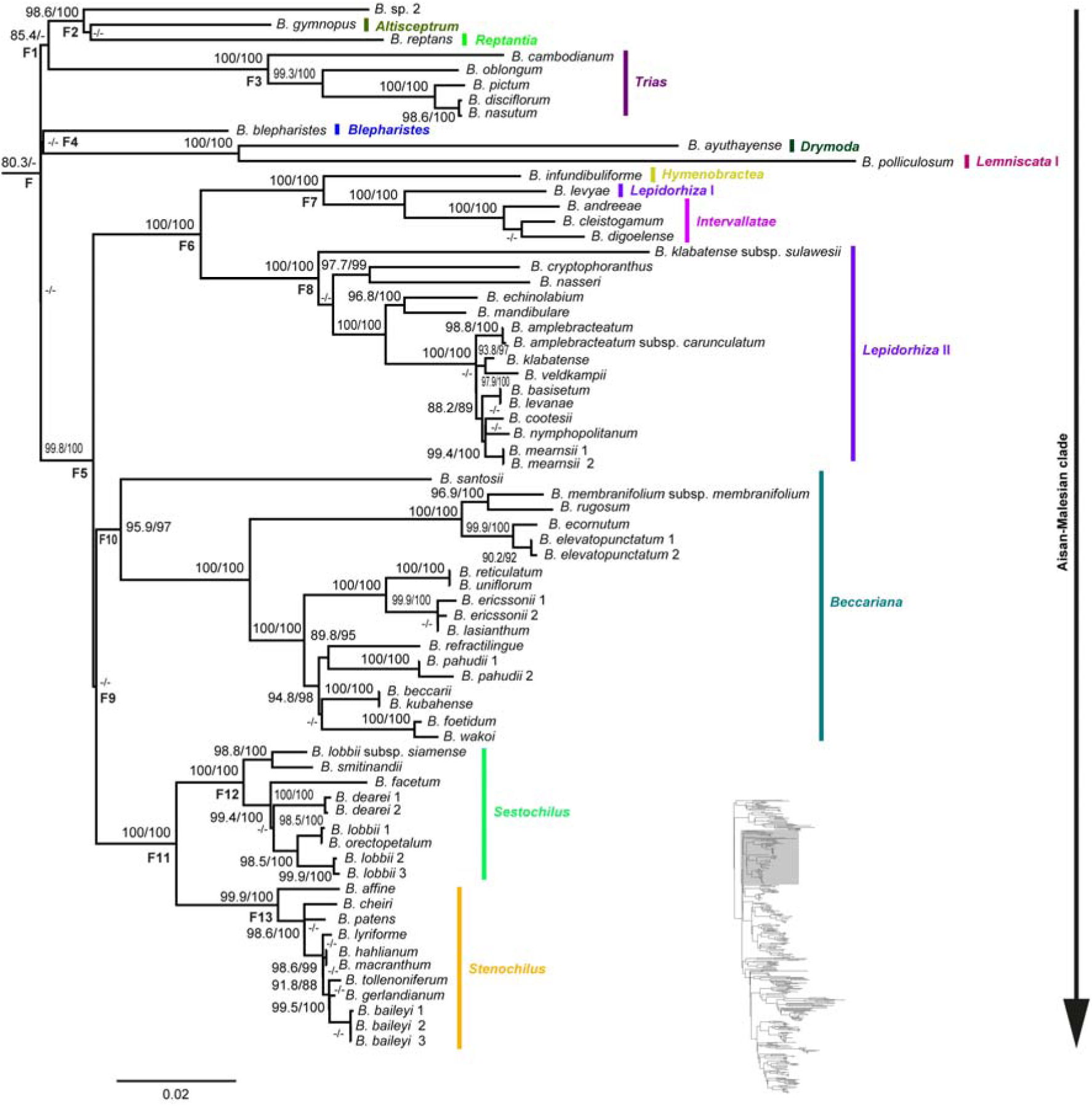
Phylogenetic relationships within the first major lineage (F) of the Asian-Malesian clade of *Bulbophyllum*. The maximum likelihood tree is based on 63 plastid loci from 468 taxa (outgroups not shown). Support values are displayed above each branch, with SH-aLRT followed by UFBoot values. Values <80 are indicated by “-/-”. The scale bar represents branch length, corresponding to 0.02 expected substitutions per site. Smaller subclades representing sectional groupings are labelled F1-F13. Section names for species are indicated in bold after the species names. **Alt text.** A maximum-likelihood phylogenetic tree showing relationships within the first major lineage (F) of the Asian-Malesian clade of *Bulbophyllum*. Smaller subclades are labelled F1-F13, representing sectional groupings, and species names are followed by their corresponding section names in bold.

**Figure 5.**
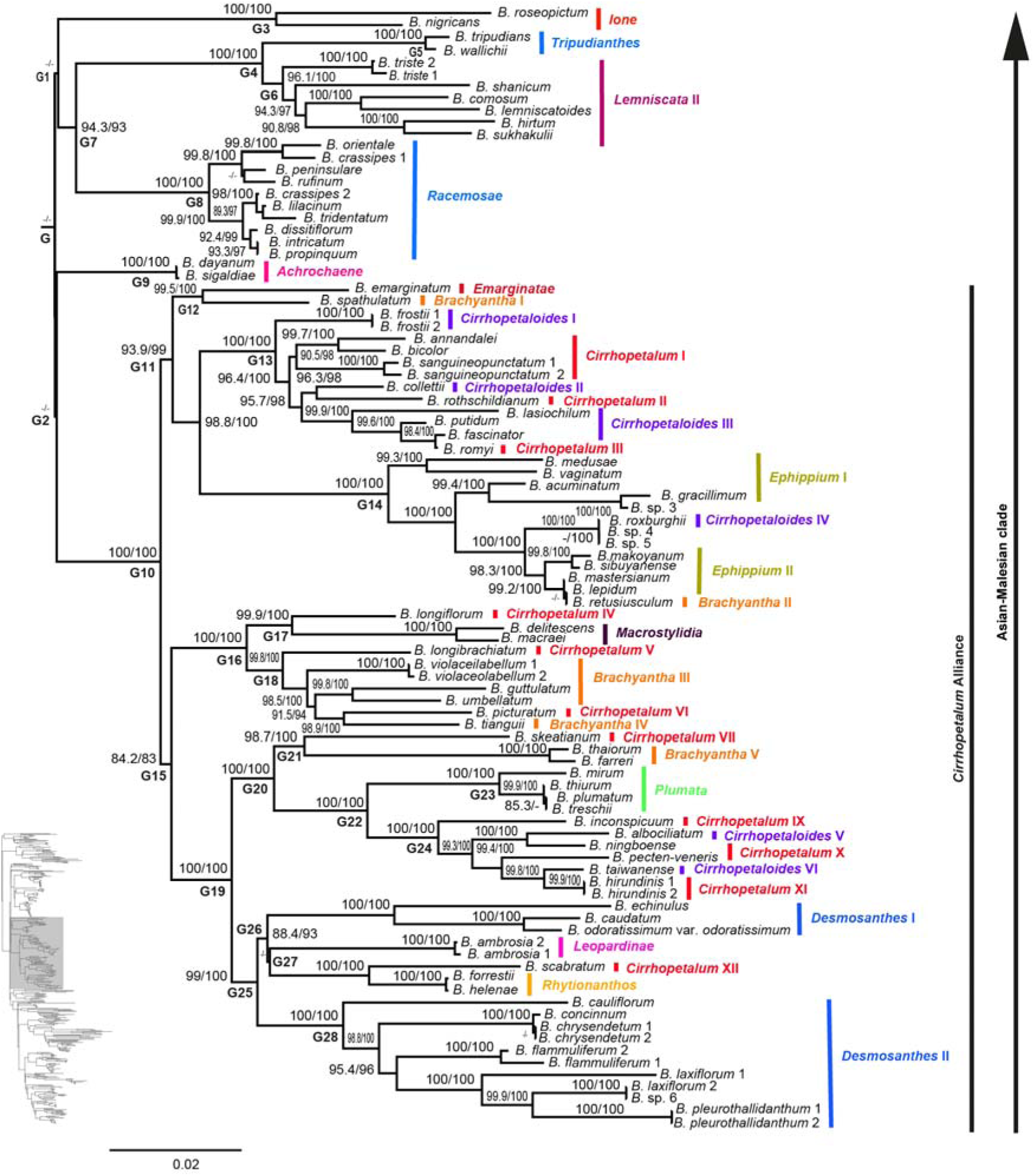
Phylogenetic relationships within the second major lineage (G) of the Asian-Malesian clade of *Bulbophyllum*. The maximum likelihood tree is based on 63 plastid loci from 468 taxa (outgroups not shown). Support values are displayed above each branch, with SH-aLRT followed by UFBoot values. Values <80 are indicated by “-/-”. The scale bar represents branch length, corresponding to 0.02 expected substitutions per site. Smaller subclades corresponding to sectional groupings are labelled G1-G28. The *Cirrhopetalum* alliance is designated as G10. Section names for species are indicated in bold after the species names. **Alt text.** A maximum-likelihood phylogeny illustrating relationships within the second major lineage (G) of the Asian-Malesian clade of *Bulbophyllum*. Subclades G1-G28 are labelled, including the *Cirrhopetalum* alliance designated as G10. Species names are followed by their corresponding section names in bold.

The second main clade within the Asian-Malesian lineage, clade G, largely comprises the *Cirrhopetalum* alliance. Subclade G1 comprises three well-supported smaller clades representing monophyletic sections *Ione*, *Tripudianthes*, and *Racemosae* Benth. & Hook.f., as well as other representatives of the non-monophyletic section *Lemniscata*.

Subclade G2 encompasses the core *Cirrhopetalum* Alliance Group (CAG) (G10: 100/100) and one satellite clade, namely section *Achrochaene* (G9: 100/100). Six major subclades, namely G12 (99.5/100), G13 (100/100), G14 (100/100), G16 (100/100), G20 (100/100), and G25 (99/100), are recognised within the core *Cirrhopetalum* alliance, each with strong nodal support (Fig. 5). These subclades comprise species from sections *Brachyantha*, *Emarginatae*, *Cirrhopetaloides*, *Cirrhopetalum*, *Ephippium*, *Macrostylidia*, *Plumata*, *Rhytionanthos*, *Leopardinae* Benth. & Hook.f., and *Desmosanthes*, all of which are non-monophyletic.

Within the *Cirrhopetalum* alliance, G12 includes the non-monophyletic sections *Emarginatae* and *Brachyantha*, supported together on the same node. Species from sections *Cirrhopetalum* and *Cirrhopetaloides* are intermixed within subclade G13. G14 consists mainly of species from section *Ephippium*, with single species of sections *Brachyantha* and *Cirrhopetaloides* nested within it. Section *Macrostylidia* is recovered within G16, embedded among species of sections *Cirrhopetalum* and *Brachyantha*. Section *Plumata* is recovered within G20, sister to G24, which comprises intermixed species from sections *Cirrhopetalum* and *Cirrhopetaloides*. Subclade G25 consists mainly of the non-monophyletic section *Desmosanthes*, with species of sections *Cirrhopetalum*, *Leopardinae*, and *Rhytionanthos* nested within it.

The second arm of the Asian clade comprises the Malesian-Papuasian clade which is composed of three well supported lineages namely E1 (99.9/100), E2 (97.5/99) (Fig. 6) and the Papuasian clade (E3: 99.5/95). Subclade E1 is composed of species from monophyletic section *Brachystachyae* Benth. & Hook.f., which forms a close relationship with sections *Hirtula* and *Stachysanthes* (Blume) Aver., both of which are monophyletic sister groups. *Bulbophyllum planibulbe* Ridl., section *Planibulbus* J.J.Verm., is recovered as an early-diverging lineage within E2.

**Figure 6.**
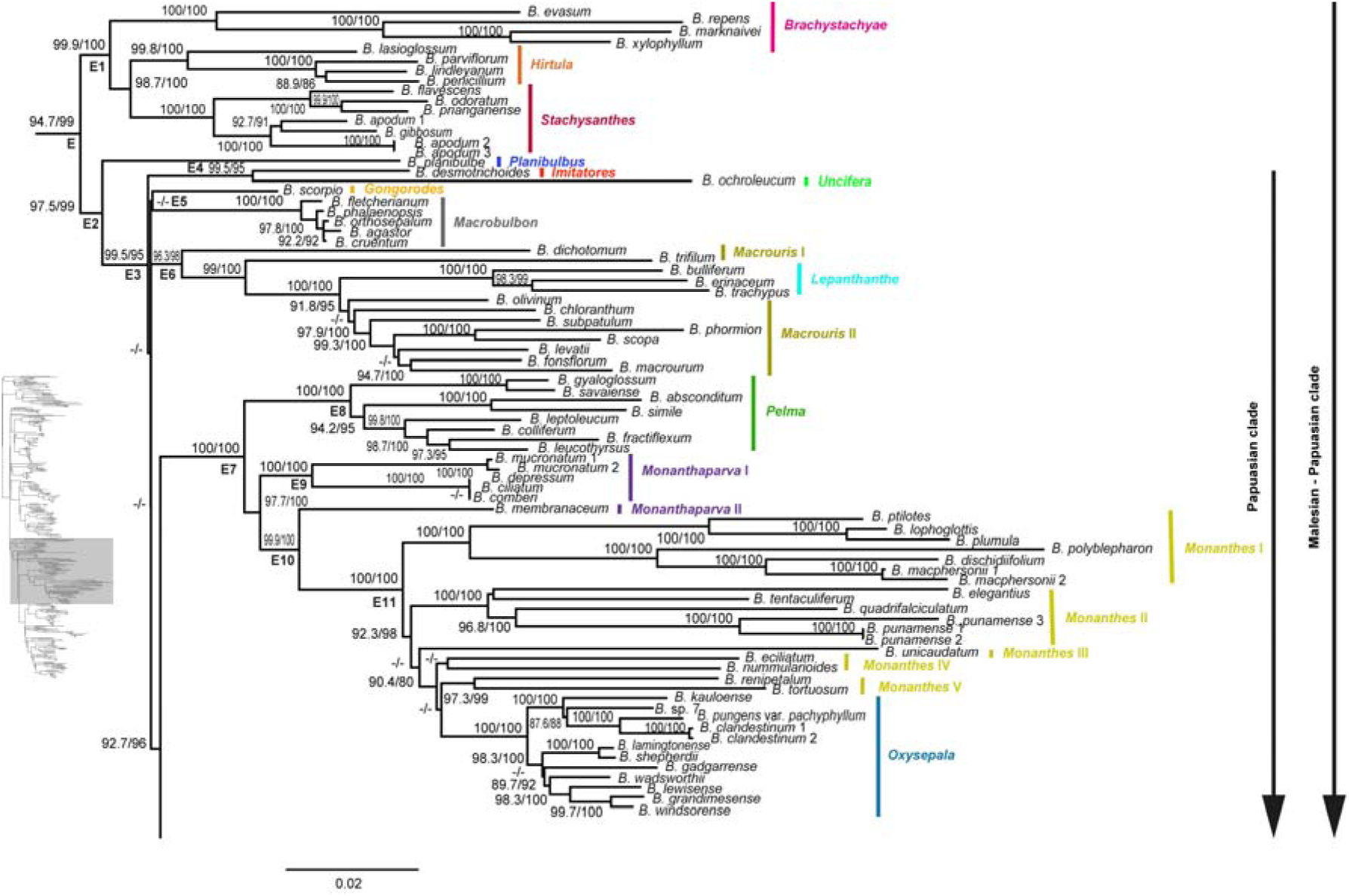
Phylogenetic relationships within the first half of the Malesian-Papuasian clade (E) of *Bulbophyllum*. The maximum likelihood tree is based on 63 plastid loci from 468 taxa (outgroups not shown). Support values are displayed above each branch, with SH-aLRT followed by UFBoot values. Values <80 are indicated by “-/-”. The scale bar represents branch length corresponding to 0.02 expected substitutions per site. Smaller subclades representing sectional groupings are labelled E1-E10. Section names for species are indicated in bold after the species names. **Alt text.** A maximum-likelihood phylogeny depicting relationships within the first half of the Malesian-Papuasian clade of *Bulbophyllum*. Smaller subclades (E1-E10) are labelled to indicate sectional groupings, and species names are followed by their corresponding section names in bold.

The Papuasian clade (E3: 99.5/95) is a well-supported lineage within the Malesian-Papuasian clade and comprises of six subclades E4-E7 (Fig. 6) and E13 and E18 (Fig. 7). *Bulbophyllum ochroleucum* Schltr. section *Uncifera* and *Bulbophyllum desmotrichoides* Schltr. section *Imitatores* J.J.Verm. are supported on the same node with strong support (E4: 99.5/95). The unsupported subclade E5 consists of *Bulbophyllum scorpio* J.J.Verm. of section *Gongorodes* J.J.Sm. and species of the monophyletic section *Macrobulbon* (100/100).

**Figure 7.**
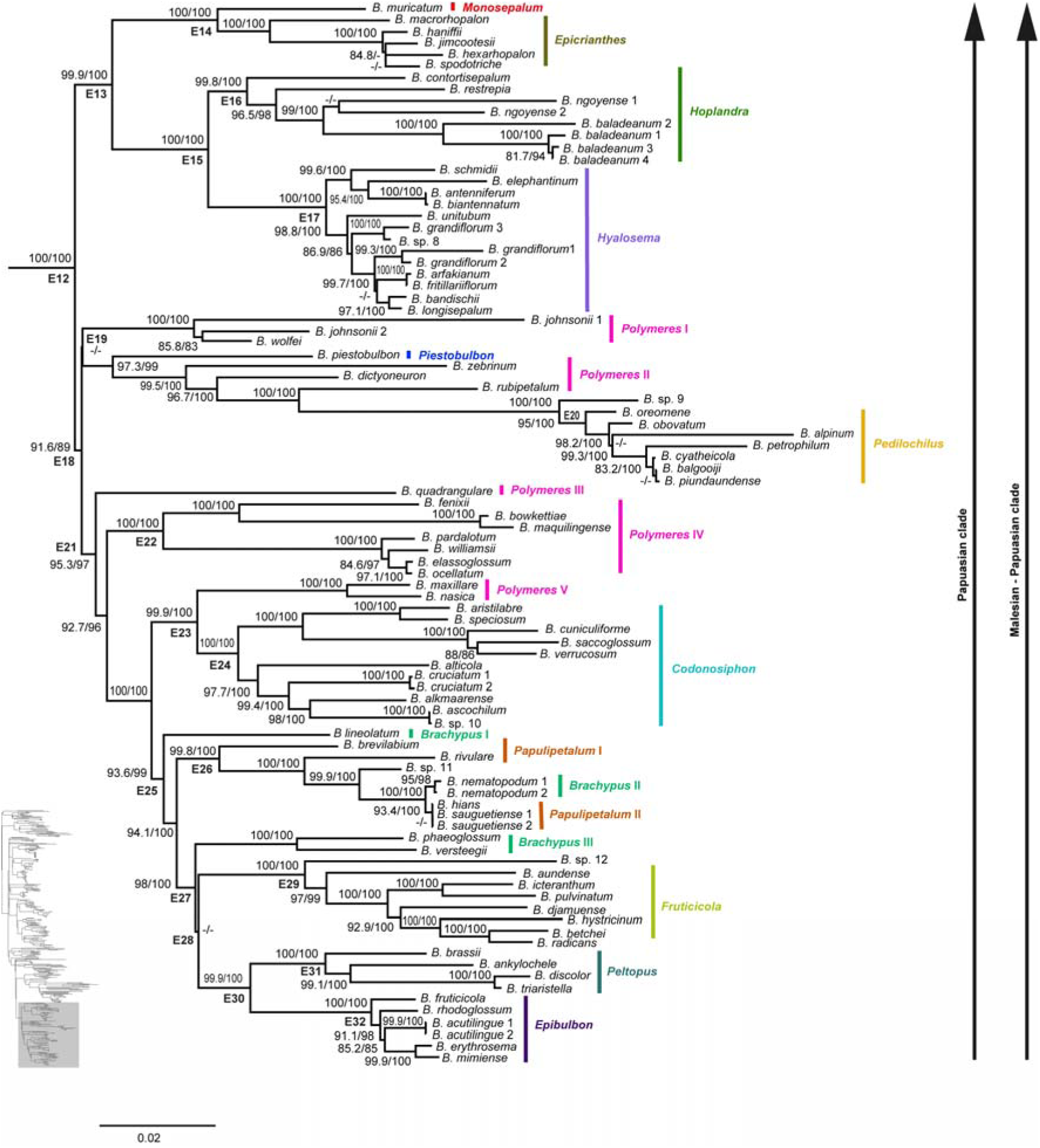
Phylogenetic relationships within the apex of the Malesian-Papuasian clade of *Bulbophyllum*. The maximum likelihood tree is based on 63 plastid loci from 468 taxa (outgroups not shown). Support values are displayed above each branch, with SH-aLRT followed by UFBoot values. Values <80 are indicated by “-/-”. The scale bar represents branch length, corresponding to 0.02 expected substitutions per site. Smaller subclades representing sectional groupings are labelled E11-E32. Section names for species are indicated in bold after the species names. **Alt text.** A maximum-likelihood phylogeny representing the apex of the Malesian-Papuasian clade of *Bulbophyllum*. Subclades E11-E32 indicate sectional groupings, and species labels are followed by their corresponding section names in bold.

Subclade E6 (96.3/98) includes mainly species of the non-monophyletic section *Macrouris* Schltr., and *Lepanthanthe*. The subsequent, well-supported subclade E7 (100/100) encompasses the early diverging section monophyletic *Pelma* Schltr., followed by the non-monophyletic sections *Monanthaparva* Ridl. and *Monanthes* (Blume) Aver. with the latter forming a grade having species of section *Oxysepala* nested within it.

Subclade E13 (99.9/100) comprises sections *Monosepalum*, *Epicrianthes*, *Hoplandra* J.J.Verm., and *Hyalosema* Schltr. (Fig. 7). Section *Monosepalum* is closely related to species of the monophyletic section *Epicrianthes*. The monophyly for section *Hoplandra* (E16) and section *Hyalosema* (E17) is well supported, as well as their sister group relationship.

Subclade E18 is poorly resolved and includes species from sections *Polymeres*, *Piestobulbon*, *Pedilochilus*, *Codonosiphon*, *Brachypus*, *Papulipetalum* Schltr., *Fruticicola*, *Peltopus* Schltr., and *Epibulbon* Schltr. (Fig. 7). Within E18, the unsupported subclade E19 comprises species of the polyphyletic section *Polymeres*, with *Bulbophyllum piestobulbon* Schltr. (section *Piestobulbon*) nested among three species of section *Polymeres* (*Bulbophyllum zebrinum* J.J.Sm., *Bulbophyllum dictyoneuron* Schltr., and *Bulbophyllum rubipetalum* P.Royen), together with species of section *Pedilochilus* (E20: 95/100).

Subclade E21 (95.3/97) includes the remaining species of the polyphyletic section *Polymeres*, with two species from this section (*Bulbophyllum maxillare* (Lindl.) Rchb.f. and *Bulbophyllum nasica* Schltr.) closely related to the monophyletic section *Codonosiphon* on subclade E23 (99.9/100).

Subclade E25 consists of an early-diverging *Bulbophyllum lineolatum* Schltr., section *Brachypus*, and two well-supported smaller clades, E26 (99.8/100) and E27 (98/100). Within E26, species of section *Papulipetalum* are intermixed with species of section *Brachypus*, rendering both sections non-monophyletic. The final subclade, E27, comprises an early-diverging lineage containing the remaining two species of the non-monophyletic section *Brachypus* (*Bulbophyllum phaeglossum* Schltr. and *Bulbophyllum versteegii* J.J.Sm.) and an unsupported subclade E28, which encompasses sections *Fruticicola* (E29: 97/99), *Peltopus* (E31: 100/100), and *Epibulbon* (E32: 100/100), all recovered as monophyletic, with the latter two recovered as successive sister groups with strong nodal support (E30: 99.9/100).

## DISCUSSION

### Monophyly of *Bulbophyllum* and its biogeographic lineages

Our phylogenomic study of *Bulbophyllum* based on 63 plastid protein-coding genes (Figs. 2-7; Supporting Information, Figs S1 and S2), establishes a broad phylogenetic framework for the genus and provides the most comprehensive understanding of the Asian lineage to date. Our results provide robust support for the monophyly of the genus and its sister relationship with *Dendrobium*, consistent with previous phylogenetic findings (Simpson et al., 2024; Pérez-Escobar et al., 2024; Thawara et al., 2024). The overall topology of the maximum likelihood phylogenetic tree exhibits a broad-level biogeographic structure with major continental lineages forming well-supported monophyletic clades, consistent with Simpson et al. (2024). The Asian clade of *Bulbophyllum* is densely sampled for the first time and is well supported (Fig. 2: Clade C), although many sections within this clade remain unresolved due to lack of support at internal nodes, highlighting the need for taxonomic revision within the genus.

Five distinct biogeographical lineages are inferred within *Bulbophyllum*, with the Australasian clade recovered as an early-diverging lineage sister to the rest of the genus (Simpson et al., 2024; Fig. 2: Clade A). The Madagascan clade is recovered as sister to a clade comprising both the African and Neotropical clades, which form a sister relationship (Gamisch and Comes, 2019; Smidt et al., 2011; Simpson et al., 2024; Fig. 2: Clade B). The final strongly supported lineage corresponds to the Asian clade of *Bulbophyllum* (Simpson et al., 2024; Fig. 2: Clade C).

While the five main biogeographic clades within *Bulbophyllum* are clearly resolved, lineages within the Asian region exhibit more complex biogeographic structure. Our phylogenetic analysis reveals a dichotomous split within the Asian clade into two biogeographic lineages: an Asian-Malesian clade (Fig. 2: Clade D) and a Malesian-Papuasian clade (Fig. 2: Clade E), supporting a biogeographic distinction identified among subclades within the Asian lineage in early phylogenetic analyses (Gravendeel et al., 2014). The Asian-Malesian clade comprises predominantly of sections found in Southeast Asia and Eastern Asia with some species from Australasia, Malesia, Papuasia, and the Pacific (Vermeulen, 2014; Figs. 4 & 5). Within the Malesian-Papuasian clade (Fig. 6), our analysis reveals a grade of West Malesian lineages terminating with a Papuasian clade comprised of lineages predominantly occurring in Papuasia. The pattern is consistent with the intricate biogeographical patterns that characterise the Malesian region, a complex transitional zone where Asian floristic elements in west Malesia grade eastward through Wallacea into the Australasian-influenced flora of east Malesia, a gradient shaped by the region’s dynamic geological history and position at the interface of major continental plates (van Steenis-Kruseman and van Steenis, 1950; Kalkman, 1955; De Boer et al., 2015). Further studies, particularly historical biogeography using ancestral area reconstruction, are needed to fully clarify the origin, diversification dynamics, and dispersal patterns of the genus specifically within the Asian clade.

### The Australasian clade

The Australasian clade comprises species primarily from Australia, New Zealand, and the Pacific, with sections *Adelopetalum* and *Minutissima* forming a sister relationship (Fig. 3: Clade A), consistent with the findings of Simpson et al. (2024). Furthermore, the *Bulbophyllum argyropus* (Endl.) Rchb.f., *Bulbophyllum bracteatum* (Fitzg.) F.M.Bailey, and *Bulbophyllum newportii* (F.M.Bailey) Rolfe clades identified by Simpson et al. (2024) within section *Adelopetalum* are recovered with strong statistical support. However, the phylogenetic placement of *Bulbophyllum pygmaeum* (Sm.) Lindl. within section *Adelopetalum* remains unresolved.

Section *Minutissima* sensu Vermeulen (2014) and Vermeulen et al. (2015) includes species previously assigned by Schlechter to section *Nematorhizis* Schltr., which are mainly found in New Guinea, alongside several Malesian and continental Asian species. Of these, we only sampled *Bulbophyllum mucronatum* (Blume) Lindl., which was recovered nesting with species of section *Monanthaparva*, far removed from the type species of section *Minutissima*. A narrower circumscription of section *Minutissima* may therefore be warranted, restricted to species from Australia and New Caledonia (Jones and Clements, 2001). Additionally, the disposition of species previously included in sect. *Nematorhizis* requires more research.

### Madagascan, African, and Neotropical clades

Within the Madagascan clade (Fig. 3: Clade B1), section *Alcistachys* Schltr. forms an early-diverging lineage, sister to a strongly supported clade that includes sections *Pachychlamys* Schltr. and *Ploiarium* Schltr. Despite limited species sampling in the present study, relationships among these sections are consistent with previous phylogenetic findings (Gamisch and Comes, 2019) (Fig. 3). Within the African clade (Fig. 3: Clade B3), sections *Megaclinium* G.A. Fischer & J.J.Verm. and *Ptiloglossum* Lindl. are recovered at the same node, although their relationship remains uncertain due to a lack of support at the corresponding internal node.

The Neotropical clade (Fig. 3: Clade B4) comprises the early-diverging *Bulbophyllum steyermarkii* Foldats (section *Furvescens* E.C. Smidt, Borba and van den Berg), which is sister to the remaining Neotropical taxa (Smidt et al., 2011; Gamisch and Comes, 2019; Zavala-Páez et al., 2020). Consistent with Smidt et al. (2011) and Zavala-Páez et al. (2020), section *Napellii* Rchb.f. forms an early-diverging lineage that is closely related to a strongly supported subclade comprising sections *Micranthae* Barb.Rodr., *Xiphizusa* Rchb.f., and *Didactyle* (Lindl.) Cogn. with high support and morphological congruence.

### The Asian clade

Within the Asian clade (Figs. 4-7), 20 sections are resolved as monophyletic: *Achrochaene*, *Beccariana*, *Brachystachyae*, *Hirtula*, *Ione*, *Racemosae*, *Sestochilus*, *Stachysanthes*, *Stenochilus*, *Trias*, *Tripudianthes*, *Codonosiphon*, *Epibulbon*, *Epicrianthes*, *Fruticicola*, *Hoplandra*, *Hyalosema*, *Macrobulbon*, *Pelma*, and *Peltopus*, which are all supported by strong statistical support based on representative sampling. Several sections, however, are identified as non-monophyletic, including *Drymoda*, *Hymenobractea*, *Intervallatae*, *Lemniscata*, *Lepidorhiza*, all 10 sections of the *Cirrhopetalum* alliance (Asian-Malesian clade), as well as *Brachypus*, *Macrouris*, *Monanthaparva*, *Monanthes*, *Papulipetalum*, and *Polymeres* (Malesian-Papuasian clade). This study also provides the first molecular evidence clarifying the phylogenetic positions of several previously unsampled or poorly known sections, including *Monosepalum*, *Macrobulbon*, *Gongorodes*, *Imitatores*, *Lepanthanthe*, *Piestobulbon*, *Planibulbus*, and *Uncifera*. Nonetheless, the placement of several sections such as *Altisceptrum*, *Blepharistes*, and *Reptantia* remains unresolved due to lack of statistical support at internal nodes. A detailed discussion of sectional concepts, inter-clade relationships based on the plastid framework, and the morphological affinities among lineages within the Asian clade of *Bulbophyllum* is provided below.

### Asian-Malesian clade

The Asian-Malesian lineage includes 26 sections of the Asian clade (Figs. 4 and 5). Within this lineage, section *Trias* forms a well-supported clade, consistent with its historical recognition as a distinct genus (Cameron et al., 1999; Chase et al., 2003; van den Berg et al., 2005), with all included species aligning with the concept of section *Trias* (Vermeulen, 2014). However, Gamisch and Comes (2019) recovered *Trias* as closely related to section *Biflorae*, which was not sampled in the current study. The phylogenetic placement of section *Biflorae* warrants further investigation. Morphologically, it appears to belong to the *Cirrhopetalum* alliance and shares little similarity with section *Trias* (Vermeulen, 2014).

The phylogenetic relationship of *Bulbophyllum blepharistes* Rchb.f. (section *Blepharistes*), *Bulbophyllum ayuthayense* J.J.Verm., Schuit. & de Vogel (section *Drymoda*), and a single species, *Bulbophyllum polliculosum* Seidenf. (section *Lemniscata*) remains ambiguous in this study due to lack of statistical support. The taxonomic placement of *B. blepharistes* is challenging due to its unique combination of floral and vegetative traits, including two-leaved pseudobulbs, a *Cirrhopetalum*-like inflorescence, and morphologically distinctive subumbellate flowers (Vermeulen, 2014; Vermeulen et al., 2014).

Section *Lemniscata* is recovered as polyphyletic, with *B. polliculosum* forming a close relationship with section *Drymoda*, clearly distinct from the remaining representatives of section *Lemniscata* (*Lemniscata* II: Fig. 5). It should be noted that *B. polliculosum* was only tentatively assigned to section *Lemniscata*, while vegetative similarities with section *Drymoda* were emphasised (Schuiteman et al., 2023). Section *Drymoda*, which has deciduous, two-leaved, discoid pseudobulbs, is traditionally recognised as a separate genus based on its unique floral traits, including a rigid non-flexible labellum, and an elongated column with the lateral sepals attached to the apex of the column-foot (Seidenfaden and Smitinand, 1961). Garay et al. (1994) transferred two species of *Bulbophyllum* to *Drymoda* based on the structure of the column-foot only. The first, *Bulbophyllum gymnopus* Hook.f. is here included in sect. *Altisceptrum*, the second, *B. digitatum* J.J.Sm. of sect. *Gongorodes* J.J.Sm., was not sampled by us but we included the related *B. scorpio*. Neither *B. gymnopus* nor *B. scorpio* are inferred as being closely related to sect. *Drymoda* or to each other in our tree. *Bulbophyllum polliculosum* is vegetatively similar to section *Drymoda* but exhibits a *Cirrhopetalum*-like floral morphology (Seidenfaden, 1973; Garay et al., 1994). Given the strong statistical support for this clade, further investigations with expanded taxon sampling (including the type of section *Drymoda*, *Bulbophyllum drymoda* J.J.Verm., Schuit. & de Vogel) are needed to clarify the placement of *B. polliculosum* relative to *Drymoda*. If *B. polliculosum* is moved from section *Lemniscata* to section *Drymoda*, both sections are here supported as monophyletic. The phylogenetic placement of *Bulbophyllum infundibuliforme* J.J.Sm. (section *Hymenobractea*), *Bulbophyllum levyae* Garay, Hamer & Siegerist (section *Lepidorhiza*), species of section *Intervallatae*, and the remaining species of section *Lepidorhiza*, aligns with findings by Simpson et al. (2024). *Bulbophyllum infundibuliforme* and species of section *Intervallatae* share morphological similarities including rather small pseudobulbs and distichously arranged flowers on the inflorescence rachis as described by De Witte and Vermeulen (2010), who included *B. infundibuliforme* within section *Intervallatae*. *Bulbophyllum levyae* has been assigned to section *Lepidorhiza* on the basis of its verrucose roots, believed to be a diagnostic character for this section (Vermeulen et al., 2015). However, its floral characters agree better with species in section *Intervallatae*, especially in the relatively very short petals and the relatively long lip compared to the length of the sepals (Vermeulen et al., 2015). The phylogenetic placement of section *Intervallatae* alongside the other species of section *Lepidorhiza* reflects their overall morphological similarity, particularly a distichous raceme, often with flowers opening in succession, only that species in section *Lepidorhiza* differ in having verrucose roots rather than smooth ones, have generally larger pseudobulbs, with more prominent floral bracts and flowers with differently proportioned petals and lip, and bear a more elongated raceme (Vermeulen, 2014; Vermeulen et al., 2015). In *Hymenobractea* the roots are papillose (Schuiteman and Lambkin, 2024), while they are glabrous in species traditionally assigned to section *Intervallatae* (Vermeulen et al., 2015). Section *Hymenobractea* is recognised by Vermeulen (2014). Taking into account its sister position to section *Intervallatae* in the current analysis, it can still be maintained based on several morphological characters, including non-resupinate flowers, infundibuliform floral bracts, strongly reduced pseudobulbs, and a simple, smooth, thin-textured lip (Vermeulen, 2014).

Sections *Beccariana*, *Sestochilus*, and *Stenochilus*, are well defined monophyletic groups with the latter two recovered as successive sister groups (Gamisch and Comes, 2019). These correspond to well-defined groups characterised by similar plant habit, often substantial in size, number of flowers per inflorescence; multiple in *Beccariana* and reduced to single flowers in sections *Sestochilus* and *Stenochilus* (Hosseini et al., 2016; Vermeulen, 2014). In sect. *Stenochilus* the flowers are non-resupinate whereas they are resupinate in sect. *Sestochilus* (Vermeulen, 2014).

Section *Ione* is characterised by distinct morphological traits, including two sets of pollinia, two viscidia, and a pollinarium with stipes and an adhesive disc (Vermeulen, 2014). Similar to Gamisch and Comes (2019), the phylogenetic position of section *Ione* in the present study remains unresolved due to lack of statistical support at internal nodes. There is, however, no doubt that the section is deeply nested within *Bulbophyllum*.

The phylogenetic placement of section *Lemniscata* + *Tripudianthes* in this study largely aligns with the findings of Gamisch and Comes (2019), who also recovered these groups as closely related to species of section *Racemosae*. Sections *Tripudianthes* and *Lemniscata* have long been considered closely related (Seidenfaden, 1979) because of their plant habit of bifoliate, deciduous pseudobulbs, similar to section *Drymoda*. *Tripudianthes* was subsumed into *Lemniscata* by Vermeulen (2014). Based on our sampling, it seems possible that *Lemniscata* and *Tripudianthes* are sister groups that could be reinstated as distinct sections. Section *Racemosae* forms a well-defined and morphologically distinct lineage as described by Hooker (1825), with single-leaved, non-deciduous pseudobulbs and racemose inflorescences. All species within this section in the present study conform to the concept of *Racemosae* as outlined by Vermeulen (2014).

Historically, section *Achrochaene* was recognised as the genus *Achrochaene* by Lindley (1853) distinguished by two sets of pollinia with bifid stipes (Vermeulen, 2014). Although section *Achrochaene* is recovered as a satellite clade of the *Cirrhopetalum* alliance group (CAG) in this study, this relationship lacks statistical support leaving the phylogenetic placement of these lineages unresolved.

### The Cirrhopetalum alliance

The CAG comprises a group of *Bulbophyllum* sections most of which used to be distinguished from *Bulbophyllum* as the genus *Cirrhopetalum* (Seidenfaden, 1973). This group is morphologically characterised by sub-umbellate racemes and elongated sepals that are inwardly twisted and partially connate (Garay et al., 1994). Earlier classifications placed several sections within the *Cirrhopetalum* alliance, namely *Biflorae*, *Blepharistes*, *Brachyantha*, *Cirrhopetaloides*, *Cirrhopetalum*, *Emarginatae*, *Ephippium*, *Eublepharon*, *Lemniscata*, *Macrostylidia*, *Plumata*, *Recurvae* (nom. inval. = *Ephippium*), *Reptantia*, and *Rhytionanthos* (Chen and Vermeulen, 2009; Vermeulen, 2014; Vermeulen et al., 2014, 2015). Our findings support a strongly supported CAG comprising the sections *Brachyantha*, *Cirrhopetaloides*, *Cirrhopetalum*, *Emarginatae*, *Ephippium*, *Macrostylidia*, *Plumata*, and *Rhytionanthos* (Fig. 5). These findings broadly correspond with previous classifications (Chen and Vermeulen, 2009; Vermeulen et al., 2014, 2015), although representatives of two sections (*Eublepharon* and *Biflorae*) previously assigned to the alliance are not included in our sampling. Additionally, this study confirms the inclusion of section *Desmosanthes* and *Bulbophyllum ambrosia* (Hance) Schltr. (assigned to section *Leopardinae* but probably not closely related to the type species of that section, see below) within the *Cirrhopetalum* alliance, consistent with previous molecular phylogenetic studies (Hu et al., 2020; Thawara et al., 2024).

Phylogenetic relationships within the *Cirrhopetalum* alliance are largely consistent with the findings of Hu et al. (2020) and Thawara et al. (2024) showing nine major divisions within the CAG designated as follows: EMA (*Emarginatae*), CIR (*Cirrhopetaloides*), EPH (*Ephippium*), CIRR I (*Cirrhopetalum* I), BRA (*Brachyantha*), PLU (*Plumata*), and CIRR II (*Cirrhopetalum* II), DES (*Desmosanthes*), EUB&RHY (*Eublepharon* and *Rhytionanthos*), corresponding with the broader taxonomic groupings identified in the present study namely: G12, G13, G14, G16, G21, G23, G24, G25 and G27 respectively (with the exception of section *Eublepharon* (EUB), which was not sampled in the present study).

The first arm of the core CAG (G11) comprises an early-diverging lineage (G12) that supports *Bulbophyllum emarginatum* (Finet) J.J.Sm. of section *Emarginatae* and *Bulbophyllum spathulatum* (Rolfe ex E.W.Cooper) Seidenf. of section *Brachyantha*, which are recovered on the same node with strong statistical support, consistent with previous studies (Hu et al., 2020; Thawara et al., 2024). The remaining taxa within G11 form two non-monophyletic subclades corresponding to sections *Cirrhopetaloides* (Hu et al., 2020) and *Ephippium* (Vermeulen et al., 2015). Additionally, *Bulbophyllum fascinator* (Rolfe) Rolfe, formerly placed in the genus *Mastigion* Garay, Hamer & Siegerist based on the *B. fascinator* group (Garay et al., 1994) is resolved within G13, indicating that the *B. fascinator* group represents only a subgroup within a larger sectional framework.

The remaining species within the CAG are grouped into two strongly supported sister clades, G16 and G19. Subclade G16 includes *Bulbophyllum longiflorum* Thouars, the designated type species of the genus *Cirrhopetalum*, thereby establishing the concept of section *Cirrhopetalum* as defined by Reichenbach (1861). This section is characterised by pseudoumbellate or capitate inflorescences, with lateral sepals that originate from a divergent base and twist once to form a convex blade, their outer margins conjoined to create a tunnel-like passage around the lip (Garay et al., 1994; Vermeulen, 2014). *Bulbophyllum macraei* (Lindl.) Rchb.f. and *Bulbophyllum delitescens* Hance are also recovered within this group, aligning morphologically with section *Cirrhopetalum* (Vermeulen, 2014). These findings differ from their earlier placement in section *Macrostylidia* by Garay et al. (1994), where they were classified based on their conspicuous, blade-like twisted stelidia. The present phylogenetic evidence suggests that this character alone is insufficient for delimiting these two sections. Furthermore, *Bulbophyllum umbellatum* Lindl., the type species of section *Brachyantha*, is also placed within this clade, illustrating that section *Brachyantha* is a synonym of section *Cirrhopetalum* in the strict sense.

Subclade G19 within CAG is subdivided into two strongly supported clades, G20 and G25. Subclade G20 comprises *Bulbophyllum skeatianum* Ridl., *Bulbophyllum thaiorum* J.J.Sm., and *Bulbophyllum farreri* (W.W.Sm.) Seidenf., representing species within section *Brachyantha*, characterised by relatively smaller, narrower flowers with obtuse, unadorned petals (Garay et al., 1994). We are unable to explain why *Bulbophyllum retusiusculum* Rchb.f., which is morphologically very similar to *B. skeatianum*, is not recovered in this clade and is instead placed in clade G13. Section *Plumata* characterised by tassel-like projections at the petal apices, elongate lateral sepals, and a longitudinally furrowed column face (Vermeulen, 2014; Vermeulen et al., 2014) is also found embedded within the broader subclade G20. Within G20, the remaining species from subclade G24 (ranging from *B. inconspicuum* to *Bulbophyllum hirundinis* (Gagnep.) Seidenf.) align morphologically with subclade G21. The phylogenetic placement of G22 aligns with findings by Hu et al. (2020) and Thawara et al. (2024), who retrieved a basal clade of section *Cirrhopetalum* (G24) closely related to two species from section *Plumata* included in their studies.

The phylogenetic placement of species from sections *Rhytionanthos*, *Cirrhopetalum*, and *B. ambrosia* section *Leopardinae* nested within section *Desmosanthes* (G25) largely aligns with previous studies by Hu et al. (2020) and Thawara et al. (2024), which also identified a close phylogenetic relationship between *B. ambrosia* and species from sections *Rhytionanthos* and *Desmosanthes*. However, in the present study, the precise phylogenetic placement of *B. ambrosia* and section *Rhytionanthos* within the broader section *Desmosanthes* remains unresolved.

Hu et al. (2020) and Thawara et al. (2024) sampled several species of section *Leopardinae*, including the type species *Bulbophyllum leopardinum* (Wall.) Lindl. ex Wall. in their analyses. Both studies recovered this group as sister to the CAG, albeit without support in both studies. These findings, consistent with the present study, suggest that *B. ambrosia* may not be a proper representative of section *Leopardinae* and support the recognition of a monospecific section for *B. ambrosia* (Hu et al., 2020; Thawara et al., 2024).

### Malesian-Papuasian clade

The second arm of the Asian clade of *Bulbophyllum* comprises the Malesian-Papuasian clade encompassing 27 sections (Fig. 6 and Fig. 7). Within this clade, sections *Brachystachyae*, *Hirtula* and *Stachysanthes* form a strongly supported lineage (E1), followed by section *Planibulbus* (E2), largely consistent with findings by Simpson et al. (2024). These four sections are primarily found in the Malesian region and are characterised by racemose inflorescences (Vermeulen, 2014; Vermeulen et al., 2015). Subclade E1 is mainly West Malesian and consists only of sections that are absent from New Guinea (section *Hirtula*) or are poorly developed there, with few or no endemic species (sections *Stachysanthes* and *Brachystachyae*), while section *Planibulbus* is West Malesian and absent from New Guinea (Vermeulen, 2014; Vermeulen et al., 2015). Sections *Brachystachyae* and *Stachysanthes* are historically linked by the presence of highly reduced or nearly absent pseudobulbs, a relationship first proposed by Schlechter (1911-1914), who recognised their morphological similarities. In contrast, section *Hirtula* is a more complex taxon, characterised by well-developed pseudobulbs in all species within the group (Vermeulen, 2014; Vermeulen et al., 2015). The pattern suggests an evolution of reduced pseudobulbs in the ancestor of the *Brachystachyae* + *Stachysanthes* + *Hirtula* clade, that is reversed within section *Hirtula*. Further work with more comprehensive sampling and ancestral character state reconstruction is required to understand the evolution of vegetative traits within these groups.

Comprising only two species, section *Planibulbus* is one of the rare sections of the genus characterised by prostrate pseudobulbs borne on an elongated, wiry rhizome and racemose inflorescences (Vermeulen et al., 2015). This small section is of special interest because it is sister to the huge Papuasian clade (E3).

### The Papuasian clade

Within the Malesian-Papuasian clade, a strongly supported Papuasian clade (E3) is recovered, encompassing the remaining 23 sections of the Asian clade (Fig. 6 and Fig. 7), all of which have the majority of their species occurring on the island of New Guinea, with several endemic to the region (Vermeulen, 2014; Vermeulen et al., 2015). Although broader phylogenetic relationships within this lineage remain uncertain due to low support for some internal nodes, several well-supported sectional alliances can be identified, as outlined below (Fig. 6 and Fig. 7).

A close relationship is identified between sections *Uncifera* and *Imitatores* (E4: 99.5/95), while the phylogenetic position of section *Gongorodes* among other lineages within the Papuasian clade remains unclear due to lack of statistical support at internal nodes. As mentioned above, there is no support for a close alliance between sects. *Gongorodes* and *Drymoda*, as proposed by Garay et al. (1994). Section *Uncifera* consists of eight species morphologically distinguished by uncinate stelidia (Vermeulen, 2014). Section *Imitatores* was previously included by Vermeulen (1993) in sect. *Macrouris* but was removed due to the stigma not protruding proximally (Vermeulen et al., 2014). Our findings support the exclusion of section *Imitatores* from section *Macrouris*. *Bulbophyllum scorpio* of sect. *Gongorodes* is a rare species within the genus, distinguished by its intricately folded or incised petal tips, an otherwise unique trait that could align it with section *Leopardinae* (Vermeulen, 2014).

Section *Macrobulbon* has been included in section *Beccariana* (Vermeulen, 2014) but our results convincingly show that these sections are quite distinct and not phylogenetically close. Our analysis includes ‘true’ species of section *Beccariana* from New Guinea, such as *Bulbophyllum foetidum* Schltr. (Fig. 4: F10), as well as five out of six known species of section *Macrobulbon* (Fig. 6). The latter forms a well resolved monophyletic group in the Papuasian clade, whereas section *Beccariana* resides in the Asian-Malesian clade. Section *Macrobulbon* is endemic to New Guinea and is characterised by the large subglobose pseudobulbs, fleshy, often pendent, pruinose leaves (Vermeulen, 1993), and with condensed racemes of fetid-smelling flowers held against the pseudobulb (Yukawa and Karasawa, 1997). Section *Macrobulbon* includes some of the largest species within *Bulbophyllum*, such as *B. phalaenopsis* J.J.Sm. (Vermeulen, 1993; Teoh, 2021), which may have the largest leaves in Orchidaceae in terms of surface area.

Our findings reveal a close relationship between sections *Macrouris* and *Lepanthanthe*, with sect. *Macrouris* rendered paraphyletic by a well-supported lineage consisting of species of *Lepanthanthe* nested within it. These relationships align with Vermeulen (2014), who identified a close relationship between these two sections characterised by a racemose inflorescence and articulated pedicels. Vermeulen (1993) also identified a close relationship between these two sections and considered section *Macrouris* as the paraphyletic root of section *Lepanthanthe*. Species of section *Lepanthanthe* as first delimited by Schlechter (1911-1914) are characterised by a pendulous habit, sepals bearing subulate projections at their tips, and a lip with lobes near its base (Vermeulen, 1993). However, the three representative species of this section included in the present study namely, *Bulbophyllum bulliferum* J.J.Sm., *Bulbophyllum trachypus* Schltr., and *Bulbophyllum erinaceum* Schltr., were originally placed by Schlechter (1911-1914) in section *Trachyrhachis* Schltr., distinguished by a creeping rather than a pendulous rhizome but with similar floral characters. Section *Trachyrhachis* was later synonymised into section *Lepanthanthe* by Vermeulen (1993, 2014). Interestingly, in the present study, *Bulbophyllum trifilum* J.J.Sm. has a pendulous habit yet resembles *Bulbophyllum fonsflorum* J.J.Verm. and *Bulbophyllum macrourum* Schltr. in floral morphology rather than any species of section *Lepanthanthe* (Vermeulen, 1993). Conversely, *Bulbophyllum olivinum* J.J.Sm., *Bulbophyllum chloranthum* Schltr., and *Bulbophyllum subpatulum* J.J.Verm. of section *Macrouris* are vegetatively much more similar to species of section *Trachyrhachis*, now recognised as *Lepanthanthe* (Vermeulen, 1993). The phylogenetic placement of *Lepanthanthe* in the present study suggests that it may represent a highly specialised lineage within section *Macrouris* characterised by its more complex flowers. However, limited taxon sampling within *Lepanthanthe* and *Macrouris* in the present study restricts broader conclusions, highlighting the need for additional studies to clarify the evolutionary relationships and identify taxonomically informative characters.

The phylogenetic placement of sections *Pelma*, *Monanthaparva*, *Monanthes*, and *Oxysepala* in this study aligns with the findings of Thawara et al. (2024), who likewise recovered the latter three sections as closely related with strong nodal support. In their study, *Bulbophyllum membranaceum* Teijsm. & Binn. section *Monanthaparva* was resolved as an early-branching lineage closely related to the *Monanthes*/*Oxysepala* group, a pattern consistent with the present results. Furthermore, the close relationship between *Monanthes* and *Oxysepala* inferred here is congruent with findings by Gamisch and Comes (2019) and Simpson et al. (2024).

Sections *Pelma* (Schlechter, 1911-1914) and *Monanthaparva* (Ridley, 1896) are morphologically distinct groups, as described by Vermeulen (2014). Sections *Monanthes* and *Oxysepala* were merged by Vermeulen (2014) under *Monanthes*, although Vermeulen et al. (2015) adopted *Oxysepala* as the accepted name for the combined section. The connate lateral sepals of *B. membranaceum* resemble those of many species in section *Monanthes* (Vermeulen et al., 2015). Additional sampling of *B. membranaceum* using different markers is needed to clarify its phylogenetic placement within the broader E10 clade.

Morphologically, section *Monanthaparva* is still poorly understood as it includes species that previous authors have included in sections *Minutissima* and *Oxysepala* (Vermeulen et al., 2015; Averyanov et al., 2024). A much broader sampling is needed here to assess relationships, and this should include species of Schlechter’s section *Nematorhizis*, which has been sunk in section *Minutissima* by Vermeulen and O’Byrne (2011) and Vermeulen (2014). Sections *Monanthaparva* and *Monanthes* possess single-flowered inflorescences, whereas section *Pelma* may have single-flowered as well as racemose inflorescences (Vermeulen, 2014; Vermeulen et al., 2015).

Section *Monanthes* comprises a large group including species historically recognised within sections *Polyblepharon* Schltr. and *Hybochilus* Schltr., with *Polyblepharon* characterised by the presence of two auricles at the base of the lip with an adjacent callus, connate lateral sepals and usually non-resupinate flowers (Schlechter, 1911-1914). Section *Oxysepala* is distinguished by its reduced pseudobulbs, fleshy leaves, free lateral sepals, and very short, single-flowered inflorescences (Bentham, 1883). This study represents the broadest sampling achieved thus far in this alliance, but species coverage remains limited to approximately 10% of the known species diversity within section *Monanthes*. Our results suggest that merging of sections *Monanthes* and *Oxysepala* is well-supported (Vermeulen, 2014).

The remainder of the Papuasian lineage aligned into one large well supported clade (E12), which includes only sections characterised by single flowered inflorescences (Vermeulen, 2014) (Fig. 7). Within this group evolutionary relationships are mostly well resolved. Sections *Monosepalum* and *Epicrianthes* are closely related in the present study and are united morphologically by the presence of clavate or ornate appendages on the petals (Vermeulen, 2014). Section *Monosepalum* is distinguished by the much larger, long-pedunculate flowers with all three sepals mutually connate (vs. small, subsessile flowers with free sepals in section *Epicrianthes*) (Vermeulen, 2014; Vermeulen et al., 2015).

Sections *Hyalosema* (Schlechter, 1911) and *Hoplandra* (Vermeulen, 2008) represent well-defined morphologically similar sections and are each resolved as monophyletic sister groups based on current sampling. The primary distinction between them lies in pollinia number, with section *Hyalosema* possessing two pollinia, whereas section *Hoplandra* has four pollinia (Vermeulen, 2014). Species of both sections usually have quadrangular pseudobulbs along a creeping rhizome (Vermeulen et al., 2015). Section *Tapeinoglossum* Schltr., which we have not been able to sample, is similar to section *Hyalosema*, but is distinguished by all three sepals being mutually connate and having much smaller flowers that are pilose inside (Vermeulen, 2014).

Section *Polymeres* is highly polyphyletic forming a basal grade, consistent with Simpson et al. (2024). *Polymeres* is characterised by creeping or patent rhizomes, single-flowered inflorescences and a widened and winged column foot (Vermeulen, 2014). Further studies including increased coverage of species and ancestral trait reconstruction are required to evaluate the taxonomic utility of these morphological characters within a phylogenetic framework.

Our findings reveal a strongly supported relationship of section *Piestobulbon* with *B. zebrinum*, *B. dictyoneuron* and *B. rubipetalum* of section *Polymeres* (Fig. 7). Section *Piestobulbon*, originally described by Schlechter (1923), is characterised by long, single-flowered inflorescences with brown papery basal bracts (Vermeulen, 2014). Further studies incorporating additional samples from section *Piestobulbon* are necessary to clarify the phylogenetic placement of this section within the broader genus *Bulbophyllum*.

Section *Pedilochilus* was described as a distinct genus by Schlechter (1905) and widely accepted since; it is morphologically characterised by a slipper-shaped lip and petals that exhibit an S-shaped twist along the midvein. Its phylogenetic placement within E19 may indicate that *Pedilochilus* represents a morphologically derived lineage within this clade.

The phylogenetic placement of *B. nasica* and *B. maxillare* (section *Polymeres*), along with taxa from section *Codonosiphon* is unexpected, given that *Codonosiphon* is distinguished by a column-structure with a reduced column-foot and translucent, curved column without long stelidia, and often with an immobile labellum (Vermeulen, 2014). These findings suggest a need for further investigation to clarify the phylogenetic placement of *B. nasica* and *B. maxillare* within *Bulbophyllum*. On morphological grounds, one would expect the latter two species to group with such species as *B. zebrinum* in clade E19 rather than with the distinctive section *Codonosiphon*.

Consistent with findings by Simpson et al. (2024), species from sections *Brachypus* and *Papulipetalum* form a basal grade. Morphologically, these sections share similarities with *Polymeres*, except for the absence of the widened and winged column foot characteristic of *Polymeres* (Vermeulen, 2014). Both groups consist of medium-sized species with clustered pseudobulbs, often bearing conspicuous bracts and one-flowered inflorescences, supporting their unification as proposed by Vermeulen (2008). Schlechter (1911-1914) also recognised their close relationship and treated them as adjacent taxa. However, the phylogenetic placement of three species of section *Brachypus* namely: *B. lineolatum*, *B. phaeglossum*, and *B. versteegii* remains unresolved in the present study, warranting further investigations.

The remaining subclade of the Papuasian lineage comprises sections *Fruticicola* together with the monophyletic sister clades *Peltopus* and *Epibulbon*, all three of which were historically recognised by Schlechter (1911-1914). Section *Peltopus* forms a well-defined clade, characterised by flowers with a bulbous lobe at the base of the column that fits into a cavity at the base of the fleshy labellum (Schlechter, 1911-1914). Sections *Fruticicola* and *Epibulbon* were subsumed into section *Polymeres* by Vermeulen and O’Byrne (2008) based on the similar column morphology (Vermeulen et al., 2020). Species of sections *Epibulbon* and *Fruticicola* always have superposed pseudobulbs on stem-like rhizomes where new shoots arise from the top of the old pseudobulb (Vermeulen, 1993), whereas most species in sections *Polymeres* and *Peltopus* have non-superposed pseudobulbs often clustered or arranged along a creeping rhizome (Vermeulen, 2014; Vermeulen et al., 2015). Extended taxon sampling across these sections is necessary to fully resolve their phylogenetic positions and clarify taxonomic relationships within and between them. Based on our results, it seems likely that sections *Epibulbon* and *Fruticicola* can be reinstated, but section *Polymeres* requires more study in order to arrive at a morphologically supported classification.

## FUTURE PROSPECTS

Our phylogenomic analyses provide strong support for key taxonomic revisions and the recognition of major evolutionary lineages within *Bulbophyllum*. In particular, the reinstatement of sections *Tripudianthes* and *Fruticicola* is strongly supported, with both sections recovered as monophyletic with maximum statistical support. The *Cirrhopetalum* alliance is likewise recovered as monophyletic with maximum support (Hu et al., 2020), but there is little support for the many sections previously recognised within this alliance. In addition, the Papuasian clade forms a well-supported lineage; the more basal members of this clade tend to have multi-flowered inflorescences, whereas the more derived members have single-flowered inflorescences (Vermeulen, 2014).

Many sections within *Bulbophyllum* differ only in relatively loose combinations of morphological characters (Hosseini et al., 2016; Hu et al., 2020). Vermeulen called these ‘polythetic sets’ (Vermeulen, 1993). In every section, there are species that lack one or more of the diagnostic traits yet still appear to belong there; conversely, species that initially seem to fit a sectional diagnosis may exhibit additional characters that contradict this affinity (Garay et al., 1994; Vermeulen, 2014; Vermeulen et al., 2015; Hosseini et al., 2016). Such inconsistencies are evident in groups such as the *Cirrhopetalum* alliance, in which all sections are recovered as non-monophyletic. This pattern is congruent with the findings of Hu et al. (2020), who, despite conducting ancestral character reconstructions within the group based on 17 selected characters, were unable to identify clear synapomorphies for most well-supported clades or subclades. A broader range of putatively diagnostic taxonomic characters should therefore be explored to enable a revised sectional classification of the *Cirrhopetalum* alliance, within which sectional boundaries and diagnostic characters can be identified with greater confidence.

Other non-monophyletic sections requiring priority taxonomic revision include *Brachypus/Papulipetalum*, *Drymoda*/*Lemniscata*, *Hymenobractea*/*Lepidorhiza*/*Intervallatae*, *Macrouris*/*Lepanthanthe*, *Monanthes*/*Oxysepala*, and *Polymeres*, as revealed by this study. These patterns likely reflect convergent evolution, heavy reliance on homoplastic characters in sectional delimitation, and historically inconsistent sectional concepts that were either overly narrow or overly broad (Seidenfaden, 1979; Seidenfaden and Wood, 1992; Vermeulen, 2014; Vermeulen et al., 2014). These sections require comprehensive recircumscription based on increased taxon sampling, as the floral and vegetative characters traditionally used to diagnose them are homoplastic and have proven unreliable.

This study is based on plastid phylogenomic data, which, while valuable for resolving deeper evolutionary relationships, captures only a single maternally inherited genealogical history (Cafasso et al., 2005; Ruhlman and Jansen, 2014; Serna-Sánchez et al., 2021). To complement the plastid phylogenetic analyses and to obtain a more complete picture of evolutionary relationships within *Bulbophyllum*, nuclear phylogenomic approaches such as target enrichment or hybrid-capture sequencing should be prioritised in future studies (Dodsworth et al., 2019; Johnson et al., 2019). Despite these limitations, the plastid phylogenomic framework presented in this study produced many groupings of species that were already believed to be related on morphological grounds and provides a strong working hypothesis for *Bulbophyllum* sectional relationships that serves as an important foundation for future hypothesis testing using nuclear-derived datasets.

## CONCLUSIONS

This phylogenomic analysis represents a substantial advancement in our understanding of evolutionary relationships within the mega-diverse genus *Bulbophyllum*. Based on 63 plastid genes, the study expanded taxonomic coverage of Asian and Pacific species to include 65 sections (eight represented in a molecular phylogenetic analysis for the first time) demonstrating complex evolutionary relationships within the diverse Asian clade. Well-supported section alliances are revealed within the Asian clade, providing a framework for assessing sectional relationships. A total of 20 sections were resolved as monophyletic based on current representative sampling (i.e., *Achrochaene*, *Beccariana*, *Brachystachyae*, *Codonosiphon*, *Epibulbon*, *Epicrianthes*, *Fruticicola*, *Hoplandra*, *Hyalosema*, *Hirtula*, *Ione*, *Macrobulbon*, *Pelma*, *Peltopus, Racemosae*, *Sestochilus*, *Stachysanthes*, *Stenochilus*, *Trias*, and *Tripudianthes*) while evidence of non-monophyly was identified in 21 sections, including *Brachypus*, all *Cirrhopetalum* alliance sections, *Lepidorhiza*, and *Polymeres*. These findings highlight the need for comprehensive sectional revision within *Bulbophyllum* supported by a careful analysis of character evolution within the genus. Further work with more extensive taxon sampling including the other 16 *Bulbophyllum* sections not included in this study, and the integration of nuclear genomic data is required to clarify broader sectional relationships and provide robust assessment of sectional taxonomy.

Beyond taxonomic implications, this phylogenetic framework provides a foundation for exploring the evolutionary processes that have shaped diversification patterns within this diverse Asian lineage. The remarkable diversity of *Bulbophyllum* species across the Asia-Pacific region represents an exceptional model system for investigating ancestral range evolution in one of the world’s most biogeographically complex zones. Our phylogenetic analysis reveals that the biogeographic structure previously observed between continental lineages within the genus is also reflected within the Asian clade, with strong support for a Papuasian lineage. Combined with the spectacular ecological and morphological diversity exhibited among species, this phylogenetic framework establishes *Bulbophyllum* as a uniquely powerful system for macroevolutionary studies. Future research employing macroevolutionary approaches including divergence dating, diversification rate analyses, and ancestral state reconstruction, while taking the geological and climatological history into account, will be essential to fully realize the potential of this intriguing study system. Such analyses will elucidate patterns of trait evolution and their associations with shifts in diversification rates, thereby providing deeper insights into the evolutionary dynamics underlying this remarkable diversity.

## Acknowledgements

The authors acknowledge the use of next-generation sequencing services and facilities of the Australian Genomic Research Facilities (AGRF, Melbourne), the Biomolecular Resource Facility (BRI, Canberra) and the contribution of Bioplatforms Australia (enabled by NCRIS) in the generation of data used in this publication. The authors thank Brendan Lepschi (CANB), Michael Mathieson (BRI), Frank Zich (CNS), Heidi Zimmer (CANBR), Alex Baribeau (KEW), and Bala Kompallii (KEW) for providing access to research collections for this study. We are grateful to Mike Fay (KEW), Ilia Leitch (KEW), and Pérez-Escobar (KEW) for their valuable support and advice at Kew. The authors acknowledge funding by the Australian Tropical Herbarium, an unincorporated joint venture of James Cook University, the State of Queensland, CSIRO and Director of National Parks. A preprint of the manuscript was released online (Nanjala et al., 2026; https://doi.org/10.64898/2026.03.30.715161).

## Author contributions

Conceptualisation: CN, LS, KN, and MC; Funding acquisition: KN, CN; Supervision: KN, LS and DC; Data curation: CN, MC, LS and SG; Resources: MC, LS, VP, AH, SG, GF, AS, and KN; Investigation and Methodology: CN, SB, and JN; Formal Analysis: CN; Writing original draft: CN; Writing-Review and editing: CN, KN, LS, DC, MC, AS, AH, VP, JN, SB, GF, and SG. All authors contributed to the article and approved the submitted version.

## Supplementary data

**Table S1.** Sectional coverage of *Bulbophyllum* species included in this study

**Table S2.** Details of plant material used in this study including voucher information and provenance.

**Table S3.** List of sample names and the specific genes included in the phylogenetic analysis.

**Figure S1.** Maximum likelihood phylogenetic tree showing the relationships between *Bulbophyllum*, its sister group *Dendrobium*, and other outgroups representing all major lineages of Orchidaceae. The tree also illustrates the relationships among major biogeographic lineages within *Bulbophyllum*. The tree is based on 468 taxa and was constructed using 63 plastid loci. Branch support values are shown at each node, with SH-aLRT (Shimodaira-Hasegawa-like Approximate Likelihood Ratio Test) values presented first, followed by UFBoot (Ultrafast Bootstrap) support values.

**Figure S2.** Maximum likelihood phylogenetic tree with original branch lengths showing the relationships between *Bulbophyllum*, its sister group *Dendrobium*, with all other outgroups pruned. The tree also illustrates the relationships among major biogeographic lineages within *Bulbophyllum*. The tree is based on 468 taxa and was constructed using 63 plastid loci. Branch support values are shown at each node, with SH-aLRT (Shimodaira-Hasegawa-like Approximate Likelihood Ratio Test) values presented first, followed by UFBoot (Ultrafast Bootstrap) support values. The scale bar indicates branch length, corresponding to 0.02 expected substitutions per site.

## Conflicts of interest

The authors declare no conflict of interest.

## Funding

This work was supported by the Hermon Slade Foundation (HSF 16-04) and the Australian Orchid Foundation (AOF 325.18, AOF 360-202). The authors acknowledge funding by the Australian Tropical Herbarium, an unincorporated joint venture of James Cook University, the State of Queensland, CSIRO and Director of National Parks.

## Data availability statement

The data used for this study has been deposited in the European Nucleotide Archive (ENA) at EMBL-EBI under accession number PRJEB87630 (https://www.ebi.ac.uk/ena/browser/view/PRJEB87630).

## Notes

### Competing Interest Statement

The authors have declared no competing interest.

### Summary of Updates

This version of the manuscript has been revised to update 2 references that were added according to the reviewers' comments

