## Supplementary Data for "Phylogenomics of the mega genus *Bulbophyllum* (Orchidaceae) with implications for infrageneric classification in the Asian clade" (from another version of this preprint: version 2, 2026-07-06)

**Table S1:** Sectional coverage of *Bulbophyllum* species included in this study

| Section | Type species | Clade | Estimated species number in the section | Reference sectional concept | Species number in the present study | Sectional coverage (%) | Type species included |
|---|---|---|---|---|---|---|---|
| *Adelopetalum* (Fitzg.) J.J.Verm. | *Bulbophyllum bracteatum* (Fitzg.) F.M.Bailey | Australasian | 12 | Pridgeon et al. (2014) | 12 | 100 | 1 |
| *Minutissima* Pfitz. | *Bulbophyllum minutissimum* (F.Muell.) F.Muell. | Australasian | 23 | Pridgeon et al. (2014) | 3 | 13 | 1 |
| *Megaclinium* Lindl. | *Bulbophyllum falcatum* (Lindl.) Rchb.f. | African | 19 | Pridgeon et al. (2014) | 4 | 21 | 1 |
| *Ptiloglossum* Lindl. | *Bulbophyllum barbigerum* Lindl. | African | 30 | Pridgeon et al. (2014) | 3 | 10 | 1 |
| *Didactyle* (Lindl.) Cogn. | *Bulbophyllum exaltatum* Lindl. | Neotropical | 7 | Pridgeon et al. (2014) | 3 | 43 | 1 |
| *Furvescens* E.C Smidt, Borba & van den Berg | *Bulbophyllum nagelii* L.O.Williams | Neotropical | 5 | Pridgeon et al. (2014) | 1 | 20 | 0 |
| *Micranthae* Barb Rodr. | *Bulbophyllum micranthum* Barb.Rodr. | Neotropical | 10 | Pridgeon et al. (2014) | 2 | 20 | 0 |
| *Napellii* Rchb.f | *Bulbophyllum napellii* Lindl. | Neotropical | 12 | Pridgeon et al. (2014) | 2 | 17 | 0 |
| *Xiphizusa* Rchb.f | *Bulbophyllum chloropterum* Rchb.f. | Neotropical | 23 | Pridgeon et al. (2014) | 1 | 4 | 0 |
| *Alcistachys* Schltr. | *Bulbophyllum occlusum* Ridl. | Madagascan | 6 | Pridgeon et al. (2014) | 2 | 33 | 1 |
| *Pachychlamys* Schltr. | *Bulbophyllum pachypus* Schltr. | Madagascan | 10 | Pridgeon et al. (2014) | 1 | 10 | 1 |
| *Ploiarium* Schltr. | *Bulbophyllum coriophorum* Ridl. | Madagascan | 75 | Pridgeon et al. (2014) | 1 | 1 | 0 |
| *Achrochaene* (Lindl.) J.J.Verm., Schuit. & de Vogel | *Bulbophyllum kingii* Hook.f. | Asian | 3 | Pridgeon et al. (2014) | 2 | 67 | 0 |
| *Altisceptrum* J.J.Sm. | *Bulbophyllum elongatum* (Blume) Hassk. | Asian | 9 | Pridgeon et al. (2014) | 1 | 11 | 0 |
| *Beccariana* Pfitz. | *Bulbophyllum beccarii* Rchb.f. | Asian | 65 | Pridgeon et al. (2014) | 15 | 23 | 1 |
| *Blepharistes* J.J.Verm., Schuit. & de Vogel | *Bulbophyllum blepharistes* Rchb.f. | Asian | 1 | Pridgeon et al. (2014) | 1 | 100 | 1 |
| *Brachyantha* Rchb.f. | *Bulbophyllum umbellatum* Lindl. | Asian | 26 | Pridgeon et al. (2014) | 8 | 31 | 1 |
| *Brachypus* Schltr. | *Bulbophyllum maxillarioides* Schltr. | Asian | 21 | Pridgeon et al. (2014) | 4 | 19 | 0 |
| *Brachystachyae* Benth. & Hook.f. | *Bulbophyllum repens* Griff. | Asian | 43 | Pridgeon et al. (2014) | 4 | 9 | 1 |
| *Cirrhopetaloides* Garay, Hamer, & Siegrist | *Bulbophyllum longissimum* (Ridl.) J.J.Sm. | Asian | 19 | Pridgeon et al. (2014) | 8 | 42 | 0 |
| *Cirrhopetalum* (Lindl.) Rchb.f | *Bulbophyllum longiflorum* Ridl. | Asian | 28 | Pridgeon et al. (2014) | 14 | 50 | 1 |
| *Codonosiphon* Schltr. | *Bulbophyllum codonanthum* Schltr. | Asian | 77 | Pridgeon et al. (2014) | 10 | 13 | 0 |
| *Desmosanthes* (Blume) J.J.Sm. | *Bulbophyllum croceum* (Blume) Lindl. | Asian | 70 | Pridgeon et al. (2014) | 9 | 13 | 0 |
| *Drymoda* (Lindl.) J.J.Verm., Schuit. & de Vogel | *Bulbophyllum drymoda* J.J.Verm., Schuit. & de Vogel | Asian | 2 | Pridgeon et al. (2014) | 1 | 50 | 0 |
| *Emarginatae* Garay, Hamer, & Siegrist | *Bulbophyllum emarginatum* (Finet) J.J.Sm. | Asian | 3 | Pridgeon et al. (2014) | 1 | 33 | 1 |
| *Ephippium* Schltr. | *Bulbophyllum lepidum* (Blume) J.J.Sm. | Asian | 36 | Pridgeon et al. (2014) | 8 | 22 | 1 |
| *Epibulbon* Schltr. | *Bulbophyllum epibulbon* Schltr. | Asian | 7 | Pridgeon et al. (2014) | 5 | 71 | 0 |
| *Epicrianthes* (Blume) Hook.f. | *Bulbophyllum epicrianthes* Hook.f. | Asian | 40 | Pridgeon et al. (2014) | 5 | 13 | 0 |
| *Fruticicola* (Schltr.) M.A Clem. & D.L Jones | *Bulbophyllum fruticicola* Schltr. | Asian | 25 | Clements & Jones (2002) | 7 | 28 | 1 |
| *Gongorodes* J.J Smith | *Bulbophyllum digitatum* J.J.Sm. | Asian | 2 | Pridgeon et al. (2014) | 1 | 50 | 0 |
| *Hirtula* Ridl. | *Bulbophyllum hirtulum* Ridl. | Asian | 40 | Pridgeon et al. (2014) | 4 | 10 | 0 |

| | | | | | | | |
|---|---|---|---|---|---|---|---|
| *Hoplandra* J.J Verm. | *Bulbophyllum restrepia* (Ridl.) Ridl. | Asian | 10 | Pridgeon et al. (2014) | 5 | 50 | 1 |
| *Hyalosema* Schltr. | *Bulbophyllum grandiflorum* Blume | Asian | 19 | Pridgeon et al. (2014) | 10 | 53 | 1 |
| *Hymenobractea* Schltr. | *Bulbophyllum infundibuliforme* J.J.Sm. | Asian | 2 | Pridgeon et al. (2014) | 1 | 50 | 1 |
| *Imitatores* J.J.Verm., Schuit. & de Vogel | *Bulbophyllum imitator* J.J.Verm. | Asian | 4 | Pridgeon et al. (2014) | 1 | 25 | 0 |
| *Intervallatae* Ridl. | *Bulbophyllum attenuatum* Rolfe | Asian | 44 | Pridgeon et al. (2014) | 3 | 7 | 0 |
| *Lemniscata* Pfitz. | *Bulbophyllum leminiscatum* C.S.P.Parish ex Hook.f. | Asian | 32 | Pridgeon et al. (2014) | 7 | 22 | 0 |
| *Leopardinae* Benth. & Hook.f. | *Bulbophyllum leopardinum* (Wall.) Lindl. ex Wall. | Asian | 7 | Pridgeon et al. (2014) | 1 | 14 | 0 |
| *Lepanthathe* Schltr. | *Bulbophyllum lepanthiflorum* Schltr. | Asian | 17 | Pridgeon et al. (2014) | 3 | 18 | 0 |
| *Lepidorhiza* Schltr. | *Bulbophyllum amplebracteatum* Teijsm. & Binn. | Asian | 28 | Pridgeon et al. (2014) | 15 | 54 | 1 |
| *Macrobulbon* J.J.Sm. | *Bulbophyllum macrobulbum* J.J.Sm. | Asian | 6 | Vermeulen et al. (1993) | 5 | 83 | 0 |
| *Macrostelydia* Garay, Hamer & Siegerist | *Bulbophyllum macraei* (Lindl.) Rchb.f. | Asian | 4 | Pridgeon et al. (2014) | 2 | 50 | 1 |
| *Macrouris* Schltr. | *Bulbophyllum macrourum* Schltr. | Asian | 28 | Pridgeon et al. (2014) | 10 | 36 | 1 |
| *Monanthaparva* Ridl. | *Bulbophyllum striatellum* Ridl. | Asian | 36 | Pridgeon et al. (2014) | 5 | 14 | 0 |
| *Monanthes* (Blume) Aver | *Bulbophyllum tortuosum* (Blume) Lindl. | Asian | 150 | Pridgeon et al. (2014) | 15 | 10 | 1 |
| *Monosepalum* (Schltr.) J.J.Sm. | *Bulbophyllum muricatum* J.J.Sm. | Asian | 3 | Pridgeon et al. (2014) | 1 | 33 | 1 |
| *Oxysepala* Benth. & Hook.f. | *Bulbophyllum clandestinum* Lindl. | Asian | 152 | Vermeulen et al. (2015) | 10 | 7 | 1 |
| *Papulipetalum* Schltr. | *Bulbophyllum papulipetalum* Schltr. | Asian | 22 | Pridgeon et al. (2014) | 4 | 18 | 0 |
| *Pedilochilus* (Schltr) J.J Verm & O'Byrne | *Bulbophyllum papuanum* (Schltr.) J.J.Verm., Schuit. & de Vogel | Asian | 33 | Pridgeon et al. (2014) | 7 | 21 | 0 |
| *Pelma* Schltr. | *Bulbophyllum absconditum* J.J.Sm. | Asian | 26 | Pridgeon et al. (2014) | 8 | 31 | 1 |
| *Peltopus* Schltr. | *Bulbophyllum peltopus* Schltr. | Asian | 33 | Pridgeon et al. (2014) | 4 | 12 | 0 |
| *Piestobulbon* Schltr. | *Bulbophyllum piestobulbon* Schltr. | Asian | 5 | Pridgeon et al. (2014) | 1 | 20 | 1 |
| *Planibulbus* J.J.Verm., Schuit. & de Vogel | *Bulbophyllum planibulbe* (Ridl.) Ridl. | Asian | 2 | Pridgeon et al. (2014) | 1 | 50 | 1 |
| *Plumata* J.J.Verm., Schuit. & de Vogel | *Bulbophyllum plumatum* Ames | Asian | 4 | Pridgeon et al. (2014) | 4 | 100 | 1 |
| *Polymeres* (Blume) J.J Verm & O'Byrne | *Bulbophyllum tenuifolium* (Blume) Lindl. | Asian | 164 | Pridgeon et al. (2014) | 15 | 9 | 0 |
| *Racemosae* Benth. & Hook.f. | *Bulbophyllum careyanum* (Hook.) Spreng. | Asian | 38 | Pridgeon et al. (2014) | 9 | 24 | 0 |
| *Reptantia* J.J.Verm. | *Bulbophyllum reptans* (Lindl.) Lindl. ex Wall. | Asian | 6 | Pridgeon et al. (2014) | 1 | 17 | 1 |
| *Rhytionanthos* Garay, Hamer & Siegerist | *Bulbophyllum helenae* (Kuntze) J.J.Sm. | Asian | 16 | Pridgeon et al. (2014) | 2 | 13 | 1 |
| *Sestochilus* Benth. & Hook.f. | *Bulbophyllum lobbii* Lindl. | Asian | 28 | Pridgeon et al. (2014) | 7 | 25 | 1 |
| *Stachysanthes* (Blume) Aver. | *Bulbophyllum gibbosum* (Blume) Lindl. | Asian | 66 | Pridgeon et al. (2014) | 5 | 8 | 1 |
| *Stenochilus* (Blume) J.J.Sm. | *Bulbophyllum macranthum* Lindl. | Asian | 12 | Pridgeon et al. (2014) | 9 | 75 | 1 |
| *Sunipia* (Lindl.) J.J.Verm., Schuit. & de Vogel | *Bulbophyllum sunipia* J.J.Verm., Schuit. & de Vogel | Asian | 22 | Pridgeon et al. (2014) | 2 | 9 | 0 |
| *Trias* (Lindl.) J.J.Verm., Schuit. & de Vogel | *Bulbophyllum oblongum* (Lindl.) Rchb.f. | Asian | 15 | Pridgeon et al. (2014) | 5 | 33 | 0 |
| *Tripudianthes* (Seidenf.) Szlach. & Kras | *Bulbophyllum tripudians* C.S.P.Parish & Rchb.f. | Asian | 11 | Nguyen et al. (2023) | 2 | 18 | 1 |
| *Uncifera* Schltr. | *Bulbophyllum ochroleucum* Schltr. | Asian | 8 | Pridgeon et al. (2014) | 1 | 13 | 1 |
| **Total** | | | | | **309** | | |

**Legend.** Type species: 1=Included; 0=Unavailable

**Table S2.A:** *Bulbophyllum* samples for phylogenetic analysis. Taxonomic concepts follow World Flora Online database (https://www.worldfloraonline.org).

| Species name | Section | DNA Number | ENA/Genbank Number | Collection Details | Voucher numbers |
|---|---|---|---|---|---|
| *Bulbophyllum absconditum* J.J.Sm. | *Pelma* | CNS_G07471 | PRJEB87630 | M.A.Clements 11242 (CANB) | CANB: 998255.1 |
| *Bulbophyllum acuminatum* (Ridl.) Ridl. | *Ephippium* | CNS_G08295 | PRJEB87630 | M.A.Clements 12862 (CANB) | CANB: 999492.1 |
| *Bulbophyllum acutilingue* J.J.Sm. 1 | *Epibulbon* | CNS_G07612 | ERR13142831 | S.Rose 25 (CANB) | CANB: 1000175.1 |
| *Bulbophyllum acutilingue* J.J.Sm. 2 | *Epibulbon* | HTCG_1923 | PRJEB87630 | S.Rose 25 (CANB) | CANB: 1000175.1 |
| *Bulbophyllum affine* Lindl. | *Stenochilus* | CNS_G07227 | PRJEB87630 | Orchid Research Group 6830 (CANB) | CANB: 1000857.1 |
| *Bulbophyllum agastor* Garay, Hamer & Siegerist | *Macrobulbon* | CNS_G07584 | PRJEB87630 | Orchid Research Group 7672 (CANB) | CANB: ORG 7672 |
| *Bulbophyllum albociliatum* (Tang S.Liu & H.Y.Su) K.Nakaj. | *Cirrhopetaloides* | _ | NC_070215 | N/A | N/A |
| *Bulbophyllum alkmaarense* J.J.Sm. | *Codonosiphon* | CNS_G05454 | ERR13142798 | G.McCraith 076 (CANB) | CANB: 1000233.1 |
| *Bulbophyllum alpinum* (P.Royen) J.J.Verm., Schuit. & de Vogel | *Pedilochilus* | CNS_G07293 | PRJEB87630 | P.VanRoyen 10814 (CANB) | CBG: 8105307.1 |
| *Bulbophyllum alticola* Schltr. | *Codonosiphon* | HTCG_1963 | PRJEB87630 | M.A.Clements 6907 (CANB) | CANB: Clements, M.A. 6907 |
| *Bulbophyllum ambrosia* (Hance) Schltr. 1 | N/A | _ | NC_088536 | N/A | N/A |
| *Bulbophyllum ambrosia* (Hance) Schltr. 2 | N/A | CNS_G08282 | PRJEB87630 | Orchid Research Group 2674, B.J Wallace (CANB) | CANB: 999463.1 |
| *Bulbophyllum amplebracteatum* Teijsm. & Binn. | *Lepidorhiza* | CNS_G08270 | PRJEB87630 | Orchid Research Group 7744, J.Wright (CANB) | CANB: 1000626.1 |
| *Bulbophyllum amplebracteatum* subsp. *carunculatum* (Garay, Hamer & Siegerist) J.J.Verm. & P.O'Byrne | *Lepidorhiza* | CNS_G08712 | PRJEB87630 | Orchid Research Group 7685 (CANB) | CANB: ORG 7685 |
| *Bulbophyllum andreeae* A.D.Hawkes | *Intervallatae* | CNS_G07593 | PRJEB87630 | M.A.Clements 6490 (CANB) | CANB: Clements, M.A. 6490 |
| *Bulbophyllum ankylochele* J.J.Verm. | *Peltopus* | CNS_G08301 | PRJEB87630 | T.M.Reeve 879 (CANB) | CANB: 238834.1 |
| *Bulbophyllum annandalei* Ridl. | *Cirrhopetalum* | CNS_G07246 | PRJEB87630 | Orchid Research Group 4801, M.Harrison (CANB) | CANB: 670922.1 |
| *Bulbophyllum antenniferum* (Lindl.) Rchb.f. | *Hyalosema* | CNS_G05294 | PRJEB87630 | T.M.Reeve 638 (CANB) | CBG: 7904522.1 |
| *Bulbophyllum apodum* Hook.f. 1 | *Stachysanthes* | CNS_G07217 | PRJEB87630 | B.J.Wallace 024/91 (CANB) | CANB: B.J.Wallace 024/91 |
| *Bulbophyllum apodum* Hook.f. 2 | *Stachysanthes* | CNS_G07219 | PRJEB87630 | M.A.Clements s.n (CANB) | CANB: M.A.Clements s.n |
| *Bulbophyllum apodum* Hook.f. 3 | *Stachysanthes* | HTCG_2054 | PRJEB87630 | M.A.Clements 13191 (CANB) | CANB: M.A.Clements 13191 |
| *Bulbophyllum arfakianum* Kraenzl. | *Hyalosema* | CNS_G05266 | PRJEB87630 | Orchid Research Group 7092 (CANB) | CANB: 1000277.1 |
| *Bulbophyllum argyropus* (Endl.) Rchb.f. | *Adelopetalum* | CNS_G03566 | ERR13142759 | Orchid Research Group 7052, D.Banks (CANB) | CANB: ORG 7052 |
| *Bulbophyllum aristilabre* J.J.Sm. | *Codonosiphon* | CNS_G07618 | PRJEB87630 | Orchid Research Group 6264, G.Backhouse (CANB) | CANB: 1000236.1 |
| *Bulbophyllum ascochilum* J.J.Verm. | *Codonosiphon* | CNS_G07503 | PRJEB87630 | T.M.Reeve 610 (CANB) | CANB: T.M.Reeve 610 |
| *Bulbophyllum aundense* Ormerod | *Fruticicola* | CNS_G07546 | PRJEB87630 | M.M.J.VanBalgooy 508 (CANB) | CANB: 183164.1 |
| *Bulbophyllum ayuthayense* J.J.Verm., Schuit. & de Vogel | *Drymoda* | CNS_G08300 | PRJEB87630 | M.A.Clements 12919 (CANB) | CANB: M.A.Clements 12919 |
| *Bulbophyllum baileyi* F.Muell. 1 | *Stenochilus* | CNS_G01572 | PRJEB87630 | D.L.Jones 8817 (CANB) | CBG: 9220103.1 |
| *Bulbophyllum baileyi* F.Muell. 2 | *Stenochilus* | CNS_G01282 | PRJEB87630 | K.Schulte 77 (CNS) | CNS: G01282 |
| *Bulbophyllum baileyi* F.Muell. 3 | *Stenochilus* | CNS_G08714 | PRJEB87630 | D.L.Jones 18707c (CANB) | CANB: 630122.1 |
| *Bulbophyllum baladeanum* J.J.Sm. 1 | *Hoplandra* | HTCG_2488 | PRJEB87630 | M.A.Clements 5721 (CANB) | CBG: 8916364.1 |
| *Bulbophyllum baladeanum* J.J.Sm. 2 | *Hoplandra* | HTCG_2518 | PRJEB87630 | M.A.Clements 5721 (CANB) | CBG: 8916364.1 |
| *Bulbophyllum baladeanum* J.J.Sm. 3 | *Hoplandra* | HTCG_2430 | PRJEB87630 | P.D.Ziesing 120 (CANB) | CANB: P.D.Ziesing 120 |
| *Bulbophyllum baladeanum* J.J.Sm. 4 | *Hoplandra* | HTCG_2534 | PRJEB87630 | P.D.Ziesing 120 (CANB) | CANB: P.D.Ziesing 120 |
| *Bulbophyllum balgooiji* J.J.Verm., Schuit. & de Vogel | *Pedilochilus* | CNS_G07542 | PRJEB87630 | R.D.Hoogland 5668 (CANB) | CANB: R.D.Hoogland 5668 |
| *Bulbophyllum bandischii* Garay, Hamer & Siegerist | *Hyalosema* | HTCG_2058 | PRJEB87630 | M.A.Clements 13095 (CANB) | CANB: M.A.Clements 13095 |
| *Bulbophyllum barbigerum* Lindl. 1 | *Ptiloglossum* | HTCG_1933 | PRJEB87630 | Orchid Reseach Group7325, P.Spence (CANB) | CANB: 1000647.1 |
| *Bulbophyllum barbigerum* Lindl. 2 | *Ptiloglossum* | CNS_G08700 | PRJEB87630 | M.A.Clements s.n (CNS) | CNS: CNS_G08700 |
| *Bulbophyllum baronii* Ridl. 1 | *Ploiarium* | CNS_G06040 | ERR13142808 | Orchid Research Group 6242 (CANB) | CANB: ORG 6242 |
| *Bulbophyllum baronii* Ridl. 2 | *Ploiarium* | CNS_G07275 | PRJEB87630 | Orchid Reseach Group 6242, D.Smedley (CANB) | CANB: ORG 6242 |
| *Bulbophyllum basisetum* J.J.Sm. | *Lepidorhiza* | HTCG_1936 | PRJEB87630 | Orchid Research Group 7657, Z.Groeneveld (CANB) | CANB: ORG 7657 |
| *Bulbophyllum beccarii* Rchb.f. | *Beccariana* | CNS_G07559 | ERR13142822 | Orchid Research Group 7678 (CANB) | CANB: ORG 7678 |

| | | | | | |
|---|---|---|---|---|---|
| *Bulbophyllum betchei* F.Muell. | *Fruticicola* | CNS_G05227 | PRJEB87630 | P.D.Ziesing 147 (CANB) | CBG: 9217816.1 |
| *Bulbophyllum biantennatum* Schltr. | *Hyalosema* | CNS_G07527 | PRJEB87630 | P.Spence SH 913 (CANB) | CBG: 8001185.1 |
| *Bulbophyllum bicolor* Lindl. | *Cirrhopetalum* | CNS_G07280 | PRJEB87630 | Orchid Reseach Group 7655, Z.Groeneveld (CANB) | CANB: ORG 7655 |
| *Bulbophyllum blepharistes* Rchb.f. | *Blepharistes* | CNS_G08305 | PRJEB87630 | M.A.Clements 8635 (CANB) | CANB: 620067.1 |
| *Bulbophyllum boonjee* B.Gray & D.L.Jones | *Adelopetalum* | CNS_G07175 | ERR13142812 | B.Gray 9761 (CANB) | CNS: 145935.1 |
| *Bulbophyllum bowkettiae* F.M.Bailey | *Polymeres* | CNS_G01618 | PRJEB87630 | B.Gray 8697 (CANB) | CANB: 596002.1 |
| *Bulbophyllum bracteatum* (Fitzg.) F.M.Bailey | *Adelopetalum* | CNS_G01553 | ERR13142752 | M.Harrison CNS048 (CANB) | CNS: G01553 |
| *Bulbophyllum brassii* J.J.Verm. | *Peltopus* | CNS_G07557 | PRJEB87630 | R.Pullen 5191 (CANB) | CANB: R.Pullen 5191 |
| *Bulbophyllum brevilabium* Schltr. | *Papulipetalum* | CNS_G07650 | PRJEB87630 | Orchid Research Group 7747, M.Harrison (CANB) | CANB: 1000744.1 |
| *Bulbophyllum bulliferum* J.J.Sm. | *Lepanthanthe* | CNS_G07555 | PRJEB87630 | S.Rose 112 (CANB) | CANB: 1000887.2 |
| *Bulbophyllum cambodianum* (Christenson) J.J.Verm., Schuit. & de Vogel | *Trias* | CNS_G05275 | ERR13142787 | R.Cowen 3077 (CANB) | CANB: 1001020.1 |
| *Bulbophyllum caudatum* Lindl. | *Desmosanthes* | CNS_G05252 | PRJEB87630 | Mark Clements s.n. (CNS) | CNS: CNS_G05252 |
| *Bulbophyllum cauliflorum* Hook.f. | *Desmosanthes* | CNS_G05243 | ERR13142776 | B.J.Wallace 020/91 (CANB) | CANB: 1000154.1 |
| *Bulbophyllum cheiri* Lindl. | *Stenochilus* | CNS_G08716 | PRJEB87630 | Orchid Research Group 7770 (CANB) | CANB: ORG 7770 |
| *Bulbophyllum chloranthum* Schltr. | *Macrouris* | CNS_G05256 | PRJEB87630 | A.N.Millar NGF23566 (CANB) | BRI: AQ0072947 |
| *Bulbophyllum chrysendetum* Ames 1 | *Desmosanthes* | CNS_G05245 | PRJEB87630 | Orchid Research Group 4017, J.Cootes (CANB) | CANB: ORG 4017 |
| *Bulbophyllum chrysendetum* Ames 2 | *Desmosanthes* | CNS_G07260 | PRJEB87630 | Orchid Research Group 4017, J.Cootes (CANB) | CANB: ORG 4017 |
| *Bulbophyllum ciliatum* (Blume) Lindl. | *Monanthaparva* | CNS_G05450 | PRJEB87630 | Orchid Research Group 4316, B.J Wallace (CANB) | CANB: 925052.1 |
| *Bulbophyllum clandestinum* Lindl. 1 | *Oxysepala* | CNS_G07307 | ERR13142786 | T.M.Reeve 735 (CANB) | CBG: 8905730.1 |
| *Bulbophyllum clandestinum* Lindl. 2 | *Oxysepala* | CNS_G05272 | ERR13142816 | Orchid Research Group 3080, R. Cowan (CANB) | CANB: ORG 3080 |
| *Bulbophyllum cleistogamum* Ridl. | *Intervallatae* | CNS_G07577 | PRJEB87630 | Orchid Research Group 7668 (CANB) | CANB: ORG 7668 |
| *Bulbophyllum collettii* King & Pantl. | *Cirrhopetaloides* | CNS_G07249 | PRJEB87630 | Orchid Research Group 2872, J.Ryan (CANB) | CANB: 680503.3 |
| *Bulbophyllum colliferum* J.J.Sm. | *Pelma* | CNS_G07352 | PRJEB87630 | T.M.Reeve 615 (CANB) | CBG: 9003614.1 |
| *Bulbophyllum comberi* J.J.Verm. | *Monanthaparva* | CNS_G05239 | PRJEB87630 | B.Wallace 0009/94 (CANB) | CBG: 9404907.5 |
| *Bulbophyllum comosum* Collett & Hemsl. | *Lemniscata* | CNS_G08291 | PRJEB87630 | Orchid Research Group 8440, Z.Groeneveld (CANB) | CANB: 998856.1 |
| *Bulbophyllum concinnum* Hook.f. | *Desmosanthes* | CNS_G01626 | PRJEB87630 | Orchid Research Group 4017, J.Cootes (CANB) | CANB: ORG 4017 |
| *Bulbophyllum contortisepalum* J.J.Sm. | *Hoplandra* | CNS_G07349 | PRJEB87630 | T.M.Reeve 4923 (CANB) | CBG: 9409388.1 |
| *Bulbophyllum cootesii* M.A.Clem. | *Lepidorhiza* | CNS_G05234 | PRJEB87630 | Orchid Research Group 7671 (CANB) | CANB: ORG 7671 |
| *Bulbophyllum corythium* N.Hallé | *Adelopetalum* | HTCG_2360 | PRJEB87630 | M.A.Clements 11238 (CANB) | CANB: 998252.1 |
| *Bulbophyllum crassipes* Hook.f. 1 | *Racemosae* | _ | NC_088537 | N/A | N/A |
| *Bulbophyllum crassipes* Hook.f. 2 | *Racemosae* | HTCG_1939 | PRJEB87630 | Orchid Research Group 7832, M.Harrison (CANB) | CANB: 999465.1 |
| *Bulbophyllum cruciatum* J.J.Sm. 1 | *Codonosiphon* | CNS_G01056 | ERR13142747 | J.M.Taylor 2464 (CANB) | CBG: 8600516.1 |
| *Bulbophyllum cruciatum* J.J.Sm. 2 | *Codonosiphon* | CNS_G07649 | PRJEB87630 | M.A.Clements 12374 (CANB) | CANB: 953855.1 |
| *Bulbophyllum cruentum* Garay, Hamer & Siegerist | *Macrobulbon* | CNS_G06039 | ERR13142807 | Orchid Research Group 5524 (CANB) | CANB: ORG 5524 |
| *Bulbophyllum cryptophoranthus* Garay | *Lepidorhiza* | HTCG_1975 | PRJEB87630 | Orchid Research Group 7972, J.Cootes (CANB) | CANB: ORG 7972 |
| *Bulbophyllum cuniculiforme* J.J.Sm. | *Codonosiphon* | CNS_G07466 | PRJEB87630 | T.M.Reeve 5702 (CANB) | CBG: 9409389.1 |
| *Bulbophyllum cyatheicola* (P.Royen) J.J.Verm., Schuit. & de Vogel | *Pedilochilus* | CNS_G07295 | PRJEB87630 | D.N.McVean, L.K.Wade ANU 7225 (CANB) | CANB: D.N.McVean, L.K.Wade ANU 7225 |
| *Bulbophyllum dayanum* Rchb.f. | *Acrochaene* | CNS_G07568 | PRJEB87630 | Orchid Research Group 7682 (CANB) | CANB: ORG 7682 |
| *Bulbophyllum dearei* W.Watson 1 | *Sestochilus* | HTCG_1949 | PRJEB87630 | Orchid Research Group 7864 (CANB) | CANB: ORG 7864 |
| *Bulbophyllum dearei* W.Watson 2 | *Sestochilus* | HTCG_1950 | PRJEB87630 | Orchid Research Group 7865 (CANB) | CANB: ORG 7865 |
| *Bulbophyllum delitescens* Hance | *Macrostelydia* | CNS_G08329 | PRJEB87630 | A.Hu KFBG 25 (KFBG) | KFBG: A.Hu KFBG 25 |
| *Bulbophyllum depressum* King & Pantl. | *Monanthaparva* | CNS_G07279 | PRJEB87630 | Orchid Reseach Group 7661 (CANB) | CANB: 948502.1 |
| *Bulbophyllum desmotrichoides* Schltr. | *Imitatores* | CNS_G07468 | PRJEB87630 | M.A.Clements 6131 (CANB) | CBG: 9010123.1 |
| *Bulbophyllum dichotomum* J.J.Sm. | *Macrouris* | HTCG_1903 | PRJEB87630 | L.A.Craven 282 (CANB) | BRI: AQ0072957 |
| *Bulbophyllum dictyoneuron* Schltr. | *Polymeres* | HTCG_1907 | PRJEB87630 | T.M.Reeve 4964 (CANB) | CBG: 9217538.1 |
| *Bulbophyllum digoelense* J.J.Sm. | *Intervallatae* | CNS_G05276 | ERR13142788 | Orchid Research Group 6246 (CANB) | CANB: ORG 6246 |

| | | | | | |
|---|---|---|---|---|---|
| *Bulbophyllum dischidiifolium* J.J.Sm. | *Monanthes* | CNS_G05278 | ERR13142789 | C.E.Carr 17087 (CANB) | CANB: 114163.1 |
| *Bulbophyllum disciflorum* Rolfe | *Trias* | CNS_G08317 | PRJEB87630 | A.Hu KFBG 13 (KFBG) | KFBG: A.Hu KFBG 13 |
| *Bulbophyllum discolor* Schltr. | *Peltopus* | CNS_G07596 | PRJEB87630 | R.D.Hoogland 9414 (CANB) | CANB: 144166.1 |
| *Bulbophyllum dissitiflorum* Seidenf. | *Racemosae* | CNS_G05203 | PRJEB87630 | Orchid Research Group 6338 (CANB) | CANB: 924957.3 |
| *Bulbophyllum djamuense* Schltr. | *Fruticicola* | HTCG_1922 | PRJEB87630 | S.Rose 86 (CANB) | CANB: 1000823.1 |
| *Bulbophyllum echinolabium* J.J.Sm. | *Lepidorhiza* | CNS_G05237 | ERR13142774 | G.McCraith 165 (CANB) | CANB: 1000610.1 |
| *Bulbophyllum echinulus* Seidenf. | *Desmosanthes* | CNS_G08306 | PRJEB87630 | A.Hu KFBG 2 (KFBG) | KFBG: A.Hu KFBG 2 |
| *Bulbophyllum eciliatum* Schltr. | *Monanthes* | CNS_G05217 | PRJEB87630 | M.A.Clements 5569B (CANB) | CANB: M.A.Clements 5569B |
| *Bulbophyllum ecornutum* (J.J.Sm.) J.J.Sm. | *Beccariana* | HTCG_1952 | PRJEB87630 | Orchid Research Group 7868 (CANB) | CANB: ORG 7868 |
| *Bulbophyllum elassoglossum* Siegerist | *Polymeres* | CNS_G05456 | ERR13142799 | M.A.Clements 11502 (CANB) | CANB: 1000195.1 |
| *Bulbophyllum elegantius* Schltr. | *Monanthes* | HTCG_1915 | PRJEB87630 | M.A.Clements 7057 (CANB) | CBG: 9013305.1 |
| *Bulbophyllum elephantinum* J.J.Sm. | *Hyalosema* | CNS_G07526 | PRJEB87630 | T.M.Reeve 106 (CANB) | CANB: T.M.Reeve 106 |
| *Bulbophyllum elevatopunctatum* J.J.Sm. 1 | *Beccariana* | CNS_G07187 | PRJEB87630 | Orchid Research Group 7047, J.Wright (CANB) | CANB: 948896.1 |
| *Bulbophyllum elevatopunctatum* J.J.Sm. 2 | *Beccariana* | HTCG_1932 | PRJEB87630 | Orchid Research Group 7047, J.Wright (CANB) | CANB: 948896.1 |
| *Bulbophyllum elisae* (F.Muell.) Benth. | *Adelopetalum* | CNS_G01548 | PRJEB87630 | L.Simpson 498 (CNS) | CNS: 154147.1 |
| *Bulbophyllum emarginatum* (Finet) J.J.Sm. | *Emarginatae* | CNS_G08705 | PRJEB87630 | A.Hu KFBG 19 (KFBG) | KFBG: A.Hu KFBG 19 |
| *Bulbophyllum epiphytum* Barb.Rodr. | *Micranthae* | _ | NC_048486 | N/A | N/A |
| *Bulbophyllum ericssonii* Kraenzl. 1 | *Beccariana* | HTCG_1931 | PRJEB87630 | Orchid Research Group 7177 (CANB) | CANB: ORG 7177 |
| *Bulbophyllum ericssonii* Kraenzl. 2 | *Beccariana* | HTCG_1957 | PRJEB87630 | Orchid Research Group 7873 (CANB) | CANB: ORG 7873 |
| *Bulbophyllum erinaceum* Schltr. | *Lepanthanthe* | CNS_G07306 | PRJEB87630 | T.M.Reeve 4382 (CANB) | CBG: 9201871.1 |
| *Bulbophyllum erythrosema* J.J.Verm. | *Epibulbon* | CNS_G07331 | PRJEB87630 | R.D.Hoogland 6255 (CANB) | CANB: 230642.1 |
| *Bulbophyllum evasum* T.E.Hunt & Rupp | *Brachystachyae* | CNS_G01247 | ERR13142749 | D.L.Jones 4352 (CANB) | CBG: 8913156.1 |
| *Bulbophyllum exaltatum* Lindl. | *Didactyle* | _ | NC_048480 | N/A | N/A |
| *Bulbophyllum exiguum* F.Muell. | *Adelopetalum* | CNS_G07178 | PRJEB87630 | D.L.Jones 19231 (CANB) | CANB: 667983.1 |
| *Bulbophyllum facetum* Garay, Hamer & Siegerist | *Sestochilus* | HTCG_1972 | PRJEB87630 | Orchid Research Group 7969, J.Cootes (CANB) | CANB: ORG 7969 |
| *Bulbophyllum falcatum* (Lindl.) Rchb.f. | *Megaclinium* | CNS_G07648 | PRJEB87630 | Orchid Research Group 7765 (CANB) | CANB: ORG 7765 |
| *Bulbophyllum farreri* (W.W.Sm.) Seidenf. | *Brachyantha* | _ | NC_088538 | N/A | N/A |
| *Bulbophyllum fascinator* (Rolfe) Rolfe | *Cirrhopetaloides* | CNS_G08685 | PRJEB87630 | G.M.McCraith 144 (CANB) | CANB: 1000671.1 |
| *Bulbophyllum fenixii* Ames | *Polymeres* | CNS_G05201 | PRJEB87630 | Orchid Research Group 6234 (CANB) | CANB: ORG 6234 |
| *Bulbophyllum flammuliferum* Ridl. 1 | *Desmosanthes* | _ | NC_084215 | N/A | N/A |
| *Bulbophyllum flammuliferum* Ridl. 2 | *Desmosanthes* | CNS_G08265 | PRJEB87630 | Orchid Research Group 7767 (CANB) | CANB: ORG 7767 |
| *Bulbophyllum flavescens* Lindl. | *Stachysanthes* | CNS_G05280 | ERR13142790 | M.A.Clements 11548 (CANB) | CANB: 1000851.1 |
| *Bulbophyllum fletcherianum* J.G.Fowler | *Macrobulbon* | CNS_G08711 | PRJEB87630 | M.A.Clements 12868 (CANB) | CANB: 1000632.2 |
| *Bulbophyllum foetidum* Schltr. | *Beccariana* | CNS_G05873 | ERR13142801 | M.A.Clements 6431 (CANB) | CANB: 1001008.1 |
| *Bulbophyllum fonsflorum* J.J.Verm. | *Macrouris* | CNS_G05257 | PRJEB87630 | M.A.Clements 9599 (CANB) | CANB: 1000662.1 |
| *Bulbophyllum forrestii* Seidenf. | *Rhytionanthos* | CNS_G06037 | ERR13142806 | Orchid Research Group 1271, R.Cowan (CANB) | CANB: 999500.1 |
| *Bulbophyllum fractiflexum* J.J.Sm. | *Pelma* | CNS_G07515 | PRJEB87630 | M.A.Clements 6340 (CANB) | CBG: 9012723.4 |
| *Bulbophyllum fritillariiflorum* J.J.Sm. | *Hyalosema* | CNS_G05264 | PRJEB87630 | M.A.Clements 13212 (CANB) | CANB: M.A.Clements 13212 |
| *Bulbophyllum frostii* Summerh. 1 | *Cirrhopetaloides* | CNS_G08307 | PRJEB87630 | A.Hu KFBG 3 (KFBG) | KFBG: A.Hu KFBG 3 |
| *Bulbophyllum frostii* Summerh. 2 | *Cirrhopetaloides* | HTCG_1978 | PRJEB87630 | Orchid Research Group 7973, J.Cootes (CANB) | CANB: ORG 7973 |
| *Bulbophyllum fruticicola* Schltr. | *Epibulbon* | CNS_G07481 | ERR13142820 | T.M.Reeve 3372 (CANB) | CBG: 9000963.1 |
| *Bulbophyllum gadgarrense* Rupp | *Oxysepala* | CNS_G01602 | PRJEB87630 | M.Harrison CM18042013-3 (CNS) | CNS: CNS_G01602 |
| *Bulbophyllum gerlandianum* Kraenzl. | *Stenochilus* | CNS_G08299 | PRJEB87630 | M.A.Clements 12895 (CANB) | CANB: 1000880.1 |
| *Bulbophyllum gibbosum* (Blume) Lindl. | *Stachysanthes* | CNS_G05279 | PRJEB87630 | M.A.Clements 10638 (CANB) | CANB: 1000852.1 |
| *Bulbophyllum gracillimum* Rolfe | *Ephippium* | CNS_G01627 | PRJEB87630 | A.R.Field ARF3512 (CNS) | CNS: A.R.Field ARF3512 |
| *Bulbophyllum grandiflorum* Blume 1 | *Hyalosema* | CNS_G07282 | PRJEB87630 | Orchid Reseach Group 7652, Z.Groeneveld (CANB) | CANB: 947107.1 |
| *Bulbophyllum grandiflorum* Blume 2 | *Hyalosema* | CNS_G08288 | PRJEB87630 | M.A.Clements 12887 (CANB) | CANB: 1000293.1 |
| *Bulbophyllum grandiflorum* Blume 3 | *Hyalosema* | CNS_G08690 | PRJEB87630 | Orchid Research Group 7419 (CANB) | CANB: 905108.1 |

| | | | | | |
|---|---|---|---|---|---|
| *Bulbophyllum grandimesense* B.Gray & D.L.Jones | *Oxysepala* | CNS_G01590 | PRJEB87630 | M.Harrison CM18042013-1 (CNS) | CNS: CNS_G01590 |
| *Bulbophyllum granulosum* Barb.Rodr. | *Napellii* | _ | NC_048481 | N/A | N/A |
| *Bulbophyllum guttulatum* (Hook.f.) N.P.Balakr. | *Brachyantha* | CNS_G05250 | ERR13142779 | Orchid Research Group 5587 (CANB) | CANB: ORG 5587 |
| *Bulbophyllum gyaloglossum* J.J.Verm. | *Pelma* | CNS_G07541 | PRJEB87630 | H.W.Millar 22554 (BRI) | BRI: AQ0358101 |
| *Bulbophyllum gymnopus* Hook.f. | *Altisceptrum* | CNS_G08327 | PRJEB87630 | A.Hu KFBG 23 (KFBG) | KFBG: A.Hu KFBG 23 |
| *Bulbophyllum hahlianum* Schltr. | *Stenochilus* | CNS_G05269 | PRJEB87630 | J.Wrigley ANBG 770881 (CANB) | CANB: 1000866.1 |
| *Bulbophyllum hamelinii* W.Watson | *Alcistachys* | HTCG_1951 | PRJEB87630 | Orchid Research Group 7866 (CANB) | CANB: ORG 7866 |
| *Bulbophyllum haniffii* Carr | *Epicrianthes* | CNS_G07566 | ERR13142824 | Orchid Research Group 7677 (CANB) | CANB: ORG 7677 |
| *Bulbophyllum helenae* (Kuntze) J.J.Sm. | *Rhytionanthos* | CNS_G08308 | PRJEB87630 | A.Hu KFBG 4 (KFBG) | KFBG: A.Hu KFBG 4 |
| *Bulbophyllum hexarhopalon* Schltr. | *Epicrianthes* | CNS_G00340 | PRJEB87630 | M.A.Clements 11214 (CANB) | CBG: 9212989 |
| *Bulbophyllum hians* Schltr. | *Papulipetalum* | CNS_G07653 | PRJEB87630 | Orchid Research Group 7741, M.Harrison (CANB) | CANB: 1000740.1 |
| *Bulbophyllum hirtum* (Sm.) Lindl. ex Wall. | *Lemniscata* | CNS_G08309 | PRJEB87630 | A.Hu KFBG 5 (KFBG) | KFBG: A.Hu KFBG 5 |
| *Bulbophyllum hirundinis* (Gagnep.) Seidenf. 1 | *Cirrhopetalum* | CNS_G08269 | PRJEB87630 | Orchid Research Group 7754 (CANB) | CANB: 1000133.1 |
| *Bulbophyllum hirundinis* (Gagnep.) Seidenf. 2 | *Cirrhopetalum* | CNS_G08713 | PRJEB87630 | Orchid Research Group 7754 (CANB) | CANB: 1000133.1 |
| *Bulbophyllum hystricinum* Schltr. | *Fruticicola* | HTCG_1921 | PRJEB87630 | S.Rose 85 (CANB) | CANB: 1000824.1 |
| *Bulbophyllum icteranthum* Schltr. | *Fruticicola* | CNS_G07355 | PRJEB87630 | T.M.Reeve 4942 (CANB) | CBG: 9217539.1 |
| *Bulbophyllum inconspicuum* Maxim. | *Cirrhopetalum* | _ | NC_046811 | N/A | N/A |
| *Bulbophyllum infundibuliforme* J.J.Sm. | *Hymenobractea* | CNS_G07305 | ERR13142815 | M.A.Clements 6616 (CANB) | CBG: 9008257.1 |
| *Bulbophyllum intricatum* Seidenf. | *Racemosae* | CNS_G05202 | ERR13142765 | G.McCraith 83 (CANB) | CANB: 999468.3 |
| *Bulbophyllum jimcootesii* (Cabactulan, M.Leon & R.B.Pimentel) R.Rice | *Epicrianthes* | HTCG_1974 | PRJEB87630 | Orchid Research Group 7971, J.Cootes (CANB) | CANB: ORG 7971 |
| *Bulbophyllum johnsonii* T.E.Hunt 1 | *Polymeres* | CNS_G00538 | PRJEB87630 | C.D.Kilgour 150 (CNS) | CNS: 144417.1 |
| *Bulbophyllum johnsonii* T.E.Hunt 2 | *Polymeres* | HTCG_1943 | PRJEB87630 | M.A.Clements 13313 (CANB) | CANB: M.A.Clements 13313 |
| *Bulbophyllum kauloense* Schltr. | *Oxysepala* | CNS_G07614 | PRJEB87630 | T.M.Reeve 6537 (CANB) | CBG: 9201823.1 |
| *Bulbophyllum keekee* N.Halle | *Minutissima* | CNS_G07329 | ERR13142818 | M.A.Clements 5675 (CANB) | CBG: 8916317.1 |
| *Bulbophyllum klabatense* Schltr. | *Lepidorhiza* | CNS_G07579 | PRJEB87630 | Orchid Research Group 7666 (CANB) | CANB: ORG 7666 |
| *Bulbophyllum klabatense* subsp. *sulawesii* (Garay, Hamer & Siegerist) J.J.Verm. & P.O'Byrne | *Lepidorhiza* | CNS_G00643 | PRJEB87630 | Orchid Research Group 6343, J.Wright (CANB) | CANB: 924959.2 |
| *Bulbophyllum kubahense* J.J.Verm. & A.Lamb | *Beccariana* | CNS_G07575 | PRJEB87630 | Orchid Research Group 7675 (CANB) | CANB: ORG 7675 |
| *Bulbophyllum lageniforme* F.M.Bailey | *Adelopetalum* | CNS_G03707 | ERR13142760 | L.Simpson LS167B (CNS) | CNS: CNS_G03707 |
| *Bulbophyllum lamingtonense* D.L.Jones | *Oxysepala* | CNS_G01013 | PRJEB87630 | Orchid Research Group 5199 (CANB) | CANB: 737251.1 |
| *Bulbophyllum lasianthum* Lindl. | *Beccariana* | CNS_G08274 | PRJEB87630 | Orchid Research Group 7768 (CANB) | CANB: ORG 7768 |
| *Bulbophyllum lasiochilum* Parish & Rchb.f. | *Cirrhopetaloides* | CNS_G07204 | ERR13142813 | M.A.Clements 7343 (CANB) | CANB: 737416.1 |
| *Bulbophyllum lasioglossum* Rolfe ex Ames | *Hirtula* | HTCG_1918 | PRJEB87630 | R.G.Phillips 1141 (CANB) | CBG: 9214366.1 |
| *Bulbophyllum laxiflorum* (Blume) Lindl. 1 | *Desmosanthes* | CNS_G08706 | PRJEB87630 | A.Hu KFBG 6 (KFBG) | KFBG: A.Hu KFBG 6 |
| *Bulbophyllum laxiflorum* (Blume) Lindl. 2 | *Desmosanthes* | CNS_G08686 | PRJEB87630 | M.A.Clements 11547 (CANB) | CANB: 1000162.1 |
| *Bulbophyllum lemniscatoides* Rolfe | *Lemniscata* | CNS_G05204 | ERR13142766 | Orchid Research Group 7153 (CANB) | CANB: 946207.1 |
| *Bulbophyllum lepidum* (Blume) J.J.Sm. | *Ephippium* | CNS_G07252 | PRJEB87630 | G.McCraith 25 (CANB) | CANB: McCraith 25 |
| *Bulbophyllum leptoleucum* Schltr. | *Pelma* | CNS_G07470 | PRJEB87630 | A.N.Millar 18668 (CANB) | CANB: 134847.1 |
| *Bulbophyllum leucothyrsus* Schltr. | *Pelma* | HTCG_2051 | PRJEB87630 | T.M.Reeve 643 (CANB) | CBG: 7904541.3 |
| *Bulbophyllum levanae* Ames | *Lepidorhiza* | CNS_G05235 | PRJEB87630 | G.McCraith 130 (CANB) | CANB: 1000613.1 |
| *Bulbophyllum levatii* Kraenzl. | *Macrouris* | CNS_G07590 | PRJEB87630 | T.M.Reeve 1054 (CANB) | CBG: 8905746.1 |
| *Bulbophyllum levyae* Garay, Hamer & Siegerist | *Lepidorhiza* | CNS_G07565 | ERR13142823 | Orchid Research Group 7674 (CANB) | CANB: ORG 7674 |
| *Bulbophyllum lewisense* B.Gray & D.L.Jones | *Oxysepala* | CNS_G01607 | PRJEB87630 | M.Harrison CM18042013-4 (CNS) | CNS: CNS_G01607 |
| *Bulbophyllum lilacinum* Ridl. | *Racemosae* | CNS_G01934 | PRJEB87630 | Orchid Research Group 7692, M.Harrison (CANB) | CANB: 948510.1 |
| *Bulbophyllum lilianae* Rendle | *Adelopetalum* | CNS_G07754 | ERR13142833 | L.Simpson 337E (CNS) | CNS: 154142.1 |
| *Bulbophyllum lindleyanum* Griff. | *Hirtula* | CNS_G06035 | ERR13142805 | Orchid Research Group 6237 (CANB) | CANB: 1000268.1 |
| *Bulbophyllum lineolatum* Schltr. | *Brachypus* | CNS_G05262 | ERR13142783 | M.A.Clements 9580 (CANB) | CANB: 1000700.1 |

| | | | | | |
|---|---|---|---|---|---|
| *Bulbophyllum lingulatum* Rendle | *Adelopetalum* | CNS_G06046 | ERR13142811 | M.A.Clements 7959 (CANB) | CBG: 9213395.3 |
| *Bulbophyllum lobbii* Lindl. 1 | *Sestochilus* | CNS_G05229 | ERR13142771 | Orchid Research Group 5166 (CANB) | CANB: 740419.3 |
| *Bulbophyllum lobbii* Lindl. 2 | *Sestochilus* | HTCG_1946 | PRJEB87630 | Orchid Research Group 7861 (CANB) | CANB: ORG 7861 |
| *Bulbophyllum lobbii* Lindl. 3 | *Sestochilus* | HTCG_1948 | PRJEB87630 | Orchid Research Group 7863 (CANB) | CANB: ORG 7863 |
| *Bulbophyllum lobbii* subsp. *siamense* | *Sestochilus* | HTCG_1954 | PRJEB87630 | Orchid Research Group 7870 (CANB) | CANB: ORG 7870 |
| *Bulbophyllum longibrachiatum* Z.H.Tsi | *Cirrhopetalum* | CNS_G07245 | PRJEB87630 | Orchid Research Group 6263, B.J.Wallace (CANB) | CANB: 1000137.1 |
| *Bulbophyllum longiflorum* Thouars | *Cirrhopetalum* | CNS_G06022 | ERR13142802 | Orchid Research Group 7758, M. Harrison (CANB) | CANB: 1000939.1 |
| *Bulbophyllum longisepalum* Rolfe | *Hyalosema* | CNS_G08287 | PRJEB87630 | M.A.Clements 12890 (CANB) | CANB: 1000294.1 |
| *Bulbophyllum lophoglottis* (Guillaumin) N.Hallé | *Monanthes* | HTCG_2523 | PRJEB87630 | P.D.Ziesing 150 (CANB) | CANB: P.D.Ziesing 150 |
| *Bulbophyllum lyriforme* J.J.Verm. & P.O'Byrne | *Stenochilus* | HTCG_1947 | PRJEB87630 | Orchid Research Group 7862 (CANB) | CANB: ORG 7862 |
| *Bulbophyllum macphersonii* Rupp 1 | *Monanthes* | HTCG_2048 | PRJEB87630 | M.A.Clements 12889 (CANB) | CANB: 1000804.1 |
| *Bulbophyllum macphersonii* Rupp 2 | *Monanthes* | HTCG_2061 | PRJEB87630 | M.A.Clements 12889 (CANB) | CANB: 1000804.1 |
| *Bulbophyllum macraei* (Lindl.) Rchb.f. | *Macrostelydia* | CNS_G06045 | ERR13142810 | M.A.Clements 12421 (CANB) | CANB: 956310.1 |
| *Bulbophyllum macranthum* Lindl. | *Stenochilus* | CNS_G05452 | ERR13142797 | CBG 770881 (CANB) | CBG: 770881 |
| *Bulbophyllum macrorhopalon* Schltr. | *Epicrianthes* | CNS_G07489 | PRJEB87630 | T.M.Reeve 1093 (CANB) | CBG: 8905758.1 |
| *Bulbophyllum macrourum* Schltr. | *Macrouris* | CNS_G07336 | PRJEB87630 | K.Paijmans 718 (CANB) | CANB: 219238.1 |
| *Bulbophyllum makoyanum* (Rchb.f.) Ridl. | *Ephippium* | CNS_G08294 | PRJEB87630 | M.A.Clements 12899 (CANB) | CANB: 1000120.1 |
| *Bulbophyllum mandibulare* Rchb.f. | *Lepidorhiza* | CNS_G07645 | PRJEB87630 | Orchid Research Group 7769 (CANB) | CANB: 1000621.1 |
| *Bulbophyllum maquilingense* Ames & Quisumb. | *Polymeres* | CNS_G05216 | PRJEB87630 | M.A.Clements 1978 (CANB) | CANB: M.A.Clements 1978 |
| *Bulbophyllum marknaivei* Cootes, R.Boos & M.Leon | *Brachystachyae* | CNS_G08293 | PRJEB87630 | M.A.Clements 12861 (CANB) | CANB: 999482.1 |
| *Bulbophyllum mastersianum* (Rolfe) J.J.Sm. | *Ephippium* | HTCG_1940 | PRJEB87630 | Orchid Research Group 7855, K.Turner (CANB) | CANB: ORG 7855 |
| *Bulbophyllum maxillare* Rchb.f. | *Polymeres* | CNS_G01299 | ERR13142750 | J.M.Taylor 284 (CANB) | CANB: J.M.Taylor 284 |
| *Bulbophyllum maximum* (Lindl.) Rchb.f. | *Megaclinium* | CNS_G05289 | ERR13142791 | Orchid Research Group 6250 (CANB) | CANB: ORG 6250 |
| *Bulbophyllum mearnsii* Ames 1 | *Lepidorhiza* | CNS_G07561 | PRJEB87630 | Orchid Research Group 7686 (CANB) | CANB: 1000619.1 |
| *Bulbophyllum mearnsii* Ames 2 | *Lepidorhiza* | HTCG_1935 | PRJEB87630 | Orchid Research Group 7686 (CANB) | CANB: 1000619.2 |
| *Bulbophyllum medusae* Rchb.f. | *Ephippium* | CNS_G07573 | ERR13142827 | Orchid Research Group 7684 (CANB) | CANB: ORG 7684 |
| *Bulbophyllum membranaceum* Teijsm. & Binn. | *Monanthaparva* | CNS_G05449 | PRJEB87630 | M.A.Clements 11550 (CANB) | CANB: 1000842.1 |
| *Bulbophyllum membranifolium* Hook.f. | *Beccariana* | HTCG_1962 | PRJEB87630 | M.A.Clements 13092 (CANB) | CANB: M.A.Clements 13092 |
| *Bulbophyllum mentosum* Barb.Rodr. | *Micranthae* | _ | NC_048482 | N/A | N/A |
| *Bulbophyllum meridense* Rchb.f. | *Didactyle* | HTCG_1959 | PRJEB87630 | Orchid Research Group 7879 (CANB) | CANB: ORG 7879 |
| *Bulbophyllum mimiense* Schltr. | *Epibulbon* | CNS_G05213 | PRJEB87630 | M.A.Clements s.n. (CNS) | CNS: CNS_G05213 |
| *Bulbophyllum minutissimum* (F.Muell.) F.Muell. | *Minutissima* | CNS_G01587 | ERR13142753 | M.Harrison s.n (CANB) | CANB: 1001570.1 |
| *Bulbophyllum mirum* J.J.Sm. | *Plumata* | CNS_G06031 | ERR13142804 | Orchid Research Group, J. Cootes 5501 (CANB) | CANB: 778009.1 |
| *Bulbophyllum mucronatum* (Blume) Lindl. 1 | *Monanthaparva* | CNS_G05440 | ERR13142794 | M.A.Clements 12379 (CANB) | CANB: 953860.1 |
| *Bulbophyllum mucronatum* (Blume) Lindl. 2 | *Monanthaparva* | HTCG_2052 | PRJEB87630 | M.A.Clements 13190 (CANB) | CANB: M.A.Clements 13190 |
| *Bulbophyllum muricatum* J.J.Sm. | *Monosepalum* | HTCG_1925 | PRJEB87630 | S.Rose 66 (CANB) | CANB: S.Rose 66 |
| *Bulbophyllum nasica* Schltr. | *Polymeres* | CNS_G07289 | PRJEB87630 | M.A.Clements 6337 (CANB) | CBG: 9012720.1 |
| *Bulbophyllum nasseri* Garay | *Lepidorhiza* | HTCG_1960 | PRJEB87630 | M.A.Clements 13090 (CANB) | CANB: M.A.Clements 13090 |
| *Bulbophyllum nasutum* Rchb.f. | *Trias* | CNS_G08318 | PRJEB87630 | A.Hu KFBG 14 (KFBG) | KFBG: A.Hu KFBG 14 |
| *Bulbophyllum nematopodum* F.Muell. 1 | *Brachypus* | CNS_G07257 | PRJEB87630 | CBG 721721 (CANB) | CANB: CBG 721721 |
| *Bulbophyllum nematopodum* F.Muell. 2 | *Brachypus* | HTCG_1977 | PRJEB87630 | K.Schulte 193a (CNS) | CNS: 136996.1 |
| *Bulbophyllum newportii* Rolfe | *Adelopetalum* | CNS_G03754 | ERR13142761 | L.Simpson 155E (CNS) | CNS: 154141.1 |
| *Bulbophyllum ngoyense* Schltr. 1 | *Hoplandra* | CNS_G05218 | PRJEB87630 | M.A.Clements 11179 (CANB) | CANB: 949284.1 |
| *Bulbophyllum ngoyense* Schltr. 2 | *Hoplandra* | HTCG_2504 | PRJEB87630 | P.D.Ziesing 170 (CANB) | CANB: P.D.Ziesing 170 |
| *Bulbophyllum nigricans* (Aver.) J.J.Verm., Schuit. & de Vogel | *Ione* | CNS_G08324 | PRJEB87630 | A.Hu KFBG 20 (KFBG) | KFBG: A.Hu KFBG 20 |
| *Bulbophyllum ningboense* G.Y.Li ex H.L.Lin & X.P.Li | *Cirrhopetalum* | _ | NC_064067 | N/A | N/A |
| *Bulbophyllum nummularioides* Schltr. | *Monanthes* | CNS_G07538 | PRJEB87630 | T.M.Reeve 5637 (CANB) | CBG: 9409407.1 |
| *Bulbophyllum nymphopolitanum* Kraenzl. | *Lepidorhiza* | CNS_G07651 | PRJEB87630 | Orchid Research Group 4019 (CANB) | CANB: ORG 6250 |

| | | | | | |
|---|---|---|---|---|---|
| *Bulbophyllum oblongum* (Lindl.) Rchb.f. | *Trias* | CNS_G08319 | PRJEB87630 | A.Hu KFBG 15 (KFBG) | KFBG: A.Hu KFBG 15 |
| *Bulbophyllum obovatum* (J.J.Sm.) J.J.Verm., Schuit. & de Vogel | *Pedilochilus* | HTCG_1924 | PRJEB87630 | S.Rose 37 (CANB) | CANB: 1000753.1 |
| *Bulbophyllum occlusum* Ridl. | *Alcistachys* | CNS_G07291 | ERR13142814 | Orchid Reseach Group 3376, D.Smedley (CANB) | CANB: 681129.1 |
| *Bulbophyllum ocellatum* Cootes & M.A.Clem. | *Polymeres* | CNS_G05293 | ERR13142792 | M.A.Clements 11603 (CANB) | CANB: 1000213.1 |
| *Bulbophyllum ochroleucum* Schltr. | *Uncifera* | CNS_G07320 | PRJEB87630 | R.D.Hoogland 9210 (CANB) | CANB: 143860.2 |
| *Bulbophyllum odoratissimum* Lindl. ex Wall. | *Desmosanthes* | CNS_G07258 | PRJEB87630 | B.J.Wallace 023/91 (CANB) | CANB: ORG 023/91 |
| *Bulbophyllum odoratum* (Blume) Lindl. | *Stachysanthes* | CNS_G05283 | PRJEB87630 | M.A.Clements 12045 (CANB) | CANB: 906120.1 |
| *Bulbophyllum olivinum* J.J.Sm. | *Macrouris* | CNS_G08719 | PRJEB87630 | L.K.Wade ANU 7585 (CANB) | CANB: 179584.1 |
| *Bulbophyllum orectopetalum* Garay, Hamer & Siegerist | *Sestochilus* | CNS_G07225 | PRJEB87630 | Orchid Research Group 4802, M.Harrison (CANB) | CANB: ORG 4802 |
| *Bulbophyllum oreomene* J.J.Verm., Schuit. & de Vogel | *Pedilochilus* | CNS_G07517 | ERR13142821 | G.Hope ANU 10850 (CANB) | CANB: G.Hope ANU 10850 |
| *Bulbophyllum orientale* Seidenf. | *Racemosae* | CNS_G05224 | PRJEB87630 | Orchid Research Group 5526 (CANB) | CANB: ORG 5526 |
| *Bulbophyllum orthosepalum* J.J.Verm. | *Macrobulbon* | HTCG_1976 | PRJEB87630 | Orchid Research Group 7854, M.Harrison (CANB) | CANB: 1000633.1 |
| *Bulbophyllum pachypus* Schltr. | *Pachychlamys* | CNS_G07740 | ERR13142832 | M.A.Clements 12774 (CANB) | CANB: Clements, M.A. 12774 |
| *Bulbophyllum pahudii* (de Vriese) Rchb.f. 1 | *Beccariana* | HTCG_1956 | PRJEB87630 | Orchid Research Group 7872, K.Turner (CANB) | CANB: OORG 7872 |
| *Bulbophyllum pahudii* (de Vriese) Rchb.f. 2 | *Beccariana* | HTCG_1961 | PRJEB87630 | M.A.Clements 13091 (CANB) | CANB: M.A.Clements 13091 |
| *Bulbophyllum pardalotum* Garay, Hamer & Siegerist | *Polymeres* | CNS_G05226 | PRJEB87630 | M.A.Clements 11503 (CANB) | CANB: ORG 11503 |
| *Bulbophyllum parviflorum* C.S.P.Parish & Rchb.f. | *Hirtula* | CNS_G08703 | PRJEB87630 | A.Hu KFBG 24 (KFBG) | KFBG: A.Hu KFBG 24 |
| *Bulbophyllum patens* King ex Hook.f. | *Stenochilus* | CNS_G05230 | PRJEB87630 | Orchid Research Group 5527 (CANB) | CANB: ORG 5527 |
| *Bulbophyllum pecten-veneris* (Gagnep.) Seidenf. | *Cirrhopetalum* | CNS_G08687 | PRJEB87630 | M.A.Clements 12338 (CANB) | CANB: M.A.Clements 12338 |
| *Bulbophyllum penicillium* C.S.P.Parish & Rchb.f. | *Hirtula* | CNS_G08261 | PRJEB87630 | M.Barret s.n. (CANB) | CANB: M.Barret s.n. |
| *Bulbophyllum peninsulare* Seidenf. | *Racemosae* | HTCG_1970 | PRJEB87630 | M.A.Clements 13211 (CANB) | CANB: 949175.1 |
| *Bulbophyllum petrophilum* (P.Royen) J.J.Verm., Schuit. & de Vogel | *Pedilochilus* | CNS_G07520 | PRJEB87630 | R.Schodde 1765 (CANB) | CANB: 107149.1 |
| *Bulbophyllum phaeoglossum* Schltr. | *Brachypus* | CNS_G08675 | PRJEB87630 | T.M.Reeve 5020 (CANB) | CANB: 570961.1 |
| *Bulbophyllum phalaenopsis* J.J.Sm. | *Macrobulbon* | CNS_G05206 | PRJEB87630 | Orchid Research Group 6982 (CANB) | CANB: ORG 6982 |
| *Bulbophyllum phormion* J.J.Verm. | *Macrouris* | CNS_G07353 | PRJEB87630 | T.M.Reeve 6220 (CANB) | CBG: 9409416.1 |
| *Bulbophyllum pictum* C.S.P.Parish & Rchb.f. | *Trias* | CNS_G07643 | PRJEB87630 | Orchid Research Group 7773 (CANB) | CANB: ORG 7773 |
| *Bulbophyllum picturatum* Rchb.f. | *Cirrhopetalum* | CNS_G07243 | PRJEB87630 | M.A.Clements 8159 (CANB) | CANB: ORG 8159 |
| *Bulbophyllum piestobulbon* Schltr. | *Piestobulbon* | CNS_G08695 | PRJEB87630 | L.Craven 75 (CANB) | CANB: 144853.1 |
| *Bulbophyllum piundaundense* (P.Royen) J.J.Verm., Schuit. & de Vogel | *Pedilochilus* | CNS_G07516 | PRJEB87630 | D.McVean, L.K.Wade ANU 7182 (CANB) | CANB: ORG 7182 |
| *Bulbophyllum planibulbe* Ridl. | *Planibulbus* | CNS_G08311 | PRJEB87630 | A.Hu KFBG 7 (KFBG) | KFBG: A.Hu KFBG 7 |
| *Bulbophyllum pleurothallidanthum* Garay 1 | *Desmosanthes* | CNS_G07567 | ERR13142825 | Orchid Research Group 7681 (CANB) | CANB: 1000159.1 |
| *Bulbophyllum pleurothallidanthum* Garay 2 | *Desmosanthes* | CNS_G01937 | PRJEB87630 | Orchid Research Group 7681 (CANB) | CANB: 1000159.1 |
| *Bulbophyllum plumosum* (Barb.Rodr.) Cogn. | *Xiphizusa* | _ | NC_048479 | N/A | N/A |
| *Bulbophyllum plumula* Schltr. | *Monanthes* | CNS_G07354 | PRJEB87630 | T.M.Reeve 109 (CANB) | CANB: 1000807.1 |
| *Bulbophyllum polliculosum* Seidenf. | *Lemniscata* | CNS_G08289 | PRJEB87630 | M.A.Clements 12872 (CANB) | CANB: OORGp 12872 |
| *Bulbophyllum polyblepharon* Schltr. | *Monanthes* | CNS_G07490 | PRJEB87630 | T.M.Reeve 2847 (CANB) | CANB: 570960.1 |
| *Bulbophyllum prianganense* J.J.Sm. | *Stachysanthes* | CNS_G08710 | PRJEB87630 | Orchid Research Group 7762 (CANB) | CANB: 1000158.1 |
| *Bulbophyllum propinquum* Kraenzl. | *Racemosae* | CNS_G07576 | ERR13142828 | Orchid Research Group 7667 (CANB) | CANB: ORG 7667 |
| *Bulbophyllum ptilotes* Schltr. | *Monanthes* | CNS_G07348 | PRJEB87630 | T.M.Reeve 1016 (CANB) | CBG: 8905761.1 |
| *Bulbophyllum pulvinatum* Schltr. | *Fruticicola* | CNS_G07473 | PRJEB87630 | T.Reeve 2973 (CANB) | CBG: 9000956.1 |
| *Bulbophyllum punamense* Schltr. 1 | *Monanthes* | CNS_G07500 | PRJEB87630 | M.A.Clements 6838 (CANB) | CBG: 9013086.3 |
| *Bulbophyllum punamense* Schltr. 2 | *Monanthes* | HTCG_1916 | PRJEB87630 | M.A.Clements 6838 (CANB) | CBG: 9013086.4 |
| *Bulbophyllum punamense* Schltr. 3 | *Monanthes* | HTCG_1927 | PRJEB87630 | M.A.Clements 9320 (CANB) | CANB: 1000815.1 |
| *Bulbophyllum pungens* Schltr. | *Oxysepala* | HTCG_1967 | PRJEB87630 | M.A.Clements 13205 (CANB) | CANB: 1000712.1 |
| *Bulbophyllum putidum* (Teijsm. & Binn.) J.J.Sm. | *Cirrhopetaloides* | CNS_G05872 | ERR13142800 | D.Banks 9199 (CANB) | CANB: 618183.2 |
| *Bulbophyllum pygmaeum* Lindl. | *Minutissima* | CNS_G04599 | ERR13142764 | B.P.J.Molloy 062/98 (CHR) | CHR: 531794 |

| | | | | | |
|---|---|---|---|---|---|
| *Bulbophyllum quadrangulare* J.J.Sm. | *Polymeres* | HTCG_1917 | PRJEB87630 | M.A.Clements 6336 (CANB) | CANB: M.A.Clements 6336 |
| *Bulbophyllum quadrifalciculatum* J.J.Sm. | *Monanthes* | HTCG_1926 | PRJEB87630 | S.Rose 74 (CANB) | CANB: 1000809.1 |
| *Bulbophyllum radicans* F.M.Bailey | *Fruticicola* | CNS_5228 | ERR13142772 | K.Schulte 149 (CNS) | CNS: 136975.1 |
| *Bulbophyllum refractilingue* J.J.Sm. | *Beccariana* | CNS_G05220 | PRJEB87630 | Orchid Research Group 7175, K.Turner (CANB) | CANB: ORG 7175 |
| *Bulbophyllum regnellii* Rchb.f. | *Napellii* | _ | NC_048483 | N/A | N/A |
| *Bulbophyllum renipetalum* Schltr. | *Monanthes* | CNS_G07475 | PRJEB87630 | T.Reeve 887 (CANB) | CBG: 8905766.1 |
| *Bulbophyllum repens* Griff. | *Brachystachyae* | CNS_G08312 | PRJEB87630 | A.Hu KFBG 8 (KFBG) | KFBG: A.Hu KFBG 8 |
| *Bulbophyllum reptans* Lindl. ex Wall. | *Reptantia* | CNS_G08313 | PRJEB87630 | A.Hu KFBG 9 (KFBG) | KFBG: A.Hu KFBG 9 |
| *Bulbophyllum restrepia* (Ridl.) Ridl. | *Hoplandra* | CNS_G08302 | PRJEB87630 | M.A.Clements s.n (CNS) | CNS: CNS_G08302 |
| *Bulbophyllum reticulatum* Bateman ex Hook.f. | *Beccariana* | HTCG_2043 | PRJEB87630 | M.A.Clements 13186 (CANB) | CANB: M.A.Clements 13186 |
| *Bulbophyllum retusiusculum* Rchb.f. | *Brachyantha* | CNS_G05253 | PRJEB87630 | G.McCraith 1270 (CANB) | CANB: 620121.2 |
| *Bulbophyllum rhodoglossum* Schltr. | *Epibulbon* | CNS_G07310 | ERR13142817 | T.M.Reeve 995 (CANB) | CBG: 8905750.1 |
| *Bulbophyllum rivulare* Schltr. | *Papulipetalum* | HTCG_1908 | PRJEB87630 | T.M.Reeve 983 (CANB) | CBG: 9003615.1 |
| *Bulbophyllum romyi* B.Thoms | *Cirrhopetalum* | HTCG_1953 | PRJEB87630 | Orchid Research Group 7869 (CANB) | CANB: ORG 7869 |
| *Bulbophyllum roseopictum* J.J.Verm., Schuit. & de Vogel | *Ione* | CNS_G03907 | ERR13142763 | M.A.Clements 12085 (CANB) | CANB: M.A.Clements 12085 |
| *Bulbophyllum rothschildianum* (O'Brien) J.J.Sm. | *Cirrhopetalum* | CNS_G05285 | PRJEB87630 | M.A.Clements 12181 (CANB) | CANB: 906169.2 |
| *Bulbophyllum roxburghii* Rchb.f. | *Cirrhopetaloides* | CNS_G07250 | PRJEB87630 | Orchid Research Group 6199, K.Groeneveld (CANB) | CANB: 1000111.1 |
| *Bulbophyllum rubipetalum* P.Royen | *Polymeres* | CNS_G07603 | PRJEB87630 | J.M.B.Smith ANU 15436 (CANB) | CANB: 240710.1 |
| *Bulbophyllum rufinum* Rchb.f. | *Racemosae* | CNS_G06395 | PRJEB87630 | M.A.Clements 12431 (CANB) | CANB: 956320.1 |
| *Bulbophyllum rugosum* Ridl. | *Beccariana* | HTCG_1964 | PRJEB87630 | M.A.Clements 13098 (CANB) | CANB: M.A.Clements 13098 |
| *Bulbophyllum saccoglossum* J.J.Verm., Schuit. & de Vogel | *Codonosiphon* | HTCG_1911 | PRJEB87630 | T.M.Reeve 6622 (CANB) | CBG: 9201831.1 |
| *Bulbophyllum saltatorium* Lindl. | *Ptiloglossum* | CNS_G08285 | PRJEB87630 | M.A.Clements 12892 (CANB) | CANB: 1000821.1 |
| *Bulbophyllum sandersonii* (Hook.f.) Rchb.f. | *Megaclinium* | CNS_G07229 | PRJEB87630 | Royal Botanic Garden Sydney 821846, P.Metcalfe | RBGS: P.Metcalfe 821846 |
| *Bulbophyllum sanguineopunctatum* Seidenf. & A.D.Kerr 1 | *Cirrhopetalum* | CNS_G07278 | PRJEB87630 | Orchid Reseach Group 7385, Z.Groeneveld (CANB) | CANB: 1000127.1 |
| *Bulbophyllum sanguineopunctatum* Seidenf. & A.D.Kerr 2 | *Cirrhopetalum* | HTCG_2046 | PRJEB87630 | M.A.Clements 13187 (CANB) | CANB: M.A.Clements 13187 |
| *Bulbophyllum santosii* Ames | *Beccariana* | CNS_G07639 | PRJEB87630 | M.A.Clements 12803 (CANB) | CANB: 949513.1 |
| *Bulbophyllum sauguetiense* Schltr. 1 | *Papulipetalum* | CNS_G05258 | ERR13142781 | M.A.Clements 7235 (CANB) | CBG: 9013484.1 |
| *Bulbophyllum sauguetiense* Schltr. 2 | *Papulipetalum* | HTCG_1965 | PRJEB87630 | M.A.Clements 13199 (CANB) | CANB: 1000737.1 |
| *Bulbophyllum savaiense* Schltr. | *Pelma* | CNS_G07483 | PRJEB87630 | R.D.Hoogland 9367 (CANB) | BRI: AQ0075037 |
| *Bulbophyllum scaberulum* (Rolfe) Bolus | *Megaclinium* | CNS_G07652 | PRJEB87630 | Orchid Research Group 6173, MH0179 (CANB) | CANB: ORG 6173 |
| *Bulbophyllum scabratum* Rchb.f. | *Cirrhopetalum* | CNS_G08330 | PRJEB87630 | A.Hu KFBG 26 (KFBG) | KFBG: A.Hu KFBG 26 |
| *Bulbophyllum schinzianum* Kraenzl. ex De Wild. & T.Durand 1 | *Ptiloglossum* | CNS_G01642 | ERR13142754 | M.A.Clements 11554 (CANB) | CANB: 1000648.1 |
| *Bulbophyllum schinzianum* Kraenzl. ex De Wild. & T.Durand 2 | *Ptiloglossum* | CNS_G06038 | PRJEB87630 | M.A.Clements 11554 (CANB) | CANB: 1000648.1 |
| *Bulbophyllum schmidii* Garay | *Hyalosema* | CNS_G07496 | PRJEB87630 | Orchid Research Group 3189 (CANB) | CANB: 680538.1 |
| *Bulbophyllum scopa* J.J.Verm. | *Macrouris* | CNS_G07511 | PRJEB87630 | T.M.Reeve 1106 (CANB) | CBG: 9201824.1 |
| *Bulbophyllum scorpio* J.J.Verm. | *Gongorodes* | CNS_G08283 | PRJEB87630 | M.A.Clements 12882 (CANB) | CANB: ORG 12882 |
| *Bulbophyllum shanicum* King & Pantl. | *Lemniscata* | _ | NC_088539 | N/A | N/A |
| *Bulbophyllum shepherdii* (F.Muell.) Rchb.f. | *Oxysepala* | CNS_G07185 | PRJEB87630 | D.L.Jones 19293 (CANB) | CANB: 672895.1 |
| *Bulbophyllum sibuyanense* Ames | *Ephippium* | HTCG_1945 | PRJEB87630 | Orchid Research Group 7860 (CANB) | CANB: ORG 7860 |
| *Bulbophyllum sigaldiae* Guillaumin | *Acrochaene* | CNS_G07581 | ERR13142830 | Orchid Research Group 7679 (CANB) | CANB: ORG 7679 |
| *Bulbophyllum simile* Schltr. | *Pelma* | CNS_G07549 | PRJEB87630 | R.D.Hoogland 9336 (CANB) | CANB: 144190.2 |
| *Bulbophyllum skeatianum* Ridl. | *Cirrhopetalum* | CNS_G08275 | PRJEB87630 | G.McCraith 126 (CANB) | CANB: 1000116.1 |
| *Bulbophyllum smitinandii* Seidenf. & Thorut | *Sestochilus* | HTCG_1955 | PRJEB87630 | Orchid Research Group 7871 (CANB) | CANB: ORG 7871 |
| *Bulbophyllum* sp. 1 | N/A | HTCG_1958 | PRJEB87630 | Orchid Research Group 7874 (CANB) | CANB: ORG 7874 |
| *Bulbophyllum* sp. 2 | N/A | HTCG_1901 | PRJEB87630 | T.G.Hartley 12136 (CANB) | CANB: 150685.1 |
| *Bulbophyllum* sp. 3 | N/A | CNS_G08701 | PRJEB87630 | M.A.Clements s.n (CNS) | CNS: CNS_G08701 |
| *Bulbophyllum* sp. 4 | N/A | CNS_G07274 | PRJEB87630 | Orchid Reseach Group 7649, Z.Groeneveld (CANB) | CANB: 947104.2 |
| *Bulbophyllum* sp. 5 | N/A | CNS_G08699 | PRJEB87630 | M.A.Clements s.n (CNS) | CNS: CNS_G08699 |

| | | | | | |
|---|---|---|---|---|---|
| *Bulbophyllum* sp. 6 | N/A | CNS_G08698 | PRJEB87630 | M.A.Clements s.n (CNS) | CNS: CNS_G08698 |
| *Bulbophyllum* sp. 7 | N/A | CNS_G07646 | PRJEB87630 | Orchid Research Group 7753, M. Harrison (CANB) | CANB: 1000717.1 |
| *Bulbophyllum* sp. 8 | N/A | CNS_G05263 | PRJEB87630 | M.A.Clements 1996 (CANB) | CANB: ORG 1996 |
| *Bulbophyllum* sp. 9 | N/A | CNS_G07592 | PRJEB87630 | L.Craven 2661 (CANB) | CANB: L.Craven 2661 |
| *Bulbophyllum* sp. 10 | N/A | CNS_G08677 | PRJEB87630 | T.M.Reeve 610 (CANB) | CANB: ORG 610 |
| *Bulbophyllum* sp. 11 | N/A | CNS_G05274 | PRJEB87630 | M.A.Clements 6847 (CANB) | CBG: 9013095.1 |
| *Bulbophyllum* sp. 12 | N/A | CNS_G07600 | PRJEB87630 | T.M.Reeve 1108 (CANB) | CBG: 8905757.1 |
| *Bulbophyllum spathulatum* (Rolfe ex E.Cooper) Seidenf. | *Brachyantha* | CNS_G08267 | PRJEB87630 | Orchid Research Group 4124 (CANB) | CANB: 1000117.1 |
| *Bulbophyllum speciosum* Schltr. | *Codonosiphon* | CNS_G07277 | PRJEB87630 | Orchid Reseach Group 7649, Z.Groeneveld (CANB) | CANB: 947104.2 |
| *Bulbophyllum spodotriche* J.J.Verm. & P.O'Byrne | *Epicrianthes* | CNS_G08303 | PRJEB87630 | M.A.Clements s.n (CNS) | CNS: CNS_G08303 |
| *Bulbophyllum steyermarkii* Foldats | *Furvescens* | _ | NC_048484 | N/A | N/A |
| *Bulbophyllum subpatulum* J.J.Verm. | *Macrouris* | CNS_G07597 | PRJEB87630 | W.Vink 16570 (CANB) | CANB: 156503.1 |
| *Bulbophyllum sukhakulii* Seidenf. | *Lemniscata* | CNS_G08314 | PRJEB87630 | A.Hu KFBG 10 (KFBG) | KFBG: A.Hu KFBG 10 |
| *Bulbophyllum taiwanense* (Fukuy.) K.Nakaj. | *Cirrhopetaloides* | CNS_G07281 | PRJEB87630 | Orchid Reseach Group 7314, Z.Groeneveld (CANB) | CANB: 1000131.1 |
| *Bulbophyllum tentaculiferum* Schltr. | *Monanthes* | CNS_G07495 | PRJEB87630 | T.M.Reeve 691 (CANB) | CANB: ORG 691 |
| *Bulbophyllum thaiorum* J.J.Sm. | *Brachyantha* | CNS_G07582 | PRJEB87630 | Orchid Research Group 7680 (CANB) | CANB: ORG 7680 |
| *Bulbophyllum thiurum* J.J.Verm. & P.O'Byrne | *Plumata* | HTCG_1941 | PRJEB87630 | Orchid Research Group 7856 (CANB) | CANB: ORG 7856 |
| *Bulbophyllum tianguii* K.Y.Lang & D.Luo | *Brachyantha* | _ | NC_060701 | N/A | N/A |
| *Bulbophyllum tollenoniferum* J.J.Sm. | *Stenochilus* | CNS_G08691 | PRJEB87630 | Orchid Research Group 7420 (CANB) | CANB: 1000878.1 |
| *Bulbophyllum tortuosum* (Blume) Lindl. | *Monanthes* | CNS_G08304 | PRJEB87630 | Orchid Reseach Group 4136, D.Banks (CANB) | CANB: ORG 4136 |
| *Bulbophyllum trachypus* Schltr. | *Lepanthanthe* | CNS_G07594 | PRJEB87630 | M.A.Clements 7073 (CANB) | CBG: 9013321.1 |
| *Bulbophyllum treschii* Jenny 1 | *Plumata* | CNS_G07578 | ERR13142829 | Z.Groeneveld (CANB) | CANB: 1000785.1 |
| *Bulbophyllum treschii* Jenny 2 | *Plumata* | HTCG_1938 | PRJEB87630 | Z.Groeneveld (CANB) | CANB: 1000785.1 |
| *Bulbophyllum triaristella* Schltr. | *Peltopus* | CNS_G01202 | ERR13142748 | Orchid Research Group 5581, B.J Wallace (CANB) | CANB: 1000693.1 |
| *Bulbophyllum tridentatum* Kraenzl. | *Racemosae* | CNS_G07261 | PRJEB87630 | M.A.Clements 8157 (CANB) | CANB: 620341.1 |
| *Bulbophyllum trifilum* J.J.Sm. | *Macrouris* | CNS_G07476 | PRJEB87630 | LAE 59489 (CANB) | BRI: AQ0359538 |
| *Bulbophyllum tripudians* C.S.P.Parish & Rchb.f. | *Tripudianthes* | CNS_G05249 | ERR13142778 | Orchid Research Group 3081, R.Cowan (CANB) | CANB: ORG 3081 |
| *Bulbophyllum triste* (Rolfe) Schltr. 1 | *Lemniscata* | _ | NC_088540 | N/A | N/A |
| *Bulbophyllum triste* Rchb.f. 2 | *Lemniscata* | CNS_G08315 | PRJEB87630 | A.Hu KFBG 11 (KFBG) | KFBG: A.Hu KFBG 11 |
| *Bulbophyllum tuberculatum* Colenso | *Adelopetalum* | HTCG_1928 | PRJEB87630 | B.P.J.Molloy 112/99 (CHR) | CHR: 531772 |
| *Bulbophyllum umbellatum* Lindl. | *Brachyantha* | CNS_G08704 | PRJEB87630 | A.Hu KFBG 22 (CANB) | KFBG: A.Hu KFBG 22 |
| *Bulbophyllum unicaudatum* Schltr. | *Monanthes* | CNS_G07512 | PRJEB87630 | T.M.Reeve 865 (CANB) | CBG: 9000251.1 |
| *Bulbophyllum uniflorum* (Blume) Hassk. | *Beccariana* | CNS_G07571 | ERR13142826 | Orchid Research Group 7670 (CANB) | CANB: 1000839.1 |
| *Bulbophyllum unitubum* J.J.Sm. | *Hyalosema* | CNS_G07580 | PRJEB87630 | Orchid Research Group 7664 (CANB) | CANB: 948504.1 |
| *Bulbophyllum vaginatum* Rchb.f. | *Ephippium* | CNS_G07262 | PRJEB87630 | Orchid Research Group 3091 (CANB) | CANB: ORGp 3091 |
| *Bulbophyllum veldkampii* J.J.Verm. & P.O'Byrne | *Lepidorhiza* | HTCG_1942 | PRJEB87630 | Orchid Research Group 7875, K.Turner (CANB) | CANB: ORG 7875 |
| *Bulbophyllum verrucosum* (L.O.Williams) J.J.Verm., Schuit. & de Vogel | *Codonosiphon* | HTCG_1912 | PRJEB87630 | T.M.Reeve 947 (CANB) | CBG: 8905744.1 |
| *Bulbophyllum versteegii* J.J.Sm. | *Brachypus* | CNS_G07595 | PRJEB87630 | M.A.Clements 6755 (CANB) | CBG: 9011947.3 |
| *Bulbophyllum violaceolabellum* Seidenf. 1 | *Brachyantha* | HTCG_1971 | PRJEB87630 | Orchid Research Group 7967, J.Cootes (CANB) | CANB: ORG 7967 |
| *Bulbophyllum violaceolabellum* Seidenf. 2 | *Brachyantha* | _ | NC_088541 | N/A | N/A |
| *Bulbophyllum wadsworthii* Dockrill | *Oxysepala* | CNS_G01601 | PRJEB87630 | M.Harrison CM18042013-2 (CNS) | CNS: CNS_G01601 |
| *Bulbophyllum wakoi* Howcroft | *Beccariana* | CNS_G05222 | PRJEB87630 | Orchid Research Group 7176, K.Turner (CANB) | CANB: ORG 7176 |
| *Bulbophyllum wallichii* Rchb.f. | *Tripudianthes* | CNS_G08298 | PRJEB87630 | M.A.Clements 13000 (CANB) | CANB: M.A.Clements 13000 |
| *Bulbophyllum weddellii* (Lindl.) Rchb.f. | *Didactyle* | _ | NC_048485 | N/A | N/A |
| *Bulbophyllum weinthalii* R.S.Rogers | *Adelopetalum* | CNS_G06024 | ERR13142803 | M.T. Mathieson 730 (BRI) | BRI: AQ0792800 |
| *Bulbophyllum williamsii* A.D.Hawkes | *Polymeres* | CNS_G08292 | PRJEB87630 | M.A.Clements 12908 (CANB) | CANB: 906503.2 |
| *Bulbophyllum windsorense* B.Gray & D.L.Jones | *Oxysepala* | CNS_G01610 | PRJEB87630 | M.Harrison 2589 (CANB) | CANB: 680446.1 |

| | | | | | |
|---|---|---|---|---|---|
| *Bulbophyllum wolfei* B.Gray & D.L.Jones | *Polymeres* | CNS_G07176 | PRJEB87630 | B.Gray 9758 (CNS) | CNS: 145936.1 |
| *Bulbophyllum xylophyllum* C.S.P.Parish & Rchb.f. | *Brachystachyae* | CNS_G08316 | PRJEB87630 | A.Hu KFBG 12 (KFBG) | KFBG: A.Hu KFBG 12 |
| *Bulbophyllum zebrinum* J.J.Sm. | *Polymeres* | CNS_G07347 | PRJEB87630 | T.M.Reeve 3320 (CANB) | CBG: 9000960.1 |

**Herbarium acronyms:** BRI: Queensland Herbarium; CANB: Australian National Herbarium; CNS: Australian Tropical Herbarium; NSW: National Herbarium of New South Wales; KFBG: Kadoori Farm Botanic Gardens; RBGS: Royal Botanic Garden, Sydney

#### **Table S2.B:** Outgroups for phylogenetic analysis

| Species name | Orchid Subfamily | DNA number | Reference |
|---|---|---|---|
| *Aa paleacea* (Kunth) Rchb.f. | Orchidoideae | _ | Givnish et al. (2015) |
| *Acriopsis emarginata* D.L.Jones & M.A.Clem. | Epidendroideae | CNS_G00305 | ERR12947594 |
| *Agapanthus coddii* F.M.Leight | Outgroup | _ | NC_035971.1 |
| *Agave americana* L. | Outgroup | _ | NC 032053.1 |
| *Angraecum sesquipedale* Thouars | Epidendroideae | _ | Givnish et al. (2015) |
| *Anoectochilus emeiensis* K.Y.Lang | Orchidoideae | _ | NC_033895.1 |
| *Aphyllorchis montana* Rchb.f. | Epidendroideae | _ | NC_030703.1 |
| *Apostasia odorata* Blume | Apostasioideae | _ | NC_030722.1 |
| *Apostasia shenzhenica* Z.J.Liu & L.J.Chen | Apostasioideae | _ | NC_039812.1 |
| *Apostasia wallichii* R.Br. | Apostasioideae | _ | NC_036260.1 |
| *Arundina graminifolia* (D.Don) Hochr | Epidendroideae | _ | MN171408.1 |
| *Asparagus officinalis* L. | Outgroup | _ | NC 028422.1 |
| *Bletilla striata* Rchb.f. | Epidendroideae | _ | Serna-Sánchez et al. (2021) |
| *Brachionidium cruziae* L.O.Williams | Epidendroideae | _ | Serna-Sánchez et al. (2021) |
| *Calanthe aristulifera* Rchb.f. | Epidendroideae | _ | MN200378 |
| *Calanthe sylvatica* Lindl. | Epidendroideae | _ | NC_044633.1 |
| *Calopogon tuberosus* (L.) Britton, Sterns & Poggenb. | Epidendroideae | _ | Givnish et al. (2015) |
| *Calypso bulbosa* var. *occidentalis* (Holz.) Cockerell | Epidendroideae | _ | NC_040980.1 |
| *Cattleya liliputana* (Pabst) Van den Berg | Epidendroideae | _ | NC_032083.1 |
| *Cephalanthera damasonium* Druce | Epidendroideae | _ | NC_041179.1 |
| *Cephalanthera rubra* (L.) Rich. | Epidendroideae | _ | NC_041181.1 |
| *Changnienia amoena* S.S.Chien | Epidendroideae | _ | NC_045402.1 |
| *Cheirostylis chinensis* Rolfe | Orchidoideae | _ | MN641483.1 |
| *Codonorchis lessonii* Lindl. | Orchidoideae | _ | Givnish et al. (2015) |
| *Coelogyne flaccida* Lindl. | Epidendroideae | _ | Givnish et al. (2015) |
| *Corallorhiza striata* Lindl. | Epidendroideae | _ | NC 040978.1 |
| *Corallorhiza trifida* Châtel | Epidendroideae | _ | NC_025662.1 |
| *Cremastra appendiculata* (D.Don) Makino | Epidendroideae | _ | NC 037439.1 |
| *Cremastra unguiculata* Finet | Epidendroideae | _ | MN200381 |
| *Cymbidium canaliculatum* R.Br. | Epidendroideae | CNS_G00165 | ERR12947595 |
| *Cypripedium calceolus* L. | Cypripedioideae | _ | NC_045400.1 |
| *Cypripedium japonicum* Thunb. | Cypripedioideae | _ | NC_027227.1 |
| *Dendrobium kingianum* Bidwill ex Lindl. | Epidendroideae | _ | LC331062.1 |
| *Dendrobium moniliforme* 1 (L.) Sw. | Epidendroideae | _ | MN200384 |
| *Dendrobium moniliforme* 2 (L.) Sw. | Epidendroideae | CNS_G00882 | ERR13142745 |
| *Dipodium pandanum* F.M.Bailey | Epidendroideae | CNS_G01262 | ERR12947593 |
| *Dipodium roseum* D.L.Jones & M.A.Clem. | Epidendroideae | HTCG_1687 | ERR12947528 |
| *Elleanthus sodiroi* Schltr. | Epidendroideae | _ | NC_027266.1 |

| | | | |
|---|---|---|---|
| *Epipactis mairei* Schltr. | Epidendroideae | _ | NC_030705.1 |
| *Epipactis thunbergia* A.Gray | Epidendroideae | _ | MN200387 |
| *Epipactis veratrifolia* Boiss. & Hohen. | Epidendroideae | _ | NC_030708.1 |
| *Eria scabrilinguis* Lindl. | Epidendroideae | _ | MN477202.1 |
| *Eulophia bicallosa* (D.Don) P.F.Hunt & Summerh. | Epidendroideae | HTCG_1696 | ERR12947533 |
| *Eulophia nuda* Lindl. | Epidendroideae | HTCG_1697 | ERR12947534 |
| *Evotella carnosa* (Lindl.) J.C.Manning & Goldblatt | Orchidoideae | _ | Givnish et al. (2015) |
| *Gastrochilus japonicus* Schltr. | Epidendroideae | _ | NC_035833.1 |
| *Goodyera fumata* Thwaites | Orchidoideae | _ | NC_026773.1 |
| *Goodyera schlechtendaliana* Rchb.f. | Orchidoideae | _ | MK134679.1 |
| *Guarianthe aurantiaca* (Bateman) Dressler & W.E.Higgins | Epidendroideae | _ | Givnish et al. (2015) |
| *Gymnadenia conopsea* (L.) R.Br. | Orchidoideae | _ | MN200391 |
| *Habenaria arenaria* Lindl. | Orchidoideae | _ | Givnish et al. (2015) |
| *Habenaria ciliolaris* Kraenzl. | Orchidoideae | _ | MN495954.1 |
| *Hexalectris warnockii* Ames & Correll | Epidendroideae | _ | MH444822.1 |
| *Holcoglossum lingulatum* (Aver.) Aver. | Epidendroideae | _ | NC_041465.1 |
| *Holcoglossum subulifolium* (Rchb.f.) Christenson | Epidendroideae | _ | NC_041519.1 |
| *Iris sanguinea* Hornem | Outgroup | _ | NC_029227.1 |
| *Lilium pensylvanicum* Ker Gawl. | Outgroup | _ | NC_043876.1 |
| *Liparis auriculata* Blume ex Miq. | Epidendroideae | _ | MN200365 |
| *Ludisia discolor* (Ker Gawl.) Blume | Orchidoideae | _ | NC_030540.1 |
| *Masdevallia coccinea* Linden ex Lindl. | Epidendroideae | _ | NC_026541.1 |
| *Neottia listeroides* Lindl. | Epidendroideae | _ | NC_030713.1 |
| *Neottia ovata* Bluff & Fingerh. | Epidendroideae | _ | NC_030712.1 |
| *Nervilia crociformis* (Zoll. & Moritzi) Seidenf. | Epidendroideae | _ | Givnish et al. (2015) |
| *Neuwiedia zollingeri* var. *singapureana* (Wall. ex Baker) de Vogel | Apostasioideae | _ | LC199503.1 |
| *Oberonia japonica* (Maxim.) Makino | Epidendroideae | _ | NC 035832.1 |
| *Ophrys fusca* subsp. *iricolor* (Desf.) K.Richt. | Orchidoideae | _ | AP018716.1 |
| *Ophrys sphegodes* Mill. | Orchidoideae | _ | AP018717.1 |
| *Palmorchis pabstii* Veyret. | Epidendroideae | _ | NC_041190.1 |
| *Paphiopedilum armeniacum* S.C.Chen & F.Y.Liu | Cypripedioideae | _ | NC_026779.1 |
| *Paphiopedilum niveum* (Rchb.f.) Stein | Cypripedioideae | _ | NC_026776.1 |
| *Pelatantheria scolopendrifolia* (Makino) Aver. | Epidendroideae | _ | NC_035829.1 |
| *Phaius tankervilleae* (Banks) Blume | Epidendroideae | _ | Givnish et al. (2015) |
| *Phalaenopsis japonica* (Rchb.f.) Kocyan & Schuit. | Epidendroideae | _ | Givnish et al. (2015) |
| *Phalaenopsis pulcherrima* (Lindl.) J.J.Sm. | Epidendroideae | _ | MG459020.1 |
| *Phragmipedium longifolium* (Rchb.f. & Warsz.) Rolfe | Cypripedioideae | _ | NC_028149.1 |
| *Platanthera mandarinorum* Rchb.f. | Orchidoideae | _ | MN200370 |
| *Platystele aurea* Garay | Epidendroideae | _ | Serna-Sánchez et al. (2021) |
| *Pleione formosana* Hayata | Epidendroideae | _ | NC_042197.1 |
| *Pogonia ophioglossoides* (L.) Ker Gawl. | Vanilloideae | _ | Givnish et al. (2015) |
| *Ponerorchis gracilis* (Blume) X.H.Jin, Schuit. & W.T.Jin | Orchidoideae | _ | MN200376 |
| *Scaphosepalum antenniferum* Rolfe | Epidendroideae | _ | Serna-Sánchez et al. (2021) |
| *Sobralia mucronata* Ames & C.Schweinf. | Epidendroideae | _ | Givnish et al. (2015) |
| *Spiranthes sinensis* (Pers.) Ames | Orchidoideae | _ | MK936427.1 |
| *Teagueia* (Luer) Luer | Epidendroideae | _ | Serna-Sánchez et al. (2021) |
| *Thelymitra cyanea* (Lindl.) Benth. | Orchidoideae | _ | Givnish et al. (2015) |

| | | | |
|---|---|---|---|
| *Thrixspermum japonicum* Rchb.f. | Epidendroideae | _ | NC 035831.2 |
| *Triphora trianthophora* (Sw.) Rydb. | Epidendroideae | _ | Givnish et al. (2015) |
| *Tropidia polystachya* Ames | Epidendroideae | _ | Givnish et al. (2015) |
| *Vanda brunnea* Rchb.f. | Epidendroideae | _ | NC_041522.1 |
| *Vanda falcata* Beer | Epidendroideae | _ | NC_036372.1 |
| *Vanilla aphylla* Blume | Vanilloideae | _ | NC_035320.1 |
| *Vanilla planifolia* Andrews | Vanilloideae | _ | NC_026778.1 |
| *Vanilla pompona* Schiede | Vanilloideae | _ | NC_036809.1 |
| *Zootrophion hirtzii* Luer | Epidendroideae | _ | Serna-Sánchez et al. (2021) |

**Table S3:** Genes included in phylogenetic analysis

| Sample name | *acc* D | *atp* A | *atp* B | *atp* E | *atp* F | *atp* H | *atp* I | *ccs* A | *cem* A | *clp* P | infA | *mat* K | *pet* A | *pet* B | *pet* D | *pet* G | *pet* L | *pet* N | *psa* A | *psa* B | *psa* C | *psa* I | *psa* J | *psb* A | *psb* B | *psb* C | *psb* D | *psb* E | *psb* F | *psb* H | *psb* I | *psb* J | *psb* K | *psb* L | *psb* M | *psb* N | *psb* T | *psb* Z | *rbc* L | *rpl* 2 | *rpl* 06 | *rpl* 20 | *rpl* 22 | *rpl* 23 | *rpl* 32 | *rpl* 33 | *rpl* 36 | *rpo* A | *rpo* B | *rpo* C1 | *rpo* C2 | *rps* 2 | *rps* 3 | *rps* 7 | *rps* 8 | *rps* 11 | *rps* 12 | *rps* 14 | *rps* 15 | *rps* 16 | *rps* 18 | *ycf* 3 | *ycf* 4 |
|---|---|---|---|---|---|---|---|---|---|---|---|---|---|---|---|---|---|---|---|---|---|---|---|---|---|---|---|---|---|---|---|---|---|---|---|---|---|---|---|---|---|---|---|---|---|---|---|---|---|---|---|---|---|---|---|---|---|---|---|---|---|---|---|
| *Bulbophyllum absconditum* J.J.Sm. | 1 | 1 | 1 | 1 | 1 | 1 | 1 | 1 | 1 | 1 | 1 | 1 | 1 | 1 | 1 | 1 | 1 | 1 | 1 | 1 | 1 | 1 | 1 | 1 | 1 | 1 | 1 | 1 | 1 | 1 | 1 | 1 | 1 | 1 | 1 | 1 | 1 | 1 | 1 | 1 | 1 | 1 | 0 | 1 | 1 | 1 | 1 | 1 | 1 | 1 | 1 | 1 | 1 | 1 | 1 | 1 | 1 | 1 | 1 | 0 | 1 | 1 | 1 |
| *Bulbophyllum acuminatum* (Ridl.) Ridl. | 1 | 1 | 1 | 1 | 1 | 1 | 1 | 1 | 1 | 1 | 1 | 1 | 1 | 1 | 1 | 1 | 1 | 1 | 1 | 1 | 1 | 1 | 1 | 1 | 1 | 1 | 1 | 1 | 1 | 1 | 1 | 1 | 0 | 1 | 1 | 1 | 1 | 1 | 1 | 1 | 1 | 1 | 1 | 1 | 1 | 1 | 1 | 1 | 1 | 0 | 1 | 1 | 1 | 1 | 1 | 1 | 0 | 1 | 1 | 1 | 1 | 1 | 1 |
| *Bulbophyllum acutilingue* J.J.Sm. 1 | 1 | 1 | 1 | 1 | 1 | 1 | 1 | 1 | 1 | 1 | 1 | 1 | 1 | 1 | 1 | 1 | 1 | 1 | 0 | 1 | 1 | 1 | 1 | 1 | 1 | 1 | 1 | 1 | 1 | 1 | 1 | 1 | 1 | 1 | 1 | 1 | 1 | 1 | 1 | 1 | 1 | 1 | 1 | 1 | 1 | 1 | 1 | 1 | 1 | 1 | 1 | 1 | 1 | 1 | 1 | 1 | 1 | 0 | 1 | 1 | 1 | 1 | 1 |
| *Bulbophyllum acutilingue* J.J.Sm. 2 | 1 | 1 | 1 | 1 | 1 | 1 | 1 | 1 | 1 | 1 | 1 | 1 | 1 | 1 | 1 | 1 | 1 | 1 | 1 | 1 | 1 | 1 | 1 | 1 | 1 | 1 | 1 | 1 | 1 | 1 | 1 | 1 | 1 | 1 | 1 | 1 | 1 | 1 | 1 | 1 | 1 | 1 | 1 | 1 | 1 | 1 | 1 | 1 | 1 | 1 | 1 | 1 | 1 | 1 | 1 | 1 | 1 | 1 | 1 | 1 | 1 | 1 | 1 |
| *Bulbophyllum affine* Lindl. | 1 | 1 | 1 | 1 | 1 | 1 | 1 | 1 | 1 | 1 | 1 | 1 | 1 | 1 | 1 | 1 | 1 | 1 | 1 | 1 | 1 | 1 | 1 | 1 | 1 | 1 | 1 | 1 | 1 | 1 | 1 | 1 | 1 | 1 | 1 | 1 | 1 | 1 | 1 | 1 | 1 | 1 | 1 | 1 | 1 | 1 | 0 | 1 | 1 | 1 | 1 | 1 | 0 | 1 | 1 | 1 | 1 | 1 | 1 | 1 | 1 | 1 | 1 |
| *Bulbophyllum agastor* Garay, Hamer & Siegerist | 1 | 1 | 1 | 1 | 1 | 1 | 1 | 1 | 1 | 1 | 1 | 1 | 1 | 1 | 1 | 1 | 1 | 1 | 1 | 1 | 1 | 1 | 1 | 1 | 1 | 0 | 0 | 1 | 1 | 1 | 1 | 1 | 1 | 1 | 1 | 1 | 1 | 1 | 1 | 1 | 1 | 1 | 1 | 1 | 0 | 1 | 1 | 1 | 1 | 1 | 1 | 1 | 1 | 1 | 1 | 1 | 1 | 1 | 1 | 1 | 1 | 1 | 1 |
| *Bulbophyllum albociliatum* (Tang S.Liu & H.Y.Su) K.Nakaj. | 1 | 1 | 1 | 1 | 1 | 1 | 1 | 1 | 1 | 1 | 1 | 1 | 1 | 1 | 1 | 1 | 1 | 1 | 1 | 1 | 1 | 1 | 1 | 1 | 1 | 1 | 1 | 1 | 1 | 1 | 1 | 1 | 1 | 1 | 1 | 1 | 1 | 1 | 1 | 1 | 1 | 1 | 1 | 1 | 1 | 1 | 1 | 1 | 1 | 1 | 1 | 1 | 1 | 1 | 1 | 1 | 0 | 1 | 1 | 1 | 1 | 1 | 1 |
| *Bulbophyllum alkmaarense* J.J.Sm. | 1 | 1 | 1 | 1 | 1 | 1 | 1 | 1 | 1 | 1 | 1 | 1 | 1 | 1 | 1 | 1 | 1 | 1 | 1 | 1 | 1 | 1 | 1 | 1 | 1 | 1 | 1 | 1 | 1 | 1 | 1 | 1 | 1 | 1 | 1 | 1 | 1 | 1 | 1 | 1 | 1 | 1 | 0 | 1 | 1 | 1 | 1 | 1 | 1 | 1 | 1 | 1 | 1 | 1 | 1 | 1 | 1 | 1 | 1 | 1 | 1 | 1 | 1 |
| *Bulbophyllum alpinum* (P.Royen) J.J.Verm., Schuit. & de Vogel | 1 | 1 | 1 | 1 | 1 | 1 | 1 | 1 | 1 | 1 | 1 | 1 | 1 | 1 | 1 | 1 | 1 | 1 | 1 | 1 | 1 | 1 | 1 | 1 | 1 | 1 | 1 | 1 | 1 | 1 | 1 | 1 | 1 | 1 | 1 | 1 | 1 | 1 | 1 | 1 | 1 | 1 | 1 | 1 | 1 | 1 | 1 | 1 | 1 | 0 | 1 | 1 | 1 | 1 | 1 | 1 | 1 | 1 | 1 | 1 | 1 | 1 | 1 |
| *Bulbophyllum alticola* Schltr. | 1 | 1 | 1 | 1 | 1 | 1 | 1 | 1 | 1 | 1 | 1 | 1 | 1 | 1 | 1 | 1 | 1 | 1 | 1 | 1 | 1 | 1 | 1 | 1 | 1 | 1 | 1 | 1 | 1 | 1 | 1 | 1 | 1 | 1 | 1 | 1 | 1 | 1 | 1 | 1 | 1 | 1 | 1 | 1 | 1 | 1 | 1 | 1 | 1 | 0 | 1 | 1 | 1 | 1 | 1 | 1 | 0 | 1 | 1 | 1 | 1 | 1 | 1 |
| *Bulbophyllum ambrosia* (Hance) Schltr. 1 | 1 | 1 | 1 | 1 | 1 | 1 | 1 | 1 | 1 | 1 | 1 | 1 | 1 | 1 | 1 | 1 | 1 | 1 | 1 | 1 | 1 | 1 | 1 | 1 | 1 | 1 | 1 | 1 | 1 | 1 | 1 | 1 | 1 | 1 | 1 | 1 | 1 | 1 | 1 | 1 | 1 | 1 | 1 | 1 | 1 | 1 | 1 | 1 | 1 | 1 | 1 | 1 | 1 | 1 | 1 | 1 | 0 | 1 | 1 | 1 | 1 | 1 | 1 |
| *Bulbophyllum ambrosia* (Hance) Schltr. 2 | 1 | 1 | 1 | 1 | 1 | 1 | 1 | 1 | 1 | 1 | 1 | 1 | 1 | 1 | 1 | 1 | 1 | 1 | 1 | 1 | 1 | 1 | 1 | 1 | 1 | 1 | 1 | 1 | 1 | 1 | 1 | 1 | 1 | 1 | 1 | 1 | 1 | 1 | 1 | 1 | 1 | 1 | 1 | 1 | 1 | 1 | 1 | 1 | 1 | 1 | 1 | 0 | 1 | 1 | 1 | 1 | 1 | 1 | 1 | 1 | 1 | 1 | 1 |
| *Bulbophyllum amplebracteatum* Teijsm. & Binn. | 1 | 1 | 1 | 1 | 1 | 1 | 1 | 1 | 1 | 1 | 1 | 1 | 1 | 1 | 1 | 1 | 1 | 1 | 1 | 1 | 1 | 1 | 1 | 1 | 1 | 0 | 1 | 1 | 1 | 1 | 1 | 1 | 1 | 1 | 1 | 1 | 1 | 1 | 1 | 1 | 1 | 1 | 1 | 1 | 1 | 1 | 1 | 1 | 1 | 1 | 1 | 1 | 1 | 1 | 1 | 1 | 1 | 1 | 1 | 1 | 1 | 1 | 1 |
| *Bulbophyllum amplebracteatum* subsp. *carunculatum* (Garay, Hamer & Siegerist) J.J.Verm. & P.O'Byrne | 1 | 1 | 1 | 1 | 1 | 1 | 1 | 1 | 1 | 1 | 1 | 1 | 1 | 1 | 1 | 1 | 1 | 1 | 1 | 1 | 1 | 1 | 1 | 1 | 1 | 1 | 1 | 1 | 1 | 1 | 1 | 1 | 1 | 1 | 1 | 1 | 1 | 1 | 1 | 1 | 1 | 1 | 1 | 1 | 1 | 1 | 1 | 1 | 1 | 1 | 1 | 1 | 1 | 1 | 1 | 1 | 1 | 1 | 1 | 1 | 1 | 1 | 1 |
| *Bulbophyllum andreeae* A.D.Hawkes | 0 | 1 | 1 | 1 | 1 | 1 | 1 | 1 | 1 | 1 | 1 | 1 | 1 | 1 | 1 | 1 | 1 | 1 | 1 | 1 | 1 | 1 | 1 | 1 | 1 | 1 | 1 | 1 | 1 | 1 | 1 | 0 | 1 | 1 | 1 | 1 | 1 | 1 | 1 | 1 | 1 | 1 | 1 | 1 | 1 | 1 | 1 | 0 | 1 | 1 | 0 | 1 | 1 | 1 | 1 | 1 | 1 | 1 | 1 | 1 | 1 | 1 | 1 |
| *Bulbophyllum ankylochele* J.J.Verm. | 1 | 1 | 1 | 1 | 1 | 1 | 1 | 1 | 1 | 1 | 1 | 1 | 1 | 1 | 1 | 1 | 1 | 1 | 1 | 1 | 1 | 1 | 1 | 1 | 1 | 1 | 1 | 1 | 1 | 1 | 1 | 1 | 1 | 1 | 1 | 1 | 1 | 1 | 1 | 1 | 1 | 1 | 1 | 1 | 1 | 1 | 1 | 1 | 1 | 1 | 1 | 1 | 1 | 1 | 1 | 1 | 1 | 1 | 1 | 1 | 1 | 1 | 1 |
| *Bulbophyllum annandalei* Ridl. | 1 | 1 | 1 | 1 | 1 | 1 | 1 | 1 | 1 | 1 | 1 | 1 | 1 | 1 | 1 | 1 | 1 | 1 | 0 | 0 | 1 | 1 | 1 | 1 | 1 | 0 | 0 | 1 | 1 | 1 | 1 | 1 | 1 | 1 | 0 | 1 | 1 | 0 | 1 | 1 | 1 | 1 | 1 | 1 | 1 | 1 | 1 | 1 | 1 | 1 | 1 | 1 | 1 | 1 | 1 | 1 | 1 | 0 | 1 | 1 | 1 | 0 | 1 |
| *Bulbophyllum antenniferum* (Lindl.) Rchb.f. | 1 | 1 | 1 | 1 | 1 | 1 | 1 | 1 | 1 | 1 | 1 | 1 | 1 | 1 | 1 | 1 | 1 | 1 | 1 | 1 | 1 | 1 | 1 | 1 | 1 | 1 | 1 | 1 | 1 | 1 | 1 | 1 | 1 | 1 | 1 | 1 | 1 | 1 | 1 | 1 | 1 | 1 | 0 | 1 | 1 | 1 | 1 | 1 | 1 | 1 | 1 | 1 | 1 | 1 | 1 | 1 | 1 | 1 | 1 | 1 | 1 | 1 | 1 |
| *Bulbophyllum apodum* Hook.f. 1 | 1 | 1 | 1 | 1 | 0 | 1 | 1 | 1 | 1 | 1 | 1 | 1 | 1 | 1 | 1 | 1 | 1 | 1 | 1 | 1 | 1 | 1 | 1 | 1 | 1 | 1 | 1 | 1 | 1 | 1 | 1 | 1 | 1 | 1 | 1 | 1 | 1 | 1 | 1 | 1 | 1 | 1 | 1 | 1 | 1 | 1 | 1 | 1 | 1 | 1 | 1 | 1 | 1 | 1 | 1 | 1 | 1 | 1 | 1 | 1 | 1 | 1 | 1 |
| *Bulbophyllum apodum* Hook.f. 2 | 1 | 1 | 1 | 1 | 1 | 1 | 1 | 1 | 1 | 1 | 1 | 1 | 1 | 1 | 1 | 1 | 1 | 1 | 1 | 1 | 1 | 1 | 1 | 1 | 1 | 1 | 1 | 1 | 1 | 1 | 1 | 1 | 1 | 1 | 1 | 1 | 1 | 1 | 1 | 1 | 1 | 1 | 0 | 1 | 1 | 1 | 1 | 1 | 1 | 1 | 1 | 1 | 1 | 1 | 1 | 1 | 1 | 1 | 1 | 1 | 1 | 1 | 1 |
| *Bulbophyllum apodum* Hook.f. 3 | 1 | 1 | 1 | 1 | 1 | 1 | 1 | 1 | 1 | 1 | 1 | 1 | 1 | 1 | 1 | 1 | 1 | 1 | 1 | 1 | 1 | 1 | 1 | 1 | 1 | 1 | 1 | 1 | 1 | 1 | 1 | 1 | 1 | 1 | 1 | 1 | 1 | 1 | 1 | 1 | 1 | 1 | 0 | 1 | 1 | 1 | 1 | 1 | 1 | 1 | 1 | 1 | 1 | 1 | 1 | 1 | 1 | 1 | 1 | 1 | 1 | 1 | 1 |
| *Bulbophyllum arfakianum* Kraenzl. | 1 | 1 | 1 | 1 | 1 | 1 | 1 | 1 | 1 | 1 | 1 | 1 | 1 | 1 | 1 | 1 | 1 | 1 | 1 | 1 | 1 | 1 | 1 | 1 | 1 | 1 | 1 | 1 | 1 | 1 | 1 | 1 | 1 | 1 | 1 | 1 | 1 | 1 | 1 | 1 | 1 | 1 | 1 | 1 | 1 | 1 | 1 | 1 | 1 | 1 | 1 | 1 | 1 | 1 | 1 | 1 | 1 | 1 | 1 | 1 | 1 | 1 | 1 |
| *Bulbophyllum argyropus* (Endl.) Rchb.f. | 1 | 1 | 1 | 1 | 1 | 1 | 1 | 1 | 1 | 1 | 1 | 1 | 1 | 1 | 1 | 1 | 1 | 1 | 1 | 1 | 1 | 1 | 1 | 1 | 1 | 1 | 1 | 1 | 1 | 1 | 1 | 1 | 1 | 1 | 1 | 1 | 1 | 1 | 1 | 1 | 1 | 1 | 1 | 1 | 1 | 1 | 1 | 1 | 1 | 1 | 1 | 1 | 1 | 1 | 1 | 0 | 1 | 1 | 1 | 0 | 1 | 1 | 1 |
| *Bulbophyllum aristilabre* J.J.Sm. | 1 | 1 | 1 | 1 | 1 | 1 | 1 | 1 | 1 | 1 | 1 | 1 | 1 | 1 | 1 | 1 | 1 | 1 | 1 | 1 | 1 | 1 | 1 | 1 | 1 | 1 | 1 | 1 | 1 | 1 | 1 | 1 | 1 | 1 | 1 | 1 | 1 | 1 | 1 | 1 | 1 | 1 | 1 | 1 | 1 | 1 | 1 | 1 | 1 | 1 | 1 | 1 | 1 | 1 | 1 | 1 | 1 | 1 | 1 | 1 | 1 | 1 | 1 |
| *Bulbophyllum ascochilum* J.J.Verm. | 1 | 1 | 1 | 1 | 1 | 1 | 1 | 1 | 1 | 1 | 1 | 1 | 1 | 1 | 1 | 1 | 1 | 1 | 1 | 1 | 1 | 1 | 1 | 1 | 1 | 1 | 1 | 1 | 1 | 1 | 1 | 1 | 1 | 1 | 1 | 1 | 1 | 1 | 1 | 1 | 1 | 1 | 1 | 1 | 1 | 1 | 1 | 1 | 1 | 1 | 1 | 1 | 1 | 1 | 1 | 1 | 1 | 1 | 1 | 1 | 1 | 1 | 1 |
| *Bulbophyllum aundense* Ormerod | 1 | 1 | 1 | 1 | 1 | 1 | 1 | 1 | 1 | 1 | 1 | 1 | 1 | 1 | 1 | 1 | 1 | 1 | 1 | 1 | 1 | 1 | 1 | 1 | 1 | 1 | 1 | 1 | 1 | 1 | 1 | 1 | 1 | 1 | 1 | 1 | 1 | 1 | 1 | 1 | 1 | 1 | 1 | 1 | 1 | 1 | 1 | 1 | 1 | 1 | 1 | 1 | 1 | 1 | 1 | 1 | 1 | 1 | 1 | 1 | 1 | 1 | 1 |
| *Bulbophyllum ayuthayense* J.J.Verm., Schuit. & de Vogel | 1 | 1 | 1 | 1 | 1 | 1 | 1 | 1 | 1 | 1 | 1 | 1 | 1 | 1 | 1 | 1 | 1 | 1 | 1 | 1 | 1 | 1 | 1 | 1 | 1 | 1 | 1 | 1 | 1 | 1 | 1 | 1 | 1 | 1 | 1 | 1 | 1 | 1 | 1 | 1 | 1 | 1 | 0 | 1 | 1 | 1 | 1 | 0 | 1 | 1 | 1 | 1 | 0 | 1 | 1 | 1 | 1 | 1 | 1 | 1 | 1 | 1 | 1 |
| *Bulbophyllum baileyi* F.Muell. 1 | 1 | 1 | 1 | 1 | 1 | 1 | 1 | 1 | 1 | 1 | 1 | 1 | 1 | 1 | 1 | 1 | 1 | 1 | 1 | 1 | 1 | 1 | 1 | 1 | 1 | 1 | 1 | 1 | 1 | 1 | 1 | 1 | 1 | 1 | 1 | 1 | 1 | 1 | 1 | 1 | 1 | 1 | 1 | 1 | 1 | 1 | 1 | 1 | 1 | 1 | 1 | 1 | 1 | 1 | 1 | 1 | 1 | 1 | 1 | 1 | 1 | 1 | 1 |
| *Bulbophyllum baileyi* F.Muell. 2 | 1 | 1 | 1 | 1 | 1 | 1 | 1 | 1 | 1 | 1 | 1 | 1 | 1 | 1 | 1 | 1 | 1 | 1 | 1 | 1 | 1 | 1 | 1 | 1 | 1 | 1 | 1 | 1 | 1 | 1 | 1 | 1 | 1 | 1 | 1 | 1 | 1 | 1 | 1 | 1 | 1 | 1 | 1 | 1 | 1 | 1 | 1 | 1 | 1 | 1 | 1 | 1 | 1 | 1 | 1 | 1 | 1 | 1 | 1 | 1 | 1 | 1 | 1 |
| *Bulbophyllum baileyi* F.Muell. 3 | 1 | 1 | 1 | 1 | 1 | 1 | 1 | 1 | 1 | 1 | 1 | 1 | 1 | 1 | 1 | 1 | 1 | 1 | 1 | 0 | 1 | 1 | 1 | 1 | 1 | 1 | 1 | 1 | 1 | 1 | 1 | 1 | 1 | 1 | 1 | 1 | 1 | 1 | 1 | 1 | 1 | 1 | 1 | 1 | 1 | 1 | 1 | 1 | 1 | 1 | 1 | 1 | 1 | 1 | 1 | 1 | 1 | 1 | 1 | 1 | 1 | 1 | 1 |
| *Bulbophyllum baladeanum* J.J.Sm. 1 | 1 | 1 | 1 | 1 | 1 | 1 | 1 | 1 | 1 | 1 | 1 | 1 | 0 | 1 | 1 | 0 | 0 | 0 | 1 | 1 | 1 | 1 | 1 | 1 | 0 | 1 | 1 | 1 | 1 | 1 | 0 | 1 | 0 | 1 | 1 | 1 | 1 | 1 | 1 | 1 | 1 | 1 | 0 | 1 | 1 | 1 | 1 | 1 | 1 | 1 | 1 | 1 | 1 | 1 | 1 | 1 | 1 | 1 | 1 | 1 | 1 | 1 | 1 |
| *Bulbophyllum baladeanum* J.J.Sm. 2 | 1 | 0 | 0 | 1 | 0 | 0 | 0 | 1 | 1 | 1 | 1 | 1 | 0 | 1 | 1 | 1 | 1 | 1 | 0 | 1 | 1 | 1 | 1 | 0 | 1 | 1 | 0 | 0 | 0 | 1 | 1 | 1 | 1 | 1 | 1 | 1 | 1 | 1 | 1 | 1 | 1 | 0 | 0 | 1 | 1 | 1 | 1 | 1 | 1 | 1 | 0 | 0 | 1 | 1 | 1 | 1 | 1 | 1 | 1 | 1 | 1 | 1 | 1 |
| *Bulbophyllum baladeanum* J.J.Sm. 3 | 1 | 1 | 1 | 1 | 1 | 1 | 1 | 1 | 1 | 1 | 1 | 1 | 1 | 1 | 1 | 1 | 1 | 1 | 1 | 1 | 1 | 1 | 1 | 1 | 1 | 1 | 1 | 1 | 1 | 1 | 1 | 1 | 1 | 1 | 1 | 1 | 1 | 1 | 1 | 1 | 1 | 1 | 1 | 1 | 1 | 1 | 1 | 1 | 1 | 1 | 1 | 1 | 1 | 1 | 1 | 1 | 1 | 1 | 1 | 1 | 1 | 1 | 1 |
| *Bulbophyllum baladeanum* J.J.Sm. 4 | 1 | 1 | 1 | 1 | 1 | 1 | 1 | 1 | 1 | 1 | 1 | 1 | 1 | 1 | 1 | 1 | 1 | 1 | 1 | 1 | 1 | 1 | 1 | 1 | 1 | 1 | 1 | 1 | 1 | 1 | 1 | 1 | 1 | 1 | 1 | 1 | 1 | 1 | 1 | 1 | 1 | 1 | 1 | 1 | 1 | 1 | 1 | 1 | 1 | 1 | 1 | 1 | 1 | 1 | 1 | 1 | 1 | 1 | 1 | 1 | 1 | 1 | 1 |
| *Bulbophyllum balgooiji* J.J.Verm., Schuit. & de Vogel | 1 | 1 | 1 | 1 | 1 | 1 | 1 | 1 | 1 | 1 | 1 | 1 | 1 | 1 | 1 | 1 | 1 | 1 | 1 | 1 | 1 | 1 | 1 | 1 | 1 | 1 | 1 | 1 | 1 | 1 | 1 | 1 | 1 | 1 | 1 | 1 | 1 | 1 | 1 | 1 | 0 | 1 | 1 | 1 | 1 | 1 | 1 | 1 | 1 | 1 | 1 | 1 | 1 | 1 | 1 | 1 | 1 | 1 | 1 | 1 | 1 | 1 | 1 |
| *Bulbophyllum bandischii* Garay, Hamer & Siegerist | 1 | 1 | 1 | 1 | 1 | 1 | 1 | 1 | 1 | 1 | 1 | 1 | 1 | 1 | 1 | 1 | 1 | 1 | 1 | 1 | 1 | 1 | 1 | 1 | 1 | 1 | 1 | 1 | 1 | 1 | 1 | 1 | 1 | 1 | 1 | 1 | 1 | 1 | 1 | 1 | 1 | 1 | 1 | 1 | 1 | 1 | 1 | 1 | 1 | 1 | 1 | 1 | 1 | 1 | 1 | 1 | 1 | 1 | 1 | 1 | 1 | 1 | 1 |
| *Bulbophyllum barbigerum* Lindl. 1 | 1 | 1 | 1 | 1 | 1 | 1 | 1 | 1 | 1 | 1 | 1 | 1 | 1 | 1 | 1 | 1 | 1 | 1 | 1 | 1 | 1 | 1 | 1 | 1 | 1 | 1 | 1 | 1 | 1 | 1 | 1 | 1 | 1 | 1 | 1 | 1 | 1 | 1 | 1 | 1 | 1 | 1 | 1 | 1 | 1 | 1 | 1 | 1 | 1 | 1 | 1 | 1 | 1 | 1 | 1 | 1 | 1 | 1 | 1 | 0 | 1 | 1 | 1 |
| *Bulbophyllum barbigerum* Lindl. 2 | 1 | 1 | 1 | 1 | 1 | 1 | 1 | 1 | 1 | 1 | 1 | 1 | 1 | 1 | 1 | 1 | 1 | 1 | 1 | 1 | 1 | 1 | 1 | 1 | 1 | 1 | 1 | 1 | 1 | 1 | 1 | 1 | 1 | 1 | 1 | 1 | 1 | 1 | 1 | 1 | 1 | 1 | 1 | 1 | 1 | 1 | 1 | 1 | 1 | 1 | 1 | 1 | 1 | 1 | 1 | 1 | 1 | 1 | 1 | 0 | 1 | 1 | 1 |
| *Bulbophyllum baronii* Ridl. 1 | 1 | 1 | 1 | 1 | 1 | 1 | 1 | 1 | 1 | 1 | 1 | 1 | 1 | 1 | 1 | 1 | 1 | 1 | 1 | 1 | 1 | 1 | 1 | 1 | 1 | 1 | 1 | 1 | 1 | 1 | 1 | 1 | 1 | 1 | 1 | 1 | 1 | 1 | 1 | 1 | 1 | 1 | 0 | 1 | 1 | 1 | 1 | 1 | 1 | 1 | 1 | 1 | 1 | 1 | 1 | 1 | 1 | 1 | 1 | 1 | 1 | 1 | 1 |
| *Bulbophyllum baronii* Ridl. 2 | 1 | 1 | 1 | 1 | 1 | 1 | 1 | 1 | 1 | 1 | 1 | 1 | 1 | 1 | 1 | 1 | 1 | 1 | 1 | 1 | 1 | 1 | 1 | 1 | 1 | 1 | 1 | 1 | 1 | 1 | 1 | 1 | 1 | 1 | 1 | 1 | 1 | 1 | 1 | 1 | 1 | 1 | 0 | 1 | 1 | 1 | 1 | 1 | 1 | 1 | 1 | 1 | 1 | 1 | 1 | 1 | 1 | 1 | 1 | 1 | 1 | 1 | 1 |
| *Bulbophyllum basisetum* J.J.Sm. | 1 | 1 | 1 | 1 | 1 | 1 | 1 | 1 | 1 | 1 | 1 | 1 | 1 | 1 | 1 | 1 | 1 | 1 | 1 | 1 | 1 | 1 | 1 | 1 | 1 | 1 | 1 | 1 | 1 | 1 | 1 | 1 | 1 | 1 | 1 | 1 | 1 | 1 | 1 | 1 | 1 | 1 | 1 | 1 | 1 | 1 | 1 | 1 | 1 | 1 | 1 | 1 | 1 | 1 | 1 | 1 | 1 | 1 | 1 | 1 | 1 | 1 | 1 |
| *Bulbophyllum beccarii* Rchb.f. | 1 | 1 | 1 | 1 | 1 | 1 | 1 | 1 | 1 | 1 | 1 | 1 | 1 | 1 | 1 | 1 | 1 | 1 | 1 | 1 | 1 | 1 | 1 | 1 | 1 | 1 | 1 | 1 | 1 | 1 | 1 | 1 | 1 | 1 | 1 | 1 | 1 | 1 | 1 | 1 | 1 | 1 | 0 | 1 | 1 | 1 | 1 | 1 | 1 | 1 | 1 | 1 | 1 | 1 | 1 | 1 | 1 | 1 | 1 | 1 | 1 | 1 | 1 |
| *Bulbophyllum betchei* F.Muell. | 1 | 1 | 1 | 1 | 1 | 1 | 1 | 1 | 1 | 1 | 1 | 1 | 1 | 1 | 1 | 1 | 1 | 1 | 1 | 1 | 1 | 1 | 1 | 1 | 1 | 1 | 1 | 1 | 1 | 1 | 1 | 1 | 1 | 1 | 1 | 1 | 1 | 1 | 1 | 1 | 1 | 1 | 0 | 1 | 1 | 1 | 1 | 1 | 1 | 1 | 1 | 1 | 1 | 1 | 1 | 1 | 1 | 1 | 1 | 1 | 1 | 1 | 1 |
| *Bulbophyllum biantennatum* Schltr. | 1 | 1 | 1 | 1 | 1 | 1 | 1 | 1 | 1 | 1 | 1 | 1 | 1 | 1 | 1 | 1 | 1 | 1 | 1 | 1 | 1 | 1 | 1 | 1 | 1 | 1 | 1 | 1 | 1 | 1 | 1 | 1 | 1 | 1 | 1 | 1 | 1 | 1 | 1 | 1 | 1 | 1 | 1 | 1 | 1 | 1 | 1 | 1 | 1 | 1 | 1 | 1 | 1 | 1 | 1 | 1 | 1 | 1 | 1 | 1 | 1 | 1 | 1 |
| *Bulbophyllum bicolor* Lindl. | 1 | 1 | 1 | 1 | 1 | 1 | 1 | 1 | 1 | 1 | 1 | 1 | 1 | 1 | 1 | 1 | 1 | 1 | 1 | 1 | 1 | 1 | 1 | 1 | 1 | 1 | 1 | 1 | 1 | 1 | 1 | 1 | 1 | 1 | 1 | 1 | 1 | 1 | 1 | 1 | 1 | 1 | 1 | 1 | 1 | 1 | 1 | 1 | 1 | 1 | 1 | 1 | 1 | 1 | 1 | 1 | 1 | 1 | 1 | 1 | 1 | 1 | 1 |
| *Bulbophyllum blepharistes* Rchb.f. | 1 | 1 | 1 | 1 | 1 | 1 | 1 | 1 | 1 | 1 | 1 | 1 | 1 | 1 | 1 | 1 | 1 | 1 | 1 | 1 | 1 | 1 | 1 | 1 | 1 | 1 | 1 | 1 | 1 | 1 | 1 | 1 | 1 | 1 | 1 | 1 | 1 | 1 | 1 | 1 | 1 | 0 | 1 | 1 | 1 | 1 | 1 | 0 | 1 | 1 | 1 | 0 | 0 | 1 | 1 | 1 | 1 | 1 | 1 | 0 | 1 | 1 | 1 |
| *Bulbophyllum boonjee* B.Gray & D.L.Jones | 1 | 1 | 1 | 1 | 1 | 1 | 1 | 1 | 1 | 1 | 1 | 1 | 1 | 1 | 1 | 1 | 1 | 1 | 1 | 1 | 1 | 1 | 1 | 1 | 1 | 1 | 1 | 1 | 1 | 1 | 1 | 1 | 1 | 1 | 1 | 1 | 1 | 1 | 1 | 1 | 1 | 1 | 1 | 1 | 1 | 1 | 1 | 1 | 1 | 0 | 1 | 1 | 1 | 1 | 1 | 1 | 0 | 1 | 1 | 1 | 1 | 1 | 1 |
| *Bulbophyllum bowkettiae* F.M.Bailey | 1 | 1 | 1 | 1 | 1 | 1 | 1 | 1 | 1 | 0 | 1 | 1 | 1 | 1 | 1 | 1 | 1 | 1 | 1 | 1 | 1 | 1 | 1 | 1 | 1 | 1 | 1 | 1 | 1 | 1 | 1 | 1 | 1 | 1 | 1 | 1 | 1 | 1 | 1 | 1 | 1 | 1 | 1 | 1 | 1 | 1 | 1 | 1 | 1 | 0 | 1 | 1 | 1 | 1 | 1 | 1 | 0 | 1 | 1 | 1 | 1 | 1 | 1 |
| *Bulbophyllum bracteatum* (Fitzg.) F.M.Bailey | 1 | 1 | 1 | 1 | 1 | 1 | 1 | 1 | 1 | 1 | 1 | 1 | 1 | 1 | 1 | 1 | 1 | 1 | 1 | 1 | 1 | 1 | 1 | 1 | 1 | 1 | 1 | 1 | 1 | 1 | 1 | 1 | 1 | 1 | 1 | 1 | 1 | 1 | 1 | 1 | 1 | 1 | 1 | 1 | 1 | 1 | 1 | 1 | 1 | 1 | 1 | 1 | 1 | 1 | 1 | 1 | 1 | 1 | 1 | 1 | 1 | 1 | 1 |
| *Bulbophyllum brassii* J.J.Verm. | 1 | 1 | 1 | 1 | 1 | 1 | 1 | 1 | 1 | 1 | 1 | 1 | 1 | 1 | 1 | 1 | 1 | 1 | 1 | 1 | 1 | 1 | 1 | 1 | 1 | 1 | 1 | 1 | 1 | 1 | 1 | 1 | 1 | 1 | 1 | 1 | 1 | 1 | 1 | 1 | 1 | 1 | 1 | 1 | 1 | 1 | 1 | 1 | 1 | 1 | 1 | 1 | 1 | 1 | 1 | 1 | 1 | 1 | 1 | 1 | 1 | 1 | 1 |
| *Bulbophyllum brevilabium* Schltr. | 1 | 1 | 1 | 1 | 1 | 1 | 1 | 1 | 1 | 1 | 1 | 1 | 1 | 1 | 1 | 1 | 1 | 1 | 1 | 1 | 1 | 1 | 1 | 1 | 1 | 1 | 1 | 1 | 1 | 1 | 1 | 1 | 1 | 1 | 1 | 1 | 1 | 1 | 0 | 1 | 1 | 1 | 1 | 1 | 1 | 1 | 1 | 1 | 1 | 1 | 1 | 1 | 1 | 1 | 1 | 1 | 1 | 1 | 1 | 1 | 1 | 1 | 1 |
| *Bulbophyllum bulliferum* J.J.Sm. | 1 | 1 | 1 | 1 | 1 | 1 | 1 | 1 | 1 | 1 | 1 | 1 | 1 | 1 | 1 | 1 | 1 | 1 | 1 | 1 | 1 | 1 | 1 | 1 | 1 | 1 | 1 | 1 | 1 | 1 | 1 | 1 | 1 | 1 | 1 | 1 | 1 | 1 | 1 | 1 | 1 | 1 | 1 | 1 | 1 | 1 | 1 | 1 | 1 | 1 | 1 | 1 | 1 | 1 | 1 | 1 | 1 | 1 | 0 | 1 | 1 | 1 | 1 |
| *Bulbophyllum cambodianum* (Christenson) J.J.Verm., Schuit. & de Vogel | 1 | 1 | 1 | 1 | 1 | 1 | 1 | 1 | 0 | 1 | 1 | 1 | 1 | 1 | 1 | 1 | 1 | 1 | 1 | 1 | 1 | 1 | 1 | 1 | 1 | 1 | 1 | 1 | 1 | 1 | 1 | 1 | 1 | 1 | 1 | 1 | 1 | 1 | 1 | 1 | 1 | 1 | 0 | 1 | 1 | 1 | 1 | 1 | 1 | 1 | 0 | 1 | 1 | 1 | 1 | 1 | 1 | 1 | 1 | 0 | 1 | 1 | 1 |
| *Bulbophyllum caudatum* Lindl. | 1 | 1 | 1 | 1 | 1 | 1 | 1 | 1 | 1 | 1 | 1 | 1 | 1 | 1 | 1 | 1 | 1 | 1 | 1 | 1 | 1 | 1 | 1 | 1 | 1 | 1 | 1 | 1 | 1 | 1 | 1 | 1 | 1 | 1 | 1 | 1 | 1 | 1 | 1 | 1 | 1 | 1 | 1 | 1 | 1 | 1 | 1 | 1 | 1 | 1 | 1 | 1 | 1 | 1 | 1 | 1 | 1 | 1 | 1 | 1 | 1 | 1 | 1 |
| *Bulbophyllum cauliflorum* Hook.f. | 1 | 1 | 1 | 1 | 1 | 1 | 1 | 1 | 1 | 1 | 1 | 1 | 0 | 1 | 1 | 1 | 1 | 1 | 1 | 1 | 1 | 1 | 1 | 1 | 1 | 1 | 1 | 1 | 1 | 1 | 1 | 1 | 1 | 1 | 1 | 1 | 1 | 1 | 1 | 1 | 1 | 1 | 0 | 1 | 1 | 1 | 1 | 1 | 1 | 1 | 0 | 1 | 1 | 1 | 1 | 1 | 1 | 1 | 1 | 1 | 1 | 1 | 1 |

| | | | | | | | | | | | | | | | | | | | | | | | | | | | | | | | | | | | | | | | | | | | | | | | | | | | | | | | | | | | | | | | |
|---|---|---|---|---|---|---|---|---|---|---|---|---|---|---|---|---|---|---|---|---|---|---|---|---|---|---|---|---|---|---|---|---|---|---|---|---|---|---|---|---|---|---|---|---|---|---|---|---|---|---|---|---|---|---|---|---|---|---|---|---|---|---|---|
| *Bulbophyllum cheiri* Lindl. | 1 | 1 | 1 | 1 | 1 | 1 | 1 | 1 | 1 | 1 | 1 | 1 | 1 | 1 | 1 | 1 | 1 | 1 | 1 | 1 | 1 | 1 | 1 | 1 | 1 | 1 | 1 | 1 | 1 | 1 | 1 | 1 | 1 | 1 | 1 | 1 | 1 | 1 | 1 | 1 | 1 | 1 | 1 | 1 | 1 | 1 | 1 | 1 | 1 | 1 | 1 | 1 | 1 | 1 | 1 | 1 | 1 | 1 | 1 | 1 | 1 | 1 | 1 |
| *Bulbophyllum chloranthum* Schltr. | 1 | 1 | 1 | 1 | 1 | 1 | 1 | 1 | 1 | 1 | 1 | 1 | 1 | 1 | 1 | 1 | 1 | 1 | 1 | 1 | 1 | 1 | 1 | 1 | 1 | 1 | 1 | 1 | 1 | 1 | 1 | 1 | 1 | 1 | 1 | 1 | 1 | 1 | 1 | 1 | 1 | 1 | 0 | 1 | 1 | 1 | 1 | 1 | 1 | 1 | 1 | 1 | 1 | 1 | 1 | 1 | 1 | 1 | 1 | 0 | 1 | 1 | 1 |
| *Bulbophyllum chrysendetum* Ames 1 | 1 | 1 | 1 | 1 | 1 | 1 | 1 | 1 | 1 | 1 | 1 | 1 | 1 | 1 | 1 | 1 | 1 | 1 | 1 | 1 | 1 | 1 | 1 | 1 | 1 | 1 | 1 | 1 | 1 | 1 | 1 | 1 | 1 | 1 | 1 | 1 | 1 | 1 | 1 | 1 | 1 | 1 | 1 | 1 | 1 | 1 | 1 | 1 | 1 | 1 | 1 | 1 | 1 | 1 | 1 | 1 | 1 | 1 | 1 | 1 | 1 | 1 | 1 |
| *Bulbophyllum chrysendetum* Ames 2 | 1 | 1 | 1 | 1 | 1 | 1 | 1 | 1 | 1 | 1 | 1 | 1 | 1 | 1 | 1 | 1 | 1 | 1 | 1 | 1 | 1 | 1 | 1 | 1 | 1 | 1 | 1 | 1 | 1 | 1 | 1 | 1 | 1 | 1 | 1 | 1 | 1 | 1 | 1 | 1 | 1 | 1 | 1 | 1 | 1 | 1 | 1 | 1 | 0 | 1 | 1 | 1 | 1 | 1 | 1 | 1 | 1 | 1 | 1 | 1 | 1 | 1 | 1 |
| *Bulbophyllum ciliatum* (Blume) Lindl. | 1 | 1 | 1 | 1 | 1 | 1 | 1 | 1 | 1 | 1 | 1 | 1 | 1 | 1 | 1 | 1 | 1 | 1 | 1 | 1 | 1 | 1 | 1 | 1 | 1 | 1 | 1 | 1 | 1 | 1 | 1 | 1 | 1 | 1 | 1 | 1 | 1 | 1 | 1 | 1 | 1 | 1 | 0 | 1 | 1 | 1 | 1 | 1 | 1 | 1 | 1 | 1 | 1 | 1 | 1 | 1 | 1 | 1 | 1 | 1 | 1 | 1 | 1 |
| *Bulbophyllum clandestinum* Lindl. 1 | 1 | 1 | 1 | 1 | 1 | 1 | 1 | 1 | 1 | 1 | 1 | 1 | 1 | 1 | 1 | 1 | 1 | 1 | 1 | 1 | 1 | 1 | 1 | 1 | 1 | 1 | 1 | 1 | 1 | 1 | 1 | 1 | 1 | 1 | 1 | 1 | 1 | 1 | 1 | 1 | 1 | 1 | 0 | 1 | 1 | 1 | 1 | 1 | 1 | 1 | 1 | 1 | 1 | 1 | 1 | 1 | 1 | 1 | 1 | 1 | 1 | 1 | 1 |
| *Bulbophyllum clandestinum* Lindl. 2 | 1 | 1 | 1 | 1 | 1 | 1 | 1 | 1 | 1 | 1 | 1 | 1 | 1 | 1 | 1 | 1 | 1 | 1 | 1 | 1 | 1 | 1 | 1 | 1 | 1 | 1 | 1 | 1 | 1 | 1 | 1 | 1 | 1 | 1 | 1 | 1 | 1 | 1 | 1 | 1 | 1 | 1 | 0 | 1 | 1 | 1 | 1 | 1 | 1 | 1 | 1 | 1 | 1 | 1 | 1 | 1 | 1 | 1 | 1 | 1 | 1 | 1 | 1 |
| *Bulbophyllum cleistogamum* Ridl. | 0 | 1 | 1 | 1 | 1 | 1 | 1 | 1 | 1 | 1 | 1 | 1 | 1 | 1 | 1 | 1 | 1 | 1 | 1 | 1 | 1 | 1 | 1 | 1 | 1 | 1 | 1 | 1 | 1 | 1 | 1 | 1 | 1 | 1 | 1 | 1 | 1 | 1 | 1 | 1 | 1 | 1 | 1 | 1 | 0 | 1 | 1 | 1 | 1 | 1 | 1 | 1 | 1 | 1 | 1 | 1 | 1 | 1 | 1 | 1 | 1 | 1 | 1 |
| *Bulbophyllum collettii* King & Pantl. | 1 | 1 | 1 | 1 | 1 | 1 | 1 | 1 | 1 | 1 | 1 | 1 | 1 | 1 | 1 | 1 | 1 | 1 | 1 | 1 | 1 | 1 | 1 | 1 | 1 | 1 | 1 | 1 | 1 | 1 | 1 | 1 | 1 | 1 | 1 | 1 | 1 | 1 | 1 | 1 | 1 | 1 | 1 | 1 | 1 | 1 | 1 | 1 | 1 | 1 | 1 | 1 | 1 | 1 | 1 | 1 | 1 | 1 | 1 | 1 | 1 | 1 | 1 |
| *Bulbophyllum colliferum* J.J.Sm. | 1 | 1 | 1 | 1 | 1 | 1 | 1 | 1 | 1 | 1 | 1 | 1 | 1 | 1 | 1 | 1 | 1 | 1 | 1 | 1 | 1 | 1 | 1 | 1 | 1 | 1 | 1 | 1 | 1 | 1 | 1 | 1 | 1 | 1 | 1 | 1 | 1 | 1 | 1 | 1 | 1 | 1 | 1 | 1 | 1 | 1 | 1 | 1 | 1 | 0 | 1 | 1 | 1 | 1 | 1 | 1 | 1 | 1 | 1 | 1 | 1 | 1 | 1 |
| *Bulbophyllum comberi* J.J.Verm. | 1 | 1 | 1 | 1 | 1 | 1 | 1 | 1 | 1 | 1 | 1 | 1 | 1 | 0 | 1 | 1 | 1 | 1 | 1 | 1 | 1 | 1 | 1 | 1 | 1 | 1 | 1 | 1 | 1 | 1 | 1 | 1 | 1 | 1 | 1 | 1 | 1 | 1 | 1 | 1 | 1 | 1 | 1 | 1 | 1 | 1 | 1 | 1 | 1 | 1 | 1 | 1 | 1 | 1 | 1 | 1 | 0 | 1 | 1 | 1 | 1 | 1 | 1 |
| *Bulbophyllum comosum* Collett & Hemsl. | 1 | 1 | 1 | 1 | 1 | 1 | 1 | 1 | 1 | 1 | 1 | 1 | 1 | 1 | 1 | 1 | 1 | 1 | 1 | 1 | 1 | 1 | 1 | 1 | 1 | 1 | 1 | 1 | 1 | 1 | 1 | 1 | 1 | 1 | 1 | 1 | 1 | 1 | 1 | 1 | 1 | 1 | 0 | 1 | 1 | 1 | 1 | 1 | 1 | 0 | 1 | 1 | 1 | 1 | 1 | 1 | 1 | 1 | 1 | 1 | 1 | 1 | 1 |
| *Bulbophyllum concinnum* Hook.f. | 1 | 1 | 1 | 1 | 1 | 1 | 1 | 1 | 1 | 1 | 1 | 1 | 1 | 1 | 1 | 1 | 1 | 1 | 1 | 1 | 1 | 1 | 1 | 1 | 1 | 1 | 1 | 1 | 1 | 1 | 1 | 1 | 1 | 1 | 1 | 1 | 1 | 1 | 1 | 1 | 1 | 1 | 1 | 1 | 1 | 1 | 1 | 1 | 1 | 1 | 1 | 1 | 1 | 1 | 1 | 1 | 1 | 1 | 1 | 1 | 1 | 1 | 1 |
| *Bulbophyllum contortisepalum* J.J.Sm. | 0 | 1 | 1 | 1 | 1 | 1 | 1 | 1 | 1 | 1 | 1 | 1 | 1 | 1 | 1 | 1 | 1 | 1 | 1 | 1 | 1 | 1 | 1 | 1 | 1 | 1 | 1 | 1 | 1 | 1 | 1 | 1 | 1 | 1 | 1 | 1 | 1 | 1 | 1 | 1 | 1 | 1 | 0 | 1 | 1 | 1 | 1 | 1 | 1 | 1 | 1 | 1 | 1 | 1 | 1 | 1 | 1 | 1 | 1 | 1 | 1 | 1 | 1 |
| *Bulbophyllum cootesii* M.A.Clem. | 1 | 1 | 1 | 1 | 1 | 1 | 1 | 1 | 1 | 1 | 1 | 1 | 1 | 1 | 1 | 1 | 1 | 1 | 1 | 1 | 1 | 1 | 1 | 1 | 1 | 1 | 1 | 1 | 1 | 1 | 1 | 1 | 1 | 1 | 1 | 1 | 1 | 1 | 1 | 1 | 1 | 1 | 1 | 1 | 1 | 1 | 1 | 1 | 1 | 1 | 1 | 1 | 1 | 1 | 1 | 1 | 1 | 1 | 1 | 1 | 1 | 1 | 1 |
| *Bulbophyllum corythium* N.Hallé | 1 | 1 | 1 | 1 | 1 | 1 | 1 | 1 | 1 | 1 | 1 | 1 | 1 | 1 | 1 | 1 | 1 | 0 | 1 | 1 | 1 | 1 | 1 | 1 | 1 | 1 | 0 | 1 | 1 | 1 | 1 | 1 | 1 | 1 | 1 | 1 | 1 | 1 | 1 | 1 | 1 | 1 | 1 | 1 | 1 | 1 | 1 | 1 | 1 | 1 | 1 | 1 | 1 | 1 | 1 | 1 | 1 | 0 | 1 | 0 | 1 | 1 | 1 |
| *Bulbophyllum crassipes* Hook.f. 1 | 1 | 1 | 1 | 1 | 1 | 1 | 1 | 1 | 1 | 1 | 1 | 1 | 1 | 1 | 1 | 1 | 1 | 1 | 1 | 1 | 1 | 1 | 1 | 1 | 1 | 1 | 1 | 1 | 1 | 1 | 1 | 1 | 1 | 1 | 1 | 1 | 1 | 1 | 1 | 1 | 1 | 1 | 1 | 1 | 1 | 1 | 1 | 1 | 1 | 1 | 1 | 1 | 1 | 1 | 1 | 1 | 0 | 1 | 1 | 1 | 1 | 1 | 1 |
| *Bulbophyllum crassipes* Hook.f. 2 | 1 | 1 | 1 | 1 | 1 | 1 | 1 | 1 | 1 | 1 | 1 | 1 | 1 | 1 | 1 | 1 | 1 | 1 | 1 | 1 | 1 | 1 | 1 | 1 | 1 | 1 | 1 | 1 | 1 | 1 | 1 | 1 | 1 | 1 | 1 | 1 | 1 | 1 | 1 | 1 | 1 | 1 | 1 | 1 | 1 | 1 | 1 | 1 | 1 | 1 | 1 | 1 | 1 | 1 | 1 | 1 | 1 | 1 | 1 | 1 | 1 | 1 | 1 |
| *Bulbophyllum cruciatum* J.J.Sm. 1 | 1 | 1 | 1 | 1 | 1 | 1 | 1 | 1 | 1 | 1 | 1 | 1 | 1 | 1 | 1 | 1 | 1 | 1 | 1 | 1 | 1 | 1 | 1 | 1 | 1 | 1 | 1 | 1 | 1 | 1 | 1 | 1 | 1 | 1 | 1 | 1 | 1 | 1 | 1 | 1 | 1 | 1 | 0 | 1 | 1 | 1 | 1 | 1 | 1 | 1 | 1 | 1 | 1 | 1 | 1 | 1 | 1 | 1 | 1 | 1 | 1 | 1 | 1 |
| *Bulbophyllum cruciatum* J.J.Sm. 2 | 1 | 1 | 1 | 1 | 1 | 1 | 1 | 1 | 1 | 1 | 1 | 1 | 1 | 1 | 1 | 1 | 1 | 1 | 1 | 1 | 1 | 1 | 1 | 0 | 1 | 1 | 0 | 1 | 1 | 1 | 1 | 1 | 1 | 1 | 1 | 1 | 1 | 1 | 1 | 1 | 1 | 1 | 1 | 1 | 1 | 1 | 1 | 1 | 1 | 1 | 1 | 1 | 1 | 1 | 1 | 1 | 1 | 0 | 1 | 1 | 1 | 1 | 1 |
| *Bulbophyllum cruentum* Garay, Hamer & Siegerist | 1 | 1 | 1 | 1 | 1 | 1 | 1 | 0 | 1 | 0 | 0 | 1 | 1 | 0 | 0 | 0 | 0 | 1 | 1 | 1 | 0 | 1 | 0 | 1 | 0 | 1 | 1 | 0 | 0 | 0 | 1 | 0 | 1 | 0 | 1 | 0 | 0 | 1 | 1 | 0 | 0 | 0 | 0 | 0 | 0 | 0 | 0 | 0 | 1 | 1 | 1 | 1 | 0 | 0 | 0 | 0 | 0 | 1 | 0 | 1 | 0 | 1 | 1 |
| *Bulbophyllum cryptophoranthus* Garay | 1 | 1 | 1 | 1 | 1 | 1 | 1 | 1 | 1 | 1 | 1 | 1 | 1 | 1 | 1 | 1 | 1 | 1 | 1 | 1 | 1 | 1 | 1 | 1 | 0 | 1 | 1 | 1 | 1 | 1 | 1 | 1 | 1 | 1 | 1 | 1 | 1 | 1 | 1 | 1 | 1 | 0 | 0 | 1 | 1 | 1 | 1 | 1 | 0 | 1 | 1 | 0 | 1 | 1 | 1 | 1 | 1 | 1 | 0 | 1 | 1 | 1 | 1 |
| *Bulbophyllum cuniculiforme* J.J.Sm. | 1 | 1 | 1 | 1 | 1 | 1 | 1 | 1 | 1 | 1 | 1 | 1 | 1 | 1 | 1 | 1 | 1 | 1 | 1 | 1 | 1 | 1 | 1 | 1 | 1 | 1 | 1 | 1 | 1 | 1 | 1 | 1 | 1 | 1 | 1 | 1 | 1 | 1 | 1 | 1 | 1 | 1 | 1 | 1 | 1 | 1 | 1 | 1 | 1 | 1 | 1 | 1 | 1 | 1 | 1 | 1 | 1 | 1 | 1 | 0 | 1 | 1 | 1 |
| *Bulbophyllum cyatheicola* (P.Royen) J.J.Verm., Schuit. & de Vogel | 1 | 1 | 1 | 1 | 1 | 1 | 1 | 1 | 1 | 1 | 1 | 1 | 1 | 1 | 1 | 1 | 1 | 1 | 1 | 1 | 1 | 1 | 1 | 1 | 1 | 1 | 1 | 1 | 1 | 1 | 1 | 1 | 1 | 1 | 1 | 1 | 1 | 1 | 1 | 1 | 0 | 1 | 1 | 1 | 1 | 1 | 1 | 1 | 1 | 1 | 1 | 1 | 1 | 1 | 1 | 1 | 1 | 1 | 1 | 1 | 1 | 1 | 1 |
| *Bulbophyllum dayanum* Rchb.f. | 1 | 1 | 1 | 1 | 1 | 1 | 1 | 1 | 1 | 1 | 1 | 1 | 1 | 1 | 1 | 1 | 1 | 1 | 1 | 1 | 1 | 1 | 1 | 1 | 0 | 1 | 1 | 1 | 1 | 1 | 1 | 1 | 1 | 1 | 1 | 1 | 1 | 1 | 1 | 1 | 1 | 1 | 1 | 1 | 1 | 1 | 1 | 0 | 1 | 1 | 1 | 1 | 1 | 1 | 1 | 1 | 1 | 1 | 1 | 1 | 1 | 1 | 1 |
| *Bulbophyllum dearei* W.Watson 1 | 1 | 1 | 1 | 1 | 1 | 1 | 1 | 1 | 1 | 1 | 1 | 1 | 1 | 1 | 1 | 1 | 1 | 1 | 1 | 1 | 1 | 1 | 1 | 1 | 1 | 1 | 1 | 1 | 1 | 1 | 1 | 1 | 1 | 1 | 1 | 1 | 1 | 1 | 1 | 1 | 1 | 1 | 1 | 1 | 1 | 1 | 1 | 1 | 1 | 1 | 1 | 1 | 1 | 1 | 1 | 1 | 1 | 1 | 1 | 1 | 1 | 1 | 1 |
| *Bulbophyllum dearei* W.Watson 2 | 1 | 1 | 1 | 1 | 1 | 1 | 1 | 1 | 1 | 1 | 1 | 1 | 1 | 1 | 1 | 1 | 1 | 1 | 1 | 1 | 1 | 1 | 1 | 1 | 1 | 1 | 1 | 1 | 1 | 1 | 1 | 1 | 1 | 1 | 1 | 1 | 1 | 1 | 1 | 1 | 1 | 1 | 1 | 1 | 1 | 1 | 1 | 1 | 1 | 1 | 1 | 1 | 1 | 1 | 1 | 1 | 1 | 1 | 1 | 1 | 1 | 1 | 1 |
| *Bulbophyllum delitescens* Hance | 1 | 1 | 1 | 1 | 1 | 1 | 1 | 1 | 1 | 1 | 1 | 1 | 1 | 1 | 1 | 1 | 1 | 1 | 1 | 1 | 1 | 1 | 1 | 1 | 1 | 1 | 1 | 1 | 1 | 1 | 1 | 1 | 1 | 1 | 1 | 1 | 1 | 1 | 1 | 1 | 1 | 1 | 1 | 1 | 1 | 1 | 1 | 1 | 1 | 1 | 1 | 1 | 1 | 1 | 1 | 1 | 1 | 1 | 1 | 1 | 1 | 1 | 1 |
| *Bulbophyllum depressum* King & Pantl. | 1 | 1 | 1 | 1 | 1 | 1 | 1 | 0 | 1 | 1 | 1 | 1 | 1 | 1 | 1 | 1 | 1 | 1 | 1 | 1 | 1 | 1 | 1 | 1 | 1 | 1 | 1 | 1 | 1 | 1 | 1 | 1 | 1 | 1 | 1 | 1 | 1 | 1 | 1 | 1 | 1 | 1 | 1 | 1 | 1 | 1 | 1 | 1 | 1 | 1 | 1 | 1 | 1 | 1 | 1 | 1 | 1 | 1 | 1 | 1 | 1 | 1 | 1 |
| *Bulbophyllum desmotrichoides* Schltr. | 1 | 1 | 1 | 1 | 1 | 1 | 1 | 1 | 1 | 1 | 1 | 1 | 1 | 0 | 1 | 1 | 1 | 1 | 1 | 1 | 1 | 1 | 1 | 1 | 1 | 1 | 1 | 1 | 1 | 1 | 1 | 1 | 1 | 1 | 1 | 1 | 1 | 1 | 1 | 1 | 1 | 1 | 1 | 1 | 1 | 1 | 1 | 1 | 1 | 1 | 1 | 1 | 1 | 1 | 1 | 1 | 0 | 1 | 1 | 1 | 1 | 1 | 1 |
| *Bulbophyllum dichotomum* J.J.Sm. | 1 | 1 | 1 | 1 | 1 | 1 | 1 | 1 | 1 | 1 | 1 | 1 | 1 | 0 | 1 | 1 | 1 | 1 | 1 | 1 | 1 | 1 | 1 | 1 | 1 | 1 | 1 | 1 | 1 | 1 | 1 | 1 | 1 | 1 | 1 | 1 | 1 | 0 | 1 | 1 | 0 | 1 | 1 | 1 | 1 | 1 | 1 | 1 | 1 | 0 | 1 | 1 | 1 | 1 | 1 | 1 | 0 | 1 | 1 | 1 | 1 | 1 | 1 |
| *Bulbophyllum dictyoneuron* Schltr. | 1 | 1 | 1 | 1 | 1 | 1 | 1 | 1 | 1 | 1 | 1 | 1 | 1 | 1 | 1 | 1 | 1 | 1 | 1 | 1 | 1 | 1 | 1 | 1 | 1 | 1 | 1 | 1 | 1 | 1 | 1 | 1 | 1 | 1 | 1 | 1 | 1 | 1 | 1 | 1 | 1 | 1 | 1 | 1 | 1 | 1 | 1 | 1 | 0 | 1 | 1 | 1 | 1 | 1 | 1 | 1 | 1 | 1 | 1 | 1 | 1 | 1 | 1 |
| *Bulbophyllum digoelense* J.J.Sm. | 1 | 1 | 1 | 1 | 1 | 1 | 1 | 1 | 1 | 1 | 1 | 1 | 1 | 0 | 1 | 1 | 1 | 1 | 1 | 1 | 1 | 1 | 1 | 1 | 1 | 1 | 1 | 1 | 1 | 1 | 1 | 1 | 1 | 1 | 1 | 1 | 1 | 1 | 1 | 1 | 1 | 1 | 1 | 1 | 1 | 1 | 1 | 1 | 1 | 1 | 1 | 1 | 1 | 1 | 1 | 1 | 0 | 1 | 1 | 1 | 1 | 1 | 1 |
| *Bulbophyllum dischidiifolium* J.J.Sm. | 1 | 1 | 1 | 1 | 1 | 1 | 1 | 1 | 1 | 1 | 1 | 1 | 1 | 1 | 1 | 1 | 1 | 1 | 1 | 1 | 1 | 1 | 1 | 1 | 1 | 1 | 1 | 1 | 1 | 1 | 1 | 1 | 1 | 1 | 1 | 1 | 1 | 1 | 1 | 1 | 1 | 1 | 0 | 1 | 1 | 1 | 1 | 1 | 1 | 1 | 1 | 1 | 1 | 1 | 1 | 1 | 1 | 1 | 1 | 1 | 1 | 1 | 1 |
| *Bulbophyllum disciflorum* Rolfe | 1 | 1 | 1 | 1 | 1 | 1 | 1 | 1 | 1 | 1 | 1 | 1 | 1 | 1 | 1 | 1 | 1 | 1 | 1 | 1 | 1 | 1 | 1 | 1 | 1 | 1 | 1 | 1 | 1 | 1 | 1 | 1 | 1 | 1 | 1 | 1 | 1 | 1 | 1 | 1 | 1 | 1 | 1 | 1 | 1 | 1 | 1 | 1 | 1 | 1 | 1 | 1 | 1 | 1 | 1 | 1 | 1 | 1 | 1 | 1 | 1 | 1 | 1 |
| *Bulbophyllum discolor* Schltr. | 0 | 1 | 1 | 1 | 1 | 1 | 1 | 1 | 1 | 1 | 1 | 1 | 1 | 1 | 1 | 1 | 1 | 1 | 0 | 1 | 1 | 1 | 1 | 1 | 1 | 1 | 1 | 1 | 1 | 1 | 1 | 1 | 1 | 1 | 1 | 1 | 1 | 1 | 1 | 1 | 1 | 1 | 1 | 1 | 1 | 1 | 1 | 1 | 0 | 1 | 1 | 1 | 1 | 1 | 1 | 1 | 1 | 1 | 1 | 1 | 1 | 1 | 1 |
| *Bulbophyllum dissitiflorum* Seidenf. | 1 | 1 | 1 | 1 | 1 | 1 | 1 | 1 | 1 | 1 | 1 | 1 | 1 | 1 | 1 | 1 | 1 | 1 | 1 | 1 | 1 | 1 | 1 | 1 | 1 | 1 | 1 | 1 | 1 | 1 | 1 | 1 | 1 | 1 | 1 | 1 | 1 | 1 | 1 | 1 | 1 | 1 | 1 | 1 | 1 | 1 | 1 | 1 | 1 | 1 | 1 | 1 | 1 | 1 | 1 | 1 | 1 | 1 | 1 | 1 | 1 | 1 | 1 |
| *Bulbophyllum djamuense* Schltr. | 1 | 1 | 1 | 1 | 1 | 1 | 1 | 1 | 1 | 1 | 1 | 1 | 1 | 1 | 1 | 1 | 1 | 1 | 1 | 1 | 1 | 1 | 1 | 1 | 1 | 1 | 1 | 1 | 1 | 1 | 1 | 1 | 1 | 1 | 1 | 1 | 1 | 1 | 1 | 1 | 1 | 1 | 1 | 1 | 1 | 0 | 1 | 1 | 1 | 1 | 1 | 1 | 1 | 1 | 1 | 1 | 1 | 1 | 1 | 1 | 1 | 1 | 1 |
| *Bulbophyllum echinolabium* J.J.Sm. | 1 | 1 | 1 | 1 | 1 | 1 | 1 | 1 | 1 | 1 | 1 | 1 | 1 | 1 | 1 | 1 | 1 | 1 | 1 | 1 | 1 | 1 | 1 | 1 | 1 | 1 | 1 | 1 | 1 | 1 | 1 | 1 | 1 | 1 | 1 | 1 | 1 | 1 | 1 | 1 | 1 | 1 | 0 | 1 | 1 | 1 | 1 | 1 | 1 | 1 | 1 | 1 | 1 | 1 | 1 | 1 | 1 | 1 | 1 | 1 | 1 | 1 | 1 |
| *Bulbophyllum echinulus* Seidenf. | 1 | 1 | 1 | 1 | 1 | 1 | 1 | 1 | 1 | 1 | 1 | 1 | 1 | 1 | 1 | 1 | 1 | 1 | 1 | 1 | 1 | 1 | 1 | 1 | 1 | 1 | 1 | 1 | 1 | 1 | 1 | 1 | 1 | 1 | 1 | 1 | 1 | 1 | 1 | 1 | 1 | 1 | 1 | 1 | 1 | 1 | 1 | 1 | 1 | 1 | 1 | 1 | 1 | 1 | 1 | 1 | 1 | 1 | 1 | 1 | 1 | 1 | 1 |
| *Bulbophyllum eciliatum* Schltr. | 1 | 1 | 1 | 1 | 1 | 1 | 1 | 1 | 1 | 1 | 0 | 1 | 1 | 0 | 1 | 1 | 0 | 0 | 1 | 1 | 1 | 1 | 1 | 1 | 1 | 1 | 1 | 1 | 0 | 1 | 1 | 1 | 0 | 0 | 1 | 1 | 1 | 0 | 1 | 1 | 0 | 1 | 1 | 1 | 1 | 1 | 1 | 1 | 1 | 0 | 1 | 1 | 1 | 1 | 1 | 1 | 0 | 1 | 1 | 1 | 1 | 1 | 1 |
| *Bulbophyllum ecornutum* (J.J.Sm.) J.J.Sm. | 1 | 1 | 1 | 1 | 1 | 1 | 1 | 1 | 1 | 1 | 1 | 1 | 1 | 1 | 1 | 1 | 1 | 1 | 1 | 1 | 1 | 1 | 1 | 1 | 1 | 1 | 1 | 1 | 1 | 1 | 1 | 1 | 1 | 1 | 1 | 1 | 1 | 1 | 1 | 1 | 1 | 1 | 1 | 1 | 1 | 1 | 1 | 1 | 1 | 1 | 1 | 1 | 1 | 1 | 1 | 1 | 1 | 1 | 1 | 1 | 1 | 1 | 1 |
| *Bulbophyllum elassoglossum* Siegerist | 1 | 1 | 1 | 1 | 1 | 1 | 1 | 1 | 1 | 1 | 1 | 1 | 1 | 1 | 1 | 1 | 1 | 1 | 1 | 1 | 1 | 1 | 1 | 1 | 1 | 1 | 1 | 1 | 1 | 1 | 1 | 1 | 1 | 1 | 1 | 1 | 1 | 1 | 1 | 1 | 1 | 1 | 1 | 1 | 1 | 1 | 1 | 1 | 1 | 1 | 1 | 1 | 1 | 1 | 1 | 1 | 1 | 1 | 1 | 1 | 1 | 1 | 1 |
| *Bulbophyllum elegantius* Schltr. | 0 | 1 | 1 | 1 | 1 | 1 | 1 | 0 | 1 | 1 | 1 | 0 | 1 | 0 | 0 | 1 | 1 | 1 | 1 | 1 | 1 | 1 | 1 | 1 | 1 | 1 | 1 | 1 | 1 | 1 | 1 | 1 | 1 | 1 | 1 | 1 | 1 | 1 | 1 | 1 | 1 | 1 | 0 | 1 | 1 | 0 | 1 | 0 | 0 | 1 | 0 | 0 | 0 | 1 | 1 | 0 | 1 | 1 | 1 | 0 | 0 | 1 | 1 |
| *Bulbophyllum elephantinum* J.J.Sm. | 1 | 1 | 1 | 1 | 1 | 1 | 1 | 1 | 1 | 1 | 1 | 1 | 1 | 0 | 1 | 1 | 1 | 1 | 1 | 1 | 1 | 1 | 1 | 1 | 1 | 1 | 1 | 1 | 1 | 1 | 1 | 1 | 0 | 1 | 1 | 1 | 1 | 1 | 1 | 1 | 1 | 1 | 1 | 1 | 1 | 1 | 1 | 1 | 1 | 0 | 1 | 1 | 1 | 1 | 1 | 1 | 0 | 1 | 1 | 1 | 1 | 0 | 1 |
| *Bulbophyllum elevatopunctatum* J.J.Sm. 1 | 1 | 1 | 1 | 1 | 1 | 1 | 1 | 1 | 1 | 1 | 1 | 1 | 1 | 1 | 1 | 1 | 1 | 1 | 1 | 1 | 1 | 1 | 1 | 1 | 1 | 1 | 1 | 1 | 1 | 1 | 1 | 1 | 1 | 1 | 1 | 1 | 1 | 1 | 1 | 1 | 1 | 1 | 0 | 1 | 1 | 1 | 1 | 1 | 1 | 1 | 1 | 1 | 1 | 1 | 1 | 1 | 1 | 1 | 1 | 1 | 1 | 1 | 1 |
| *Bulbophyllum elevatopunctatum* J.J.Sm. 2 | 1 | 1 | 1 | 1 | 1 | 1 | 1 | 1 | 1 | 1 | 1 | 1 | 1 | 1 | 1 | 1 | 1 | 1 | 1 | 1 | 1 | 1 | 1 | 1 | 1 | 1 | 1 | 1 | 1 | 1 | 1 | 1 | 1 | 1 | 1 | 1 | 1 | 1 | 1 | 1 | 1 | 1 | 1 | 1 | 1 | 1 | 1 | 1 | 1 | 1 | 1 | 1 | 1 | 1 | 1 | 1 | 1 | 1 | 1 | 1 | 1 | 1 | 1 |
| *Bulbophyllum elisae* (F.Muell.) Benth. | 1 | 1 | 1 | 1 | 1 | 1 | 1 | 1 | 1 | 1 | 1 | 1 | 1 | 1 | 1 | 1 | 1 | 1 | 1 | 1 | 1 | 1 | 1 | 0 | 1 | 1 | 1 | 1 | 1 | 1 | 1 | 1 | 1 | 1 | 1 | 1 | 1 | 1 | 1 | 1 | 1 | 0 | 1 | 1 | 1 | 1 | 1 | 0 | 1 | 0 | 1 | 1 | 0 | 1 | 0 | 0 | 1 | 1 | 1 | 1 | 1 | 1 | 1 |
| *Bulbophyllum emarginatum* (Finet) J.J.Sm. | 1 | 1 | 1 | 1 | 1 | 1 | 1 | 1 | 1 | 1 | 1 | 1 | 1 | 1 | 1 | 1 | 1 | 1 | 1 | 1 | 1 | 1 | 1 | 1 | 1 | 1 | 1 | 1 | 1 | 1 | 1 | 1 | 1 | 1 | 1 | 1 | 1 | 1 | 1 | 1 | 1 | 1 | 1 | 1 | 1 | 1 | 1 | 1 | 1 | 1 | 1 | 1 | 1 | 1 | 1 | 1 | 1 | 1 | 1 | 0 | 1 | 1 | 1 |
| *Bulbophyllum epiphytum* Barb.Rodr. | 1 | 1 | 1 | 1 | 1 | 1 | 1 | 1 | 1 | 1 | 1 | 1 | 1 | 1 | 1 | 1 | 1 | 1 | 1 | 1 | 1 | 1 | 1 | 1 | 1 | 1 | 1 | 1 | 1 | 1 | 1 | 1 | 1 | 1 | 1 | 1 | 1 | 1 | 1 | 1 | 1 | 1 | 1 | 1 | 1 | 1 | 1 | 1 | 1 | 1 | 1 | 1 | 1 | 1 | 1 | 1 | 0 | 1 | 1 | 1 | 1 | 1 | 1 |
| *Bulbophyllum ericssonii* Kraenzl. 1 | 1 | 1 | 1 | 1 | 1 | 1 | 1 | 1 | 1 | 1 | 1 | 1 | 1 | 1 | 1 | 1 | 1 | 1 | 1 | 0 | 1 | 1 | 1 | 1 | 1 | 1 | 1 | 1 | 1 | 1 | 1 | 1 | 1 | 1 | 1 | 1 | 1 | 1 | 1 | 1 | 1 | 1 | 1 | 1 | 1 | 1 | 1 | 1 | 1 | 1 | 1 | 1 | 1 | 1 | 1 | 1 | 1 | 1 | 1 | 0 | 1 | 1 | 1 |
| *Bulbophyllum ericssonii* Kraenzl. 2 | 1 | 1 | 1 | 1 | 1 | 1 | 1 | 1 | 1 | 1 | 1 | 1 | 1 | 1 | 1 | 1 | 1 | 1 | 1 | 1 | 1 | 1 | 1 | 1 | 1 | 1 | 1 | 1 | 1 | 1 | 1 | 1 | 1 | 1 | 1 | 1 | 1 | 1 | 1 | 1 | 1 | 1 | 1 | 1 | 1 | 1 | 1 | 1 | 1 | 1 | 1 | 1 | 1 | 1 | 1 | 1 | 1 | 1 | 1 | 0 | 1 | 1 | 1 |
| *Bulbophyllum erinaceum* Schltr. | 1 | 0 | 1 | 1 | 1 | 1 | 1 | 1 | 1 | 1 | 1 | 1 | 1 | 1 | 1 | 1 | 1 | 1 | 1 | 1 | 1 | 1 | 1 | 1 | 1 | 1 | 1 | 1 | 1 | 1 | 1 | 1 | 1 | 1 | 1 | 1 | 1 | 1 | 1 | 1 | 1 | 1 | 1 | 1 | 1 | 1 | 1 | 1 | 1 | 1 | 1 | 1 | 1 | 1 | 1 | 1 | 1 | 1 | 1 | 1 | 1 | 1 | 1 |
| *Bulbophyllum erythrosema* J.J.Verm. | 1 | 1 | 1 | 1 | 1 | 1 | 1 | 1 | 1 | 1 | 1 | 1 | 1 | 1 | 1 | 1 | 1 | 1 | 1 | 1 | 1 | 1 | 1 | 1 | 1 | 1 | 0 | 1 | 1 | 1 | 1 | 1 | 1 | 1 | 1 | 1 | 1 | 1 | 1 | 1 | 1 | 1 | 1 | 1 | 1 | 1 | 1 | 1 | 1 | 1 | 1 | 1 | 1 | 1 | 1 | 1 | 1 | 0 | 1 | 1 | 1 | 1 | 1 |
| *Bulbophyllum evasum* T.E.Hunt & Rupp | 1 | 1 | 1 | 1 | 1 | 1 | 1 | 1 | 1 | 1 | 1 | 1 | 1 | 1 | 1 | 1 | 1 | 1 | 1 | 1 | 1 | 1 | 1 | 1 | 1 | 1 | 1 | 1 | 1 | 1 | 1 | 1 | 1 | 1 | 1 | 1 | 1 | 1 | 1 | 1 | 1 | 1 | 1 | 1 | 1 | 1 | 1 | 1 | 1 | 1 | 1 | 1 | 1 | 1 | 1 | 1 | 0 | 1 | 1 | 1 | 1 | 1 | 1 |
| *Bulbophyllum exaltatum* Lindl. | 1 | 1 | 1 | 1 | 1 | 1 | 1 | 1 | 1 | 1 | 1 | 1 | 1 | 1 | 1 | 1 | 1 | 1 | 1 | 1 | 1 | 1 | 1 | 1 | 1 | 1 | 1 | 1 | 1 | 1 | 1 | 1 | 1 | 1 | 1 | 1 | 1 | 1 | 1 | 1 | 1 | 1 | 1 | 1 | 1 | 1 | 1 | 1 | 1 | 1 | 1 | 1 | 1 | 1 | 1 | 1 | 0 | 1 | 1 | 1 | 1 | 1 | 1 |
| *Bulbophyllum exiguum* F.Muell. | 1 | 1 | 1 | 1 | 1 | 1 | 1 | 1 | 1 | 1 | 1 | 1 | 1 | 1 | 1 | 1 | 1 | 1 | 1 | 1 | 1 | 1 | 1 | 1 | 1 | 1 | 1 | 1 | 1 | 1 | 1 | 1 | 1 | 1 | 1 | 1 | 1 | 1 | 1 | 1 | 1 | 1 | 0 | 1 | 1 | 1 | 1 | 1 | 1 | 1 | 1 | 1 | 1 | 1 | 1 | 1 | 1 | 1 | 1 | 1 | 1 | 1 | 1 |
| *Bulbophyllum facetum* Garay, Hamer & Siegerist | 1 | 1 | 1 | 1 | 1 | 1 | 1 | 1 | 1 | 1 | 1 | 1 | 1 | 1 | 1 | 1 | 1 | 1 | 1 | 1 | 1 | 1 | 1 | 1 | 1 | 1 | 1 | 1 | 1 | 1 | 1 | 1 | 1 | 1 | 1 | 1 | 1 | 1 | 1 | 1 | 1 | 1 | 1 | 1 | 1 | 1 | 1 | 1 | 1 | 1 | 1 | 1 | 1 | 1 | 1 | 1 | 1 | 1 | 1 | 1 | 1 | 1 | 1 |
| *Bulbophyllum falcatum* (Lindl.) Rchb.f. | 1 | 1 | 1 | 1 | 1 | 1 | 1 | 1 | 1 | 1 | 1 | 1 | 1 | 1 | 0 | 1 | 1 | 1 | 1 | 1 | 1 | 1 | 1 | 1 | 1 | 1 | 1 | 1 | 1 | 0 | 1 | 1 | 1 | 1 | 1 | 1 | 1 | 1 | 1 | 1 | 1 | 1 | 1 | 1 | 1 | 1 | 1 | 1 | 1 | 1 | 1 | 1 | 1 | 1 | 1 | 0 | 1 | 1 | 1 | 0 | 1 | 1 | 0 |
| *Bulbophyllum farreri* (W.W.Sm.) Seidenf. | 1 | 1 | 1 | 1 | 1 | 1 | 1 | 1 | 1 | 1 | 1 | 1 | 1 | 1 | 1 | 1 | 1 | 1 | 1 | 1 | 1 | 1 | 1 | 1 | 1 | 1 | 1 | 1 | 1 | 1 | 1 | 1 | 1 | 1 | 1 | 1 | 1 | 1 | 1 | 1 | 1 | 1 | 1 | 1 | 1 | 1 | 1 | 1 | 1 | 1 | 1 | 1 | 1 | 1 | 1 | 1 | 0 | 1 | 1 | 1 | 1 | 1 | 1 |
| *Bulbophyllum fascinator* (Rolfe) Rolfe | 1 | 1 | 1 | 1 | 1 | 1 | 1 | 1 | 1 | 1 | 1 | 1 | 1 | 1 | 1 | 1 | 1 | 1 | 1 | 1 | 1 | 1 | 1 | 0 | 1 | 1 | 1 | 1 | 1 | 1 | 1 | 1 | 1 | 1 | 1 | 1 | 1 | 1 | 1 | 1 | 1 | 1 | 1 | 1 | 1 | 1 | 1 | 1 | 1 | 1 | 1 | 1 | 1 | 1 | 1 | 1 | 1 | 1 | 1 | 1 | 1 | 1 | 1 |
| *Bulbophyllum fenixii* Ames | 1 | 1 | 1 | 1 | 1 | 1 | 1 | 1 | 1 | 1 | 1 | 1 | 1 | 1 | 1 | 1 | 1 | 1 | 1 | 1 | 1 | 1 | 1 | 1 | 1 | 1 | 1 | 1 | 1 | 1 | 1 | 1 | 1 | 1 | 1 | 1 | 1 | 1 | 1 | 1 | 1 | 1 | 1 | 1 | 1 | 1 | 1 | 1 | 1 | 1 | 1 | 1 | 1 | 1 | 1 | 1 | 1 | 1 | 1 | 1 | 1 | 1 | 1 |
| *Bulbophyllum flammuliferum* Ridl. 1 | 1 | 1 | 1 | 1 | 1 | 1 | 1 | 1 | 1 | 1 | 1 | 1 | 1 | 1 | 1 | 1 | 1 | 1 | 1 | 1 | 1 | 1 | 1 | 1 | 1 | 1 | 1 | 1 | 1 | 1 | 1 | 1 | 1 | 1 | 1 | 0 | 1 | 1 | 1 | 1 | 1 | 1 | 1 | 1 | 1 | 1 | 1 | 1 | 1 | 1 | 1 | 1 | 1 | 1 | 1 | 1 | 0 | 1 | 1 | 1 | 1 | 0 | 0 |

| | | | | | | | | | | | | | | | | | | | | | | | | | | | | | | | | | | | | | | | | | | | | | | | | | | | | | | | | | | | | | | | |
|---|---|---|---|---|---|---|---|---|---|---|---|---|---|---|---|---|---|---|---|---|---|---|---|---|---|---|---|---|---|---|---|---|---|---|---|---|---|---|---|---|---|---|---|---|---|---|---|---|---|---|---|---|---|---|---|---|---|---|---|---|---|---|---|
| *Bulbophyllum flammuliferum* Ridl. 2 | 1 | 1 | 1 | 1 | 1 | 1 | 1 | 1 | 1 | 1 | 1 | 1 | 1 | 1 | 1 | 1 | 1 | 1 | 1 | 1 | 1 | 1 | 1 | 1 | 1 | 1 | 1 | 1 | 1 | 1 | 1 | 1 | 1 | 1 | 1 | 1 | 1 | 1 | 1 | 1 | 1 | 1 | 1 | 1 | 1 | 1 | 1 | 1 | 1 | 1 | 1 | 1 | 1 | 1 | 1 | 1 | 1 | 1 | 1 | 1 | 1 | 1 | 1 |
| *Bulbophyllum flavescens* Lindl. | 1 | 1 | 1 | 1 | 1 | 1 | 1 | 1 | 1 | 1 | 1 | 1 | 1 | 1 | 1 | 1 | 1 | 1 | 1 | 1 | 1 | 1 | 1 | 1 | 1 | 1 | 1 | 1 | 1 | 1 | 1 | 1 | 1 | 1 | 1 | 1 | 1 | 1 | 1 | 1 | 1 | 1 | 0 | 1 | 1 | 1 | 1 | 1 | 1 | 1 | 1 | 1 | 1 | 1 | 0 | 1 | 1 | 1 | 1 | 1 | 1 | 1 | 1 |
| *Bulbophyllum fletcherianum* J.G.Fowler | 1 | 1 | 1 | 1 | 1 | 1 | 1 | 1 | 1 | 1 | 1 | 1 | 1 | 1 | 1 | 1 | 1 | 1 | 1 | 1 | 1 | 1 | 1 | 1 | 1 | 1 | 1 | 1 | 1 | 1 | 1 | 1 | 1 | 1 | 1 | 1 | 1 | 1 | 1 | 1 | 1 | 1 | 1 | 1 | 1 | 1 | 1 | 1 | 1 | 1 | 1 | 1 | 1 | 1 | 1 | 1 | 1 | 1 | 1 | 1 | 1 | 1 | 1 |
| *Bulbophyllum foetidum* Schltr. | 1 | 1 | 1 | 1 | 1 | 1 | 1 | 1 | 1 | 1 | 1 | 1 | 1 | 1 | 1 | 1 | 1 | 1 | 1 | 1 | 1 | 1 | 1 | 1 | 1 | 1 | 1 | 1 | 1 | 1 | 1 | 1 | 1 | 1 | 1 | 1 | 1 | 1 | 1 | 1 | 1 | 1 | 0 | 1 | 1 | 1 | 1 | 0 | 1 | 1 | 1 | 1 | 1 | 1 | 1 | 1 | 1 | 1 | 1 | 1 | 1 | 1 | 1 |
| *Bulbophyllum fonsflorum* J.J.Verm. | 1 | 1 | 1 | 1 | 1 | 1 | 1 | 1 | 1 | 1 | 1 | 1 | 1 | 1 | 1 | 1 | 1 | 1 | 1 | 1 | 1 | 1 | 1 | 1 | 1 | 1 | 1 | 1 | 1 | 1 | 1 | 1 | 1 | 1 | 1 | 1 | 1 | 1 | 1 | 1 | 1 | 1 | 1 | 1 | 1 | 1 | 1 | 1 | 1 | 1 | 1 | 1 | 1 | 1 | 1 | 1 | 1 | 1 | 1 | 1 | 1 | 1 | 1 |
| *Bulbophyllum forrestii* Seidenf. | 1 | 1 | 1 | 1 | 1 | 1 | 1 | 1 | 1 | 1 | 1 | 1 | 1 | 1 | 1 | 1 | 1 | 1 | 1 | 1 | 1 | 1 | 1 | 1 | 1 | 1 | 1 | 1 | 1 | 1 | 1 | 1 | 1 | 1 | 1 | 1 | 1 | 1 | 1 | 1 | 1 | 1 | 0 | 1 | 1 | 1 | 1 | 1 | 1 | 1 | 1 | 1 | 1 | 1 | 1 | 1 | 1 | 1 | 1 | 1 | 1 | 1 | 1 |
| *Bulbophyllum fractiflexum* J.J.Sm. | 1 | 1 | 1 | 1 | 1 | 1 | 1 | 1 | 1 | 1 | 1 | 1 | 1 | 1 | 1 | 1 | 1 | 1 | 1 | 1 | 1 | 1 | 1 | 1 | 1 | 1 | 1 | 1 | 1 | 1 | 1 | 1 | 1 | 1 | 1 | 1 | 1 | 1 | 1 | 1 | 0 | 1 | 0 | 1 | 1 | 1 | 1 | 1 | 1 | 1 | 0 | 1 | 1 | 1 | 1 | 1 | 1 | 1 | 1 | 1 | 1 | 1 | 1 |
| *Bulbophyllum fritillariiflorum* J.J.Sm. | 1 | 1 | 1 | 1 | 1 | 1 | 1 | 1 | 1 | 1 | 1 | 1 | 1 | 1 | 1 | 1 | 1 | 1 | 1 | 1 | 1 | 1 | 1 | 1 | 1 | 1 | 1 | 1 | 1 | 1 | 1 | 1 | 1 | 1 | 1 | 1 | 1 | 1 | 1 | 1 | 1 | 1 | 1 | 1 | 1 | 1 | 1 | 1 | 1 | 1 | 1 | 1 | 1 | 1 | 1 | 1 | 1 | 1 | 1 | 1 | 1 | 1 | 1 |
| *Bulbophyllum frostii* Summerh. 1 | 1 | 1 | 1 | 1 | 1 | 1 | 1 | 1 | 1 | 1 | 1 | 1 | 1 | 1 | 1 | 1 | 1 | 1 | 1 | 1 | 1 | 1 | 1 | 1 | 1 | 1 | 1 | 1 | 1 | 1 | 1 | 1 | 1 | 1 | 1 | 1 | 1 | 1 | 1 | 1 | 1 | 1 | 1 | 1 | 1 | 1 | 1 | 1 | 1 | 1 | 1 | 1 | 1 | 1 | 1 | 1 | 1 | 1 | 1 | 1 | 1 | 1 | 1 |
| *Bulbophyllum frostii* Summerh. 2 | 1 | 1 | 1 | 1 | 1 | 1 | 1 | 1 | 1 | 1 | 1 | 1 | 1 | 1 | 1 | 1 | 1 | 1 | 1 | 1 | 1 | 1 | 1 | 1 | 1 | 1 | 1 | 1 | 1 | 1 | 1 | 1 | 1 | 1 | 1 | 1 | 1 | 1 | 1 | 1 | 1 | 1 | 1 | 1 | 1 | 1 | 1 | 1 | 1 | 1 | 1 | 1 | 1 | 1 | 1 | 1 | 1 | 1 | 1 | 1 | 1 | 1 | 1 |
| *Bulbophyllum fruticicola* Schltr. | 1 | 1 | 1 | 1 | 1 | 1 | 1 | 1 | 1 | 1 | 1 | 1 | 1 | 1 | 1 | 1 | 1 | 1 | 1 | 1 | 1 | 1 | 1 | 1 | 1 | 1 | 1 | 1 | 1 | 1 | 1 | 1 | 1 | 1 | 1 | 1 | 1 | 1 | 1 | 1 | 1 | 1 | 0 | 1 | 1 | 1 | 1 | 1 | 1 | 1 | 1 | 1 | 1 | 1 | 1 | 1 | 1 | 1 | 1 | 1 | 1 | 1 | 1 |
| *Bulbophyllum gadgarrense* Rupp | 1 | 1 | 1 | 1 | 1 | 1 | 1 | 1 | 1 | 1 | 1 | 1 | 1 | 1 | 1 | 1 | 1 | 1 | 1 | 1 | 1 | 1 | 1 | 1 | 1 | 0 | 0 | 1 | 1 | 1 | 1 | 1 | 1 | 1 | 1 | 1 | 1 | 1 | 1 | 1 | 1 | 1 | 1 | 1 | 1 | 1 | 1 | 1 | 1 | 1 | 1 | 1 | 1 | 1 | 1 | 1 | 1 | 1 | 1 | 1 | 1 | 1 | 1 |
| *Bulbophyllum gerlandianum* Kraenzl. | 1 | 1 | 1 | 1 | 1 | 1 | 1 | 1 | 1 | 1 | 1 | 1 | 1 | 1 | 1 | 1 | 1 | 1 | 1 | 1 | 1 | 1 | 1 | 1 | 1 | 1 | 1 | 1 | 1 | 1 | 1 | 1 | 1 | 1 | 1 | 1 | 1 | 1 | 1 | 1 | 1 | 1 | 1 | 1 | 1 | 1 | 1 | 1 | 1 | 1 | 1 | 1 | 1 | 1 | 1 | 1 | 1 | 1 | 1 | 1 | 1 | 1 | 1 |
| *Bulbophyllum gibbosum* (Blume) Lindl. | 1 | 1 | 1 | 1 | 1 | 1 | 1 | 1 | 1 | 1 | 1 | 1 | 1 | 1 | 1 | 1 | 1 | 1 | 1 | 1 | 1 | 1 | 1 | 1 | 1 | 1 | 1 | 1 | 1 | 1 | 1 | 1 | 1 | 1 | 1 | 1 | 1 | 1 | 1 | 1 | 1 | 1 | 1 | 1 | 1 | 1 | 1 | 1 | 1 | 1 | 1 | 1 | 1 | 1 | 1 | 1 | 1 | 1 | 1 | 1 | 1 | 1 | 1 |
| *Bulbophyllum gracillimum* Rolfe | 1 | 1 | 1 | 0 | 1 | 1 | 0 | 1 | 1 | 1 | 0 | 1 | 1 | 1 | 1 | 1 | 1 | 1 | 1 | 1 | 1 | 1 | 1 | 0 | 1 | 1 | 0 | 1 | 1 | 1 | 1 | 1 | 1 | 1 | 1 | 1 | 1 | 1 | 1 | 1 | 0 | 0 | 1 | 1 | 1 | 1 | 1 | 1 | 1 | 1 | 1 | 1 | 1 | 1 | 0 | 1 | 1 | 1 | 1 | 1 | 1 | 1 | 1 |
| *Bulbophyllum grandiflorum* Blume 1 | 1 | 1 | 1 | 1 | 1 | 1 | 1 | 1 | 1 | 1 | 1 | 0 | 1 | 1 | 1 | 1 | 1 | 1 | 1 | 1 | 1 | 1 | 1 | 0 | 1 | 1 | 1 | 1 | 1 | 1 | 1 | 1 | 0 | 1 | 1 | 1 | 1 | 1 | 1 | 1 | 1 | 1 | 1 | 1 | 1 | 1 | 1 | 1 | 1 | 1 | 1 | 1 | 1 | 1 | 1 | 1 | 1 | 1 | 1 | 0 | 1 | 1 | 1 |
| *Bulbophyllum grandiflorum* Blume 2 | 1 | 1 | 1 | 1 | 1 | 1 | 1 | 1 | 1 | 1 | 1 | 1 | 1 | 1 | 1 | 1 | 1 | 1 | 1 | 1 | 1 | 1 | 1 | 1 | 1 | 1 | 1 | 1 | 1 | 1 | 1 | 1 | 1 | 1 | 1 | 1 | 1 | 1 | 1 | 1 | 1 | 1 | 1 | 1 | 1 | 1 | 1 | 1 | 1 | 1 | 1 | 1 | 1 | 1 | 1 | 1 | 1 | 1 | 1 | 1 | 1 | 1 | 1 |
| *Bulbophyllum grandiflorum* Blume 3 | 1 | 1 | 1 | 1 | 1 | 1 | 1 | 1 | 1 | 1 | 1 | 1 | 1 | 1 | 1 | 1 | 1 | 1 | 1 | 1 | 1 | 1 | 1 | 1 | 1 | 1 | 1 | 1 | 1 | 1 | 1 | 1 | 1 | 1 | 1 | 1 | 1 | 1 | 1 | 1 | 1 | 1 | 1 | 1 | 1 | 1 | 1 | 1 | 1 | 1 | 1 | 1 | 1 | 1 | 1 | 1 | 1 | 1 | 1 | 1 | 1 | 1 | 1 |
| *Bulbophyllum grandimesense* B.Gray & D.L.Jones | 1 | 1 | 1 | 1 | 1 | 1 | 1 | 1 | 1 | 1 | 1 | 1 | 1 | 1 | 1 | 1 | 1 | 1 | 1 | 1 | 1 | 1 | 1 | 1 | 1 | 1 | 1 | 1 | 1 | 1 | 1 | 1 | 1 | 1 | 1 | 1 | 1 | 1 | 1 | 1 | 1 | 1 | 1 | 1 | 1 | 1 | 1 | 1 | 1 | 1 | 1 | 1 | 1 | 1 | 1 | 1 | 1 | 1 | 1 | 1 | 1 | 1 | 1 |
| *Bulbophyllum granulosum* Barb.Rodr. | 1 | 1 | 1 | 1 | 1 | 1 | 1 | 1 | 1 | 1 | 1 | 1 | 1 | 1 | 1 | 1 | 1 | 1 | 1 | 1 | 1 | 1 | 1 | 1 | 1 | 1 | 1 | 1 | 1 | 1 | 1 | 1 | 1 | 1 | 1 | 1 | 1 | 1 | 1 | 1 | 1 | 1 | 1 | 1 | 1 | 1 | 1 | 1 | 1 | 1 | 1 | 1 | 1 | 1 | 1 | 1 | 0 | 1 | 1 | 1 | 1 | 1 | 1 |
| *Bulbophyllum guttulatum* (Hook.f.) N.P.Balakr. | 1 | 1 | 1 | 1 | 1 | 1 | 1 | 1 | 1 | 1 | 1 | 1 | 1 | 1 | 1 | 1 | 1 | 1 | 1 | 1 | 1 | 1 | 1 | 1 | 1 | 1 | 1 | 1 | 1 | 1 | 1 | 1 | 1 | 1 | 1 | 1 | 1 | 1 | 1 | 1 | 1 | 1 | 0 | 1 | 1 | 0 | 1 | 1 | 1 | 1 | 1 | 1 | 1 | 1 | 1 | 1 | 1 | 1 | 1 | 1 | 1 | 1 | 1 |
| *Bulbophyllum gyaloglossum* J.J.Verm. | 1 | 1 | 1 | 1 | 1 | 1 | 1 | 1 | 1 | 1 | 1 | 1 | 1 | 1 | 1 | 1 | 1 | 1 | 1 | 1 | 1 | 1 | 1 | 1 | 1 | 1 | 1 | 1 | 1 | 1 | 1 | 1 | 1 | 1 | 1 | 1 | 1 | 1 | 1 | 1 | 1 | 1 | 1 | 1 | 1 | 1 | 1 | 1 | 1 | 0 | 1 | 1 | 1 | 1 | 1 | 1 | 1 | 1 | 1 | 1 | 1 | 1 | 1 |
| *Bulbophyllum gymnopus* Hook.f. | 1 | 1 | 1 | 1 | 1 | 1 | 1 | 1 | 1 | 1 | 1 | 1 | 1 | 1 | 1 | 1 | 1 | 1 | 1 | 1 | 1 | 1 | 1 | 1 | 1 | 1 | 1 | 1 | 1 | 0 | 1 | 1 | 1 | 1 | 1 | 1 | 1 | 1 | 1 | 1 | 1 | 1 | 1 | 1 | 1 | 1 | 1 | 1 | 1 | 1 | 0 | 1 | 1 | 1 | 1 | 1 | 1 | 1 | 1 | 1 | 1 | 1 | 0 |
| *Bulbophyllum hahlianum* Schltr. | 1 | 1 | 1 | 1 | 1 | 1 | 1 | 1 | 1 | 1 | 1 | 1 | 1 | 1 | 1 | 1 | 1 | 1 | 1 | 1 | 1 | 1 | 1 | 1 | 1 | 1 | 1 | 1 | 1 | 1 | 1 | 1 | 1 | 1 | 1 | 1 | 1 | 1 | 1 | 1 | 1 | 1 | 1 | 1 | 1 | 1 | 1 | 1 | 1 | 1 | 1 | 1 | 1 | 1 | 1 | 1 | 0 | 1 | 1 | 1 | 1 | 0 | 1 |
| *Bulbophyllum hamelinii* W.Watson | 1 | 1 | 1 | 1 | 1 | 1 | 1 | 1 | 1 | 1 | 1 | 1 | 1 | 1 | 1 | 1 | 1 | 1 | 1 | 1 | 1 | 1 | 1 | 1 | 1 | 1 | 1 | 1 | 1 | 0 | 1 | 1 | 1 | 1 | 1 | 1 | 1 | 1 | 1 | 1 | 1 | 1 | 1 | 1 | 1 | 1 | 1 | 0 | 0 | 1 | 0 | 1 | 1 | 1 | 1 | 1 | 1 | 1 | 0 | 1 | 1 | 1 | 1 |
| *Bulbophyllum haniffii* Carr | 1 | 1 | 1 | 1 | 1 | 1 | 1 | 1 | 1 | 1 | 1 | 1 | 1 | 1 | 1 | 1 | 1 | 1 | 1 | 1 | 1 | 1 | 1 | 1 | 1 | 1 | 1 | 1 | 1 | 1 | 1 | 1 | 1 | 1 | 1 | 1 | 1 | 1 | 1 | 1 | 1 | 1 | 1 | 1 | 1 | 1 | 1 | 1 | 1 | 0 | 1 | 1 | 1 | 1 | 1 | 1 | 0 | 1 | 1 | 1 | 1 | 1 | 1 |
| *Bulbophyllum helenae* (Kuntze) J.J.Sm. | 1 | 1 | 1 | 1 | 1 | 1 | 1 | 1 | 1 | 1 | 1 | 1 | 1 | 1 | 1 | 1 | 1 | 1 | 1 | 1 | 1 | 1 | 1 | 1 | 1 | 1 | 1 | 1 | 1 | 1 | 1 | 1 | 1 | 1 | 1 | 1 | 1 | 1 | 1 | 1 | 1 | 1 | 1 | 1 | 1 | 1 | 1 | 1 | 1 | 1 | 1 | 1 | 1 | 1 | 1 | 1 | 1 | 1 | 1 | 1 | 1 | 1 | 1 |
| *Bulbophyllum hexarhopalon* Schltr. | 1 | 1 | 1 | 1 | 1 | 1 | 1 | 1 | 1 | 1 | 1 | 1 | 1 | 1 | 1 | 1 | 1 | 1 | 1 | 1 | 1 | 1 | 1 | 1 | 1 | 1 | 1 | 1 | 1 | 1 | 1 | 1 | 1 | 1 | 1 | 1 | 1 | 1 | 1 | 1 | 1 | 1 | 1 | 1 | 1 | 1 | 1 | 1 | 1 | 1 | 1 | 1 | 1 | 1 | 1 | 1 | 1 | 1 | 1 | 1 | 1 | 1 | 1 |
| *Bulbophyllum hians* Schltr. | 1 | 1 | 1 | 1 | 1 | 1 | 1 | 1 | 1 | 1 | 1 | 1 | 1 | 1 | 1 | 1 | 1 | 1 | 1 | 1 | 1 | 1 | 1 | 1 | 1 | 1 | 1 | 1 | 1 | 1 | 1 | 1 | 1 | 1 | 1 | 1 | 1 | 1 | 1 | 1 | 1 | 1 | 1 | 1 | 1 | 1 | 1 | 1 | 1 | 1 | 1 | 1 | 1 | 1 | 1 | 1 | 1 | 1 | 1 | 1 | 1 | 1 | 1 |
| *Bulbophyllum hirtum* (Sm.) Lindl. ex Wall. | 1 | 1 | 1 | 1 | 1 | 1 | 1 | 1 | 1 | 1 | 1 | 1 | 1 | 1 | 1 | 1 | 1 | 1 | 1 | 1 | 1 | 1 | 1 | 1 | 0 | 1 | 1 | 1 | 1 | 1 | 1 | 1 | 1 | 1 | 1 | 1 | 1 | 1 | 1 | 1 | 1 | 1 | 1 | 1 | 1 | 0 | 1 | 1 | 1 | 1 | 1 | 1 | 1 | 1 | 1 | 1 | 1 | 1 | 1 | 0 | 1 | 1 | 1 |
| *Bulbophyllum hirundinis* (Gagnep.) Seidenf. 1 | 1 | 1 | 1 | 1 | 1 | 1 | 1 | 1 | 1 | 1 | 1 | 1 | 1 | 1 | 1 | 1 | 1 | 1 | 1 | 1 | 1 | 1 | 1 | 1 | 1 | 1 | 1 | 1 | 1 | 1 | 1 | 1 | 1 | 1 | 1 | 1 | 1 | 1 | 1 | 1 | 1 | 1 | 1 | 1 | 1 | 1 | 1 | 1 | 1 | 1 | 1 | 1 | 1 | 1 | 1 | 1 | 1 | 1 | 1 | 1 | 1 | 1 | 1 |
| *Bulbophyllum hirundinis* (Gagnep.) Seidenf. 2 | 1 | 1 | 1 | 1 | 1 | 1 | 1 | 1 | 1 | 1 | 1 | 1 | 1 | 1 | 1 | 1 | 1 | 1 | 1 | 1 | 1 | 1 | 1 | 1 | 1 | 1 | 1 | 1 | 1 | 1 | 1 | 1 | 1 | 1 | 1 | 1 | 1 | 1 | 1 | 1 | 1 | 1 | 1 | 1 | 1 | 1 | 1 | 1 | 1 | 1 | 1 | 1 | 1 | 1 | 1 | 1 | 1 | 1 | 1 | 1 | 1 | 1 | 1 |
| *Bulbophyllum hystricinum* Schltr. | 1 | 1 | 1 | 1 | 1 | 1 | 1 | 1 | 1 | 1 | 1 | 1 | 1 | 1 | 1 | 1 | 1 | 1 | 1 | 1 | 1 | 1 | 1 | 1 | 1 | 1 | 1 | 1 | 1 | 1 | 1 | 1 | 1 | 1 | 1 | 1 | 1 | 1 | 1 | 1 | 1 | 1 | 1 | 1 | 1 | 1 | 1 | 1 | 1 | 1 | 1 | 1 | 1 | 1 | 1 | 1 | 1 | 1 | 1 | 0 | 1 | 1 | 1 |
| *Bulbophyllum icteranthum* Schltr. | 1 | 1 | 1 | 1 | 1 | 1 | 1 | 1 | 1 | 1 | 1 | 1 | 1 | 1 | 1 | 1 | 1 | 1 | 1 | 1 | 1 | 1 | 1 | 1 | 1 | 1 | 1 | 1 | 1 | 1 | 1 | 1 | 1 | 1 | 1 | 1 | 1 | 1 | 1 | 1 | 1 | 1 | 1 | 1 | 1 | 1 | 1 | 1 | 1 | 1 | 1 | 1 | 1 | 1 | 1 | 1 | 1 | 1 | 1 | 1 | 1 | 1 | 1 |
| *Bulbophyllum inconspicuum* Maxim. | 1 | 1 | 1 | 1 | 1 | 1 | 1 | 1 | 1 | 0 | 1 | 1 | 1 | 1 | 1 | 1 | 1 | 1 | 1 | 1 | 1 | 0 | 1 | 1 | 1 | 1 | 1 | 1 | 1 | 1 | 1 | 1 | 1 | 1 | 1 | 1 | 1 | 1 | 1 | 1 | 1 | 1 | 1 | 1 | 1 | 1 | 1 | 1 | 1 | 1 | 0 | 1 | 1 | 1 | 1 | 1 | 1 | 1 | 1 | 1 | 1 | 1 | 1 |
| *Bulbophyllum infundibuliforme* J.J.Sm. | 1 | 1 | 1 | 1 | 1 | 1 | 1 | 1 | 1 | 1 | 1 | 1 | 1 | 1 | 1 | 1 | 1 | 1 | 1 | 1 | 1 | 1 | 1 | 1 | 1 | 1 | 1 | 1 | 1 | 1 | 1 | 1 | 1 | 1 | 1 | 1 | 1 | 1 | 1 | 1 | 1 | 1 | 1 | 1 | 1 | 1 | 1 | 1 | 1 | 0 | 1 | 1 | 1 | 1 | 1 | 1 | 0 | 1 | 1 | 1 | 1 | 1 | 1 |
| *Bulbophyllum intricatum* Seidenf. | 1 | 1 | 1 | 1 | 1 | 1 | 1 | 1 | 1 | 1 | 1 | 1 | 1 | 1 | 1 | 1 | 1 | 1 | 1 | 1 | 1 | 1 | 1 | 1 | 1 | 1 | 1 | 1 | 1 | 1 | 1 | 1 | 1 | 1 | 1 | 1 | 1 | 1 | 1 | 1 | 1 | 1 | 0 | 1 | 1 | 1 | 1 | 1 | 1 | 1 | 1 | 1 | 1 | 1 | 1 | 1 | 1 | 1 | 1 | 1 | 1 | 1 | 1 |
| *Bulbophyllum jimcootesii* (Cabactulan, M.Leon & R.B.Pimentel) R.Rice | 1 | 1 | 1 | 1 | 1 | 1 | 1 | 1 | 1 | 1 | 1 | 1 | 1 | 1 | 1 | 1 | 1 | 1 | 1 | 1 | 1 | 1 | 1 | 1 | 1 | 1 | 1 | 1 | 1 | 1 | 1 | 1 | 1 | 1 | 1 | 1 | 1 | 1 | 1 | 1 | 1 | 1 | 1 | 1 | 1 | 1 | 1 | 1 | 1 | 0 | 1 | 1 | 1 | 1 | 1 | 1 | 0 | 1 | 1 | 1 | 1 | 1 | 1 |
| *Bulbophyllum johnsonii* T.E.Hunt 1 | 1 | 1 | 1 | 1 | 1 | 1 | 1 | 1 | 1 | 1 | 1 | 1 | 1 | 0 | 1 | 1 | 1 | 1 | 1 | 1 | 1 | 1 | 1 | 1 | 1 | 1 | 1 | 1 | 1 | 1 | 1 | 0 | 1 | 1 | 1 | 1 | 1 | 1 | 1 | 0 | 1 | 1 | 0 | 0 | 1 | 1 | 1 | 1 | 1 | 0 | 1 | 1 | 1 | 1 | 1 | 1 | 0 | 1 | 1 | 1 | 1 | 1 | 1 |
| *Bulbophyllum johnsonii* T.E.Hunt 2 | 0 | 1 | 1 | 1 | 0 | 1 | 1 | 1 | 1 | 1 | 1 | 0 | 1 | 0 | 0 | 1 | 1 | 1 | 0 | 0 | 1 | 0 | 1 | 0 | 0 | 1 | 0 | 1 | 0 | 1 | 0 | 0 | 1 | 0 | 1 | 0 | 1 | 0 | 1 | 1 | 1 | 1 | 0 | 1 | 1 | 1 | 1 | 1 | 0 | 1 | 1 | 1 | 1 | 1 | 1 | 1 | 0 | 0 | 1 | 1 | 0 | 1 | 1 |
| *Bulbophyllum kauloense* Schltr. | 1 | 1 | 1 | 1 | 1 | 1 | 1 | 1 | 1 | 1 | 1 | 1 | 1 | 1 | 1 | 1 | 1 | 1 | 1 | 1 | 1 | 1 | 1 | 1 | 1 | 1 | 1 | 1 | 1 | 1 | 1 | 1 | 1 | 1 | 1 | 1 | 1 | 1 | 1 | 1 | 1 | 1 | 1 | 1 | 1 | 0 | 1 | 1 | 1 | 1 | 1 | 0 | 1 | 1 | 1 | 1 | 1 | 1 | 1 | 1 | 1 | 1 | 1 |
| *Bulbophyllum keekee* N.Halle | 1 | 1 | 1 | 1 | 1 | 1 | 1 | 0 | 0 | 1 | 1 | 0 | 1 | 1 | 0 | 1 | 1 | 1 | 0 | 0 | 1 | 1 | 1 | 0 | 0 | 0 | 1 | 1 | 1 | 0 | 1 | 1 | 1 | 1 | 1 | 1 | 1 | 1 | 1 | 1 | 1 | 0 | 1 | 1 | 1 | 0 | 1 | 0 | 0 | 1 | 0 | 0 | 0 | 1 | 0 | 1 | 1 | 1 | 0 | 0 | 1 | 1 | 0 |
| *Bulbophyllum klabatense* Schltr. | 1 | 1 | 1 | 1 | 1 | 1 | 1 | 1 | 1 | 1 | 1 | 1 | 1 | 1 | 1 | 1 | 1 | 1 | 1 | 1 | 1 | 1 | 1 | 1 | 1 | 1 | 1 | 1 | 1 | 1 | 1 | 1 | 1 | 1 | 1 | 1 | 1 | 1 | 1 | 1 | 1 | 1 | 1 | 1 | 1 | 1 | 1 | 1 | 1 | 1 | 1 | 1 | 1 | 1 | 1 | 1 | 1 | 1 | 1 | 1 | 1 | 1 | 1 |
| *Bulbophyllum klabatense* subsp. *sulawesii* (Garay, Hamer & Siegerist) J.J.Verm. & P.O'Byrne | 1 | 1 | 1 | 1 | 1 | 1 | 1 | 1 | 1 | 1 | 1 | 1 | 1 | 0 | 1 | 1 | 1 | 1 | 0 | 1 | 1 | 1 | 1 | 1 | 1 | 1 | 1 | 1 | 1 | 1 | 1 | 1 | 1 | 1 | 1 | 1 | 1 | 1 | 1 | 1 | 1 | 1 | 1 | 1 | 1 | 1 | 1 | 1 | 1 | 0 | 1 | 1 | 1 | 1 | 1 | 1 | 0 | 1 | 1 | 1 | 1 | 1 | 1 |
| *Bulbophyllum kubahense* J.J.Verm. & A.Lamb | 1 | 1 | 0 | 1 | 1 | 1 | 1 | 1 | 1 | 1 | 1 | 1 | 1 | 1 | 1 | 1 | 1 | 1 | 1 | 1 | 1 | 0 | 1 | 1 | 1 | 1 | 1 | 1 | 1 | 1 | 1 | 1 | 1 | 1 | 1 | 1 | 1 | 1 | 1 | 1 | 1 | 1 | 1 | 1 | 1 | 1 | 1 | 1 | 1 | 1 | 1 | 1 | 1 | 1 | 1 | 1 | 1 | 1 | 1 | 1 | 1 | 1 | 1 |
| *Bulbophyllum lageniforme* F.M.Bailey | 1 | 1 | 1 | 1 | 1 | 1 | 1 | 1 | 1 | 1 | 1 | 1 | 1 | 1 | 1 | 1 | 1 | 1 | 1 | 1 | 1 | 1 | 1 | 1 | 1 | 1 | 1 | 1 | 1 | 1 | 1 | 1 | 1 | 1 | 1 | 1 | 1 | 1 | 1 | 1 | 1 | 1 | 0 | 1 | 1 | 1 | 1 | 0 | 0 | 1 | 1 | 1 | 1 | 1 | 1 | 1 | 1 | 1 | 0 | 1 | 1 | 1 | 1 |
| *Bulbophyllum lamingtonense* D.L.Jones | 1 | 1 | 1 | 1 | 1 | 1 | 1 | 1 | 1 | 1 | 1 | 1 | 1 | 1 | 1 | 1 | 1 | 1 | 1 | 1 | 1 | 1 | 1 | 1 | 1 | 1 | 1 | 1 | 1 | 1 | 1 | 1 | 1 | 1 | 1 | 1 | 1 | 1 | 1 | 1 | 1 | 1 | 1 | 1 | 1 | 1 | 1 | 1 | 1 | 1 | 1 | 1 | 1 | 1 | 1 | 1 | 1 | 1 | 1 | 1 | 1 | 1 | 1 |
| *Bulbophyllum lasianthum* Lindl. | 1 | 1 | 1 | 1 | 1 | 1 | 1 | 1 | 1 | 1 | 1 | 1 | 1 | 1 | 1 | 1 | 1 | 1 | 1 | 1 | 1 | 1 | 1 | 1 | 1 | 1 | 1 | 1 | 1 | 1 | 1 | 1 | 1 | 1 | 1 | 1 | 1 | 1 | 1 | 1 | 1 | 1 | 1 | 1 | 1 | 1 | 1 | 1 | 1 | 1 | 1 | 1 | 1 | 1 | 1 | 1 | 1 | 1 | 1 | 0 | 1 | 1 | 1 |
| *Bulbophyllum lasiochilum* Parish & Rchb.f. | 1 | 1 | 1 | 1 | 1 | 1 | 1 | 1 | 1 | 1 | 1 | 0 | 1 | 1 | 1 | 1 | 1 | 0 | 1 | 1 | 1 | 1 | 1 | 0 | 1 | 1 | 1 | 1 | 1 | 1 | 1 | 1 | 1 | 1 | 0 | 1 | 1 | 1 | 1 | 1 | 1 | 1 | 0 | 1 | 1 | 1 | 1 | 1 | 1 | 1 | 1 | 1 | 0 | 1 | 1 | 1 | 1 | 0 | 1 | 0 | 1 | 1 | 0 |
| *Bulbophyllum lasioglossum* Rolfe ex Ames | 1 | 1 | 1 | 1 | 1 | 1 | 1 | 1 | 1 | 1 | 1 | 1 | 1 | 0 | 1 | 1 | 1 | 1 | 1 | 1 | 1 | 1 | 1 | 1 | 0 | 1 | 1 | 1 | 1 | 1 | 1 | 1 | 1 | 1 | 1 | 1 | 1 | 1 | 1 | 1 | 1 | 0 | 0 | 1 | 1 | 1 | 1 | 1 | 0 | 1 | 1 | 1 | 1 | 1 | 0 | 0 | 1 | 1 | 0 | 1 | 1 | 1 | 1 |
| *Bulbophyllum laxiflorum* (Blume) Lindl. 1 | 1 | 1 | 1 | 1 | 1 | 1 | 1 | 1 | 1 | 1 | 1 | 1 | 1 | 1 | 1 | 1 | 1 | 1 | 1 | 1 | 1 | 1 | 1 | 1 | 1 | 1 | 1 | 1 | 1 | 1 | 1 | 1 | 1 | 1 | 1 | 1 | 1 | 1 | 1 | 1 | 1 | 1 | 1 | 1 | 1 | 1 | 1 | 1 | 1 | 1 | 0 | 1 | 1 | 1 | 1 | 1 | 1 | 1 | 1 | 0 | 1 | 1 | 1 |
| *Bulbophyllum laxiflorum* (Blume) Lindl. 2 | 1 | 1 | 1 | 1 | 1 | 1 | 1 | 1 | 1 | 1 | 1 | 1 | 1 | 1 | 1 | 1 | 1 | 1 | 1 | 1 | 1 | 1 | 1 | 1 | 1 | 1 | 1 | 1 | 1 | 1 | 1 | 1 | 1 | 1 | 1 | 1 | 1 | 1 | 1 | 1 | 1 | 1 | 1 | 1 | 1 | 1 | 1 | 1 | 1 | 1 | 1 | 1 | 1 | 1 | 1 | 1 | 1 | 1 | 1 | 1 | 1 | 1 | 1 |
| *Bulbophyllum lemniscatoides* Rolfe | 1 | 1 | 1 | 1 | 1 | 1 | 1 | 1 | 1 | 1 | 1 | 1 | 1 | 1 | 1 | 1 | 1 | 1 | 1 | 1 | 1 | 1 | 1 | 1 | 1 | 1 | 1 | 1 | 1 | 1 | 1 | 1 | 1 | 1 | 1 | 1 | 1 | 1 | 1 | 1 | 1 | 1 | 0 | 1 | 1 | 1 | 1 | 1 | 1 | 1 | 1 | 1 | 1 | 1 | 1 | 1 | 1 | 1 | 1 | 1 | 1 | 1 | 1 |
| *Bulbophyllum lepidum* (Blume) J.J.Sm. | 1 | 1 | 1 | 1 | 1 | 1 | 1 | 1 | 1 | 1 | 1 | 1 | 1 | 1 | 1 | 1 | 1 | 1 | 1 | 1 | 1 | 1 | 1 | 1 | 1 | 1 | 1 | 1 | 1 | 1 | 1 | 1 | 1 | 1 | 1 | 1 | 1 | 1 | 1 | 1 | 1 | 1 | 1 | 1 | 1 | 1 | 1 | 1 | 1 | 1 | 1 | 1 | 1 | 1 | 1 | 1 | 1 | 1 | 1 | 1 | 1 | 1 | 1 |
| *Bulbophyllum leptoleucum* Schltr. | 1 | 1 | 1 | 1 | 1 | 1 | 1 | 1 | 1 | 1 | 1 | 1 | 1 | 1 | 1 | 1 | 1 | 1 | 1 | 1 | 1 | 1 | 1 | 1 | 1 | 1 | 1 | 1 | 1 | 1 | 1 | 1 | 1 | 1 | 1 | 1 | 1 | 1 | 1 | 1 | 1 | 1 | 1 | 1 | 1 | 1 | 1 | 1 | 1 | 1 | 1 | 1 | 1 | 1 | 1 | 1 | 1 | 1 | 1 | 1 | 1 | 1 | 1 |
| *Bulbophyllum leucothyrsus* Schltr. | 1 | 1 | 1 | 1 | 1 | 1 | 1 | 1 | 1 | 1 | 1 | 1 | 1 | 1 | 1 | 1 | 1 | 1 | 1 | 1 | 1 | 1 | 1 | 1 | 1 | 1 | 1 | 1 | 1 | 1 | 1 | 1 | 1 | 1 | 1 | 1 | 1 | 1 | 1 | 1 | 0 | 1 | 0 | 1 | 1 | 1 | 1 | 1 | 1 | 1 | 1 | 1 | 1 | 1 | 1 | 1 | 1 | 1 | 1 | 1 | 1 | 1 | 1 |
| *Bulbophyllum levanae* Ames | 1 | 1 | 1 | 1 | 1 | 1 | 1 | 1 | 1 | 1 | 1 | 1 | 1 | 1 | 1 | 1 | 1 | 1 | 1 | 1 | 1 | 1 | 1 | 1 | 1 | 1 | 1 | 1 | 1 | 1 | 1 | 1 | 1 | 1 | 1 | 1 | 1 | 1 | 1 | 1 | 1 | 1 | 1 | 1 | 1 | 1 | 1 | 1 | 1 | 1 | 1 | 1 | 1 | 1 | 1 | 1 | 1 | 1 | 1 | 1 | 1 | 1 | 1 |
| *Bulbophyllum levatii* Kraenzl. | 1 | 1 | 1 | 1 | 1 | 1 | 1 | 0 | 1 | 1 | 1 | 1 | 1 | 1 | 1 | 1 | 1 | 1 | 1 | 1 | 0 | 1 | 1 | 1 | 1 | 1 | 1 | 1 | 1 | 1 | 1 | 1 | 1 | 1 | 1 | 1 | 1 | 1 | 1 | 0 | 1 | 1 | 1 | 0 | 0 | 1 | 1 | 1 | 1 | 1 | 1 | 1 | 1 | 0 | 1 | 1 | 0 | 1 | 0 | 1 | 1 | 1 | 1 |
| *Bulbophyllum levyae* Garay, Hamer & Siegerist | 1 | 1 | 1 | 1 | 1 | 1 | 1 | 1 | 1 | 1 | 1 | 1 | 1 | 1 | 1 | 1 | 1 | 1 | 1 | 1 | 1 | 1 | 1 | 1 | 1 | 1 | 1 | 1 | 1 | 1 | 1 | 1 | 1 | 1 | 1 | 1 | 1 | 1 | 1 | 1 | 1 | 1 | 1 | 1 | 1 | 1 | 1 | 1 | 1 | 0 | 1 | 1 | 1 | 1 | 1 | 1 | 0 | 1 | 1 | 1 | 1 | 1 | 1 |
| *Bulbophyllum lewisense* B.Gray & D.L.Jones | 1 | 1 | 1 | 1 | 1 | 1 | 0 | 0 | 1 | 1 | 1 | 1 | 1 | 1 | 1 | 0 | 0 | 1 | 1 | 1 | 0 | 1 | 0 | 1 | 1 | 1 | 0 | 0 | 1 | 1 | 1 | 1 | 1 | 1 | 1 | 1 | 1 | 1 | 1 | 1 | 1 | 1 | 1 | 1 | 0 | 0 | 1 | 1 | 1 | 1 | 0 | 0 | 0 | 1 | 1 | 1 | 1 | 0 | 0 | 1 | 0 | 1 | 1 |
| *Bulbophyllum lilacinum* Ridl. | 1 | 1 | 1 | 1 | 1 | 1 | 1 | 1 | 1 | 1 | 1 | 1 | 1 | 1 | 1 | 1 | 1 | 1 | 1 | 1 | 1 | 1 | 1 | 1 | 1 | 1 | 1 | 1 | 1 | 1 | 1 | 1 | 1 | 1 | 1 | 1 | 1 | 1 | 1 | 1 | 1 | 1 | 1 | 1 | 1 | 1 | 1 | 1 | 1 | 1 | 1 | 1 | 1 | 1 | 1 | 1 | 1 | 1 | 1 | 1 | 1 | 1 | 1 |
| *Bulbophyllum lilianae* Rendle | 1 | 1 | 1 | 1 | 1 | 1 | 1 | 1 | 1 | 1 | 1 | 1 | 1 | 1 | 1 | 1 | 1 | 1 | 0 | 0 | 1 | 1 | 1 | 1 | 1 | 1 | 0 | 1 | 1 | 1 | 1 | 1 | 1 | 1 | 1 | 1 | 1 | 1 | 1 | 1 | 1 | 1 | 0 | 1 | 1 | 1 | 1 | 1 | 1 | 1 | 1 | 1 | 0 | 1 | 1 | 1 | 1 | 0 | 0 | 1 | 1 | 1 | 1 |

| | | | | | | | | | | | | | | | | | | | | | | | | | | | | | | | | | | | | | | | | | | | | | | | | | | | | | | | | | | | | | | | |
|---|---|---|---|---|---|---|---|---|---|---|---|---|---|---|---|---|---|---|---|---|---|---|---|---|---|---|---|---|---|---|---|---|---|---|---|---|---|---|---|---|---|---|---|---|---|---|---|---|---|---|---|---|---|---|---|---|---|---|---|---|---|---|---|
| *Bulbophyllum lindleyanum* Griff. | 1 | 1 | 1 | 1 | 1 | 1 | 1 | 1 | 1 | 1 | 1 | 1 | 1 | 1 | 1 | 1 | 1 | 1 | 1 | 1 | 1 | 1 | 1 | 1 | 1 | 1 | 1 | 1 | 1 | 1 | 1 | 1 | 1 | 1 | 1 | 1 | 1 | 1 | 1 | 1 | 1 | 1 | 1 | 1 | 1 | 1 | 1 | 1 | 1 | 1 | 1 | 1 | 1 | 1 | 1 | 1 | 1 | 1 | 1 | 1 | 1 | 1 | 1 |
| *Bulbophyllum lineolatum* Schltr. | 1 | 1 | 1 | 1 | 1 | 1 | 1 | 0 | 1 | 1 | 1 | 1 | 1 | 1 | 1 | 1 | 1 | 1 | 1 | 1 | 0 | 1 | 1 | 1 | 1 | 1 | 1 | 1 | 1 | 1 | 1 | 1 | 1 | 1 | 1 | 1 | 1 | 1 | 1 | 0 | 1 | 1 | 0 | 0 | 0 | 1 | 1 | 1 | 1 | 1 | 1 | 1 | 1 | 0 | 1 | 1 | 0 | 1 | 0 | 1 | 1 | 1 | 1 |
| *Bulbophyllum lingulatum* Rendle | 1 | 1 | 1 | 1 | 1 | 1 | 1 | 1 | 1 | 1 | 1 | 1 | 1 | 1 | 1 | 1 | 1 | 1 | 1 | 1 | 1 | 1 | 1 | 1 | 1 | 1 | 1 | 1 | 1 | 1 | 1 | 1 | 1 | 1 | 1 | 1 | 1 | 1 | 1 | 1 | 1 | 1 | 1 | 1 | 1 | 1 | 1 | 1 | 1 | 1 | 1 | 1 | 1 | 1 | 1 | 1 | 1 | 1 | 1 | 1 | 1 | 1 | 1 |
| *Bulbophyllum lobbii* Lindl. 1 | 1 | 1 | 1 | 1 | 1 | 1 | 1 | 1 | 1 | 1 | 1 | 1 | 1 | 1 | 1 | 1 | 1 | 1 | 1 | 1 | 1 | 1 | 1 | 1 | 1 | 1 | 1 | 1 | 1 | 1 | 1 | 1 | 1 | 1 | 1 | 1 | 1 | 1 | 1 | 1 | 1 | 1 | 0 | 1 | 1 | 1 | 1 | 1 | 1 | 1 | 1 | 1 | 1 | 1 | 1 | 1 | 1 | 1 | 1 | 1 | 1 | 1 | 1 |
| *Bulbophyllum lobbii* Lindl. 2 | 1 | 1 | 1 | 1 | 1 | 1 | 1 | 1 | 1 | 1 | 1 | 1 | 1 | 1 | 1 | 1 | 1 | 1 | 1 | 1 | 1 | 1 | 1 | 1 | 1 | 1 | 1 | 1 | 1 | 1 | 1 | 1 | 1 | 1 | 1 | 1 | 1 | 1 | 1 | 1 | 1 | 1 | 1 | 1 | 1 | 1 | 1 | 1 | 1 | 1 | 1 | 1 | 1 | 1 | 1 | 1 | 1 | 1 | 1 | 1 | 1 | 1 | 1 |
| *Bulbophyllum lobbii* Lindl. 3 | 1 | 1 | 1 | 1 | 1 | 1 | 1 | 1 | 1 | 1 | 1 | 1 | 1 | 1 | 1 | 1 | 1 | 1 | 1 | 1 | 1 | 1 | 1 | 1 | 1 | 1 | 1 | 1 | 1 | 1 | 1 | 1 | 1 | 1 | 1 | 1 | 1 | 1 | 1 | 1 | 1 | 1 | 1 | 1 | 1 | 1 | 1 | 1 | 1 | 1 | 1 | 1 | 1 | 1 | 1 | 1 | 1 | 1 | 1 | 1 | 1 | 1 | 1 |
| *Bulbophyllum lobbii* subsp. *siamense* (Rchb.f.) Mangal, F.Velazquez & J.J.Verm. | 1 | 1 | 1 | 1 | 1 | 1 | 1 | 1 | 1 | 1 | 1 | 1 | 1 | 1 | 1 | 1 | 1 | 1 | 1 | 1 | 1 | 1 | 1 | 1 | 1 | 1 | 1 | 1 | 1 | 1 | 1 | 1 | 1 | 1 | 1 | 1 | 1 | 1 | 1 | 1 | 1 | 1 | 1 | 1 | 1 | 1 | 1 | 1 | 1 | 1 | 1 | 1 | 1 | 1 | 1 | 1 | 1 | 1 | 1 | 1 | 1 | 1 | 1 |
| *Bulbophyllum longibrachiatum* Z.H.Tsi | 1 | 1 | 1 | 1 | 1 | 1 | 1 | 1 | 1 | 1 | 1 | 1 | 1 | 1 | 1 | 1 | 1 | 1 | 1 | 1 | 1 | 1 | 1 | 1 | 1 | 1 | 1 | 1 | 1 | 1 | 1 | 1 | 1 | 1 | 1 | 1 | 1 | 1 | 1 | 1 | 1 | 1 | 1 | 1 | 1 | 1 | 1 | 1 | 1 | 1 | 1 | 1 | 1 | 1 | 1 | 1 | 1 | 1 | 1 | 1 | 1 | 1 | 1 |
| *Bulbophyllum longiflorum* Thouars | 1 | 1 | 1 | 1 | 1 | 1 | 1 | 1 | 1 | 1 | 1 | 1 | 1 | 1 | 1 | 1 | 1 | 1 | 1 | 1 | 1 | 1 | 1 | 1 | 1 | 1 | 1 | 1 | 1 | 1 | 1 | 1 | 1 | 1 | 1 | 1 | 1 | 1 | 1 | 1 | 1 | 1 | 0 | 1 | 1 | 1 | 1 | 1 | 1 | 1 | 1 | 1 | 1 | 1 | 1 | 1 | 1 | 1 | 1 | 0 | 1 | 1 | 1 |
| *Bulbophyllum longisepalum* Rolfe | 1 | 1 | 1 | 1 | 1 | 1 | 1 | 1 | 1 | 1 | 1 | 1 | 1 | 1 | 1 | 1 | 1 | 1 | 1 | 1 | 1 | 1 | 1 | 1 | 1 | 1 | 1 | 1 | 1 | 1 | 1 | 1 | 1 | 1 | 1 | 1 | 1 | 1 | 1 | 1 | 1 | 1 | 1 | 1 | 1 | 1 | 1 | 1 | 1 | 0 | 1 | 1 | 1 | 1 | 1 | 1 | 0 | 1 | 1 | 1 | 1 | 1 | 1 |
| *Bulbophyllum lophoglottis* (Guillaumin) N.Hallé | 1 | 1 | 1 | 1 | 1 | 1 | 1 | 1 | 1 | 1 | 1 | 1 | 1 | 1 | 1 | 1 | 1 | 1 | 1 | 1 | 1 | 1 | 1 | 1 | 1 | 1 | 1 | 1 | 1 | 1 | 1 | 1 | 0 | 1 | 1 | 1 | 1 | 1 | 1 | 1 | 1 | 1 | 1 | 1 | 1 | 1 | 1 | 1 | 1 | 1 | 1 | 1 | 1 | 1 | 1 | 1 | 0 | 1 | 1 | 1 | 1 | 1 | 1 |
| *Bulbophyllum lyriforme* J.J.Verm. & P.O'Byrne | 1 | 1 | 1 | 1 | 1 | 1 | 1 | 1 | 1 | 1 | 1 | 1 | 1 | 1 | 1 | 1 | 1 | 1 | 1 | 1 | 1 | 1 | 1 | 1 | 1 | 1 | 1 | 1 | 1 | 1 | 1 | 1 | 1 | 1 | 1 | 1 | 1 | 1 | 1 | 1 | 1 | 1 | 1 | 1 | 1 | 1 | 1 | 1 | 1 | 1 | 1 | 1 | 1 | 1 | 1 | 1 | 1 | 1 | 1 | 1 | 1 | 1 | 1 |
| *Bulbophyllum macphersonii* Rupp 1 | 1 | 1 | 1 | 1 | 1 | 1 | 1 | 1 | 1 | 1 | 1 | 1 | 1 | 1 | 1 | 1 | 1 | 1 | 1 | 1 | 1 | 1 | 1 | 1 | 1 | 1 | 1 | 1 | 1 | 1 | 1 | 1 | 1 | 1 | 1 | 1 | 1 | 1 | 1 | 1 | 1 | 1 | 0 | 1 | 1 | 1 | 1 | 1 | 1 | 1 | 1 | 1 | 1 | 1 | 1 | 1 | 1 | 1 | 1 | 1 | 1 | 1 | 1 |
| *Bulbophyllum macphersonii* Rupp 2 | 0 | 1 | 1 | 1 | 1 | 0 | 1 | 1 | 0 | 1 | 1 | 1 | 1 | 0 | 1 | 1 | 1 | 1 | 1 | 0 | 1 | 0 | 1 | 0 | 0 | 1 | 0 | 1 | 0 | 1 | 0 | 0 | 1 | 0 | 1 | 0 | 1 | 0 | 1 | 1 | 1 | 0 | 0 | 1 | 1 | 1 | 1 | 1 | 1 | 1 | 1 | 1 | 1 | 1 | 1 | 1 | 1 | 0 | 1 | 1 | 1 | 1 | 1 |
| *Bulbophyllum macraei* (Lindl.) Rchb.f. | 1 | 1 | 1 | 1 | 1 | 1 | 1 | 1 | 1 | 1 | 1 | 1 | 1 | 1 | 1 | 1 | 1 | 1 | 1 | 1 | 1 | 1 | 1 | 1 | 1 | 1 | 1 | 1 | 1 | 1 | 1 | 1 | 1 | 1 | 1 | 1 | 1 | 1 | 1 | 1 | 1 | 1 | 0 | 1 | 1 | 1 | 1 | 1 | 1 | 1 | 1 | 1 | 1 | 1 | 1 | 1 | 1 | 1 | 1 | 1 | 1 | 1 | 1 |
| *Bulbophyllum macranthum* Lindl. | 1 | 1 | 1 | 1 | 1 | 1 | 1 | 1 | 1 | 1 | 1 | 1 | 1 | 1 | 1 | 1 | 1 | 1 | 1 | 1 | 1 | 1 | 1 | 1 | 1 | 1 | 1 | 1 | 1 | 1 | 1 | 1 | 1 | 1 | 1 | 1 | 1 | 1 | 1 | 1 | 1 | 1 | 0 | 1 | 1 | 1 | 1 | 1 | 1 | 1 | 1 | 1 | 1 | 1 | 1 | 1 | 1 | 1 | 1 | 1 | 1 | 1 | 1 |
| *Bulbophyllum macrorhopalon* Schltr. | 1 | 1 | 1 | 1 | 1 | 1 | 1 | 1 | 1 | 0 | 1 | 1 | 1 | 1 | 1 | 1 | 1 | 1 | 1 | 1 | 1 | 1 | 1 | 1 | 1 | 1 | 1 | 1 | 1 | 1 | 1 | 1 | 1 | 1 | 1 | 1 | 1 | 1 | 1 | 1 | 1 | 1 | 1 | 1 | 1 | 1 | 1 | 1 | 1 | 0 | 1 | 1 | 1 | 1 | 1 | 1 | 1 | 1 | 1 | 1 | 1 | 1 | 1 |
| *Bulbophyllum macrourum* Schltr. | 1 | 1 | 1 | 1 | 1 | 1 | 1 | 1 | 1 | 1 | 1 | 1 | 1 | 1 | 1 | 1 | 1 | 1 | 1 | 1 | 1 | 1 | 1 | 1 | 1 | 1 | 1 | 1 | 1 | 1 | 1 | 1 | 1 | 1 | 1 | 1 | 1 | 1 | 1 | 1 | 1 | 1 | 1 | 1 | 1 | 1 | 1 | 1 | 1 | 1 | 1 | 1 | 1 | 1 | 1 | 1 | 1 | 1 | 1 | 1 | 1 | 1 | 1 |
| *Bulbophyllum makoyanum* (Rchb.f.) Ridl. | 1 | 1 | 1 | 1 | 1 | 1 | 1 | 1 | 1 | 1 | 1 | 1 | 1 | 1 | 1 | 1 | 1 | 1 | 1 | 1 | 1 | 1 | 1 | 1 | 1 | 1 | 1 | 1 | 1 | 1 | 1 | 1 | 1 | 1 | 1 | 1 | 1 | 1 | 1 | 1 | 1 | 1 | 1 | 1 | 1 | 1 | 1 | 1 | 1 | 1 | 1 | 1 | 1 | 1 | 1 | 1 | 1 | 1 | 1 | 1 | 1 | 1 | 1 |
| *Bulbophyllum mandibulare* Rchb.f. | 1 | 1 | 1 | 1 | 1 | 1 | 1 | 1 | 0 | 1 | 1 | 1 | 1 | 1 | 1 | 1 | 1 | 1 | 1 | 1 | 1 | 1 | 1 | 1 | 1 | 1 | 1 | 1 | 1 | 1 | 1 | 1 | 1 | 1 | 1 | 1 | 1 | 1 | 1 | 1 | 1 | 1 | 1 | 1 | 1 | 1 | 1 | 1 | 1 | 1 | 1 | 1 | 1 | 1 | 1 | 1 | 1 | 1 | 1 | 1 | 1 | 1 | 1 |
| *Bulbophyllum maquilingense* Ames & Quisumb. | 1 | 1 | 1 | 1 | 1 | 1 | 1 | 1 | 1 | 1 | 1 | 1 | 1 | 1 | 1 | 1 | 1 | 1 | 1 | 1 | 1 | 1 | 1 | 1 | 1 | 1 | 1 | 1 | 1 | 1 | 1 | 1 | 1 | 1 | 1 | 1 | 1 | 1 | 1 | 1 | 1 | 1 | 1 | 1 | 1 | 1 | 1 | 1 | 1 | 1 | 1 | 1 | 1 | 1 | 1 | 1 | 1 | 1 | 1 | 1 | 1 | 1 | 1 |
| *Bulbophyllum marknaivei* Cootes, R.Boos & M.Leon | 1 | 1 | 1 | 1 | 1 | 1 | 1 | 1 | 1 | 1 | 1 | 1 | 1 | 1 | 1 | 1 | 1 | 1 | 1 | 1 | 1 | 1 | 1 | 1 | 1 | 1 | 1 | 1 | 1 | 1 | 1 | 1 | 1 | 1 | 1 | 1 | 1 | 1 | 1 | 1 | 1 | 1 | 1 | 1 | 1 | 1 | 1 | 1 | 1 | 0 | 1 | 1 | 1 | 1 | 1 | 1 | 1 | 1 | 1 | 1 | 1 | 1 | 1 |
| *Bulbophyllum mastersianum* (Rolfe) J.J.Sm. | 1 | 1 | 1 | 1 | 1 | 1 | 1 | 1 | 1 | 1 | 1 | 1 | 1 | 1 | 1 | 1 | 1 | 1 | 1 | 1 | 1 | 1 | 1 | 1 | 1 | 1 | 1 | 1 | 1 | 1 | 1 | 1 | 1 | 1 | 1 | 1 | 1 | 1 | 1 | 1 | 1 | 1 | 1 | 1 | 1 | 1 | 1 | 1 | 1 | 1 | 1 | 1 | 1 | 1 | 1 | 1 | 1 | 1 | 1 | 1 | 1 | 1 | 1 |
| *Bulbophyllum maxillare* Rchb.f. | 1 | 1 | 1 | 1 | 1 | 1 | 1 | 1 | 1 | 1 | 1 | 1 | 1 | 1 | 1 | 1 | 1 | 1 | 1 | 1 | 1 | 1 | 1 | 1 | 1 | 1 | 1 | 1 | 1 | 1 | 1 | 1 | 1 | 1 | 1 | 1 | 1 | 1 | 1 | 1 | 1 | 1 | 0 | 1 | 1 | 1 | 1 | 1 | 1 | 1 | 1 | 1 | 1 | 1 | 1 | 1 | 1 | 1 | 1 | 1 | 1 | 1 | 1 |
| *Bulbophyllum maximum* (Lindl.) Rchb.f. | 1 | 1 | 1 | 1 | 1 | 1 | 1 | 1 | 1 | 1 | 1 | 1 | 1 | 1 | 0 | 1 | 1 | 1 | 1 | 1 | 1 | 1 | 1 | 1 | 1 | 1 | 1 | 1 | 1 | 1 | 1 | 1 | 1 | 1 | 1 | 1 | 1 | 1 | 1 | 1 | 1 | 1 | 1 | 1 | 1 | 1 | 1 | 1 | 1 | 1 | 1 | 1 | 1 | 1 | 1 | 1 | 1 | 1 | 1 | 0 | 1 | 1 | 1 |
| *Bulbophyllum mearnsii* Ames 1 | 1 | 1 | 1 | 1 | 1 | 1 | 1 | 1 | 1 | 1 | 1 | 1 | 1 | 1 | 1 | 1 | 1 | 1 | 1 | 1 | 1 | 1 | 1 | 1 | 1 | 1 | 1 | 1 | 1 | 1 | 1 | 1 | 1 | 1 | 1 | 1 | 1 | 1 | 1 | 1 | 1 | 1 | 0 | 1 | 1 | 1 | 1 | 1 | 1 | 1 | 1 | 1 | 1 | 1 | 1 | 1 | 1 | 1 | 1 | 1 | 1 | 1 | 1 |
| *Bulbophyllum mearnsii* Ames 2 | 1 | 1 | 1 | 1 | 1 | 1 | 1 | 1 | 1 | 1 | 1 | 1 | 1 | 1 | 1 | 1 | 1 | 1 | 1 | 1 | 1 | 1 | 1 | 1 | 1 | 1 | 1 | 1 | 1 | 1 | 1 | 1 | 1 | 1 | 1 | 1 | 1 | 1 | 1 | 1 | 1 | 0 | 1 | 1 | 1 | 1 | 1 | 1 | 1 | 1 | 1 | 1 | 1 | 1 | 1 | 1 | 1 | 1 | 1 | 1 | 1 | 1 | 1 |
| *Bulbophyllum medusae* Rchb.f. | 1 | 1 | 1 | 1 | 1 | 1 | 1 | 1 | 1 | 1 | 1 | 1 | 1 | 1 | 1 | 1 | 1 | 1 | 1 | 1 | 1 | 1 | 1 | 1 | 1 | 1 | 1 | 1 | 1 | 1 | 1 | 1 | 1 | 1 | 1 | 1 | 1 | 1 | 1 | 1 | 1 | 1 | 0 | 1 | 1 | 1 | 1 | 1 | 1 | 1 | 1 | 1 | 1 | 1 | 1 | 1 | 1 | 1 | 1 | 1 | 1 | 1 | 1 |
| *Bulbophyllum membranaceum* Teijsm. & Binn. | 1 | 1 | 1 | 1 | 1 | 1 | 1 | 1 | 1 | 1 | 1 | 1 | 1 | 1 | 1 | 1 | 1 | 1 | 1 | 1 | 1 | 1 | 1 | 1 | 1 | 1 | 1 | 1 | 1 | 1 | 1 | 1 | 1 | 1 | 1 | 1 | 1 | 1 | 1 | 1 | 1 | 1 | 1 | 1 | 1 | 1 | 1 | 1 | 1 | 1 | 1 | 1 | 1 | 1 | 1 | 1 | 1 | 1 | 1 | 1 | 1 | 1 | 1 |
| *Bulbophyllum membranifolium* Hook.f. | 0 | 1 | 1 | 1 | 1 | 1 | 1 | 1 | 1 | 1 | 1 | 1 | 1 | 1 | 1 | 1 | 1 | 1 | 1 | 1 | 1 | 1 | 1 | 1 | 1 | 1 | 1 | 1 | 1 | 1 | 1 | 1 | 1 | 1 | 1 | 1 | 1 | 1 | 1 | 1 | 1 | 1 | 1 | 1 | 1 | 1 | 1 | 1 | 1 | 1 | 0 | 1 | 1 | 1 | 1 | 1 | 1 | 1 | 1 | 1 | 1 | 1 | 1 |
| *Bulbophyllum mentosum* Barb.Rodr. | 1 | 1 | 1 | 1 | 1 | 1 | 1 | 1 | 1 | 1 | 1 | 1 | 1 | 1 | 1 | 1 | 1 | 1 | 1 | 1 | 1 | 1 | 1 | 1 | 1 | 1 | 1 | 1 | 1 | 1 | 1 | 1 | 1 | 1 | 1 | 1 | 1 | 1 | 1 | 1 | 1 | 1 | 1 | 1 | 1 | 1 | 1 | 1 | 1 | 1 | 1 | 1 | 1 | 1 | 1 | 1 | 0 | 1 | 1 | 1 | 1 | 1 | 1 |
| *Bulbophyllum meridense* Rchb.f. | 0 | 1 | 1 | 1 | 1 | 1 | 1 | 1 | 0 | 1 | 1 | 0 | 1 | 1 | 1 | 1 | 1 | 1 | 0 | 1 | 1 | 1 | 1 | 0 | 1 | 0 | 1 | 1 | 1 | 0 | 1 | 1 | 1 | 1 | 1 | 1 | 1 | 1 | 1 | 1 | 1 | 0 | 0 | 0 | 0 | 1 | 0 | 0 | 0 | 1 | 0 | 0 | 0 | 1 | 1 | 1 | 1 | 1 | 0 | 0 | 1 | 1 | 0 |
| *Bulbophyllum mimiense* Schltr. | 1 | 1 | 1 | 1 | 1 | 1 | 1 | 1 | 1 | 1 | 1 | 1 | 1 | 1 | 1 | 1 | 1 | 1 | 1 | 1 | 1 | 1 | 1 | 1 | 1 | 1 | 1 | 1 | 1 | 1 | 1 | 1 | 1 | 1 | 1 | 1 | 1 | 1 | 1 | 1 | 1 | 1 | 1 | 1 | 1 | 1 | 1 | 1 | 1 | 1 | 1 | 1 | 1 | 1 | 1 | 1 | 1 | 1 | 1 | 1 | 1 | 1 | 1 |
| *Bulbophyllum minutissimum* (F.Muell.) F.Muell. | 1 | 1 | 0 | 1 | 0 | 1 | 1 | 0 | 0 | 1 | 0 | 0 | 0 | 0 | 1 | 1 | 1 | 1 | 0 | 0 | 1 | 0 | 1 | 1 | 0 | 0 | 1 | 1 | 0 | 0 | 1 | 1 | 1 | 1 | 1 | 1 | 1 | 1 | 1 | 1 | 0 | 0 | 1 | 1 | 0 | 0 | 1 | 0 | 0 | 1 | 0 | 0 | 0 | 1 | 0 | 0 | 1 | 0 | 0 | 1 | 1 | 1 | 0 |
| *Bulbophyllum mirum* J.J.Sm. | 1 | 1 | 1 | 1 | 1 | 1 | 1 | 1 | 1 | 1 | 1 | 1 | 1 | 1 | 1 | 1 | 1 | 1 | 1 | 1 | 1 | 1 | 1 | 1 | 1 | 1 | 1 | 1 | 1 | 1 | 1 | 1 | 1 | 1 | 1 | 1 | 1 | 1 | 1 | 1 | 1 | 1 | 1 | 1 | 1 | 1 | 1 | 1 | 1 | 1 | 1 | 1 | 1 | 1 | 1 | 1 | 1 | 1 | 1 | 1 | 1 | 1 | 1 |
| *Bulbophyllum mucronatum* (Blume) Lindl. 1 | 1 | 1 | 1 | 1 | 1 | 1 | 1 | 1 | 1 | 1 | 1 | 1 | 1 | 1 | 1 | 1 | 1 | 1 | 1 | 1 | 1 | 1 | 1 | 1 | 1 | 1 | 1 | 1 | 1 | 1 | 1 | 1 | 1 | 1 | 1 | 1 | 1 | 1 | 1 | 1 | 1 | 0 | 1 | 1 | 1 | 1 | 1 | 1 | 1 | 1 | 1 | 1 | 1 | 1 | 1 | 1 | 1 | 1 | 1 | 1 | 1 | 1 | 1 |
| *Bulbophyllum mucronatum* (Blume) Lindl. 2 | 1 | 1 | 1 | 1 | 1 | 1 | 1 | 1 | 1 | 1 | 1 | 1 | 1 | 1 | 1 | 1 | 1 | 1 | 1 | 1 | 1 | 1 | 1 | 1 | 1 | 1 | 1 | 1 | 1 | 1 | 1 | 1 | 1 | 1 | 1 | 1 | 1 | 1 | 1 | 1 | 1 | 0 | 1 | 1 | 1 | 1 | 1 | 1 | 1 | 1 | 1 | 1 | 1 | 1 | 1 | 1 | 1 | 1 | 1 | 1 | 1 | 1 | 1 |
| *Bulbophyllum muricatum* J.J.Sm. | 1 | 1 | 1 | 1 | 1 | 1 | 1 | 1 | 1 | 1 | 1 | 1 | 1 | 1 | 1 | 1 | 1 | 1 | 1 | 1 | 1 | 1 | 1 | 1 | 1 | 1 | 1 | 1 | 1 | 1 | 1 | 1 | 1 | 1 | 1 | 1 | 1 | 1 | 1 | 1 | 1 | 1 | 1 | 1 | 1 | 1 | 1 | 1 | 1 | 0 | 1 | 1 | 1 | 1 | 1 | 1 | 1 | 1 | 1 | 1 | 1 | 1 | 1 |
| *Bulbophyllum nasica* Schltr. | 1 | 1 | 1 | 1 | 1 | 1 | 1 | 1 | 1 | 1 | 1 | 1 | 1 | 1 | 1 | 1 | 1 | 1 | 1 | 1 | 1 | 1 | 1 | 1 | 1 | 1 | 1 | 1 | 1 | 1 | 1 | 1 | 1 | 1 | 1 | 1 | 1 | 1 | 0 | 1 | 1 | 1 | 1 | 1 | 1 | 1 | 1 | 1 | 1 | 1 | 1 | 1 | 1 | 1 | 1 | 1 | 1 | 1 | 1 | 1 | 1 | 1 | 1 |
| *Bulbophyllum nasseri* Garay | 1 | 1 | 1 | 1 | 1 | 1 | 1 | 1 | 1 | 1 | 1 | 1 | 1 | 1 | 1 | 1 | 1 | 1 | 1 | 1 | 1 | 1 | 1 | 1 | 1 | 1 | 1 | 1 | 1 | 1 | 1 | 1 | 1 | 1 | 1 | 1 | 1 | 1 | 1 | 1 | 1 | 1 | 1 | 1 | 1 | 1 | 1 | 1 | 1 | 1 | 1 | 1 | 1 | 1 | 1 | 1 | 1 | 1 | 1 | 1 | 1 | 1 | 1 |
| *Bulbophyllum nasutum* Rchb.f. | 1 | 1 | 1 | 1 | 1 | 1 | 1 | 1 | 1 | 1 | 1 | 1 | 1 | 1 | 1 | 1 | 1 | 1 | 1 | 1 | 1 | 1 | 1 | 1 | 1 | 1 | 1 | 1 | 1 | 1 | 1 | 1 | 1 | 1 | 1 | 1 | 1 | 1 | 1 | 1 | 1 | 1 | 1 | 1 | 1 | 1 | 1 | 1 | 1 | 1 | 1 | 1 | 1 | 1 | 1 | 1 | 1 | 1 | 1 | 1 | 1 | 1 | 1 |
| *Bulbophyllum nematopodum* F.Muell. 1 | 1 | 1 | 1 | 1 | 1 | 1 | 1 | 1 | 1 | 1 | 1 | 1 | 1 | 1 | 1 | 1 | 1 | 1 | 1 | 1 | 1 | 1 | 1 | 1 | 1 | 1 | 1 | 1 | 1 | 1 | 1 | 1 | 1 | 1 | 1 | 1 | 1 | 1 | 1 | 1 | 1 | 1 | 1 | 1 | 1 | 1 | 1 | 1 | 1 | 1 | 1 | 1 | 1 | 1 | 1 | 1 | 1 | 1 | 1 | 1 | 1 | 1 | 1 |
| *Bulbophyllum nematopodum* F.Muell. 2 | 1 | 1 | 1 | 1 | 1 | 1 | 1 | 1 | 1 | 1 | 1 | 1 | 1 | 1 | 1 | 1 | 1 | 1 | 1 | 1 | 1 | 1 | 1 | 1 | 1 | 1 | 1 | 1 | 1 | 1 | 1 | 1 | 1 | 1 | 1 | 1 | 1 | 1 | 1 | 1 | 1 | 1 | 1 | 1 | 1 | 1 | 1 | 1 | 1 | 1 | 1 | 1 | 1 | 1 | 1 | 1 | 1 | 1 | 1 | 1 | 1 | 1 | 1 |
| *Bulbophyllum newportii* Rolfe | 1 | 1 | 1 | 0 | 1 | 1 | 1 | 1 | 1 | 1 | 1 | 1 | 1 | 1 | 1 | 1 | 1 | 1 | 1 | 1 | 1 | 1 | 1 | 1 | 0 | 1 | 1 | 1 | 1 | 1 | 1 | 1 | 1 | 1 | 1 | 1 | 1 | 1 | 1 | 1 | 1 | 0 | 1 | 1 | 1 | 1 | 1 | 1 | 1 | 1 | 1 | 1 | 1 | 1 | 1 | 1 | 1 | 1 | 1 | 1 | 1 | 1 | 1 |
| *Bulbophyllum ngoyense* Schltr. 1 | 1 | 1 | 1 | 1 | 1 | 1 | 1 | 0 | 0 | 1 | 1 | 1 | 0 | 1 | 1 | 1 | 1 | 1 | 1 | 1 | 1 | 1 | 1 | 1 | 1 | 1 | 0 | 1 | 1 | 1 | 1 | 1 | 1 | 1 | 1 | 1 | 1 | 1 | 1 | 1 | 1 | 1 | 1 | 1 | 1 | 1 | 1 | 1 | 1 | 1 | 1 | 1 | 1 | 0 | 1 | 1 | 0 | 0 | 1 | 1 | 1 | 1 | 0 |
| *Bulbophyllum ngoyense* Schltr. 2 | 1 | 1 | 0 | 1 | 1 | 1 | 1 | 1 | 1 | 1 | 1 | 1 | 1 | 1 | 1 | 1 | 1 | 1 | 1 | 1 | 1 | 1 | 1 | 1 | 1 | 1 | 1 | 1 | 1 | 1 | 1 | 1 | 1 | 1 | 1 | 1 | 1 | 1 | 0 | 1 | 1 | 1 | 0 | 1 | 1 | 1 | 1 | 0 | 1 | 1 | 1 | 1 | 1 | 1 | 1 | 1 | 1 | 1 | 1 | 1 | 1 | 1 | 1 |
| *Bulbophyllum nigricans* (Aver.) J.J.Verm., Schuit. & de Vogel | 0 | 1 | 1 | 1 | 1 | 1 | 1 | 1 | 1 | 1 | 1 | 1 | 1 | 1 | 1 | 1 | 1 | 1 | 1 | 0 | 1 | 1 | 1 | 0 | 0 | 0 | 1 | 1 | 1 | 1 | 1 | 1 | 1 | 1 | 1 | 1 | 1 | 1 | 1 | 1 | 1 | 1 | 1 | 1 | 0 | 1 | 1 | 0 | 0 | 1 | 1 | 1 | 0 | 1 | 1 | 0 | 1 | 1 | 1 | 0 | 0 | 1 | 0 |
| *Bulbophyllum ningboense* G.Y.Li ex H.L.Lin & X.P.Li | 1 | 1 | 1 | 1 | 1 | 1 | 1 | 1 | 1 | 1 | 1 | 1 | 1 | 1 | 1 | 1 | 1 | 1 | 1 | 1 | 1 | 1 | 1 | 1 | 1 | 1 | 1 | 1 | 1 | 1 | 1 | 1 | 1 | 1 | 1 | 1 | 1 | 1 | 1 | 1 | 1 | 1 | 1 | 1 | 1 | 1 | 1 | 1 | 1 | 1 | 1 | 1 | 1 | 1 | 1 | 1 | 0 | 1 | 1 | 1 | 1 | 1 | 1 |
| *Bulbophyllum nummularioides* Schltr. | 0 | 1 | 1 | 1 | 1 | 1 | 1 | 1 | 1 | 1 | 1 | 1 | 1 | 1 | 1 | 1 | 1 | 1 | 1 | 1 | 1 | 1 | 1 | 1 | 1 | 1 | 1 | 1 | 1 | 1 | 1 | 1 | 1 | 1 | 1 | 1 | 1 | 1 | 1 | 1 | 1 | 1 | 1 | 1 | 1 | 1 | 1 | 1 | 1 | 1 | 1 | 1 | 1 | 1 | 1 | 1 | 1 | 1 | 1 | 1 | 1 | 1 | 1 |
| *Bulbophyllum nymphopolitanum* Kraenzl. | 1 | 1 | 1 | 1 | 1 | 1 | 1 | 1 | 1 | 1 | 1 | 1 | 1 | 1 | 1 | 1 | 1 | 1 | 1 | 1 | 1 | 1 | 1 | 1 | 1 | 1 | 1 | 1 | 1 | 1 | 1 | 1 | 1 | 1 | 1 | 1 | 1 | 1 | 1 | 1 | 1 | 1 | 1 | 1 | 1 | 1 | 1 | 1 | 1 | 1 | 1 | 1 | 1 | 1 | 1 | 1 | 1 | 1 | 1 | 1 | 1 | 1 | 1 |
| *Bulbophyllum oblongum* (Lindl.) Rchb.f. | 1 | 1 | 1 | 1 | 1 | 1 | 1 | 1 | 1 | 1 | 1 | 1 | 1 | 1 | 1 | 1 | 1 | 1 | 1 | 1 | 1 | 1 | 1 | 1 | 0 | 1 | 1 | 1 | 1 | 1 | 1 | 1 | 1 | 1 | 1 | 1 | 1 | 1 | 1 | 1 | 1 | 1 | 1 | 1 | 1 | 1 | 1 | 1 | 1 | 1 | 1 | 1 | 0 | 1 | 1 | 1 | 1 | 1 | 1 | 1 | 1 | 1 | 1 |
| *Bulbophyllum obovatum* (J.J.Sm.) J.J.Verm., Schuit. & de Vogel | 1 | 1 | 1 | 1 | 1 | 1 | 1 | 1 | 1 | 1 | 1 | 1 | 1 | 1 | 1 | 1 | 1 | 1 | 1 | 1 | 1 | 1 | 1 | 1 | 1 | 1 | 1 | 1 | 1 | 1 | 1 | 1 | 1 | 1 | 1 | 1 | 1 | 1 | 1 | 1 | 0 | 1 | 1 | 1 | 1 | 1 | 1 | 1 | 1 | 1 | 1 | 1 | 1 | 1 | 1 | 1 | 1 | 1 | 1 | 1 | 1 | 1 | 1 |
| *Bulbophyllum occlusum* Ridl. | 1 | 1 | 1 | 1 | 1 | 1 | 1 | 1 | 1 | 1 | 1 | 0 | 1 | 0 | 1 | 1 | 1 | 1 | 1 | 1 | 1 | 1 | 1 | 1 | 1 | 1 | 1 | 1 | 1 | 1 | 1 | 1 | 1 | 1 | 1 | 1 | 0 | 1 | 1 | 1 | 1 | 1 | 1 | 1 | 1 | 1 | 1 | 1 | 1 | 0 | 1 | 1 | 1 | 1 | 1 | 1 | 0 | 1 | 0 | 1 | 1 | 1 | 1 |
| *Bulbophyllum ocellatum* Cootes & M.A.Clem. | 1 | 1 | 1 | 1 | 1 | 1 | 1 | 1 | 1 | 1 | 1 | 1 | 1 | 1 | 1 | 1 | 1 | 1 | 1 | 1 | 1 | 1 | 1 | 1 | 1 | 1 | 1 | 1 | 1 | 1 | 1 | 1 | 1 | 1 | 1 | 1 | 1 | 1 | 1 | 1 | 1 | 1 | 1 | 1 | 1 | 1 | 1 | 1 | 1 | 1 | 1 | 1 | 1 | 1 | 1 | 1 | 1 | 1 | 1 | 1 | 1 | 1 | 1 |
| *Bulbophyllum ochroleucum* Schltr. | 1 | 1 | 1 | 1 | 1 | 1 | 1 | 1 | 1 | 0 | 1 | 1 | 1 | 1 | 1 | 1 | 1 | 1 | 1 | 1 | 1 | 1 | 1 | 1 | 1 | 1 | 1 | 1 | 1 | 1 | 1 | 1 | 1 | 1 | 1 | 1 | 1 | 0 | 1 | 1 | 0 | 1 | 1 | 1 | 1 | 1 | 1 | 1 | 1 | 0 | 0 | 1 | 1 | 1 | 1 | 1 | 0 | 0 | 1 | 1 | 1 | 0 | 1 |
| *Bulbophyllum odoratissimum* Lindl. ex Wall. | 1 | 1 | 1 | 1 | 1 | 1 | 1 | 1 | 1 | 1 | 1 | 1 | 1 | 1 | 1 | 1 | 1 | 1 | 1 | 1 | 1 | 1 | 1 | 1 | 1 | 1 | 1 | 1 | 1 | 1 | 1 | 1 | 1 | 1 | 1 | 1 | 1 | 1 | 1 | 1 | 1 | 1 | 1 | 1 | 1 | 1 | 1 | 1 | 1 | 1 | 1 | 1 | 1 | 1 | 1 | 1 | 1 | 1 | 1 | 1 | 1 | 1 | 1 |
| *Bulbophyllum odoratum* (Blume) Lindl. | 1 | 1 | 1 | 1 | 1 | 1 | 1 | 1 | 1 | 1 | 1 | 1 | 1 | 1 | 1 | 1 | 1 | 1 | 1 | 1 | 1 | 1 | 1 | 1 | 1 | 1 | 1 | 1 | 1 | 1 | 1 | 1 | 1 | 1 | 1 | 1 | 1 | 1 | 1 | 1 | 1 | 1 | 1 | 1 | 1 | 1 | 1 | 1 | 1 | 1 | 1 | 1 | 1 | 1 | 1 | 1 | 1 | 1 | 1 | 1 | 1 | 1 | 1 |
| *Bulbophyllum olivinum* J.J.Sm. | 1 | 1 | 1 | 1 | 1 | 1 | 1 | 1 | 1 | 1 | 1 | 1 | 1 | 1 | 1 | 1 | 1 | 1 | 1 | 1 | 1 | 1 | 1 | 1 | 1 | 1 | 1 | 1 | 1 | 1 | 1 | 1 | 1 | 1 | 1 | 1 | 1 | 1 | 1 | 1 | 1 | 1 | 1 | 1 | 1 | 1 | 1 | 1 | 1 | 1 | 1 | 1 | 1 | 1 | 1 | 1 | 1 | 1 | 1 | 1 | 1 | 1 | 1 |
| *Bulbophyllum orectopetalum* Garay, Hamer & Siegerist | 1 | 1 | 1 | 1 | 1 | 1 | 1 | 1 | 1 | 1 | 1 | 1 | 1 | 1 | 1 | 1 | 1 | 1 | 1 | 1 | 1 | 1 | 1 | 1 | 1 | 1 | 1 | 1 | 1 | 1 | 1 | 1 | 1 | 1 | 1 | 1 | 1 | 1 | 1 | 1 | 1 | 1 | 1 | 1 | 1 | 1 | 1 | 1 | 1 | 1 | 1 | 1 | 1 | 1 | 1 | 1 | 1 | 1 | 1 | 1 | 1 | 1 | 1 |
| *Bulbophyllum oreomene* J.J.Verm., Schuit. & de Vogel | 1 | 1 | 0 | 1 | 1 | 1 | 1 | 0 | 1 | 1 | 1 | 1 | 1 | 1 | 1 | 1 | 1 | 1 | 1 | 1 | 1 | 1 | 1 | 1 | 1 | 1 | 1 | 1 | 1 | 1 | 1 | 1 | 1 | 1 | 1 | 1 | 1 | 1 | 1 | 1 | 0 | 1 | 0 | 1 | 1 | 1 | 1 | 1 | 1 | 1 | 1 | 1 | 1 | 1 | 1 | 1 | 1 | 1 | 1 | 1 | 1 | 1 | 1 |
| *Bulbophyllum orientale* Seidenf. | 1 | 1 | 1 | 1 | 1 | 1 | 1 | 1 | 1 | 1 | 1 | 1 | 1 | 1 | 1 | 1 | 1 | 1 | 1 | 1 | 1 | 1 | 1 | 1 | 1 | 1 | 1 | 1 | 1 | 1 | 1 | 1 | 1 | 1 | 1 | 1 | 1 | 1 | 1 | 1 | 1 | 1 | 1 | 1 | 1 | 1 | 1 | 1 | 1 | 1 | 1 | 1 | 1 | 1 | 1 | 1 | 1 | 1 | 1 | 1 | 1 | 1 | 1 |
| *Bulbophyllum orthosepalum* J.J.Verm. | 1 | 1 | 1 | 1 | 1 | 1 | 1 | 1 | 1 | 1 | 1 | 1 | 1 | 1 | 1 | 1 | 1 | 1 | 1 | 1 | 1 | 1 | 1 | 1 | 1 | 1 | 1 | 1 | 1 | 1 | 1 | 1 | 1 | 1 | 1 | 1 | 1 | 1 | 1 | 1 | 1 | 1 | 1 | 1 | 1 | 1 | 1 | 1 | 1 | 1 | 1 | 1 | 1 | 1 | 1 | 1 | 1 | 1 | 1 | 1 | 1 | 1 | 1 |

| | | | | | | | | | | | | | | | | | | | | | | | | | | | | | | | | | | | | | | | | | | | | | | | | | | | | | | | | | | | | | | | |
|---|---|---|---|---|---|---|---|---|---|---|---|---|---|---|---|---|---|---|---|---|---|---|---|---|---|---|---|---|---|---|---|---|---|---|---|---|---|---|---|---|---|---|---|---|---|---|---|---|---|---|---|---|---|---|---|---|---|---|---|---|---|---|---|
| *Bulbophyllum pachypus* Schltr. | 1 | 1 | 1 | 1 | 1 | 1 | 1 | 1 | 1 | 1 | 1 | 1 | 1 | 1 | 1 | 1 | 1 | 1 | 1 | 1 | 1 | 1 | 1 | 1 | 1 | 1 | 1 | 1 | 1 | 1 | 1 | 1 | 1 | 1 | 1 | 1 | 1 | 1 | 1 | 1 | 1 | 1 | 0 | 1 | 1 | 1 | 1 | 1 | 1 | 1 | 1 | 1 | 1 | 1 | 1 | 1 | 1 | 1 | 1 | 1 | 1 | 1 | 1 |
| *Bulbophyllum pahudii* (de Vriese) Rchb.f. 1 | 1 | 1 | 1 | 1 | 1 | 1 | 1 | 1 | 1 | 1 | 1 | 1 | 1 | 1 | 1 | 1 | 1 | 1 | 1 | 1 | 1 | 1 | 1 | 1 | 1 | 1 | 1 | 1 | 1 | 1 | 1 | 1 | 1 | 1 | 1 | 1 | 1 | 1 | 1 | 1 | 1 | 1 | 1 | 1 | 1 | 1 | 1 | 1 | 1 | 0 | 1 | 1 | 1 | 1 | 1 | 1 | 0 | 1 | 1 | 1 | 1 | 1 | 1 |
| *Bulbophyllum pahudii* (de Vriese) Rchb.f. 2 | 1 | 1 | 1 | 1 | 1 | 1 | 1 | 1 | 1 | 1 | 1 | 1 | 1 | 1 | 1 | 1 | 1 | 1 | 1 | 1 | 1 | 1 | 1 | 1 | 1 | 1 | 1 | 1 | 1 | 1 | 1 | 1 | 1 | 1 | 1 | 1 | 1 | 1 | 1 | 1 | 1 | 1 | 1 | 1 | 1 | 1 | 1 | 1 | 1 | 1 | 1 | 1 | 1 | 1 | 1 | 1 | 0 | 1 | 1 | 1 | 1 | 1 | 1 |
| *Bulbophyllum pardalotum* Garay, Hamer & Siegerist | 1 | 1 | 1 | 1 | 1 | 1 | 1 | 1 | 1 | 1 | 1 | 1 | 1 | 1 | 1 | 1 | 1 | 1 | 0 | 1 | 1 | 1 | 1 | 1 | 1 | 1 | 1 | 1 | 1 | 1 | 1 | 1 | 1 | 1 | 1 | 1 | 1 | 1 | 1 | 1 | 1 | 1 | 1 | 1 | 1 | 1 | 1 | 1 | 1 | 1 | 1 | 1 | 1 | 1 | 1 | 1 | 1 | 1 | 1 | 1 | 1 | 1 | 1 |
| *Bulbophyllum parviflorum* C.S.P.Parish & Rchb.f. | 1 | 1 | 1 | 1 | 1 | 1 | 1 | 0 | 1 | 1 | 1 | 1 | 1 | 1 | 1 | 1 | 1 | 1 | 1 | 1 | 1 | 1 | 1 | 1 | 0 | 1 | 1 | 1 | 1 | 1 | 1 | 1 | 1 | 1 | 1 | 1 | 1 | 1 | 1 | 1 | 1 | 0 | 1 | 1 | 1 | 1 | 1 | 1 | 1 | 1 | 1 | 0 | 1 | 1 | 1 | 1 | 1 | 1 | 1 | 1 | 1 | 1 | 0 |
| *Bulbophyllum patens* King ex Hook.f. | 1 | 1 | 1 | 1 | 1 | 1 | 1 | 1 | 1 | 1 | 1 | 1 | 1 | 1 | 1 | 1 | 1 | 1 | 1 | 1 | 1 | 1 | 1 | 1 | 1 | 1 | 1 | 1 | 1 | 1 | 1 | 1 | 1 | 1 | 1 | 1 | 1 | 1 | 1 | 1 | 1 | 1 | 1 | 1 | 1 | 1 | 1 | 1 | 1 | 1 | 1 | 1 | 1 | 1 | 1 | 1 | 1 | 1 | 1 | 1 | 1 | 1 | 1 |
| *Bulbophyllum pecten-veneris* (Gagnep.) Seidenf. | 1 | 1 | 1 | 1 | 1 | 1 | 1 | 1 | 1 | 1 | 1 | 1 | 1 | 1 | 1 | 1 | 1 | 1 | 1 | 1 | 1 | 1 | 1 | 1 | 1 | 1 | 1 | 1 | 1 | 1 | 1 | 1 | 1 | 1 | 1 | 1 | 1 | 1 | 1 | 1 | 1 | 1 | 1 | 1 | 1 | 1 | 1 | 1 | 1 | 1 | 1 | 1 | 1 | 1 | 1 | 1 | 1 | 1 | 1 | 1 | 1 | 1 | 1 |
| *Bulbophyllum penicillium* C.S.P.Parish & Rchb.f. | 1 | 1 | 1 | 1 | 1 | 1 | 1 | 1 | 1 | 1 | 1 | 1 | 1 | 1 | 1 | 1 | 1 | 1 | 1 | 1 | 1 | 1 | 1 | 1 | 0 | 1 | 1 | 1 | 1 | 1 | 1 | 1 | 1 | 1 | 1 | 1 | 1 | 1 | 1 | 1 | 1 | 0 | 1 | 1 | 1 | 1 | 1 | 1 | 1 | 1 | 1 | 1 | 0 | 1 | 1 | 1 | 1 | 1 | 1 | 1 | 1 | 1 | 1 |
| *Bulbophyllum peninsulare* Seidenf. | 1 | 1 | 1 | 1 | 1 | 1 | 1 | 1 | 1 | 1 | 1 | 1 | 1 | 1 | 1 | 1 | 1 | 1 | 1 | 1 | 1 | 1 | 1 | 1 | 1 | 1 | 1 | 1 | 1 | 1 | 1 | 1 | 1 | 1 | 1 | 1 | 1 | 1 | 1 | 1 | 1 | 1 | 1 | 1 | 1 | 1 | 1 | 1 | 1 | 1 | 1 | 1 | 1 | 1 | 1 | 1 | 1 | 1 | 1 | 1 | 1 | 1 | 1 |
| *Bulbophyllum petrophilum* (P.Royen) J.J.Verm., Schuit. & de Vogel | 1 | 1 | 1 | 1 | 1 | 1 | 1 | 0 | 1 | 1 | 1 | 1 | 1 | 1 | 1 | 1 | 1 | 1 | 1 | 1 | 1 | 1 | 1 | 1 | 1 | 1 | 1 | 1 | 1 | 1 | 1 | 1 | 1 | 1 | 1 | 1 | 1 | 1 | 1 | 1 | 1 | 1 | 1 | 1 | 1 | 1 | 1 | 1 | 1 | 1 | 1 | 1 | 1 | 1 | 1 | 1 | 0 | 1 | 1 | 1 | 1 | 0 | 1 |
| *Bulbophyllum phaeoglossum* Schltr. | 1 | 1 | 1 | 1 | 1 | 0 | 1 | 1 | 1 | 1 | 1 | 1 | 1 | 1 | 1 | 1 | 1 | 1 | 1 | 1 | 1 | 1 | 1 | 1 | 1 | 1 | 1 | 1 | 1 | 1 | 1 | 1 | 1 | 1 | 1 | 0 | 0 | 1 | 1 | 1 | 0 | 1 | 1 | 1 | 1 | 1 | 1 | 1 | 1 | 0 | 1 | 1 | 1 | 1 | 1 | 1 | 0 | 1 | 1 | 1 | 1 | 1 | 1 |
| *Bulbophyllum phalaenopsis* J.J.Sm. | 1 | 1 | 1 | 1 | 1 | 1 | 1 | 1 | 1 | 1 | 1 | 1 | 1 | 1 | 1 | 1 | 1 | 1 | 1 | 1 | 1 | 1 | 1 | 1 | 1 | 1 | 1 | 1 | 1 | 1 | 1 | 1 | 1 | 1 | 1 | 1 | 1 | 1 | 1 | 1 | 1 | 1 | 1 | 1 | 1 | 1 | 1 | 1 | 1 | 1 | 1 | 1 | 1 | 1 | 1 | 1 | 1 | 1 | 1 | 1 | 1 | 1 | 1 |
| *Bulbophyllum phormion* J.J.Verm. | 1 | 1 | 1 | 1 | 1 | 1 | 1 | 1 | 0 | 1 | 1 | 1 | 0 | 1 | 1 | 1 | 1 | 1 | 1 | 1 | 1 | 1 | 1 | 1 | 0 | 0 | 1 | 1 | 1 | 1 | 1 | 1 | 1 | 1 | 1 | 1 | 1 | 1 | 0 | 1 | 1 | 1 | 1 | 1 | 1 | 1 | 1 | 1 | 0 | 1 | 0 | 0 | 1 | 1 | 1 | 1 | 1 | 1 | 1 | 0 | 1 | 1 | 0 |
| *Bulbophyllum pictum* C.S.P.Parish & Rchb.f. | 1 | 1 | 1 | 1 | 1 | 1 | 1 | 1 | 1 | 1 | 1 | 1 | 1 | 1 | 1 | 1 | 1 | 1 | 1 | 1 | 1 | 1 | 1 | 1 | 1 | 1 | 1 | 1 | 1 | 1 | 1 | 1 | 1 | 1 | 1 | 1 | 1 | 1 | 1 | 1 | 1 | 1 | 1 | 1 | 1 | 1 | 1 | 1 | 1 | 1 | 1 | 1 | 1 | 1 | 1 | 1 | 1 | 1 | 1 | 1 | 1 | 1 | 1 |
| *Bulbophyllum picturatum* Rchb.f. | 1 | 1 | 1 | 1 | 1 | 1 | 1 | 1 | 1 | 1 | 1 | 1 | 0 | 1 | 1 | 1 | 1 | 1 | 1 | 1 | 1 | 1 | 1 | 1 | 1 | 1 | 1 | 1 | 1 | 1 | 1 | 1 | 1 | 1 | 1 | 1 | 1 | 1 | 1 | 1 | 1 | 1 | 1 | 1 | 1 | 1 | 1 | 1 | 1 | 1 | 1 | 1 | 1 | 1 | 1 | 1 | 1 | 1 | 1 | 1 | 1 | 1 | 1 |
| *Bulbophyllum piestobulbon* Schltr. | 0 | 1 | 1 | 1 | 1 | 1 | 1 | 0 | 0 | 0 | 0 | 1 | 0 | 0 | 0 | 0 | 0 | 1 | 1 | 1 | 0 | 0 | 0 | 1 | 0 | 1 | 1 | 0 | 0 | 0 | 1 | 0 | 1 | 0 | 1 | 0 | 0 | 1 | 1 | 0 | 0 | 0 | 0 | 0 | 0 | 0 | 0 | 0 | 1 | 1 | 1 | 1 | 0 | 0 | 0 | 0 | 0 | 1 | 0 | 1 | 0 | 1 | 0 |
| *Bulbophyllum piundaundense* (P.Royen) J.J.Verm., Schuit. & de Vogel | 1 | 1 | 1 | 1 | 1 | 1 | 1 | 1 | 1 | 1 | 1 | 1 | 1 | 1 | 1 | 1 | 1 | 1 | 1 | 1 | 1 | 1 | 1 | 1 | 1 | 1 | 1 | 1 | 1 | 1 | 1 | 1 | 1 | 1 | 1 | 1 | 1 | 0 | 1 | 1 | 1 | 1 | 1 | 1 | 1 | 1 | 1 | 1 | 1 | 0 | 1 | 1 | 1 | 1 | 1 | 1 | 1 | 1 | 1 | 1 | 1 | 1 | 1 |
| *Bulbophyllum planibulbe* Ridl. | 1 | 1 | 0 | 1 | 1 | 1 | 1 | 1 | 0 | 1 | 1 | 0 | 0 | 1 | 1 | 1 | 1 | 1 | 1 | 1 | 1 | 1 | 1 | 0 | 1 | 1 | 0 | 1 | 1 | 1 | 1 | 0 | 1 | 0 | 1 | 1 | 1 | 1 | 0 | 1 | 1 | 1 | 1 | 1 | 1 | 1 | 1 | 1 | 1 | 1 | 0 | 1 | 1 | 1 | 1 | 1 | 1 | 0 | 1 | 1 | 1 | 1 | 1 |
| *Bulbophyllum pleurothallidanthum* Garay 1 | 1 | 1 | 1 | 0 | 1 | 1 | 1 | 1 | 1 | 1 | 1 | 1 | 1 | 1 | 1 | 1 | 1 | 1 | 1 | 1 | 1 | 1 | 1 | 1 | 1 | 1 | 1 | 1 | 1 | 1 | 1 | 1 | 1 | 1 | 1 | 1 | 1 | 1 | 1 | 1 | 1 | 1 | 0 | 1 | 1 | 1 | 1 | 1 | 1 | 1 | 1 | 1 | 1 | 1 | 1 | 1 | 1 | 1 | 1 | 1 | 1 | 1 | 1 |
| *Bulbophyllum pleurothallidanthum* Garay 2 | 1 | 1 | 1 | 1 | 1 | 1 | 1 | 1 | 1 | 1 | 1 | 1 | 1 | 1 | 1 | 1 | 1 | 1 | 1 | 1 | 1 | 1 | 1 | 1 | 1 | 1 | 1 | 1 | 1 | 1 | 1 | 1 | 1 | 1 | 1 | 1 | 1 | 1 | 1 | 1 | 1 | 1 | 1 | 1 | 1 | 1 | 1 | 1 | 1 | 1 | 1 | 1 | 1 | 1 | 1 | 1 | 1 | 1 | 1 | 1 | 1 | 1 | 1 |
| *Bulbophyllum plumosum* (Barb.Rodr.) Cogn. | 1 | 1 | 1 | 1 | 1 | 1 | 1 | 1 | 1 | 1 | 1 | 1 | 1 | 1 | 1 | 1 | 1 | 1 | 1 | 1 | 1 | 1 | 1 | 1 | 1 | 1 | 1 | 1 | 1 | 1 | 1 | 1 | 1 | 1 | 1 | 1 | 1 | 1 | 1 | 1 | 1 | 1 | 1 | 1 | 1 | 1 | 1 | 1 | 1 | 1 | 1 | 1 | 1 | 1 | 1 | 1 | 0 | 1 | 1 | 1 | 1 | 1 | 1 |
| *Bulbophyllum plumula* Schltr. | 1 | 1 | 0 | 1 | 1 | 1 | 1 | 1 | 1 | 1 | 1 | 1 | 1 | 0 | 1 | 1 | 1 | 1 | 1 | 1 | 1 | 1 | 1 | 1 | 1 | 1 | 1 | 1 | 1 | 1 | 1 | 1 | 1 | 1 | 1 | 1 | 1 | 1 | 0 | 1 | 0 | 1 | 1 | 1 | 1 | 1 | 1 | 1 | 1 | 0 | 1 | 1 | 1 | 1 | 1 | 1 | 1 | 1 | 1 | 1 | 1 | 0 | 0 |
| *Bulbophyllum polliculosum* Seidenf. | 1 | 1 | 1 | 1 | 1 | 1 | 1 | 1 | 1 | 1 | 1 | 1 | 1 | 1 | 1 | 1 | 1 | 1 | 1 | 1 | 1 | 1 | 1 | 1 | 1 | 1 | 1 | 1 | 1 | 1 | 1 | 1 | 1 | 1 | 1 | 1 | 1 | 1 | 1 | 1 | 1 | 1 | 1 | 1 | 1 | 1 | 1 | 1 | 1 | 1 | 1 | 1 | 0 | 1 | 1 | 1 | 1 | 1 | 1 | 1 | 1 | 1 | 1 |
| *Bulbophyllum polyblepharon* Schltr. | 1 | 1 | 1 | 1 | 1 | 1 | 1 | 1 | 1 | 1 | 1 | 1 | 1 | 1 | 1 | 1 | 1 | 1 | 1 | 1 | 1 | 1 | 1 | 1 | 1 | 1 | 1 | 1 | 1 | 1 | 1 | 1 | 0 | 1 | 1 | 1 | 1 | 1 | 1 | 1 | 1 | 1 | 1 | 1 | 1 | 1 | 1 | 1 | 1 | 0 | 1 | 1 | 1 | 1 | 1 | 1 | 0 | 1 | 1 | 1 | 1 | 1 | 1 |
| *Bulbophyllum prianganense* J.J.Sm. | 1 | 1 | 1 | 1 | 1 | 1 | 1 | 1 | 1 | 1 | 1 | 1 | 1 | 1 | 1 | 1 | 1 | 1 | 1 | 1 | 1 | 1 | 1 | 1 | 1 | 1 | 1 | 1 | 1 | 1 | 1 | 1 | 1 | 1 | 1 | 1 | 1 | 1 | 1 | 1 | 1 | 1 | 1 | 1 | 1 | 1 | 1 | 1 | 1 | 1 | 1 | 1 | 1 | 1 | 1 | 1 | 1 | 1 | 1 | 1 | 1 | 1 | 1 |
| *Bulbophyllum propinquum* Kraenzl. | 1 | 1 | 1 | 0 | 1 | 1 | 1 | 1 | 1 | 1 | 1 | 1 | 1 | 1 | 1 | 1 | 1 | 1 | 1 | 1 | 1 | 1 | 1 | 1 | 1 | 1 | 1 | 1 | 1 | 1 | 1 | 1 | 1 | 1 | 1 | 1 | 1 | 1 | 1 | 1 | 1 | 1 | 1 | 1 | 1 | 1 | 1 | 1 | 1 | 0 | 1 | 1 | 1 | 1 | 1 | 1 | 0 | 1 | 1 | 1 | 1 | 1 | 1 |
| *Bulbophyllum ptilotes* Schltr. | 1 | 1 | 1 | 1 | 1 | 1 | 1 | 1 | 1 | 1 | 1 | 1 | 1 | 1 | 1 | 1 | 1 | 1 | 1 | 1 | 1 | 1 | 1 | 1 | 1 | 1 | 1 | 1 | 1 | 1 | 1 | 1 | 1 | 1 | 1 | 1 | 1 | 1 | 1 | 1 | 1 | 1 | 1 | 1 | 1 | 1 | 1 | 1 | 1 | 0 | 1 | 1 | 1 | 1 | 1 | 1 | 0 | 1 | 1 | 1 | 1 | 1 | 1 |
| *Bulbophyllum pulvinatum* Schltr. | 1 | 1 | 1 | 1 | 1 | 1 | 1 | 1 | 1 | 1 | 1 | 1 | 1 | 1 | 1 | 1 | 1 | 1 | 1 | 1 | 1 | 1 | 1 | 1 | 1 | 1 | 1 | 1 | 1 | 1 | 1 | 1 | 1 | 1 | 1 | 1 | 1 | 1 | 1 | 1 | 1 | 1 | 1 | 1 | 1 | 1 | 1 | 1 | 1 | 1 | 1 | 1 | 1 | 1 | 1 | 1 | 1 | 1 | 1 | 1 | 1 | 1 | 1 |
| *Bulbophyllum punamense* Schltr. 1 | 0 | 0 | 1 | 1 | 0 | 0 | 0 | 0 | 1 | 1 | 0 | 0 | 1 | 0 | 0 | 1 | 1 | 1 | 1 | 1 | 0 | 1 | 1 | 0 | 0 | 1 | 1 | 1 | 1 | 0 | 0 | 1 | 0 | 1 | 1 | 0 | 0 | 1 | 1 | 0 | 0 | 1 | 0 | 0 | 0 | 1 | 0 | 0 | 1 | 1 | 0 | 0 | 0 | 0 | 0 | 0 | 0 | 1 | 0 | 0 | 1 | 1 | 1 |
| *Bulbophyllum punamense* Schltr. 2 | 1 | 1 | 1 | 1 | 1 | 1 | 1 | 1 | 1 | 1 | 1 | 1 | 1 | 1 | 1 | 1 | 1 | 1 | 1 | 1 | 1 | 1 | 1 | 1 | 1 | 1 | 1 | 1 | 1 | 1 | 1 | 1 | 1 | 1 | 1 | 1 | 1 | 1 | 1 | 1 | 1 | 1 | 0 | 1 | 1 | 1 | 1 | 1 | 1 | 1 | 1 | 1 | 1 | 1 | 1 | 1 | 1 | 1 | 1 | 0 | 1 | 1 | 1 |
| *Bulbophyllum punamense* Schltr. 3 | 1 | 1 | 1 | 1 | 1 | 1 | 1 | 1 | 1 | 1 | 1 | 1 | 1 | 1 | 1 | 1 | 1 | 1 | 1 | 1 | 1 | 1 | 1 | 1 | 1 | 1 | 1 | 1 | 1 | 1 | 1 | 1 | 1 | 1 | 1 | 1 | 1 | 1 | 1 | 1 | 1 | 1 | 0 | 1 | 0 | 1 | 1 | 1 | 1 | 1 | 1 | 1 | 1 | 1 | 1 | 1 | 1 | 1 | 1 | 0 | 1 | 1 | 1 |
| *Bulbophyllum pungens* Schltr. | 0 | 1 | 1 | 1 | 1 | 1 | 1 | 1 | 1 | 1 | 1 | 1 | 1 | 1 | 1 | 1 | 1 | 1 | 1 | 1 | 1 | 1 | 1 | 1 | 1 | 1 | 1 | 1 | 1 | 1 | 1 | 1 | 1 | 1 | 1 | 1 | 1 | 1 | 1 | 1 | 1 | 1 | 1 | 1 | 1 | 1 | 1 | 1 | 1 | 1 | 1 | 0 | 1 | 1 | 1 | 1 | 1 | 1 | 1 | 1 | 1 | 1 | 1 |
| *Bulbophyllum putidum* (Teijsm. & Binn.) J.J.Sm. | 1 | 1 | 1 | 1 | 1 | 1 | 1 | 1 | 0 | 1 | 0 | 0 | 0 | 1 | 0 | 1 | 1 | 1 | 1 | 0 | 1 | 1 | 1 | 0 | 1 | 1 | 1 | 1 | 1 | 1 | 1 | 1 | 1 | 1 | 1 | 1 | 1 | 1 | 1 | 0 | 0 | 1 | 0 | 0 | 1 | 1 | 0 | 0 | 1 | 1 | 1 | 1 | 0 | 0 | 0 | 0 | 0 | 0 | 1 | 1 | 1 | 1 | 0 |
| *Bulbophyllum pygmaeum* Lindl. | 1 | 1 | 1 | 1 | 1 | 1 | 1 | 1 | 1 | 1 | 1 | 1 | 1 | 1 | 1 | 1 | 1 | 1 | 1 | 0 | 1 | 1 | 1 | 1 | 1 | 1 | 1 | 1 | 1 | 1 | 1 | 1 | 1 | 1 | 1 | 1 | 1 | 1 | 1 | 1 | 1 | 1 | 1 | 1 | 1 | 0 | 1 | 0 | 0 | 1 | 0 | 1 | 0 | 1 | 1 | 1 | 1 | 0 | 1 | 1 | 1 | 1 | 0 |
| *Bulbophyllum quadrangulare* J.J.Sm. | 1 | 1 | 1 | 1 | 1 | 1 | 1 | 1 | 1 | 1 | 1 | 1 | 1 | 1 | 1 | 1 | 1 | 1 | 1 | 1 | 1 | 1 | 1 | 1 | 1 | 1 | 1 | 1 | 1 | 1 | 1 | 1 | 1 | 1 | 1 | 1 | 1 | 1 | 1 | 1 | 1 | 1 | 1 | 1 | 1 | 1 | 1 | 0 | 1 | 1 | 0 | 1 | 1 | 1 | 1 | 1 | 1 | 1 | 1 | 1 | 1 | 1 | 1 |
| *Bulbophyllum quadrifalciculatum* J.J.Sm. | 1 | 1 | 1 | 1 | 1 | 1 | 1 | 0 | 0 | 1 | 1 | 0 | 1 | 1 | 1 | 1 | 1 | 1 | 0 | 1 | 1 | 1 | 1 | 1 | 1 | 1 | 1 | 1 | 1 | 1 | 1 | 1 | 1 | 1 | 1 | 1 | 1 | 1 | 1 | 1 | 1 | 1 | 1 | 1 | 1 | 1 | 1 | 1 | 1 | 1 | 0 | 1 | 1 | 1 | 1 | 1 | 1 | 1 | 0 | 1 | 1 | 1 | 0 |
| *Bulbophyllum radicans* F.M.Bailey | 1 | 1 | 1 | 1 | 1 | 1 | 1 | 1 | 1 | 1 | 1 | 0 | 1 | 1 | 0 | 1 | 1 | 1 | 1 | 1 | 1 | 1 | 1 | 1 | 1 | 1 | 1 | 1 | 1 | 1 | 1 | 1 | 1 | 1 | 1 | 1 | 1 | 1 | 1 | 1 | 1 | 1 | 1 | 1 | 1 | 1 | 1 | 1 | 1 | 1 | 1 | 0 | 1 | 1 | 1 | 1 | 1 | 0 | 1 | 1 | 1 | 1 | 1 |
| *Bulbophyllum refractilingue* J.J.Sm. | 1 | 1 | 0 | 1 | 1 | 1 | 1 | 1 | 1 | 1 | 1 | 1 | 1 | 1 | 1 | 1 | 1 | 1 | 1 | 1 | 1 | 1 | 1 | 1 | 1 | 1 | 1 | 1 | 1 | 1 | 1 | 1 | 1 | 1 | 1 | 1 | 1 | 1 | 1 | 1 | 1 | 1 | 1 | 1 | 1 | 1 | 1 | 1 | 1 | 1 | 1 | 1 | 0 | 1 | 1 | 1 | 1 | 1 | 1 | 1 | 1 | 1 | 1 |
| *Bulbophyllum regnellii* Rchb.f. | 1 | 1 | 1 | 1 | 1 | 1 | 1 | 1 | 1 | 1 | 1 | 1 | 1 | 1 | 1 | 1 | 1 | 1 | 1 | 1 | 1 | 1 | 1 | 1 | 1 | 1 | 1 | 1 | 1 | 1 | 1 | 1 | 1 | 1 | 1 | 1 | 1 | 1 | 1 | 1 | 1 | 1 | 1 | 1 | 1 | 1 | 1 | 1 | 1 | 1 | 1 | 1 | 1 | 1 | 1 | 1 | 0 | 1 | 1 | 1 | 1 | 1 | 1 |
| *Bulbophyllum renipetalum* Schltr. | 1 | 1 | 1 | 1 | 1 | 1 | 1 | 0 | 1 | 1 | 1 | 1 | 1 | 1 | 1 | 1 | 1 | 1 | 1 | 1 | 1 | 1 | 1 | 1 | 1 | 1 | 1 | 1 | 1 | 0 | 1 | 1 | 1 | 1 | 1 | 1 | 1 | 1 | 1 | 1 | 1 | 1 | 1 | 1 | 1 | 1 | 1 | 1 | 1 | 1 | 0 | 1 | 1 | 1 | 1 | 1 | 1 | 1 | 1 | 0 | 1 | 1 | 1 |
| *Bulbophyllum repens* Griff. | 1 | 0 | 1 | 1 | 1 | 1 | 1 | 1 | 1 | 1 | 1 | 1 | 1 | 1 | 1 | 1 | 1 | 1 | 1 | 1 | 1 | 1 | 1 | 1 | 1 | 1 | 1 | 1 | 1 | 1 | 1 | 1 | 1 | 1 | 1 | 1 | 1 | 1 | 1 | 1 | 1 | 1 | 1 | 1 | 1 | 1 | 1 | 1 | 1 | 0 | 1 | 1 | 1 | 1 | 1 | 1 | 1 | 1 | 1 | 1 | 1 | 1 | 1 |
| *Bulbophyllum reptans* Lindl. ex Wall. | 1 | 1 | 1 | 1 | 1 | 1 | 1 | 1 | 1 | 1 | 1 | 1 | 1 | 1 | 1 | 1 | 1 | 1 | 1 | 1 | 1 | 1 | 1 | 1 | 1 | 1 | 0 | 1 | 1 | 1 | 1 | 1 | 1 | 1 | 1 | 1 | 0 | 1 | 1 | 1 | 1 | 0 | 1 | 1 | 1 | 1 | 1 | 1 | 1 | 0 | 1 | 1 | 1 | 1 | 1 | 1 | 1 | 1 | 1 | 1 | 1 | 1 | 1 |
| *Bulbophyllum restrepia* (Ridl.) Ridl. | 1 | 1 | 1 | 1 | 1 | 1 | 1 | 1 | 1 | 1 | 1 | 0 | 0 | 0 | 0 | 0 | 0 | 1 | 1 | 0 | 1 | 1 | 0 | 0 | 1 | 1 | 0 | 0 | 0 | 0 | 0 | 0 | 0 | 0 | 0 | 0 | 0 | 0 | 1 | 1 | 1 | 0 | 1 | 1 | 1 | 0 | 1 | 1 | 1 | 1 | 1 | 0 | 1 | 1 | 1 | 1 | 1 | 0 | 1 | 0 | 0 | 1 | 1 |
| *Bulbophyllum reticulatum* Bateman ex Hook.f. | 1 | 1 | 1 | 1 | 1 | 1 | 1 | 1 | 1 | 1 | 1 | 1 | 1 | 1 | 1 | 1 | 1 | 1 | 1 | 1 | 1 | 1 | 1 | 1 | 1 | 1 | 1 | 1 | 1 | 1 | 1 | 1 | 1 | 1 | 1 | 1 | 1 | 1 | 1 | 1 | 1 | 1 | 1 | 1 | 1 | 1 | 1 | 1 | 1 | 1 | 1 | 1 | 1 | 1 | 1 | 1 | 1 | 1 | 1 | 1 | 1 | 1 | 1 |
| *Bulbophyllum retusiusculum* Rchb.f. | 1 | 1 | 1 | 1 | 1 | 1 | 1 | 1 | 1 | 1 | 1 | 1 | 1 | 1 | 1 | 1 | 1 | 1 | 1 | 1 | 1 | 1 | 1 | 1 | 1 | 1 | 1 | 1 | 1 | 1 | 1 | 1 | 1 | 1 | 1 | 1 | 1 | 1 | 1 | 1 | 1 | 1 | 1 | 1 | 1 | 1 | 1 | 1 | 1 | 1 | 1 | 1 | 1 | 1 | 1 | 1 | 1 | 1 | 1 | 1 | 1 | 1 | 1 |
| *Bulbophyllum rhodoglossum* Schltr. | 1 | 1 | 1 | 1 | 1 | 1 | 1 | 1 | 1 | 1 | 1 | 1 | 1 | 1 | 1 | 1 | 1 | 1 | 1 | 1 | 1 | 1 | 1 | 1 | 1 | 1 | 1 | 1 | 1 | 1 | 1 | 1 | 1 | 1 | 1 | 1 | 1 | 1 | 1 | 1 | 1 | 1 | 1 | 1 | 1 | 1 | 1 | 1 | 1 | 1 | 1 | 1 | 1 | 1 | 1 | 1 | 1 | 0 | 1 | 1 | 1 | 1 | 1 |
| *Bulbophyllum rivulare* Schltr. | 0 | 1 | 1 | 1 | 1 | 1 | 1 | 1 | 1 | 1 | 1 | 1 | 1 | 1 | 1 | 1 | 1 | 1 | 1 | 1 | 1 | 1 | 1 | 1 | 1 | 1 | 1 | 1 | 1 | 1 | 1 | 1 | 1 | 1 | 1 | 1 | 1 | 1 | 1 | 1 | 1 | 1 | 1 | 1 | 1 | 1 | 1 | 1 | 1 | 1 | 1 | 1 | 1 | 1 | 1 | 1 | 1 | 1 | 1 | 1 | 1 | 1 | 1 |
| *Bulbophyllum romyi* B.Thoms | 1 | 1 | 1 | 1 | 1 | 1 | 1 | 1 | 1 | 1 | 1 | 1 | 1 | 1 | 1 | 1 | 1 | 1 | 1 | 1 | 1 | 1 | 1 | 1 | 1 | 1 | 1 | 1 | 1 | 1 | 1 | 1 | 1 | 1 | 1 | 1 | 1 | 1 | 1 | 1 | 1 | 1 | 1 | 1 | 1 | 1 | 1 | 1 | 1 | 1 | 1 | 1 | 1 | 1 | 1 | 1 | 1 | 1 | 1 | 1 | 1 | 1 | 1 |
| *Bulbophyllum roseopictum* J.J.Verm., Schuit. & de Vogel | 1 | 1 | 1 | 1 | 1 | 1 | 1 | 1 | 1 | 1 | 1 | 0 | 1 | 1 | 1 | 1 | 1 | 1 | 1 | 1 | 1 | 1 | 1 | 1 | 1 | 1 | 1 | 1 | 1 | 1 | 1 | 1 | 1 | 1 | 1 | 1 | 1 | 1 | 1 | 1 | 1 | 1 | 0 | 1 | 1 | 1 | 1 | 1 | 1 | 1 | 1 | 1 | 1 | 1 | 1 | 1 | 1 | 1 | 1 | 0 | 1 | 1 | 1 |
| *Bulbophyllum rothschildianum* (O'Brien) J.J.Sm. | 1 | 1 | 1 | 1 | 1 | 1 | 1 | 1 | 1 | 1 | 1 | 1 | 1 | 1 | 1 | 1 | 1 | 1 | 1 | 1 | 1 | 1 | 1 | 1 | 1 | 1 | 1 | 1 | 1 | 1 | 1 | 1 | 1 | 1 | 1 | 1 | 1 | 1 | 1 | 1 | 1 | 1 | 1 | 1 | 1 | 1 | 1 | 1 | 1 | 1 | 1 | 1 | 1 | 1 | 1 | 1 | 1 | 1 | 1 | 1 | 1 | 1 | 1 |
| *Bulbophyllum roxburghii* Rchb.f. | 1 | 1 | 1 | 1 | 1 | 1 | 1 | 1 | 1 | 1 | 1 | 1 | 1 | 1 | 1 | 1 | 1 | 1 | 1 | 1 | 1 | 1 | 1 | 1 | 1 | 1 | 1 | 1 | 1 | 1 | 1 | 1 | 1 | 1 | 1 | 1 | 1 | 1 | 1 | 1 | 1 | 1 | 1 | 1 | 1 | 1 | 1 | 1 | 1 | 0 | 1 | 1 | 1 | 1 | 1 | 1 | 0 | 1 | 1 | 1 | 1 | 1 | 1 |
| *Bulbophyllum rubipetalum* P.Royen | 1 | 1 | 1 | 1 | 1 | 1 | 1 | 1 | 1 | 1 | 1 | 1 | 1 | 1 | 1 | 1 | 1 | 1 | 1 | 1 | 1 | 1 | 1 | 1 | 1 | 1 | 1 | 1 | 1 | 1 | 1 | 1 | 1 | 1 | 1 | 1 | 1 | 1 | 1 | 1 | 1 | 1 | 1 | 1 | 1 | 1 | 1 | 1 | 1 | 1 | 1 | 1 | 1 | 1 | 1 | 1 | 1 | 0 | 1 | 0 | 1 | 1 | 1 |
| *Bulbophyllum rufinum* Rchb.f. | 1 | 1 | 1 | 1 | 1 | 1 | 1 | 1 | 1 | 1 | 1 | 1 | 1 | 1 | 1 | 1 | 1 | 1 | 1 | 1 | 1 | 1 | 1 | 1 | 1 | 1 | 1 | 1 | 1 | 1 | 1 | 1 | 1 | 1 | 1 | 1 | 1 | 1 | 1 | 1 | 1 | 1 | 1 | 0 | 1 | 1 | 1 | 1 | 1 | 1 | 1 | 1 | 1 | 1 | 1 | 1 | 1 | 1 | 1 | 1 | 1 | 1 | 1 |
| *Bulbophyllum rugosum* Ridl. | 1 | 1 | 1 | 1 | 1 | 1 | 1 | 1 | 1 | 1 | 1 | 1 | 1 | 1 | 1 | 1 | 1 | 1 | 1 | 1 | 1 | 1 | 1 | 1 | 1 | 1 | 1 | 1 | 1 | 1 | 1 | 1 | 1 | 1 | 1 | 1 | 1 | 1 | 1 | 1 | 1 | 1 | 0 | 1 | 1 | 1 | 1 | 1 | 1 | 1 | 1 | 1 | 1 | 1 | 1 | 1 | 1 | 1 | 1 | 1 | 1 | 1 | 1 |
| *Bulbophyllum saccoglossum* J.J.Verm., Schuit. & de Vogel | 1 | 1 | 1 | 1 | 1 | 1 | 1 | 1 | 1 | 1 | 1 | 1 | 1 | 1 | 1 | 1 | 1 | 1 | 1 | 1 | 1 | 1 | 1 | 1 | 1 | 1 | 1 | 1 | 1 | 1 | 1 | 1 | 1 | 1 | 1 | 1 | 1 | 1 | 1 | 1 | 1 | 1 | 1 | 1 | 1 | 1 | 1 | 1 | 1 | 0 | 1 | 1 | 1 | 1 | 1 | 1 | 0 | 1 | 1 | 1 | 1 | 1 | 1 |
| *Bulbophyllum saltatorium* Lindl. | 1 | 1 | 1 | 1 | 1 | 1 | 1 | 1 | 1 | 1 | 1 | 1 | 1 | 1 | 1 | 1 | 1 | 1 | 1 | 1 | 1 | 1 | 1 | 1 | 1 | 0 | 1 | 1 | 1 | 1 | 1 | 1 | 1 | 1 | 1 | 1 | 1 | 1 | 1 | 1 | 1 | 0 | 1 | 1 | 1 | 1 | 1 | 0 | 1 | 1 | 1 | 1 | 0 | 1 | 1 | 1 | 1 | 1 | 0 | 0 | 0 | 1 | 0 |
| *Bulbophyllum sandersonii* (Hook.f.) Rchb.f. | 1 | 1 | 1 | 1 | 1 | 1 | 1 | 1 | 1 | 1 | 1 | 1 | 1 | 1 | 1 | 1 | 1 | 1 | 1 | 1 | 1 | 1 | 1 | 1 | 1 | 1 | 1 | 1 | 1 | 1 | 1 | 1 | 1 | 1 | 1 | 1 | 1 | 1 | 1 | 1 | 1 | 1 | 1 | 1 | 1 | 1 | 1 | 1 | 1 | 1 | 1 | 1 | 1 | 1 | 1 | 1 | 1 | 1 | 1 | 0 | 1 | 1 | 1 |
| *Bulbophyllum sanguineopunctatum* Seidenf. & A.D.Kerr 1 | 1 | 1 | 1 | 1 | 1 | 1 | 1 | 1 | 1 | 1 | 1 | 1 | 1 | 1 | 1 | 1 | 1 | 1 | 1 | 1 | 1 | 1 | 1 | 1 | 1 | 1 | 1 | 1 | 1 | 1 | 1 | 1 | 1 | 1 | 1 | 1 | 1 | 1 | 1 | 1 | 1 | 1 | 0 | 1 | 1 | 1 | 1 | 1 | 1 | 1 | 1 | 1 | 1 | 1 | 1 | 1 | 1 | 1 | 1 | 1 | 1 | 1 | 1 |
| *Bulbophyllum sanguineopunctatum* Seidenf. & A.D.Kerr 2 | 1 | 1 | 1 | 1 | 1 | 1 | 1 | 1 | 1 | 1 | 1 | 1 | 1 | 1 | 1 | 1 | 1 | 1 | 1 | 1 | 1 | 1 | 1 | 1 | 1 | 1 | 1 | 1 | 1 | 1 | 1 | 1 | 1 | 1 | 1 | 1 | 1 | 1 | 1 | 1 | 1 | 1 | 0 | 1 | 1 | 1 | 1 | 1 | 1 | 1 | 1 | 1 | 1 | 1 | 1 | 1 | 1 | 1 | 1 | 1 | 1 | 1 | 1 |
| *Bulbophyllum santosii* Ames | 1 | 1 | 1 | 1 | 1 | 1 | 1 | 1 | 1 | 1 | 1 | 1 | 1 | 1 | 1 | 1 | 1 | 1 | 1 | 1 | 1 | 1 | 1 | 1 | 0 | 1 | 1 | 1 | 1 | 1 | 1 | 1 | 1 | 1 | 1 | 1 | 1 | 1 | 1 | 1 | 1 | 1 | 1 | 1 | 1 | 1 | 1 | 1 | 0 | 1 | 1 | 1 | 1 | 1 | 1 | 1 | 1 | 1 | 1 | 1 | 1 | 1 | 0 |
| *Bulbophyllum sauguetiense* Schltr. 1 | 1 | 1 | 1 | 1 | 1 | 1 | 1 | 1 | 1 | 1 | 1 | 1 | 1 | 1 | 1 | 1 | 1 | 1 | 1 | 1 | 1 | 1 | 1 | 1 | 1 | 1 | 1 | 1 | 1 | 1 | 1 | 1 | 1 | 1 | 1 | 1 | 1 | 1 | 1 | 1 | 1 | 1 | 0 | 1 | 1 | 1 | 1 | 1 | 1 | 1 | 1 | 1 | 1 | 1 | 1 | 1 | 1 | 1 | 1 | 1 | 1 | 1 | 1 |
| *Bulbophyllum sauguetiense* Schltr. 2 | 1 | 1 | 1 | 1 | 1 | 1 | 1 | 1 | 1 | 1 | 1 | 1 | 1 | 1 | 1 | 1 | 1 | 1 | 1 | 1 | 1 | 1 | 1 | 1 | 1 | 1 | 1 | 1 | 1 | 1 | 1 | 1 | 1 | 1 | 1 | 1 | 1 | 1 | 1 | 1 | 1 | 1 | 0 | 1 | 1 | 1 | 1 | 1 | 1 | 1 | 1 | 1 | 1 | 1 | 1 | 1 | 1 | 1 | 1 | 1 | 1 | 1 | 1 |

| | | | | | | | | | | | | | | | | | | | | | | | | | | | | | | | | | | | | | | | | | | | | | | | | | | | | | | | | | | | | | | | |
|---|---|---|---|---|---|---|---|---|---|---|---|---|---|---|---|---|---|---|---|---|---|---|---|---|---|---|---|---|---|---|---|---|---|---|---|---|---|---|---|---|---|---|---|---|---|---|---|---|---|---|---|---|---|---|---|---|---|---|---|---|---|---|---|
| *Bulbophyllum savaiense* Schltr. | 1 | 1 | 1 | 1 | 1 | 1 | 1 | 1 | 1 | 1 | 1 | 0 | 1 | 1 | 1 | 1 | 1 | 1 | 1 | 1 | 1 | 1 | 1 | 1 | 1 | 1 | 1 | 1 | 1 | 1 | 1 | 1 | 1 | 1 | 1 | 1 | 1 | 1 | 1 | 1 | 1 | 1 | 1 | 1 | 1 | 1 | 1 | 1 | 1 | 1 | 0 | 1 | 1 | 1 | 1 | 1 | 1 | 1 | 1 | 1 | 1 | 1 | 1 |
| *Bulbophyllum scaberulum* (Rolfe) Bolus | 0 | 1 | 1 | 1 | 1 | 1 | 1 | 1 | 1 | 1 | 1 | 1 | 1 | 1 | 0 | 1 | 1 | 1 | 1 | 1 | 1 | 1 | 1 | 1 | 1 | 1 | 1 | 1 | 1 | 1 | 1 | 1 | 1 | 1 | 1 | 1 | 1 | 1 | 1 | 1 | 1 | 1 | 1 | 1 | 1 | 1 | 1 | 1 | 1 | 1 | 1 | 1 | 1 | 1 | 1 | 1 | 1 | 1 | 1 | 0 | 1 | 1 | 0 |
| *Bulbophyllum scabratum* Rchb.f. | 1 | 1 | 1 | 1 | 1 | 1 | 1 | 1 | 1 | 1 | 1 | 1 | 1 | 1 | 1 | 1 | 1 | 1 | 1 | 1 | 1 | 1 | 1 | 1 | 1 | 1 | 1 | 1 | 1 | 1 | 1 | 1 | 1 | 1 | 1 | 1 | 1 | 1 | 1 | 1 | 1 | 1 | 1 | 1 | 1 | 1 | 1 | 1 | 1 | 1 | 1 | 1 | 1 | 1 | 1 | 1 | 1 | 1 | 1 | 1 | 1 | 1 | 1 |
| *Bulbophyllum schinzianum* Kraenzl. ex De Wild. & T.Durand 1 | 1 | 1 | 1 | 1 | 1 | 1 | 1 | 1 | 1 | 1 | 1 | 1 | 1 | 1 | 1 | 1 | 1 | 1 | 1 | 1 | 1 | 1 | 1 | 1 | 1 | 1 | 1 | 1 | 1 | 1 | 1 | 1 | 1 | 1 | 1 | 1 | 1 | 1 | 1 | 1 | 1 | 1 | 1 | 1 | 1 | 1 | 1 | 1 | 1 | 1 | 1 | 1 | 1 | 1 | 1 | 1 | 1 | 1 | 1 | 0 | 1 | 1 | 1 |
| *Bulbophyllum schinzianum* Kraenzl. ex De Wild. & T.Durand 2 | 1 | 1 | 1 | 1 | 1 | 1 | 1 | 1 | 1 | 1 | 1 | 1 | 1 | 1 | 1 | 1 | 1 | 1 | 1 | 1 | 1 | 1 | 1 | 1 | 1 | 1 | 1 | 1 | 1 | 1 | 1 | 1 | 1 | 1 | 1 | 1 | 1 | 1 | 1 | 1 | 1 | 1 | 1 | 1 | 1 | 1 | 1 | 1 | 1 | 1 | 1 | 1 | 1 | 1 | 0 | 1 | 1 | 1 | 1 | 0 | 1 | 1 | 1 |
| *Bulbophyllum schmidii* Garay | 1 | 1 | 1 | 1 | 1 | 1 | 1 | 1 | 1 | 1 | 1 | 1 | 1 | 1 | 1 | 1 | 1 | 1 | 1 | 1 | 1 | 1 | 1 | 1 | 1 | 1 | 1 | 1 | 1 | 1 | 1 | 1 | 1 | 1 | 1 | 1 | 1 | 1 | 1 | 1 | 1 | 1 | 1 | 1 | 1 | 1 | 1 | 1 | 1 | 1 | 1 | 1 | 1 | 1 | 1 | 1 | 1 | 1 | 1 | 1 | 1 | 1 | 1 |
| *Bulbophyllum scopa* J.J.Verm. | 1 | 1 | 1 | 1 | 1 | 1 | 1 | 1 | 1 | 1 | 1 | 1 | 1 | 1 | 1 | 1 | 1 | 1 | 1 | 1 | 1 | 1 | 1 | 1 | 1 | 1 | 1 | 1 | 1 | 1 | 1 | 1 | 1 | 1 | 1 | 1 | 1 | 1 | 1 | 1 | 1 | 1 | 1 | 1 | 1 | 1 | 1 | 1 | 1 | 1 | 1 | 1 | 1 | 1 | 1 | 1 | 1 | 1 | 1 | 1 | 1 | 1 | 1 |
| *Bulbophyllum scorpio* J.J.Verm. | 1 | 1 | 1 | 1 | 1 | 1 | 1 | 1 | 1 | 1 | 1 | 1 | 1 | 1 | 1 | 1 | 1 | 1 | 1 | 1 | 1 | 1 | 1 | 1 | 1 | 1 | 1 | 1 | 1 | 1 | 1 | 1 | 1 | 1 | 1 | 1 | 1 | 1 | 1 | 1 | 1 | 1 | 1 | 1 | 1 | 1 | 1 | 1 | 1 | 1 | 1 | 1 | 1 | 1 | 1 | 1 | 1 | 1 | 1 | 0 | 1 | 1 | 1 |
| *Bulbophyllum shanicum* King & Pantl. | 1 | 1 | 1 | 1 | 1 | 1 | 1 | 1 | 1 | 1 | 1 | 1 | 1 | 1 | 1 | 1 | 1 | 1 | 1 | 1 | 1 | 1 | 1 | 1 | 1 | 1 | 1 | 1 | 1 | 1 | 1 | 1 | 1 | 1 | 1 | 1 | 1 | 1 | 1 | 1 | 1 | 1 | 1 | 1 | 1 | 1 | 1 | 1 | 1 | 1 | 1 | 1 | 1 | 1 | 1 | 1 | 0 | 1 | 1 | 1 | 1 | 1 | 1 |
| *Bulbophyllum shepherdii* (F.Muell.) Rchb.f. | 1 | 1 | 1 | 1 | 1 | 1 | 1 | 1 | 1 | 1 | 1 | 1 | 1 | 1 | 1 | 1 | 1 | 1 | 1 | 1 | 1 | 1 | 1 | 1 | 1 | 1 | 1 | 1 | 1 | 1 | 1 | 1 | 1 | 1 | 1 | 1 | 1 | 1 | 1 | 1 | 1 | 1 | 1 | 1 | 1 | 1 | 1 | 1 | 1 | 1 | 1 | 1 | 1 | 1 | 1 | 1 | 1 | 1 | 1 | 1 | 1 | 1 | 1 |
| *Bulbophyllum sibuyanense* Ames | 1 | 1 | 1 | 1 | 1 | 1 | 1 | 1 | 1 | 1 | 1 | 1 | 1 | 1 | 1 | 1 | 1 | 1 | 1 | 1 | 1 | 1 | 1 | 1 | 1 | 0 | 0 | 1 | 1 | 1 | 1 | 1 | 1 | 1 | 1 | 1 | 1 | 0 | 1 | 1 | 1 | 1 | 1 | 1 | 1 | 1 | 1 | 1 | 1 | 1 | 1 | 1 | 1 | 1 | 1 | 1 | 0 | 0 | 1 | 1 | 1 | 1 | 1 |
| *Bulbophyllum sigaldiae* Guillaumin | 1 | 1 | 1 | 1 | 1 | 1 | 1 | 1 | 1 | 1 | 1 | 1 | 1 | 1 | 1 | 1 | 1 | 1 | 1 | 1 | 1 | 1 | 1 | 1 | 1 | 1 | 1 | 1 | 1 | 1 | 1 | 1 | 1 | 1 | 1 | 1 | 1 | 1 | 1 | 1 | 1 | 1 | 0 | 1 | 1 | 1 | 1 | 0 | 1 | 1 | 1 | 1 | 1 | 1 | 1 | 1 | 1 | 1 | 1 | 1 | 1 | 1 | 1 |
| *Bulbophyllum simile* Schltr. | 1 | 1 | 1 | 1 | 1 | 1 | 1 | 1 | 1 | 1 | 1 | 1 | 1 | 1 | 1 | 1 | 1 | 1 | 1 | 1 | 1 | 1 | 1 | 1 | 1 | 0 | 1 | 1 | 1 | 1 | 1 | 1 | 1 | 1 | 1 | 1 | 1 | 1 | 1 | 1 | 1 | 1 | 1 | 1 | 1 | 1 | 1 | 1 | 1 | 0 | 1 | 1 | 1 | 1 | 1 | 1 | 1 | 1 | 1 | 1 | 1 | 0 | 1 |
| *Bulbophyllum skeatianum* Ridl. | 1 | 1 | 1 | 1 | 1 | 1 | 1 | 1 | 1 | 1 | 1 | 1 | 1 | 1 | 1 | 1 | 1 | 1 | 1 | 1 | 1 | 1 | 1 | 1 | 1 | 1 | 1 | 1 | 1 | 1 | 1 | 1 | 1 | 1 | 1 | 1 | 1 | 1 | 1 | 1 | 1 | 1 | 1 | 1 | 1 | 1 | 1 | 1 | 1 | 1 | 1 | 1 | 1 | 1 | 1 | 1 | 1 | 1 | 1 | 1 | 1 | 1 | 1 |
| *Bulbophyllum smitinandii* Seidenf. & Thorut | 1 | 1 | 1 | 1 | 1 | 1 | 1 | 1 | 1 | 1 | 1 | 1 | 1 | 1 | 1 | 1 | 1 | 1 | 1 | 1 | 1 | 1 | 1 | 1 | 1 | 1 | 1 | 1 | 1 | 1 | 1 | 1 | 1 | 1 | 1 | 1 | 1 | 1 | 1 | 1 | 0 | 1 | 1 | 1 | 1 | 1 | 1 | 1 | 1 | 0 | 1 | 1 | 1 | 1 | 1 | 1 | 0 | 1 | 1 | 1 | 1 | 0 | 1 |
| *Bulbophyllum* sp. 1 | 1 | 1 | 1 | 1 | 1 | 1 | 1 | 1 | 1 | 1 | 1 | 1 | 1 | 1 | 1 | 1 | 1 | 1 | 1 | 1 | 1 | 1 | 1 | 1 | 0 | 1 | 1 | 1 | 1 | 0 | 1 | 1 | 1 | 1 | 1 | 1 | 1 | 1 | 1 | 1 | 1 | 1 | 1 | 1 | 1 | 1 | 1 | 1 | 0 | 1 | 0 | 1 | 1 | 1 | 1 | 1 | 1 | 1 | 0 | 0 | 1 | 1 | 0 |
| *Bulbophyllum* sp. 2 | 1 | 1 | 1 | 1 | 1 | 1 | 1 | 1 | 1 | 1 | 1 | 1 | 1 | 1 | 1 | 1 | 1 | 1 | 1 | 1 | 1 | 1 | 1 | 1 | 1 | 1 | 1 | 0 | 1 | 0 | 1 | 1 | 1 | 1 | 1 | 1 | 1 | 1 | 1 | 1 | 1 | 1 | 1 | 1 | 0 | 1 | 1 | 1 | 1 | 1 | 0 | 0 | 1 | 1 | 1 | 1 | 1 | 1 | 1 | 0 | 1 | 1 | 0 |
| *Bulbophyllum* sp. 3 | 0 | 1 | 0 | 0 | 1 | 1 | 1 | 1 | 0 | 0 | 1 | 1 | 0 | 1 | 1 | 0 | 0 | 1 | 1 | 1 | 1 | 0 | 0 | 1 | 0 | 1 | 1 | 0 | 0 | 1 | 1 | 0 | 1 | 0 | 1 | 1 | 1 | 1 | 0 | 1 | 1 | 0 | 1 | 1 | 1 | 0 | 1 | 1 | 1 | 1 | 1 | 1 | 1 | 1 | 1 | 1 | 1 | 1 | 1 | 1 | 0 | 1 | 0 |
| *Bulbophyllum* sp. 4 | 1 | 1 | 1 | 1 | 1 | 1 | 1 | 1 | 1 | 1 | 1 | 1 | 1 | 1 | 1 | 1 | 1 | 1 | 1 | 1 | 1 | 1 | 1 | 1 | 1 | 1 | 1 | 1 | 1 | 1 | 0 | 1 | 1 | 1 | 1 | 1 | 1 | 1 | 1 | 1 | 1 | 1 | 1 | 1 | 1 | 1 | 1 | 1 | 1 | 1 | 1 | 1 | 1 | 1 | 1 | 1 | 1 | 1 | 1 | 1 | 1 | 1 | 1 |
| *Bulbophyllum* sp. 5 | 1 | 1 | 1 | 1 | 1 | 1 | 1 | 1 | 1 | 1 | 1 | 1 | 1 | 1 | 1 | 1 | 1 | 1 | 1 | 1 | 1 | 1 | 1 | 1 | 1 | 1 | 1 | 1 | 1 | 1 | 0 | 1 | 1 | 1 | 1 | 1 | 1 | 1 | 1 | 1 | 1 | 1 | 1 | 1 | 1 | 1 | 1 | 1 | 1 | 1 | 1 | 1 | 1 | 1 | 1 | 1 | 1 | 1 | 1 | 1 | 1 | 1 | 1 |
| *Bulbophyllum* sp. 6 | 1 | 1 | 1 | 1 | 1 | 1 | 1 | 1 | 1 | 1 | 1 | 1 | 1 | 1 | 1 | 1 | 1 | 1 | 1 | 1 | 1 | 1 | 1 | 1 | 1 | 1 | 1 | 1 | 1 | 1 | 1 | 1 | 1 | 1 | 1 | 1 | 1 | 1 | 1 | 1 | 1 | 1 | 1 | 1 | 1 | 1 | 1 | 1 | 1 | 1 | 1 | 1 | 1 | 1 | 1 | 1 | 1 | 1 | 1 | 1 | 1 | 1 | 1 |
| *Bulbophyllum* sp. 7 | 1 | 1 | 1 | 1 | 1 | 1 | 1 | 1 | 1 | 1 | 1 | 1 | 1 | 1 | 1 | 1 | 1 | 1 | 1 | 1 | 1 | 1 | 1 | 1 | 1 | 1 | 1 | 1 | 1 | 1 | 1 | 1 | 1 | 1 | 1 | 1 | 1 | 1 | 1 | 1 | 1 | 1 | 1 | 1 | 1 | 1 | 1 | 1 | 1 | 1 | 1 | 1 | 1 | 1 | 1 | 1 | 1 | 1 | 1 | 1 | 1 | 1 | 1 |
| *Bulbophyllum* sp. 8 | 1 | 1 | 1 | 1 | 1 | 1 | 1 | 1 | 1 | 1 | 1 | 1 | 1 | 1 | 1 | 1 | 1 | 1 | 1 | 1 | 1 | 1 | 1 | 1 | 1 | 1 | 1 | 1 | 1 | 1 | 1 | 1 | 1 | 1 | 1 | 1 | 1 | 1 | 1 | 1 | 1 | 1 | 1 | 1 | 1 | 1 | 1 | 1 | 1 | 1 | 1 | 1 | 1 | 1 | 1 | 1 | 1 | 1 | 1 | 1 | 1 | 1 | 1 |
| *Bulbophyllum* sp. 9 | 1 | 1 | 1 | 1 | 1 | 1 | 1 | 0 | 1 | 1 | 1 | 1 | 1 | 1 | 1 | 1 | 1 | 0 | 1 | 1 | 1 | 1 | 1 | 1 | 1 | 1 | 1 | 1 | 1 | 1 | 1 | 1 | 1 | 1 | 1 | 1 | 1 | 1 | 1 | 1 | 1 | 1 | 1 | 1 | 1 | 1 | 1 | 1 | 0 | 0 | 1 | 1 | 1 | 1 | 1 | 1 | 0 | 1 | 1 | 1 | 1 | 1 | 1 |
| *Bulbophyllum* sp. 10 | 1 | 1 | 1 | 1 | 1 | 1 | 1 | 1 | 1 | 1 | 1 | 1 | 1 | 1 | 1 | 1 | 1 | 1 | 1 | 1 | 1 | 1 | 1 | 1 | 1 | 1 | 1 | 1 | 1 | 1 | 1 | 1 | 1 | 1 | 1 | 1 | 1 | 1 | 1 | 1 | 1 | 1 | 1 | 1 | 1 | 1 | 1 | 1 | 1 | 1 | 1 | 1 | 1 | 1 | 1 | 1 | 1 | 1 | 1 | 1 | 1 | 1 | 1 |
| *Bulbophyllum* sp. 11 | 1 | 1 | 1 | 1 | 1 | 1 | 1 | 1 | 1 | 1 | 1 | 1 | 1 | 1 | 1 | 1 | 1 | 1 | 1 | 1 | 1 | 1 | 1 | 1 | 1 | 1 | 1 | 1 | 1 | 1 | 1 | 1 | 1 | 1 | 1 | 1 | 1 | 1 | 1 | 1 | 1 | 1 | 1 | 1 | 1 | 1 | 1 | 1 | 1 | 1 | 1 | 1 | 1 | 1 | 1 | 1 | 1 | 1 | 1 | 1 | 1 | 1 | 1 |
| *Bulbophyllum* sp. 12 | 1 | 1 | 1 | 1 | 1 | 1 | 1 | 1 | 1 | 1 | 1 | 1 | 1 | 1 | 1 | 1 | 1 | 1 | 1 | 1 | 1 | 1 | 1 | 1 | 1 | 1 | 1 | 1 | 1 | 1 | 1 | 1 | 1 | 1 | 1 | 1 | 1 | 1 | 1 | 1 | 1 | 1 | 1 | 1 | 1 | 1 | 1 | 1 | 1 | 1 | 1 | 1 | 1 | 1 | 1 | 1 | 1 | 1 | 1 | 1 | 1 | 1 | 1 |
| *Bulbophyllum spathulatum* (Rolfe ex E.Cooper) Seidenf. | 1 | 1 | 1 | 1 | 1 | 1 | 1 | 1 | 1 | 1 | 1 | 1 | 1 | 1 | 1 | 1 | 1 | 1 | 1 | 1 | 1 | 1 | 1 | 1 | 1 | 1 | 1 | 1 | 1 | 1 | 1 | 1 | 1 | 1 | 1 | 1 | 1 | 1 | 1 | 1 | 1 | 1 | 1 | 1 | 1 | 1 | 1 | 1 | 1 | 0 | 1 | 1 | 1 | 1 | 1 | 1 | 0 | 1 | 1 | 1 | 1 | 1 | 1 |
| *Bulbophyllum speciosum* Schltr. | 1 | 1 | 1 | 1 | 1 | 1 | 1 | 1 | 1 | 1 | 1 | 1 | 1 | 1 | 1 | 1 | 1 | 1 | 1 | 1 | 1 | 1 | 1 | 1 | 1 | 1 | 1 | 1 | 1 | 1 | 1 | 1 | 1 | 1 | 1 | 1 | 1 | 1 | 1 | 1 | 1 | 1 | 1 | 1 | 1 | 1 | 1 | 1 | 1 | 0 | 1 | 1 | 1 | 1 | 1 | 1 | 0 | 1 | 1 | 1 | 1 | 1 | 1 |
| *Bulbophyllum spodotriche* J.J.Verm. & P.O'Byrne | 1 | 1 | 1 | 1 | 1 | 1 | 1 | 1 | 1 | 1 | 1 | 1 | 1 | 1 | 1 | 1 | 1 | 1 | 1 | 1 | 1 | 1 | 1 | 1 | 1 | 1 | 1 | 1 | 1 | 1 | 1 | 1 | 1 | 1 | 1 | 1 | 1 | 1 | 1 | 1 | 1 | 1 | 1 | 1 | 1 | 1 | 1 | 1 | 1 | 1 | 1 | 1 | 1 | 1 | 1 | 1 | 1 | 1 | 1 | 1 | 1 | 1 | 1 |
| *Bulbophyllum steyermarkii* Foldats | 1 | 1 | 1 | 1 | 1 | 1 | 1 | 1 | 1 | 1 | 1 | 1 | 1 | 1 | 1 | 1 | 1 | 1 | 1 | 1 | 1 | 1 | 1 | 1 | 1 | 1 | 1 | 1 | 1 | 1 | 1 | 1 | 1 | 1 | 1 | 1 | 1 | 1 | 1 | 1 | 1 | 1 | 1 | 1 | 1 | 1 | 1 | 1 | 1 | 1 | 1 | 1 | 1 | 1 | 1 | 1 | 0 | 1 | 1 | 1 | 1 | 1 | 1 |
| *Bulbophyllum subpatulum* J.J.Verm. | 1 | 1 | 1 | 1 | 1 | 1 | 1 | 1 | 1 | 1 | 1 | 1 | 1 | 1 | 1 | 1 | 1 | 1 | 1 | 1 | 1 | 1 | 1 | 1 | 1 | 1 | 1 | 1 | 1 | 1 | 1 | 1 | 1 | 1 | 1 | 1 | 1 | 1 | 1 | 1 | 1 | 1 | 0 | 1 | 1 | 1 | 1 | 1 | 1 | 1 | 1 | 1 | 1 | 1 | 1 | 1 | 1 | 1 | 1 | 0 | 1 | 1 | 1 |
| *Bulbophyllum sukhakulii* Seidenf. | 1 | 1 | 1 | 1 | 1 | 1 | 1 | 1 | 1 | 1 | 1 | 1 | 1 | 1 | 1 | 1 | 1 | 1 | 1 | 1 | 1 | 1 | 1 | 1 | 1 | 1 | 1 | 1 | 1 | 1 | 1 | 1 | 1 | 1 | 1 | 1 | 1 | 1 | 1 | 1 | 1 | 1 | 1 | 1 | 1 | 1 | 1 | 1 | 1 | 1 | 1 | 1 | 1 | 1 | 1 | 1 | 1 | 1 | 1 | 1 | 1 | 1 | 1 |
| *Bulbophyllum taiwanense* (Fukuy.) K.Nakaj. | 1 | 1 | 1 | 1 | 1 | 1 | 1 | 1 | 1 | 1 | 1 | 1 | 1 | 1 | 1 | 1 | 1 | 1 | 1 | 1 | 1 | 0 | 1 | 1 | 1 | 1 | 1 | 1 | 1 | 1 | 1 | 1 | 1 | 1 | 1 | 1 | 1 | 1 | 1 | 1 | 1 | 1 | 1 | 1 | 1 | 1 | 1 | 1 | 1 | 0 | 1 | 1 | 1 | 1 | 1 | 1 | 1 | 1 | 1 | 1 | 1 | 1 | 1 |
| *Bulbophyllum tentaculiferum* Schltr. | 1 | 1 | 1 | 1 | 1 | 1 | 1 | 1 | 1 | 1 | 1 | 1 | 1 | 1 | 1 | 1 | 1 | 1 | 1 | 1 | 1 | 1 | 1 | 1 | 1 | 1 | 1 | 1 | 1 | 1 | 1 | 1 | 0 | 1 | 1 | 1 | 1 | 1 | 1 | 1 | 1 | 1 | 1 | 1 | 1 | 1 | 1 | 1 | 1 | 0 | 1 | 1 | 1 | 1 | 1 | 1 | 0 | 1 | 1 | 1 | 1 | 1 | 1 |
| *Bulbophyllum thaiorum* J.J.Sm. | 1 | 1 | 1 | 1 | 1 | 1 | 1 | 1 | 1 | 1 | 1 | 1 | 1 | 1 | 1 | 1 | 1 | 1 | 1 | 1 | 1 | 1 | 1 | 1 | 1 | 1 | 1 | 1 | 1 | 1 | 1 | 1 | 1 | 1 | 1 | 1 | 1 | 1 | 1 | 1 | 1 | 1 | 1 | 1 | 1 | 1 | 1 | 1 | 1 | 1 | 1 | 1 | 1 | 1 | 1 | 1 | 1 | 1 | 1 | 1 | 1 | 1 | 1 |
| *Bulbophyllum thiurum* J.J.Verm. & P.O'Byrne | 1 | 1 | 1 | 1 | 1 | 1 | 1 | 1 | 1 | 1 | 1 | 1 | 1 | 1 | 1 | 1 | 1 | 1 | 1 | 1 | 1 | 1 | 1 | 1 | 1 | 1 | 1 | 1 | 1 | 1 | 1 | 1 | 1 | 1 | 1 | 1 | 1 | 1 | 1 | 1 | 1 | 1 | 1 | 1 | 1 | 1 | 1 | 1 | 1 | 1 | 1 | 1 | 1 | 1 | 1 | 1 | 1 | 1 | 1 | 1 | 1 | 1 | 1 |
| *Bulbophyllum tianguii* K.Y.Lang & D.Luo | 1 | 1 | 1 | 1 | 1 | 1 | 1 | 1 | 1 | 1 | 1 | 1 | 1 | 1 | 1 | 1 | 1 | 1 | 1 | 1 | 1 | 1 | 1 | 1 | 1 | 1 | 1 | 1 | 1 | 1 | 1 | 1 | 1 | 1 | 1 | 1 | 1 | 1 | 1 | 1 | 1 | 1 | 1 | 1 | 1 | 1 | 1 | 1 | 1 | 1 | 1 | 1 | 1 | 1 | 1 | 1 | 0 | 1 | 1 | 1 | 1 | 1 | 1 |
| *Bulbophyllum tollenoniferum* J.J.Sm. | 1 | 1 | 1 | 1 | 1 | 1 | 1 | 1 | 1 | 1 | 1 | 1 | 1 | 1 | 1 | 1 | 1 | 1 | 1 | 1 | 1 | 1 | 1 | 1 | 1 | 1 | 1 | 1 | 1 | 1 | 1 | 1 | 1 | 1 | 1 | 1 | 1 | 1 | 1 | 1 | 1 | 1 | 1 | 1 | 1 | 1 | 1 | 1 | 1 | 1 | 1 | 1 | 1 | 1 | 1 | 1 | 1 | 1 | 1 | 1 | 1 | 1 | 1 |
| *Bulbophyllum tortuosum* (Blume) Lindl. | 0 | 1 | 1 | 1 | 0 | 1 | 1 | 0 | 1 | 1 | 1 | 0 | 1 | 1 | 1 | 1 | 1 | 1 | 1 | 1 | 1 | 1 | 1 | 1 | 1 | 1 | 1 | 1 | 1 | 1 | 1 | 1 | 0 | 1 | 1 | 1 | 1 | 1 | 1 | 1 | 1 | 1 | 0 | 1 | 0 | 1 | 1 | 1 | 1 | 1 | 0 | 0 | 1 | 1 | 1 | 1 | 1 | 1 | 0 | 0 | 1 | 1 | 1 |
| *Bulbophyllum trachypus* Schltr. | 1 | 1 | 1 | 1 | 1 | 1 | 1 | 0 | 1 | 0 | 0 | 1 | 1 | 0 | 0 | 1 | 1 | 1 | 1 | 1 | 0 | 1 | 1 | 0 | 0 | 1 | 1 | 1 | 1 | 0 | 1 | 1 | 1 | 1 | 1 | 0 | 0 | 1 | 1 | 0 | 0 | 0 | 0 | 0 | 0 | 0 | 0 | 0 | 1 | 1 | 1 | 1 | 0 | 0 | 0 | 0 | 0 | 1 | 0 | 0 | 0 | 1 | 1 |
| *Bulbophyllum treschii* Jenny 1 | 1 | 1 | 1 | 1 | 1 | 1 | 1 | 1 | 0 | 1 | 0 | 0 | 1 | 1 | 1 | 1 | 1 | 0 | 1 | 0 | 1 | 1 | 1 | 0 | 1 | 0 | 0 | 1 | 1 | 1 | 0 | 1 | 1 | 1 | 0 | 1 | 1 | 0 | 1 | 0 | 0 | 1 | 0 | 0 | 1 | 1 | 1 | 1 | 1 | 1 | 1 | 1 | 0 | 1 | 0 | 1 | 1 | 0 | 1 | 1 | 1 | 1 | 0 |
| *Bulbophyllum treschii* Jenny 2 | 1 | 1 | 1 | 1 | 1 | 1 | 1 | 1 | 1 | 1 | 1 | 1 | 1 | 1 | 1 | 1 | 1 | 1 | 1 | 1 | 1 | 1 | 1 | 1 | 1 | 1 | 1 | 1 | 1 | 1 | 1 | 1 | 1 | 1 | 1 | 1 | 1 | 1 | 1 | 1 | 1 | 1 | 1 | 1 | 1 | 1 | 1 | 1 | 1 | 1 | 1 | 1 | 1 | 1 | 1 | 1 | 1 | 1 | 1 | 1 | 1 | 1 | 1 |
| *Bulbophyllum triaristella* Schltr. | 1 | 1 | 1 | 1 | 1 | 1 | 1 | 1 | 1 | 1 | 1 | 1 | 1 | 1 | 1 | 1 | 1 | 1 | 1 | 1 | 1 | 1 | 1 | 1 | 1 | 1 | 1 | 1 | 1 | 1 | 1 | 1 | 1 | 1 | 1 | 1 | 1 | 1 | 1 | 1 | 1 | 1 | 0 | 1 | 1 | 1 | 1 | 1 | 1 | 1 | 1 | 1 | 1 | 1 | 1 | 1 | 1 | 1 | 1 | 1 | 1 | 1 | 1 |
| *Bulbophyllum tridentatum* Kraenzl. | 1 | 1 | 1 | 1 | 1 | 1 | 1 | 1 | 1 | 1 | 1 | 1 | 1 | 0 | 1 | 1 | 1 | 1 | 1 | 1 | 1 | 1 | 1 | 1 | 1 | 1 | 1 | 1 | 1 | 1 | 1 | 1 | 1 | 1 | 1 | 1 | 1 | 1 | 1 | 1 | 1 | 1 | 1 | 1 | 1 | 1 | 1 | 1 | 1 | 0 | 1 | 1 | 1 | 1 | 1 | 1 | 1 | 1 | 1 | 1 | 1 | 1 | 1 |
| *Bulbophyllum trifilum* J.J.Sm. | 1 | 0 | 1 | 1 | 1 | 1 | 1 | 1 | 1 | 0 | 1 | 1 | 0 | 0 | 1 | 1 | 1 | 1 | 0 | 1 | 0 | 0 | 1 | 1 | 1 | 1 | 1 | 1 | 1 | 1 | 1 | 1 | 1 | 1 | 1 | 0 | 1 | 1 | 0 | 1 | 1 | 1 | 1 | 1 | 1 | 1 | 1 | 1 | 1 | 0 | 1 | 1 | 1 | 1 | 1 | 1 | 1 | 1 | 1 | 1 | 1 | 1 | 1 |
| *Bulbophyllum tripudians* C.S.P.Parish & Rchb.f. | 1 | 1 | 1 | 1 | 1 | 1 | 1 | 1 | 1 | 1 | 1 | 1 | 1 | 1 | 1 | 1 | 1 | 1 | 1 | 1 | 1 | 1 | 1 | 1 | 1 | 1 | 1 | 1 | 1 | 1 | 1 | 1 | 1 | 1 | 1 | 1 | 1 | 1 | 1 | 1 | 1 | 1 | 0 | 1 | 1 | 1 | 1 | 1 | 1 | 1 | 0 | 1 | 1 | 1 | 1 | 1 | 1 | 1 | 1 | 1 | 1 | 1 | 1 |
| *Bulbophyllum triste* (Rolfe) Schltr. 1 | 1 | 1 | 1 | 1 | 1 | 1 | 1 | 1 | 1 | 1 | 1 | 1 | 1 | 1 | 1 | 1 | 1 | 1 | 1 | 1 | 1 | 1 | 1 | 1 | 1 | 1 | 1 | 1 | 1 | 1 | 1 | 1 | 1 | 1 | 1 | 1 | 1 | 1 | 1 | 1 | 1 | 1 | 1 | 1 | 1 | 1 | 1 | 1 | 1 | 1 | 1 | 1 | 1 | 1 | 1 | 1 | 0 | 1 | 1 | 1 | 1 | 1 | 1 |
| *Bulbophyllum triste* Rchb.f. 2 | 1 | 1 | 1 | 1 | 1 | 1 | 1 | 1 | 1 | 1 | 1 | 1 | 1 | 1 | 1 | 1 | 1 | 1 | 1 | 1 | 1 | 1 | 0 | 1 | 1 | 1 | 1 | 1 | 1 | 1 | 1 | 1 | 1 | 1 | 1 | 1 | 1 | 1 | 1 | 1 | 1 | 0 | 1 | 1 | 1 | 1 | 1 | 1 | 1 | 1 | 1 | 1 | 1 | 1 | 1 | 1 | 1 | 1 | 1 | 1 | 1 | 1 | 1 |
| *Bulbophyllum tuberculatum* Colenso | 1 | 1 | 1 | 1 | 1 | 1 | 1 | 1 | 1 | 1 | 1 | 1 | 1 | 1 | 1 | 1 | 1 | 1 | 1 | 1 | 1 | 1 | 1 | 1 | 1 | 1 | 1 | 1 | 1 | 1 | 1 | 1 | 1 | 1 | 1 | 1 | 1 | 1 | 1 | 1 | 1 | 1 | 1 | 1 | 1 | 1 | 1 | 1 | 1 | 1 | 0 | 1 | 1 | 1 | 1 | 0 | 1 | 1 | 1 | 0 | 1 | 1 | 1 |
| *Bulbophyllum umbellatum* Lindl. | 0 | 1 | 1 | 1 | 1 | 1 | 1 | 1 | 1 | 1 | 1 | 1 | 1 | 1 | 1 | 1 | 1 | 1 | 1 | 1 | 1 | 1 | 1 | 1 | 1 | 1 | 1 | 1 | 1 | 1 | 1 | 1 | 1 | 1 | 1 | 1 | 1 | 1 | 1 | 1 | 1 | 1 | 1 | 1 | 1 | 1 | 1 | 1 | 1 | 1 | 1 | 1 | 1 | 1 | 1 | 1 | 1 | 1 | 1 | 1 | 1 | 1 | 1 |
| *Bulbophyllum unicaudatum* Schltr. | 1 | 1 | 1 | 1 | 1 | 1 | 1 | 1 | 1 | 1 | 1 | 1 | 1 | 0 | 1 | 1 | 1 | 0 | 1 | 0 | 1 | 1 | 0 | 1 | 1 | 1 | 1 | 1 | 1 | 1 | 1 | 0 | 0 | 1 | 1 | 1 | 1 | 1 | 1 | 1 | 0 | 1 | 1 | 1 | 1 | 1 | 1 | 1 | 1 | 0 | 1 | 1 | 1 | 1 | 1 | 1 | 0 | 1 | 1 | 1 | 1 | 1 | 1 |
| *Bulbophyllum uniflorum* (Blume) Hassk. | 1 | 0 | 1 | 1 | 0 | 0 | 1 | 1 | 1 | 1 | 1 | 1 | 1 | 1 | 1 | 1 | 1 | 1 | 1 | 1 | 1 | 1 | 1 | 0 | 1 | 0 | 0 | 1 | 1 | 1 | 0 | 1 | 1 | 1 | 1 | 1 | 1 | 1 | 1 | 1 | 1 | 1 | 1 | 1 | 1 | 1 | 1 | 1 | 1 | 1 | 1 | 1 | 1 | 1 | 1 | 1 | 1 | 0 | 0 | 1 | 1 | 1 | 0 |
| *Bulbophyllum unitubum* J.J.Sm. | 1 | 1 | 1 | 1 | 1 | 1 | 1 | 0 | 1 | 1 | 1 | 1 | 1 | 1 | 1 | 1 | 1 | 1 | 1 | 1 | 0 | 1 | 1 | 1 | 1 | 1 | 1 | 1 | 1 | 1 | 1 | 1 | 1 | 1 | 1 | 1 | 1 | 1 | 1 | 0 | 1 | 1 | 1 | 0 | 0 | 1 | 1 | 1 | 1 | 1 | 1 | 1 | 1 | 0 | 1 | 1 | 0 | 1 | 0 | 1 | 1 | 1 | 1 |
| *Bulbophyllum vaginatum* Rchb.f. | 1 | 1 | 1 | 1 | 1 | 1 | 0 | 1 | 1 | 1 | 1 | 1 | 1 | 1 | 1 | 1 | 1 | 1 | 1 | 1 | 1 | 1 | 1 | 1 | 1 | 1 | 1 | 1 | 1 | 1 | 1 | 1 | 1 | 1 | 1 | 1 | 1 | 1 | 0 | 1 | 1 | 1 | 1 | 1 | 1 | 1 | 1 | 1 | 1 | 1 | 1 | 1 | 1 | 1 | 1 | 1 | 1 | 1 | 1 | 1 | 1 | 1 | 1 |
| *Bulbophyllum veldkampii* J.J.Verm. & P.O'Byrne | 1 | 1 | 1 | 1 | 1 | 1 | 1 | 1 | 1 | 1 | 1 | 1 | 1 | 1 | 1 | 1 | 1 | 1 | 1 | 1 | 1 | 1 | 1 | 1 | 1 | 1 | 1 | 1 | 1 | 1 | 1 | 1 | 1 | 1 | 1 | 1 | 1 | 1 | 1 | 1 | 1 | 1 | 1 | 1 | 1 | 1 | 1 | 1 | 1 | 1 | 1 | 1 | 1 | 1 | 1 | 1 | 1 | 1 | 1 | 1 | 1 | 1 | 1 |
| *Bulbophyllum verrucosum* (L.O.Williams) J.J.Verm., Schuit. & de Vogel | 1 | 1 | 1 | 1 | 1 | 1 | 1 | 1 | 1 | 1 | 1 | 1 | 1 | 1 | 1 | 1 | 1 | 1 | 1 | 1 | 1 | 1 | 1 | 1 | 1 | 1 | 1 | 1 | 1 | 1 | 1 | 1 | 1 | 1 | 1 | 1 | 1 | 1 | 1 | 1 | 0 | 1 | 1 | 1 | 1 | 1 | 1 | 1 | 1 | 1 | 1 | 1 | 1 | 1 | 1 | 1 | 1 | 1 | 1 | 1 | 1 | 1 | 1 |
| *Bulbophyllum versteegii* J.J.Sm. | 1 | 1 | 1 | 1 | 1 | 1 | 1 | 1 | 1 | 1 | 1 | 1 | 1 | 1 | 1 | 1 | 1 | 1 | 1 | 1 | 1 | 1 | 1 | 1 | 1 | 1 | 1 | 1 | 1 | 1 | 1 | 1 | 1 | 1 | 1 | 1 | 1 | 1 | 1 | 1 | 1 | 1 | 1 | 1 | 1 | 1 | 1 | 1 | 1 | 0 | 1 | 1 | 1 | 1 | 1 | 1 | 0 | 1 | 1 | 1 | 1 | 1 | 1 |
| *Bulbophyllum violaceolabellum* Seidenf. 1 | 1 | 1 | 1 | 1 | 1 | 1 | 1 | 1 | 1 | 1 | 1 | 1 | 1 | 1 | 1 | 1 | 1 | 1 | 1 | 1 | 1 | 1 | 1 | 1 | 1 | 1 | 1 | 1 | 1 | 1 | 1 | 1 | 1 | 1 | 1 | 1 | 1 | 1 | 1 | 1 | 1 | 1 | 1 | 1 | 1 | 1 | 1 | 1 | 1 | 0 | 1 | 1 | 1 | 1 | 1 | 1 | 0 | 1 | 1 | 1 | 1 | 1 | 1 |
| *Bulbophyllum violaceolabellum* Seidenf. 2 | 1 | 1 | 1 | 1 | 0 | 1 | 1 | 1 | 1 | 1 | 1 | 1 | 1 | 1 | 1 | 1 | 1 | 1 | 1 | 1 | 1 | 1 | 1 | 1 | 1 | 1 | 1 | 1 | 1 | 1 | 1 | 1 | 1 | 1 | 1 | 1 | 1 | 1 | 1 | 1 | 1 | 1 | 1 | 1 | 1 | 1 | 1 | 1 | 1 | 1 | 1 | 1 | 1 | 1 | 1 | 1 | 0 | 1 | 1 | 1 | 1 | 1 | 1 |
| *Bulbophyllum wadsworthii* Dockrill | 1 | 1 | 1 | 1 | 1 | 1 | 1 | 1 | 1 | 1 | 1 | 1 | 1 | 1 | 1 | 1 | 1 | 1 | 1 | 1 | 1 | 1 | 1 | 1 | 1 | 1 | 1 | 1 | 1 | 1 | 1 | 1 | 1 | 1 | 1 | 1 | 1 | 1 | 1 | 1 | 1 | 1 | 1 | 1 | 1 | 1 | 1 | 1 | 1 | 1 | 1 | 1 | 1 | 1 | 1 | 1 | 1 | 1 | 1 | 1 | 1 | 1 | 1 |

| | | | | | | | | | | | | | | | | | | | | | | | | | | | | | | | | | | | | | | | | | | | | | | | | | | | | | | | | | | | | | | | |
|---|---|---|---|---|---|---|---|---|---|---|---|---|---|---|---|---|---|---|---|---|---|---|---|---|---|---|---|---|---|---|---|---|---|---|---|---|---|---|---|---|---|---|---|---|---|---|---|---|---|---|---|---|---|---|---|---|---|---|---|---|---|---|---|
| *Bulbophyllum wakoi* Howcroft | 1 | 1 | 1 | 1 | 1 | 1 | 1 | 1 | 1 | 1 | 1 | 1 | 1 | 1 | 1 | 1 | 1 | 1 | 1 | 1 | 1 | 1 | 1 | 1 | 1 | 1 | 1 | 1 | 1 | 1 | 1 | 1 | 1 | 1 | 1 | 1 | 1 | 1 | 1 | 1 | 1 | 1 | 1 | 1 | 1 | 1 | 1 | 1 | 1 | 0 | 1 | 1 | 1 | 1 | 1 | 1 | 0 | 1 | 1 | 1 | 1 | 1 | 1 |
| *Bulbophyllum wallichii* Rchb.f. | 1 | 1 | 1 | 1 | 1 | 1 | 1 | 1 | 1 | 1 | 1 | 1 | 1 | 1 | 1 | 1 | 1 | 1 | 1 | 1 | 1 | 1 | 1 | 0 | 1 | 0 | 1 | 1 | 1 | 1 | 1 | 1 | 1 | 1 | 1 | 1 | 1 | 1 | 1 | 1 | 1 | 0 | 1 | 1 | 1 | 1 | 1 | 1 | 0 | 1 | 1 | 1 | 0 | 1 | 1 | 1 | 1 | 1 | 1 | 1 | 1 | 1 | 1 |
| *Bulbophyllum weddellii* (Lindl.) Rchb.f. | 1 | 1 | 1 | 1 | 1 | 1 | 1 | 1 | 1 | 1 | 1 | 1 | 1 | 1 | 1 | 1 | 1 | 1 | 1 | 1 | 1 | 1 | 1 | 1 | 1 | 1 | 1 | 1 | 1 | 1 | 1 | 1 | 1 | 1 | 1 | 1 | 1 | 1 | 1 | 1 | 1 | 1 | 1 | 1 | 1 | 1 | 1 | 1 | 1 | 1 | 1 | 1 | 1 | 1 | 1 | 1 | 0 | 1 | 1 | 1 | 1 | 1 | 1 |
| *Bulbophyllum weinthalii* R.S.Rogers | 1 | 1 | 1 | 1 | 1 | 1 | 1 | 1 | 1 | 1 | 1 | 1 | 1 | 1 | 0 | 1 | 1 | 1 | 1 | 1 | 1 | 1 | 1 | 1 | 1 | 1 | 1 | 1 | 1 | 1 | 1 | 1 | 1 | 1 | 1 | 1 | 1 | 1 | 1 | 1 | 1 | 0 | 1 | 1 | 1 | 1 | 1 | 1 | 1 | 1 | 1 | 1 | 1 | 1 | 1 | 1 | 1 | 1 | 1 | 1 | 1 | 1 | 0 |
| *Bulbophyllum williamsii* A.D.Hawkes | 1 | 1 | 1 | 1 | 1 | 1 | 1 | 1 | 1 | 1 | 1 | 1 | 1 | 1 | 1 | 1 | 1 | 1 | 1 | 1 | 1 | 1 | 1 | 1 | 1 | 1 | 1 | 1 | 1 | 1 | 1 | 1 | 1 | 1 | 1 | 1 | 1 | 1 | 1 | 1 | 1 | 1 | 1 | 1 | 1 | 1 | 1 | 1 | 1 | 1 | 1 | 1 | 1 | 1 | 1 | 1 | 1 | 1 | 1 | 1 | 1 | 1 | 1 |
| *Bulbophyllum windsorense* B.Gray & D.L.Jones | 1 | 1 | 1 | 1 | 1 | 1 | 1 | 1 | 1 | 1 | 1 | 1 | 1 | 1 | 1 | 1 | 1 | 1 | 1 | 1 | 1 | 1 | 1 | 1 | 1 | 1 | 1 | 1 | 1 | 1 | 1 | 1 | 1 | 1 | 1 | 1 | 1 | 1 | 1 | 1 | 1 | 1 | 1 | 1 | 1 | 1 | 1 | 1 | 1 | 0 | 1 | 1 | 1 | 1 | 1 | 1 | 1 | 1 | 1 | 1 | 1 | 1 | 1 |
| *Bulbophyllum wolfei* B.Gray & D.L.Jones | 1 | 1 | 1 | 1 | 1 | 1 | 1 | 1 | 1 | 1 | 1 | 1 | 1 | 1 | 1 | 1 | 1 | 1 | 1 | 1 | 1 | 1 | 1 | 1 | 1 | 1 | 1 | 1 | 1 | 1 | 1 | 1 | 1 | 1 | 1 | 1 | 1 | 1 | 1 | 1 | 1 | 1 | 1 | 1 | 1 | 1 | 1 | 1 | 1 | 1 | 1 | 1 | 1 | 1 | 1 | 1 | 1 | 1 | 1 | 1 | 1 | 1 | 1 |
| *Bulbophyllum xylophyllum* C.S.P.Parish & Rchb.f. | 1 | 1 | 0 | 1 | 1 | 1 | 1 | 0 | 1 | 1 | 1 | 1 | 1 | 1 | 1 | 1 | 1 | 1 | 1 | 1 | 1 | 1 | 1 | 1 | 1 | 1 | 1 | 1 | 1 | 1 | 1 | 1 | 1 | 1 | 1 | 1 | 1 | 1 | 1 | 1 | 1 | 1 | 0 | 1 | 1 | 1 | 1 | 0 | 1 | 1 | 0 | 1 | 0 | 1 | 1 | 1 | 1 | 1 | 0 | 0 | 1 | 1 | 1 |
| *Bulbophyllum zebrinum* J.J.Sm. | 1 | 1 | 1 | 1 | 1 | 1 | 1 | 1 | 1 | 1 | 1 | 1 | 1 | 1 | 1 | 1 | 1 | 1 | 1 | 1 | 1 | 1 | 1 | 1 | 1 | 1 | 1 | 1 | 1 | 1 | 1 | 1 | 1 | 1 | 1 | 1 | 1 | 1 | 1 | 1 | 1 | 1 | 1 | 1 | 1 | 1 | 1 | 1 | 1 | 0 | 1 | 1 | 1 | 1 | 1 | 1 | 0 | 1 | 1 | 1 | 1 | 1 | 1 |
| **Outgroups** | | | | | | | | | | | | | | | | | | | | | | | | | | | | | | | | | | | | | | | | | | | | | | | | | | | | | | | | | | | | | | | |
| *Aa paleacea* (Kunth) Rchb.f. | 1 | 1 | 1 | 1 | 1 | 1 | 1 | 1 | 1 | 1 | 1 | 1 | 1 | 1 | 1 | 1 | 1 | 1 | 1 | 1 | 1 | 1 | 1 | 1 | 1 | 1 | 1 | 1 | 1 | 1 | 1 | 1 | 1 | 1 | 1 | 1 | 1 | 1 | 1 | 1 | 0 | 1 | 1 | 1 | 1 | 1 | 1 | 1 | 1 | 1 | 1 | 1 | 1 | 1 | 1 | 1 | 1 | 1 | 1 | 1 | 1 | 1 | 1 |
| *Acriopsis emarginata* D.L.Jones & M.A.Clem. | 1 | 1 | 1 | 1 | 1 | 1 | 1 | 1 | 1 | 1 | 1 | 1 | 1 | 1 | 1 | 1 | 1 | 1 | 1 | 1 | 1 | 1 | 1 | 1 | 1 | 1 | 1 | 1 | 1 | 1 | 1 | 1 | 1 | 1 | 1 | 1 | 1 | 1 | 1 | 1 | 1 | 1 | 1 | 1 | 1 | 1 | 1 | 1 | 1 | 0 | 1 | 1 | 1 | 1 | 1 | 1 | 0 | 1 | 1 | 1 | 1 | 1 | 1 |
| *Agapanthus coddii* F.M.Leight | 1 | 1 | 1 | 1 | 1 | 1 | 1 | 1 | 1 | 1 | 1 | 1 | 1 | 1 | 1 | 1 | 1 | 1 | 1 | 1 | 1 | 1 | 1 | 1 | 1 | 1 | 1 | 1 | 1 | 1 | 1 | 1 | 1 | 1 | 1 | 1 | 1 | 1 | 1 | 1 | 1 | 1 | 1 | 1 | 1 | 1 | 1 | 1 | 1 | 0 | 1 | 1 | 1 | 1 | 1 | 1 | 0 | 1 | 1 | 1 | 1 | 1 | 1 |
| *Agave americana* L. | 1 | 1 | 1 | 1 | 0 | 1 | 1 | 1 | 1 | 1 | 1 | 1 | 1 | 1 | 1 | 1 | 1 | 1 | 1 | 1 | 1 | 1 | 1 | 1 | 1 | 1 | 1 | 1 | 1 | 1 | 1 | 1 | 1 | 1 | 1 | 1 | 1 | 1 | 1 | 1 | 1 | 1 | 1 | 1 | 1 | 1 | 1 | 1 | 1 | 1 | 1 | 1 | 1 | 1 | 1 | 1 | 0 | 1 | 1 | 1 | 1 | 1 | 1 |
| *Angraecum sesquipedale* Thouars | 1 | 1 | 1 | 1 | 1 | 1 | 1 | 1 | 1 | 1 | 1 | 1 | 1 | 1 | 1 | 1 | 1 | 1 | 1 | 1 | 1 | 1 | 1 | 1 | 1 | 1 | 1 | 1 | 1 | 1 | 1 | 1 | 1 | 1 | 1 | 1 | 1 | 1 | 1 | 1 | 1 | 1 | 1 | 1 | 1 | 1 | 1 | 1 | 1 | 1 | 1 | 1 | 1 | 1 | 1 | 1 | 1 | 1 | 1 | 1 | 1 | 1 | 1 |
| *Anoectochilus emeiensis* K.Y.Lang | 1 | 1 | 1 | 1 | 1 | 1 | 1 | 1 | 1 | 1 | 1 | 1 | 1 | 1 | 1 | 1 | 1 | 1 | 1 | 1 | 1 | 1 | 1 | 1 | 1 | 1 | 1 | 1 | 1 | 1 | 1 | 1 | 1 | 1 | 1 | 1 | 1 | 1 | 1 | 1 | 1 | 1 | 1 | 1 | 1 | 1 | 1 | 1 | 1 | 0 | 1 | 1 | 1 | 1 | 1 | 1 | 0 | 1 | 1 | 1 | 1 | 1 | 1 |
| *Aphyllorchis montana* Rchb.f. | 1 | 1 | 1 | 1 | 1 | 1 | 1 | 1 | 1 | 1 | 1 | 1 | 1 | 1 | 1 | 1 | 1 | 1 | 1 | 1 | 1 | 1 | 1 | 0 | 1 | 0 | 1 | 1 | 1 | 1 | 1 | 1 | 1 | 1 | 1 | 1 | 1 | 1 | 1 | 1 | 1 | 0 | 1 | 1 | 1 | 1 | 1 | 1 | 0 | 1 | 1 | 1 | 0 | 1 | 1 | 1 | 1 | 1 | 1 | 1 | 1 | 1 | 1 |
| *Apostasia odorata* Blume | 1 | 1 | 1 | 1 | 1 | 1 | 1 | 1 | 1 | 1 | 1 | 1 | 1 | 1 | 1 | 1 | 1 | 1 | 1 | 1 | 1 | 1 | 1 | 1 | 1 | 1 | 1 | 1 | 1 | 1 | 1 | 1 | 1 | 1 | 1 | 1 | 1 | 1 | 1 | 1 | 1 | 1 | 1 | 1 | 1 | 1 | 1 | 1 | 1 | 1 | 1 | 1 | 1 | 1 | 1 | 1 | 0 | 1 | 1 | 1 | 1 | 1 | 1 |
| *Apostasia shenzhenica* Z.J.Liu & L.J.Chen | 1 | 1 | 1 | 1 | 1 | 1 | 1 | 1 | 1 | 1 | 1 | 1 | 1 | 1 | 0 | 1 | 1 | 1 | 1 | 1 | 1 | 1 | 1 | 1 | 1 | 1 | 1 | 1 | 1 | 1 | 1 | 1 | 1 | 1 | 1 | 1 | 1 | 1 | 1 | 1 | 1 | 0 | 1 | 1 | 1 | 1 | 1 | 1 | 1 | 1 | 1 | 1 | 1 | 1 | 1 | 1 | 1 | 1 | 1 | 1 | 1 | 1 | 0 |
| *Apostasia wallichii* R.Br. | 1 | 1 | 1 | 1 | 1 | 1 | 1 | 1 | 1 | 1 | 1 | 1 | 1 | 1 | 1 | 1 | 1 | 1 | 1 | 1 | 1 | 1 | 1 | 1 | 1 | 1 | 1 | 1 | 1 | 1 | 1 | 1 | 1 | 1 | 1 | 1 | 1 | 1 | 1 | 1 | 1 | 1 | 1 | 1 | 1 | 1 | 1 | 1 | 1 | 1 | 1 | 1 | 1 | 1 | 1 | 1 | 1 | 1 | 1 | 1 | 1 | 1 | 1 |
| *Arundina graminifolia* (D.Don) Hochr | 1 | 1 | 1 | 1 | 1 | 1 | 1 | 1 | 1 | 1 | 1 | 1 | 1 | 1 | 1 | 1 | 1 | 1 | 1 | 1 | 1 | 1 | 1 | 1 | 1 | 1 | 1 | 1 | 1 | 1 | 1 | 1 | 1 | 1 | 1 | 1 | 1 | 1 | 1 | 1 | 1 | 1 | 1 | 1 | 1 | 1 | 1 | 1 | 1 | 0 | 1 | 1 | 1 | 1 | 1 | 1 | 1 | 1 | 1 | 1 | 1 | 1 | 1 |
| *Asparagus officinalis* L. | 1 | 1 | 1 | 1 | 1 | 1 | 1 | 1 | 1 | 1 | 1 | 1 | 1 | 1 | 1 | 1 | 1 | 1 | 1 | 1 | 1 | 1 | 1 | 1 | 1 | 1 | 1 | 1 | 1 | 1 | 1 | 1 | 1 | 1 | 1 | 1 | 1 | 1 | 1 | 1 | 1 | 1 | 1 | 1 | 1 | 1 | 1 | 1 | 1 | 1 | 1 | 1 | 1 | 1 | 1 | 1 | 1 | 1 | 1 | 1 | 1 | 1 | 1 |
| *Bletilla striata* Rchb.f. | 1 | 1 | 0 | 1 | 1 | 1 | 1 | 0 | 1 | 1 | 1 | 1 | 1 | 1 | 1 | 1 | 1 | 1 | 1 | 1 | 1 | 1 | 1 | 1 | 1 | 1 | 1 | 1 | 1 | 1 | 1 | 1 | 1 | 1 | 1 | 1 | 1 | 1 | 1 | 1 | 1 | 1 | 0 | 1 | 1 | 1 | 1 | 0 | 1 | 1 | 0 | 1 | 0 | 1 | 1 | 1 | 1 | 1 | 0 | 0 | 1 | 1 | 1 |
| *Brachionidium cruziae* L.O.Williams | 1 | 1 | 1 | 1 | 1 | 1 | 1 | 1 | 1 | 1 | 1 | 1 | 1 | 1 | 1 | 1 | 1 | 1 | 1 | 1 | 1 | 1 | 1 | 1 | 1 | 1 | 1 | 1 | 1 | 1 | 1 | 1 | 1 | 1 | 1 | 1 | 1 | 1 | 1 | 1 | 1 | 1 | 1 | 1 | 1 | 1 | 1 | 1 | 1 | 0 | 1 | 1 | 1 | 1 | 1 | 1 | 0 | 1 | 1 | 1 | 1 | 1 | 1 |
| *Calanthe aristulifera* Rchb.f. | 1 | 1 | 1 | 1 | 0 | 1 | 1 | 1 | 1 | 0 | 1 | 1 | 1 | 1 | 1 | 1 | 1 | 1 | 1 | 1 | 1 | 0 | 1 | 1 | 1 | 1 | 1 | 1 | 1 | 1 | 1 | 1 | 1 | 1 | 1 | 1 | 1 | 1 | 1 | 1 | 1 | 1 | 1 | 1 | 1 | 1 | 1 | 1 | 1 | 1 | 1 | 1 | 1 | 1 | 1 | 1 | 1 | 1 | 1 | 1 | 1 | 1 | 1 |
| *Calanthe sylvatica* Lindl. | 1 | 1 | 1 | 1 | 0 | 1 | 1 | 1 | 1 | 1 | 1 | 1 | 1 | 1 | 1 | 1 | 1 | 1 | 1 | 1 | 1 | 0 | 1 | 1 | 1 | 1 | 1 | 1 | 1 | 1 | 1 | 1 | 1 | 1 | 1 | 1 | 1 | 1 | 1 | 1 | 1 | 1 | 1 | 1 | 1 | 1 | 1 | 1 | 1 | 1 | 1 | 1 | 1 | 1 | 1 | 1 | 1 | 1 | 1 | 1 | 1 | 1 | 1 |
| *Calopogon tuberosus* (L.) Britton, Sterns & Poggenb. | 0 | 1 | 1 | 1 | 1 | 1 | 1 | 1 | 1 | 0 | 1 | 1 | 1 | 1 | 1 | 1 | 1 | 1 | 0 | 1 | 1 | 0 | 1 | 1 | 1 | 1 | 1 | 1 | 1 | 1 | 1 | 1 | 1 | 1 | 1 | 1 | 1 | 0 | 1 | 1 | 1 | 1 | 1 | 1 | 1 | 1 | 1 | 1 | 1 | 1 | 0 | 1 | 1 | 1 | 1 | 1 | 1 | 1 | 1 | 1 | 1 | 1 | 1 |
| *Calypso bulbosa* var. *occidentalis* (Holz.) Cockerell | 0 | 1 | 1 | 1 | 1 | 1 | 1 | 1 | 1 | 0 | 1 | 1 | 1 | 1 | 1 | 1 | 1 | 1 | 1 | 1 | 1 | 0 | 1 | 1 | 0 | 1 | 1 | 1 | 1 | 1 | 1 | 1 | 1 | 1 | 1 | 1 | 0 | 0 | 1 | 1 | 1 | 0 | 1 | 1 | 1 | 0 | 1 | 1 | 1 | 1 | 0 | 1 | 1 | 1 | 0 | 1 | 1 | 1 | 0 | 1 | 0 | 1 | 1 |
| *Cattleya liliputana* (Pabst) Van den Berg | 1 | 1 | 1 | 1 | 1 | 1 | 1 | 1 | 1 | 1 | 1 | 1 | 1 | 1 | 1 | 1 | 1 | 1 | 1 | 1 | 1 | 0 | 1 | 1 | 1 | 1 | 1 | 1 | 1 | 1 | 1 | 1 | 1 | 1 | 1 | 1 | 1 | 1 | 1 | 0 | 1 | 1 | 1 | 1 | 1 | 1 | 1 | 1 | 1 | 1 | 1 | 1 | 1 | 1 | 1 | 1 | 1 | 1 | 1 | 0 | 1 | 1 | 1 |
| *Cephalanthera damasonium* Druce | 1 | 1 | 1 | 1 | 1 | 1 | 1 | 1 | 1 | 0 | 1 | 1 | 1 | 1 | 1 | 1 | 1 | 1 | 1 | 1 | 1 | 0 | 1 | 1 | 1 | 1 | 1 | 1 | 1 | 1 | 1 | 1 | 1 | 1 | 1 | 1 | 1 | 1 | 1 | 1 | 1 | 1 | 1 | 1 | 1 | 1 | 1 | 1 | 1 | 1 | 1 | 1 | 1 | 1 | 1 | 1 | 1 | 1 | 1 | 1 | 1 | 1 | 1 |
| *Cephalanthera rubra* (L.) Rich. | 1 | 1 | 1 | 1 | 1 | 1 | 1 | 1 | 1 | 0 | 1 | 1 | 1 | 1 | 1 | 1 | 1 | 1 | 1 | 1 | 1 | 0 | 1 | 1 | 1 | 1 | 1 | 1 | 1 | 1 | 1 | 1 | 1 | 1 | 1 | 1 | 1 | 1 | 1 | 1 | 1 | 1 | 1 | 1 | 1 | 1 | 1 | 1 | 1 | 1 | 1 | 1 | 1 | 1 | 1 | 1 | 1 | 1 | 1 | 1 | 1 | 1 | 1 |
| *Changnienia amoena* S.S.Chien | 1 | 1 | 1 | 1 | 1 | 1 | 1 | 1 | 1 | 1 | 1 | 1 | 1 | 1 | 1 | 1 | 1 | 1 | 1 | 1 | 1 | 0 | 1 | 1 | 1 | 1 | 1 | 1 | 1 | 1 | 1 | 1 | 1 | 1 | 1 | 1 | 1 | 1 | 1 | 1 | 1 | 1 | 1 | 1 | 1 | 1 | 1 | 1 | 1 | 1 | 1 | 1 | 1 | 1 | 1 | 1 | 1 | 1 | 1 | 1 | 1 | 1 | 1 |
| *Cheirostylis chinensis* Rolfe | 0 | 1 | 1 | 1 | 0 | 1 | 1 | 1 | 1 | 0 | 1 | 0 | 1 | 1 | 1 | 1 | 1 | 1 | 1 | 1 | 1 | 0 | 1 | 1 | 1 | 1 | 1 | 1 | 1 | 1 | 1 | 1 | 1 | 1 | 1 | 1 | 1 | 1 | 1 | 1 | 1 | 1 | 1 | 1 | 1 | 1 | 1 | 1 | 1 | 0 | 0 | 1 | 1 | 1 | 1 | 1 | 1 | 1 | 1 | 1 | 1 | 1 | 1 |
| *Codonorchis lessonii* Lindl. | 0 | 1 | 1 | 1 | 1 | 1 | 1 | 1 | 1 | 0 | 1 | 1 | 1 | 1 | 1 | 1 | 1 | 1 | 1 | 1 | 1 | 0 | 1 | 1 | 0 | 1 | 1 | 1 | 1 | 1 | 1 | 1 | 1 | 1 | 1 | 1 | 1 | 0 | 1 | 1 | 1 | 1 | 1 | 1 | 1 | 1 | 1 | 1 | 1 | 1 | 0 | 1 | 1 | 1 | 1 | 1 | 1 | 1 | 1 | 1 | 1 | 1 | 1 |
| *Coelogyne flaccida* Lindl. | 0 | 1 | 1 | 1 | 1 | 1 | 1 | 0 | 1 | 0 | 1 | 1 | 1 | 1 | 1 | 1 | 1 | 1 | 1 | 1 | 0 | 0 | 1 | 1 | 1 | 1 | 1 | 1 | 1 | 1 | 1 | 1 | 1 | 1 | 1 | 1 | 1 | 0 | 1 | 1 | 1 | 1 | 1 | 1 | 0 | 1 | 1 | 1 | 0 | 1 | 0 | 1 | 1 | 1 | 1 | 1 | 1 | 1 | 0 | 1 | 1 | 1 | 1 |
| *Corallorhiza striata* Lindl. | 1 | 1 | 1 | 1 | 1 | 1 | 1 | 0 | 0 | 0 | 1 | 1 | 0 | 0 | 0 | 0 | 1 | 0 | 0 | 0 | 0 | 0 | 1 | 0 | 0 | 1 | 0 | 0 | 1 | 1 | 0 | 0 | 1 | 0 | 1 | 1 | 1 | 1 | 0 | 1 | 1 | 1 | 1 | 1 | 1 | 1 | 1 | 0 | 0 | 0 | 0 | 1 | 1 | 1 | 1 | 1 | 1 | 1 | 1 | 1 | 1 | 0 | 0 |
| *Corallorhiza trifida* Châtel | 1 | 1 | 1 | 0 | 1 | 1 | 1 | 1 | 1 | 0 | 1 | 1 | 1 | 1 | 1 | 1 | 1 | 1 | 1 | 1 | 1 | 0 | 1 | 1 | 1 | 1 | 1 | 1 | 1 | 1 | 1 | 1 | 1 | 1 | 1 | 1 | 1 | 1 | 1 | 1 | 1 | 1 | 1 | 1 | 1 | 1 | 1 | 1 | 1 | 1 | 1 | 1 | 1 | 1 | 1 | 1 | 1 | 1 | 1 | 1 | 1 | 1 | 1 |
| *Cremastra appendiculata* (D.Don) Makino | 0 | 1 | 1 | 1 | 1 | 1 | 1 | 1 | 1 | 1 | 1 | 1 | 1 | 1 | 1 | 1 | 1 | 1 | 1 | 1 | 1 | 0 | 1 | 1 | 1 | 1 | 1 | 1 | 1 | 1 | 1 | 1 | 1 | 1 | 1 | 1 | 1 | 1 | 1 | 1 | 1 | 1 | 1 | 1 | 1 | 1 | 1 | 1 | 1 | 1 | 1 | 1 | 1 | 1 | 1 | 1 | 1 | 1 | 1 | 0 | 1 | 1 | 1 |
| *Cremastra unguiculata* Finet | 1 | 1 | 1 | 1 | 1 | 1 | 1 | 1 | 1 | 0 | 1 | 1 | 1 | 1 | 1 | 1 | 0 | 1 | 1 | 1 | 1 | 0 | 1 | 1 | 1 | 1 | 1 | 1 | 1 | 1 | 1 | 1 | 1 | 1 | 1 | 1 | 1 | 1 | 1 | 1 | 1 | 1 | 1 | 1 | 1 | 1 | 1 | 1 | 1 | 1 | 1 | 1 | 1 | 1 | 1 | 1 | 1 | 1 | 1 | 1 | 1 | 1 | 1 |
| *Cymbidium canaliculatum* R.Br. | 1 | 1 | 1 | 1 | 1 | 1 | 1 | 1 | 1 | 1 | 1 | 1 | 1 | 1 | 1 | 1 | 1 | 1 | 1 | 1 | 1 | 1 | 1 | 1 | 1 | 1 | 1 | 1 | 1 | 1 | 1 | 1 | 1 | 1 | 1 | 1 | 1 | 1 | 1 | 1 | 1 | 1 | 1 | 1 | 1 | 1 | 1 | 1 | 1 | 1 | 1 | 1 | 1 | 1 | 1 | 1 | 1 | 1 | 1 | 1 | 1 | 1 | 1 |
| *Cypripedium calceolus* L. | 1 | 1 | 1 | 1 | 1 | 1 | 1 | 1 | 0 | 0 | 1 | 1 | 1 | 1 | 1 | 1 | 1 | 1 | 1 | 1 | 1 | 0 | 1 | 1 | 1 | 1 | 1 | 1 | 1 | 1 | 1 | 1 | 1 | 1 | 1 | 1 | 1 | 1 | 1 | 1 | 1 | 1 | 1 | 1 | 1 | 0 | 1 | 1 | 1 | 0 | 1 | 1 | 1 | 1 | 1 | 1 | 1 | 1 | 1 | 1 | 1 | 1 | 1 |
| *Cypripedium japonicum* Thunb. | 1 | 1 | 1 | 1 | 1 | 1 | 1 | 1 | 0 | 0 | 1 | 0 | 1 | 1 | 1 | 1 | 1 | 1 | 1 | 1 | 1 | 0 | 1 | 1 | 1 | 1 | 1 | 1 | 1 | 1 | 1 | 1 | 1 | 1 | 1 | 1 | 1 | 1 | 1 | 1 | 1 | 1 | 1 | 1 | 1 | 1 | 1 | 1 | 1 | 0 | 1 | 1 | 1 | 1 | 1 | 1 | 1 | 1 | 1 | 1 | 1 | 1 | 0 |
| *Dendrobium kingianum* Bidwill ex Lindl. | 1 | 1 | 1 | 1 | 1 | 1 | 1 | 1 | 1 | 0 | 1 | 1 | 0 | 1 | 1 | 1 | 1 | 1 | 1 | 1 | 1 | 0 | 1 | 1 | 1 | 1 | 1 | 1 | 1 | 1 | 0 | 1 | 0 | 1 | 1 | 1 | 1 | 1 | 1 | 0 | 1 | 1 | 1 | 1 | 1 | 1 | 1 | 1 | 1 | 1 | 1 | 1 | 1 | 1 | 1 | 1 | 0 | 1 | 1 | 1 | 1 | 1 | 1 |
| *Dendrobium moniliforme* 1 (L.) Sw. | 1 | 1 | 1 | 1 | 1 | 1 | 1 | 1 | 1 | 0 | 1 | 1 | 1 | 1 | 1 | 1 | 1 | 1 | 1 | 1 | 1 | 0 | 1 | 1 | 1 | 1 | 1 | 1 | 1 | 1 | 1 | 1 | 1 | 1 | 1 | 1 | 1 | 1 | 1 | 1 | 1 | 1 | 1 | 1 | 1 | 1 | 1 | 1 | 1 | 1 | 0 | 1 | 1 | 1 | 1 | 1 | 1 | 1 | 1 | 1 | 1 | 1 | 1 |
| *Dendrobium moniliforme* 2 (L.) Sw. | 1 | 1 | 1 | 1 | 1 | 1 | 1 | 0 | 1 | 1 | 1 | 1 | 1 | 1 | 1 | 1 | 1 | 1 | 1 | 1 | 1 | 1 | 1 | 1 | 1 | 1 | 1 | 1 | 1 | 1 | 1 | 1 | 1 | 1 | 1 | 1 | 1 | 1 | 1 | 1 | 1 | 0 | 1 | 1 | 1 | 1 | 1 | 1 | 0 | 1 | 1 | 0 | 1 | 1 | 1 | 1 | 1 | 1 | 1 | 1 | 1 | 1 | 1 |
| *Dipodium pandanum* F.M.Bailey | 1 | 1 | 1 | 1 | 1 | 1 | 1 | 1 | 1 | 1 | 1 | 1 | 1 | 1 | 1 | 1 | 1 | 1 | 1 | 1 | 1 | 1 | 1 | 1 | 1 | 1 | 1 | 1 | 1 | 1 | 1 | 1 | 1 | 1 | 1 | 1 | 1 | 1 | 1 | 1 | 1 | 1 | 1 | 1 | 1 | 1 | 1 | 1 | 1 | 1 | 1 | 1 | 1 | 1 | 1 | 1 | 1 | 1 | 1 | 1 | 1 | 1 | 1 |
| *Dipodium roseum* D.L.Jones & M.A.Clem. | 1 | 1 | 1 | 1 | 1 | 1 | 1 | 1 | 1 | 1 | 1 | 1 | 1 | 1 | 1 | 1 | 1 | 1 | 1 | 1 | 1 | 1 | 1 | 1 | 1 | 1 | 1 | 1 | 1 | 1 | 1 | 1 | 1 | 1 | 1 | 1 | 0 | 1 | 1 | 1 | 1 | 1 | 1 | 1 | 1 | 1 | 1 | 1 | 1 | 1 | 1 | 1 | 1 | 1 | 1 | 1 | 1 | 1 | 1 | 1 | 1 | 1 | 1 |
| *Elleanthus sodiroi* Schltr. | 1 | 1 | 1 | 1 | 1 | 1 | 1 | 1 | 1 | 0 | 1 | 1 | 1 | 1 | 1 | 1 | 1 | 1 | 1 | 1 | 1 | 0 | 1 | 1 | 1 | 1 | 1 | 1 | 1 | 1 | 1 | 1 | 1 | 1 | 1 | 1 | 1 | 1 | 1 | 1 | 1 | 1 | 1 | 1 | 1 | 1 | 1 | 1 | 1 | 1 | 1 | 1 | 1 | 1 | 1 | 1 | 1 | 1 | 1 | 1 | 1 | 1 | 1 |
| *Epipactis mairei* Schltr. | 1 | 1 | 1 | 1 | 1 | 1 | 1 | 1 | 1 | 0 | 1 | 1 | 1 | 1 | 1 | 1 | 1 | 1 | 1 | 1 | 1 | 0 | 1 | 1 | 1 | 1 | 1 | 1 | 1 | 1 | 1 | 1 | 1 | 1 | 1 | 1 | 1 | 1 | 1 | 1 | 1 | 1 | 1 | 1 | 1 | 1 | 1 | 1 | 1 | 1 | 1 | 1 | 1 | 1 | 1 | 1 | 1 | 1 | 1 | 1 | 1 | 1 | 1 |
| *Epipactis thunbergia* A.Gray | 1 | 1 | 1 | 1 | 1 | 1 | 1 | 1 | 1 | 0 | 1 | 1 | 1 | 1 | 1 | 1 | 1 | 1 | 1 | 1 | 1 | 0 | 1 | 1 | 1 | 1 | 1 | 1 | 1 | 1 | 1 | 1 | 1 | 1 | 1 | 1 | 1 | 1 | 1 | 1 | 1 | 1 | 1 | 1 | 1 | 1 | 1 | 1 | 1 | 1 | 1 | 1 | 1 | 1 | 1 | 1 | 1 | 1 | 1 | 1 | 1 | 1 | 1 |
| *Epipactis veratrifolia* Boiss. & Hohen. | 1 | 1 | 1 | 1 | 1 | 1 | 1 | 1 | 1 | 0 | 1 | 1 | 1 | 1 | 1 | 1 | 1 | 1 | 1 | 1 | 1 | 0 | 1 | 1 | 1 | 1 | 1 | 1 | 1 | 1 | 1 | 1 | 1 | 1 | 1 | 1 | 1 | 1 | 1 | 1 | 1 | 1 | 1 | 1 | 1 | 1 | 1 | 1 | 1 | 1 | 1 | 1 | 1 | 1 | 1 | 1 | 1 | 1 | 1 | 1 | 1 | 1 | 1 |
| *Eria scabrilinguis* Lindl. | 1 | 1 | 1 | 1 | 1 | 1 | 1 | 1 | 0 | 0 | 1 | 1 | 1 | 1 | 1 | 1 | 1 | 1 | 1 | 1 | 1 | 0 | 1 | 1 | 1 | 1 | 1 | 1 | 1 | 1 | 1 | 1 | 1 | 1 | 1 | 1 | 1 | 1 | 1 | 1 | 1 | 1 | 1 | 1 | 1 | 1 | 1 | 1 | 1 | 1 | 1 | 1 | 1 | 1 | 1 | 1 | 1 | 1 | 1 | 1 | 1 | 1 | 1 |
| *Eulophia bicallosa* (D.Don) P.F.Hunt & Summerh. | 1 | 1 | 1 | 1 | 1 | 1 | 1 | 1 | 1 | 1 | 1 | 1 | 1 | 1 | 1 | 1 | 1 | 1 | 1 | 1 | 1 | 1 | 1 | 1 | 1 | 1 | 1 | 1 | 1 | 1 | 1 | 1 | 1 | 1 | 1 | 1 | 1 | 1 | 1 | 1 | 1 | 1 | 1 | 1 | 1 | 1 | 1 | 1 | 1 | 1 | 1 | 1 | 1 | 1 | 1 | 1 | 1 | 1 | 1 | 1 | 1 | 1 | 1 |
| *Eulophia nuda* Lindl. | 1 | 1 | 1 | 1 | 1 | 1 | 1 | 1 | 1 | 1 | 1 | 1 | 1 | 1 | 1 | 1 | 1 | 1 | 1 | 1 | 1 | 0 | 1 | 1 | 1 | 1 | 1 | 1 | 1 | 1 | 1 | 1 | 1 | 1 | 1 | 1 | 1 | 1 | 1 | 1 | 1 | 1 | 1 | 1 | 1 | 1 | 1 | 1 | 1 | 1 | 1 | 1 | 1 | 1 | 1 | 1 | 1 | 1 | 1 | 1 | 1 | 1 | 1 |
| *Evotella carnosa* (Lindl.) J.C.Manning & Goldblatt | 0 | 0 | 1 | 1 | 1 | 1 | 1 | 1 | 1 | 0 | 1 | 1 | 1 | 1 | 1 | 1 | 1 | 1 | 0 | 1 | 0 | 0 | 1 | 1 | 0 | 1 | 1 | 1 | 1 | 1 | 0 | 1 | 0 | 1 | 1 | 1 | 1 | 0 | 1 | 1 | 1 | 1 | 1 | 1 | 0 | 1 | 1 | 1 | 1 | 1 | 0 | 1 | 1 | 1 | 1 | 1 | 1 | 1 | 1 | 1 | 1 | 1 | 1 |
| *Gastrochilus japonicus* Schltr. | 1 | 1 | 1 | 1 | 1 | 1 | 1 | 1 | 1 | 0 | 1 | 1 | 1 | 1 | 1 | 1 | 1 | 1 | 1 | 1 | 1 | 0 | 1 | 1 | 1 | 1 | 1 | 1 | 1 | 1 | 1 | 1 | 1 | 1 | 1 | 1 | 1 | 1 | 1 | 1 | 1 | 1 | 1 | 1 | 1 | 1 | 1 | 1 | 1 | 0 | 1 | 1 | 1 | 1 | 1 | 1 | 1 | 1 | 1 | 1 | 1 | 1 | 1 |
| *Goodyera fumata* Thwaites | 1 | 1 | 1 | 1 | 0 | 1 | 1 | 0 | 1 | 0 | 1 | 0 | 1 | 1 | 1 | 1 | 1 | 1 | 1 | 1 | 1 | 0 | 1 | 1 | 1 | 1 | 1 | 1 | 1 | 1 | 1 | 1 | 1 | 1 | 1 | 1 | 1 | 1 | 1 | 1 | 1 | 1 | 1 | 1 | 1 | 1 | 1 | 1 | 1 | 0 | 0 | 1 | 1 | 1 | 1 | 1 | 1 | 1 | 1 | 1 | 1 | 1 | 1 |
| *Goodyera schlechtendaliana* Rchb.f. | 1 | 1 | 1 | 1 | 1 | 1 | 1 | 0 | 1 | 0 | 1 | 0 | 1 | 1 | 0 | 1 | 1 | 1 | 1 | 1 | 1 | 0 | 1 | 1 | 1 | 1 | 1 | 1 | 1 | 1 | 1 | 1 | 1 | 1 | 1 | 1 | 1 | 1 | 1 | 1 | 1 | 1 | 1 | 1 | 1 | 1 | 1 | 1 | 1 | 0 | 0 | 1 | 1 | 1 | 1 | 1 | 1 | 1 | 1 | 1 | 1 | 1 | 1 |
| *Guarianthe aurantiaca* (Bateman) Dressler & W.E.Higgins | 0 | 1 | 1 | 1 | 1 | 1 | 1 | 1 | 1 | 0 | 1 | 1 | 1 | 1 | 1 | 1 | 1 | 1 | 1 | 1 | 1 | 0 | 1 | 1 | 1 | 1 | 1 | 1 | 1 | 1 | 1 | 1 | 1 | 1 | 1 | 1 | 1 | 0 | 1 | 1 | 1 | 1 | 1 | 1 | 1 | 1 | 1 | 1 | 1 | 1 | 0 | 1 | 1 | 1 | 1 | 1 | 1 | 1 | 1 | 1 | 1 | 1 | 1 |
| *Gymnadenia conopsea* (L.) R.Br. | 1 | 1 | 1 | 1 | 0 | 1 | 1 | 1 | 1 | 0 | 1 | 1 | 1 | 1 | 1 | 1 | 1 | 1 | 1 | 1 | 1 | 0 | 1 | 1 | 1 | 1 | 1 | 1 | 1 | 1 | 1 | 1 | 1 | 1 | 1 | 1 | 1 | 1 | 1 | 1 | 1 | 1 | 1 | 1 | 1 | 1 | 1 | 1 | 1 | 0 | 0 | 1 | 1 | 1 | 1 | 1 | 1 | 1 | 1 | 1 | 1 | 1 | 1 |
| *Habenaria arenaria* Lindl. | 1 | 1 | 1 | 1 | 0 | 1 | 1 | 1 | 0 | 0 | 1 | 1 | 1 | 1 | 1 | 1 | 1 | 1 | 1 | 1 | 1 | 0 | 1 | 1 | 1 | 1 | 1 | 1 | 1 | 1 | 1 | 1 | 1 | 1 | 1 | 0 | 1 | 1 | 1 | 1 | 1 | 1 | 1 | 1 | 1 | 1 | 1 | 1 | 1 | 1 | 0 | 1 | 1 | 1 | 1 | 1 | 1 | 1 | 1 | 1 | 1 | 1 | 1 |
| *Habenaria ciliolaris* Kraenzl. | 1 | 1 | 1 | 1 | 1 | 1 | 1 | 1 | 1 | 0 | 1 | 1 | 1 | 1 | 1 | 1 | 1 | 1 | 1 | 1 | 1 | 0 | 1 | 1 | 1 | 1 | 1 | 1 | 1 | 1 | 1 | 1 | 1 | 1 | 1 | 1 | 1 | 1 | 1 | 1 | 1 | 1 | 1 | 1 | 1 | 1 | 1 | 1 | 1 | 1 | 0 | 1 | 1 | 1 | 1 | 1 | 1 | 1 | 1 | 1 | 1 | 1 | 1 |
| *Hexalectris warnockii* Ames & Correll | 1 | 0 | 0 | 0 | 0 | 1 | 0 | 0 | 0 | 0 | 1 | 1 | 0 | 0 | 0 | 1 | 1 | 0 | 0 | 0 | 0 | 0 | 0 | 0 | 0 | 0 | 0 | 1 | 0 | 0 | 0 | 1 | 0 | 1 | 1 | 0 | 0 | 1 | 0 | 1 | 1 | 1 | 1 | 1 | 1 | 1 | 1 | 0 | 0 | 0 | 0 | 1 | 1 | 1 | 1 | 1 | 1 | 1 | 1 | 1 | 1 | 0 | 0 |
| *Holcoglossum lingulatum* (Aver.) Aver. | 1 | 1 | 1 | 1 | 1 | 1 | 1 | 1 | 1 | 1 | 1 | 1 | 1 | 1 | 1 | 1 | 1 | 1 | 1 | 1 | 1 | 0 | 1 | 1 | 1 | 1 | 1 | 1 | 1 | 1 | 1 | 1 | 1 | 1 | 1 | 1 | 1 | 1 | 1 | 1 | 1 | 1 | 1 | 1 | 1 | 1 | 1 | 1 | 1 | 1 | 1 | 1 | 1 | 1 | 1 | 1 | 1 | 1 | 1 | 1 | 1 | 1 | 1 |

| | | | | | | | | | | | | | | | | | | | | | | | | | | | | | | | | | | | | | | | | | | | | | | | | | | | | | | | | | | | | | | | |
|---|---|---|---|---|---|---|---|---|---|---|---|---|---|---|---|---|---|---|---|---|---|---|---|---|---|---|---|---|---|---|---|---|---|---|---|---|---|---|---|---|---|---|---|---|---|---|---|---|---|---|---|---|---|---|---|---|---|---|---|---|---|---|---|
| *Holcoglossum subulifolium* (Rchb.f.) Christenson | 1 | 1 | 1 | 1 | 1 | 1 | 1 | 1 | 1 | 1 | 1 | 1 | 1 | 1 | 1 | 1 | 1 | 1 | 1 | 1 | 1 | 0 | 1 | 1 | 1 | 1 | 1 | 1 | 1 | 1 | 1 | 1 | 1 | 1 | 1 | 1 | 1 | 1 | 1 | 1 | 1 | 1 | 1 | 1 | 1 | 1 | 1 | 1 | 1 | 1 | 1 | 1 | 1 | 1 | 1 | 1 | 1 | 1 | 1 | 1 | 1 | 1 | 1 |
| *Iris sanguinea* Hornem | 1 | 1 | 1 | 1 | 0 | 1 | 1 | 0 | 1 | 0 | 1 | 1 | 1 | 1 | 1 | 1 | 1 | 1 | 1 | 1 | 1 | 0 | 1 | 1 | 1 | 1 | 1 | 1 | 1 | 1 | 1 | 1 | 0 | 1 | 1 | 1 | 1 | 1 | 1 | 0 | 1 | 1 | 1 | 1 | 1 | 1 | 1 | 1 | 1 | 0 | 1 | 1 | 1 | 1 | 1 | 1 | 1 | 1 | 1 | 1 | 1 | 1 | 1 |
| *Lilium pensylvanicum* Ker Gawl. | 1 | 1 | 1 | 1 | 0 | 1 | 1 | 0 | 1 | 1 | 0 | 0 | 1 | 1 | 1 | 1 | 1 | 1 | 1 | 1 | 1 | 0 | 1 | 1 | 1 | 1 | 1 | 1 | 1 | 1 | 1 | 1 | 1 | 1 | 1 | 1 | 1 | 1 | 1 | 1 | 1 | 1 | 1 | 1 | 1 | 1 | 1 | 1 | 1 | 0 | 1 | 1 | 1 | 1 | 1 | 1 | 1 | 1 | 1 | 1 | 1 | 0 | 1 |
| *Liparis auriculata* Blume ex Miq. | 1 | 1 | 1 | 1 | 1 | 1 | 1 | 1 | 1 | 0 | 1 | 0 | 1 | 1 | 1 | 1 | 1 | 1 | 1 | 1 | 1 | 0 | 1 | 1 | 1 | 1 | 1 | 1 | 1 | 1 | 1 | 1 | 1 | 1 | 1 | 1 | 1 | 1 | 1 | 1 | 1 | 1 | 1 | 1 | 1 | 1 | 1 | 1 | 1 | 0 | 1 | 1 | 1 | 1 | 1 | 1 | 1 | 1 | 1 | 1 | 1 | 1 | 1 |
| *Ludisia discolor* (Ker Gawl.) Blume | 0 | 1 | 1 | 1 | 1 | 1 | 1 | 0 | 1 | 1 | 1 | 0 | 1 | 1 | 1 | 1 | 1 | 1 | 1 | 1 | 1 | 0 | 1 | 1 | 0 | 1 | 1 | 1 | 1 | 1 | 1 | 1 | 1 | 1 | 1 | 1 | 0 | 0 | 1 | 1 | 1 | 1 | 1 | 1 | 1 | 1 | 1 | 1 | 1 | 1 | 0 | 1 | 1 | 1 | 1 | 1 | 1 | 1 | 1 | 1 | 1 | 1 | 1 |
| *Masdevallia coccinea* Linden ex Lindl. | 1 | 1 | 1 | 1 | 1 | 1 | 1 | 1 | 1 | 0 | 1 | 1 | 1 | 1 | 1 | 1 | 1 | 1 | 1 | 1 | 1 | 0 | 1 | 1 | 1 | 1 | 1 | 1 | 1 | 1 | 1 | 1 | 1 | 1 | 1 | 1 | 1 | 1 | 1 | 1 | 1 | 1 | 1 | 1 | 1 | 1 | 1 | 1 | 1 | 1 | 1 | 1 | 1 | 1 | 1 | 1 | 1 | 1 | 1 | 1 | 1 | 1 | 1 |
| *Neottia listeroides* Lindl. | 1 | 0 | 0 | 0 | 0 | 0 | 0 | 0 | 0 | 1 | 1 | 1 | 0 | 0 | 0 | 0 | 0 | 0 | 0 | 0 | 0 | 0 | 0 | 0 | 0 | 0 | 0 | 0 | 0 | 0 | 0 | 0 | 0 | 0 | 0 | 0 | 0 | 0 | 0 | 1 | 1 | 1 | 1 | 1 | 1 | 1 | 1 | 0 | 0 | 0 | 0 | 1 | 1 | 1 | 1 | 1 | 1 | 1 | 0 | 1 | 1 | 0 | 0 |
| *Neottia ovata* Bluff & Fingerh. | 1 | 1 | 1 | 1 | 1 | 1 | 1 | 1 | 1 | 0 | 1 | 1 | 1 | 1 | 1 | 1 | 1 | 1 | 1 | 1 | 1 | 0 | 1 | 1 | 1 | 1 | 1 | 1 | 1 | 1 | 1 | 1 | 1 | 1 | 1 | 1 | 1 | 1 | 1 | 1 | 1 | 1 | 1 | 1 | 1 | 1 | 1 | 1 | 1 | 1 | 1 | 1 | 1 | 1 | 1 | 1 | 1 | 1 | 1 | 1 | 1 | 1 | 1 |
| *Nervilia crociformis* (Zoll. & Moritzi) Seidenf. | 0 | 0 | 1 | 1 | 0 | 0 | 1 | 0 | 0 | 0 | 1 | 0 | 0 | 1 | 1 | 1 | 0 | 0 | 1 | 1 | 0 | 0 | 0 | 1 | 1 | 0 | 0 | 1 | 1 | 1 | 0 | 0 | 1 | 1 | 0 | 1 | 0 | 0 | 1 | 1 | 1 | 0 | 1 | 1 | 0 | 1 | 0 | 1 | 0 | 1 | 0 | 1 | 1 | 1 | 0 | 1 | 1 | 1 | 0 | 0 | 0 | 1 | 1 |
| *Neuwiedia zollingeri* var. *singapureana* (Wall. ex Baker) de Vogel | 0 | 1 | 1 | 1 | 1 | 1 | 1 | 0 | 1 | 1 | 1 | 0 | 1 | 1 | 1 | 1 | 1 | 1 | 1 | 1 | 1 | 0 | 1 | 1 | 1 | 1 | 1 | 1 | 1 | 1 | 1 | 1 | 1 | 1 | 1 | 1 | 1 | 1 | 1 | 0 | 1 | 1 | 1 | 1 | 1 | 1 | 1 | 1 | 1 | 1 | 1 | 1 | 1 | 1 | 1 | 1 | 0 | 1 | 1 | 0 | 1 | 0 | 1 |
| *Oberonia japonica* (Maxim.) Makino | 1 | 1 | 1 | 1 | 1 | 1 | 1 | 1 | 1 | 0 | 1 | 1 | 1 | 1 | 1 | 1 | 1 | 1 | 1 | 1 | 1 | 0 | 1 | 1 | 1 | 1 | 1 | 1 | 1 | 1 | 1 | 1 | 1 | 1 | 1 | 1 | 1 | 1 | 1 | 1 | 1 | 1 | 1 | 1 | 1 | 1 | 1 | 1 | 1 | 1 | 1 | 1 | 1 | 1 | 1 | 1 | 1 | 1 | 1 | 1 | 1 | 1 | 1 |
| *Ophrys fusca* subsp. *iricolor* (Desf.) K.Richt. | 1 | 1 | 1 | 1 | 1 | 1 | 1 | 1 | 1 | 1 | 1 | 1 | 1 | 1 | 1 | 1 | 1 | 1 | 1 | 1 | 1 | 0 | 1 | 1 | 1 | 1 | 1 | 1 | 1 | 1 | 1 | 1 | 1 | 1 | 1 | 1 | 1 | 1 | 1 | 1 | 1 | 1 | 0 | 1 | 1 | 1 | 1 | 1 | 0 | 0 | 1 | 1 | 1 | 1 | 1 | 1 | 1 | 1 | 1 | 1 | 1 | 1 | 1 |
| *Ophrys sphegodes* Mill. | 1 | 1 | 1 | 1 | 1 | 1 | 1 | 1 | 1 | 1 | 1 | 1 | 1 | 1 | 1 | 1 | 1 | 1 | 1 | 1 | 1 | 0 | 1 | 1 | 1 | 1 | 1 | 1 | 1 | 1 | 1 | 1 | 1 | 1 | 1 | 1 | 1 | 1 | 1 | 1 | 1 | 1 | 0 | 1 | 1 | 1 | 1 | 1 | 0 | 0 | 1 | 1 | 1 | 1 | 1 | 1 | 1 | 1 | 1 | 1 | 1 | 1 | 1 |
| *Palmorchis pabstii* Veyret. | 1 | 1 | 1 | 1 | 1 | 1 | 1 | 1 | 1 | 0 | 1 | 1 | 1 | 1 | 1 | 1 | 1 | 1 | 1 | 1 | 1 | 0 | 1 | 1 | 1 | 1 | 1 | 1 | 1 | 1 | 1 | 1 | 1 | 1 | 1 | 1 | 1 | 1 | 1 | 1 | 1 | 1 | 1 | 1 | 1 | 1 | 1 | 1 | 1 | 1 | 1 | 1 | 1 | 1 | 1 | 1 | 1 | 1 | 1 | 1 | 1 | 1 | 1 |
| *Paphiopedilum armeniacum* S.C.Chen & F.Y.Liu | 0 | 1 | 1 | 1 | 1 | 1 | 1 | 1 | 1 | 0 | 1 | 1 | 1 | 1 | 1 | 1 | 1 | 1 | 1 | 1 | 1 | 0 | 1 | 1 | 1 | 1 | 1 | 1 | 1 | 1 | 1 | 1 | 1 | 1 | 1 | 1 | 1 | 1 | 1 | 1 | 1 | 1 | 1 | 1 | 1 | 1 | 1 | 1 | 1 | 1 | 1 | 1 | 1 | 1 | 1 | 1 | 1 | 1 | 1 | 1 | 1 | 1 | 1 |
| *Paphiopedilum niveum* (Rchb.f.) Stein | 0 | 1 | 1 | 1 | 1 | 1 | 1 | 1 | 1 | 1 | 0 | 1 | 1 | 1 | 1 | 1 | 1 | 1 | 1 | 1 | 1 | 0 | 1 | 1 | 1 | 1 | 1 | 1 | 1 | 1 | 1 | 1 | 1 | 1 | 1 | 1 | 1 | 1 | 1 | 1 | 1 | 1 | 1 | 1 | 1 | 1 | 1 | 1 | 1 | 0 | 1 | 1 | 1 | 1 | 1 | 1 | 1 | 1 | 1 | 1 | 1 | 1 | 1 |
| *Pelatantheria scolopendrifolia* (Makino) Aver. | 1 | 1 | 1 | 1 | 1 | 1 | 1 | 1 | 1 | 0 | 1 | 1 | 1 | 1 | 1 | 1 | 1 | 1 | 1 | 1 | 1 | 0 | 1 | 1 | 1 | 1 | 1 | 1 | 1 | 1 | 1 | 1 | 1 | 1 | 1 | 1 | 1 | 1 | 1 | 1 | 1 | 1 | 1 | 1 | 1 | 1 | 1 | 1 | 1 | 0 | 1 | 1 | 1 | 1 | 1 | 1 | 1 | 1 | 1 | 1 | 1 | 1 | 1 |
| *Phaius tankervilleae* (Banks) Blume | 0 | 1 | 1 | 1 | 1 | 1 | 1 | 1 | 1 | 0 | 1 | 1 | 1 | 1 | 1 | 1 | 1 | 1 | 1 | 1 | 1 | 0 | 1 | 1 | 0 | 1 | 1 | 1 | 1 | 1 | 1 | 1 | 1 | 1 | 1 | 1 | 1 | 0 | 1 | 1 | 1 | 1 | 1 | 1 | 1 | 1 | 1 | 1 | 1 | 1 | 0 | 1 | 1 | 1 | 1 | 1 | 1 | 1 | 1 | 1 | 1 | 1 | 1 |
| *Phalaenopsis japonica* (Rchb.f.) Kocyan & Schuit. | 1 | 1 | 1 | 1 | 1 | 1 | 1 | 1 | 1 | 0 | 1 | 1 | 1 | 1 | 1 | 1 | 1 | 1 | 1 | 1 | 1 | 0 | 1 | 1 | 1 | 1 | 1 | 1 | 1 | 1 | 1 | 1 | 1 | 1 | 1 | 1 | 1 | 1 | 1 | 1 | 1 | 1 | 1 | 1 | 1 | 1 | 1 | 1 | 1 | 1 | 1 | 1 | 1 | 1 | 1 | 1 | 1 | 1 | 1 | 1 | 1 | 1 | 1 |
| *Phalaenopsis pulcherrima* (Lindl.) J.J.Sm. | 0 | 1 | 1 | 1 | 1 | 1 | 1 | 1 | 1 | 0 | 1 | 1 | 1 | 1 | 1 | 1 | 1 | 1 | 1 | 1 | 1 | 0 | 1 | 1 | 1 | 1 | 1 | 1 | 1 | 1 | 1 | 1 | 1 | 1 | 1 | 1 | 1 | 0 | 1 | 1 | 0 | 1 | 1 | 1 | 1 | 1 | 1 | 1 | 1 | 0 | 1 | 1 | 1 | 1 | 1 | 1 | 1 | 1 | 1 | 1 | 1 | 1 | 1 |
| *Phragmipedium longifolium* (Rchb.f. & Warsz.) Rolfe | 0 | 1 | 1 | 1 | 1 | 1 | 1 | 1 | 1 | 1 | 1 | 1 | 1 | 1 | 1 | 1 | 1 | 1 | 1 | 1 | 1 | 0 | 1 | 1 | 1 | 1 | 1 | 1 | 1 | 1 | 1 | 1 | 1 | 1 | 1 | 1 | 1 | 1 | 1 | 1 | 1 | 1 | 1 | 1 | 1 | 1 | 1 | 1 | 1 | 1 | 1 | 1 | 1 | 1 | 1 | 1 | 1 | 1 | 1 | 0 | 1 | 1 | 1 |
| *Platanthera mandarinorum* Rchb.f. | 1 | 1 | 1 | 1 | 1 | 1 | 1 | 1 | 1 | 0 | 1 | 1 | 1 | 1 | 1 | 1 | 1 | 1 | 1 | 1 | 1 | 0 | 1 | 1 | 1 | 1 | 1 | 1 | 1 | 1 | 1 | 1 | 1 | 1 | 1 | 1 | 1 | 1 | 1 | 1 | 1 | 1 | 1 | 1 | 1 | 1 | 1 | 1 | 1 | 1 | 1 | 1 | 1 | 1 | 1 | 1 | 1 | 1 | 1 | 1 | 1 | 1 | 1 |
| *Platystele aurea* Garay | 0 | 1 | 1 | 1 | 1 | 1 | 1 | 1 | 1 | 0 | 1 | 1 | 1 | 1 | 1 | 1 | 1 | 1 | 1 | 1 | 1 | 0 | 1 | 1 | 1 | 1 | 1 | 1 | 1 | 1 | 1 | 1 | 1 | 1 | 1 | 0 | 1 | 1 | 1 | 1 | 1 | 1 | 1 | 1 | 1 | 1 | 1 | 1 | 1 | 1 | 1 | 1 | 1 | 1 | 1 | 1 | 1 | 1 | 1 | 1 | 1 | 1 | 1 |
| *Pleione formosana* Hayata | 1 | 1 | 1 | 1 | 1 | 1 | 1 | 1 | 1 | 1 | 1 | 1 | 1 | 1 | 1 | 1 | 1 | 1 | 1 | 1 | 1 | 0 | 1 | 1 | 1 | 1 | 1 | 1 | 1 | 1 | 1 | 1 | 1 | 1 | 1 | 1 | 1 | 1 | 1 | 1 | 1 | 1 | 1 | 1 | 1 | 1 | 1 | 1 | 1 | 1 | 1 | 1 | 1 | 1 | 1 | 1 | 1 | 1 | 1 | 1 | 1 | 1 | 1 |
| *Pogonia ophioglossoides* (L.) Ker Gawl. | 0 | 1 | 1 | 1 | 1 | 1 | 1 | 0 | 0 | 0 | 1 | 1 | 1 | 1 | 1 | 1 | 1 | 1 | 1 | 1 | 0 | 0 | 1 | 1 | 0 | 1 | 1 | 1 | 1 | 0 | 0 | 1 | 1 | 1 | 1 | 1 | 0 | 0 | 1 | 1 | 1 | 0 | 1 | 1 | 0 | 1 | 1 | 1 | 1 | 1 | 0 | 1 | 1 | 1 | 1 | 1 | 1 | 1 | 1 | 1 | 0 | 0 | 1 |
| *Ponerorchis gracilis* (Blume) X.H.Jin, Schuit. & W.T.Jin | 1 | 1 | 1 | 1 | 1 | 1 | 1 | 1 | 1 | 0 | 1 | 1 | 1 | 1 | 1 | 1 | 1 | 1 | 1 | 1 | 1 | 0 | 1 | 1 | 1 | 1 | 1 | 1 | 1 | 1 | 1 | 1 | 1 | 1 | 1 | 1 | 1 | 1 | 1 | 1 | 1 | 1 | 0 | 1 | 1 | 1 | 1 | 1 | 1 | 1 | 0 | 1 | 1 | 1 | 1 | 1 | 1 | 1 | 1 | 1 | 1 | 1 | 1 |
| *Scaphosepalum antenniferum* Rolfe | 1 | 1 | 1 | 1 | 1 | 1 | 1 | 1 | 1 | 1 | 1 | 1 | 1 | 0 | 0 | 1 | 1 | 1 | 1 | 1 | 1 | 0 | 1 | 1 | 1 | 1 | 0 | 0 | 0 | 0 | 1 | 1 | 1 | 1 | 1 | 1 | 1 | 1 | 1 | 1 | 0 | 1 | 1 | 1 | 1 | 1 | 1 | 1 | 1 | 1 | 1 | 1 | 1 | 1 | 1 | 1 | 0 | 1 | 1 | 1 | 1 | 1 | 1 |
| *Sobralia mucronata* Ames & C.Schweinf. | 1 | 1 | 1 | 1 | 1 | 1 | 1 | 1 | 1 | 1 | 1 | 1 | 1 | 0 | 0 | 1 | 1 | 1 | 1 | 1 | 1 | 0 | 1 | 1 | 1 | 1 | 0 | 0 | 0 | 0 | 1 | 1 | 1 | 1 | 1 | 1 | 1 | 1 | 1 | 1 | 0 | 1 | 1 | 1 | 1 | 1 | 1 | 1 | 1 | 1 | 1 | 1 | 1 | 1 | 1 | 1 | 0 | 1 | 1 | 1 | 1 | 1 | 1 |
| *Spiranthes sinensis* (Pers.) Ames | 1 | 1 | 1 | 1 | 1 | 1 | 1 | 1 | 1 | 0 | 1 | 0 | 1 | 1 | 1 | 1 | 1 | 1 | 1 | 1 | 1 | 0 | 1 | 1 | 1 | 1 | 1 | 1 | 1 | 1 | 1 | 1 | 1 | 1 | 1 | 1 | 1 | 1 | 1 | 1 | 1 | 1 | 1 | 1 | 1 | 1 | 1 | 1 | 1 | 1 | 0 | 1 | 1 | 1 | 1 | 1 | 1 | 1 | 0 | 1 | 1 | 1 | 1 |
| *Teagueia* (Luer) Luer | 1 | 1 | 1 | 1 | 1 | 1 | 1 | 1 | 1 | 1 | 1 | 1 | 1 | 0 | 0 | 1 | 1 | 1 | 1 | 1 | 1 | 0 | 1 | 1 | 1 | 1 | 0 | 0 | 0 | 0 | 1 | 1 | 1 | 1 | 1 | 1 | 1 | 1 | 1 | 1 | 0 | 1 | 1 | 1 | 1 | 1 | 1 | 1 | 1 | 1 | 1 | 1 | 1 | 1 | 1 | 1 | 0 | 1 | 1 | 1 | 1 | 1 | 1 |
| *Thelymitra cyanea* (Lindl.) Benth. | 0 | 1 | 1 | 1 | 1 | 1 | 1 | 1 | 1 | 0 | 1 | 0 | 1 | 1 | 1 | 1 | 1 | 1 | 1 | 1 | 0 | 0 | 1 | 1 | 1 | 1 | 1 | 1 | 1 | 1 | 1 | 1 | 1 | 1 | 1 | 1 | 1 | 0 | 1 | 1 | 1 | 1 | 1 | 1 | 1 | 1 | 1 | 1 | 1 | 1 | 0 | 1 | 1 | 1 | 1 | 1 | 1 | 1 | 0 | 1 | 1 | 1 | 1 |
| *Thrixspermum japonicum* Rchb.f. | 1 | 1 | 1 | 1 | 1 | 1 | 1 | 1 | 1 | 0 | 1 | 1 | 1 | 1 | 1 | 1 | 1 | 1 | 1 | 1 | 1 | 0 | 1 | 1 | 1 | 1 | 1 | 1 | 1 | 1 | 1 | 1 | 1 | 1 | 1 | 1 | 1 | 1 | 1 | 1 | 1 | 1 | 1 | 1 | 1 | 1 | 1 | 1 | 1 | 0 | 1 | 1 | 1 | 1 | 1 | 1 | 1 | 1 | 1 | 1 | 1 | 1 | 1 |
| *Triphora trianthophora* (Sw.) Rydb. | 0 | 1 | 1 | 1 | 1 | 1 | 1 | 1 | 1 | 0 | 1 | 0 | 1 | 1 | 1 | 1 | 1 | 1 | 1 | 1 | 1 | 0 | 1 | 1 | 0 | 1 | 1 | 1 | 1 | 1 | 1 | 1 | 1 | 1 | 1 | 1 | 1 | 0 | 1 | 1 | 1 | 1 | 1 | 1 | 1 | 1 | 1 | 1 | 1 | 1 | 0 | 1 | 1 | 1 | 0 | 1 | 1 | 1 | 1 | 1 | 1 | 1 | 1 |
| *Tropidia polystachya* Ames | 0 | 1 | 1 | 1 | 0 | 1 | 1 | 1 | 1 | 0 | 0 | 0 | 1 | 1 | 1 | 1 | 0 | 0 | 1 | 1 | 1 | 0 | 0 | 1 | 1 | 1 | 1 | 1 | 1 | 1 | 0 | 1 | 0 | 1 | 0 | 1 | 0 | 0 | 1 | 1 | 1 | 1 | 1 | 1 | 0 | 1 | 1 | 1 | 1 | 1 | 0 | 1 | 1 | 1 | 0 | 1 | 1 | 1 | 0 | 0 | 0 | 1 | 1 |
| *Vanda brunnea* Rchb.f. | 1 | 1 | 1 | 1 | 1 | 1 | 1 | 1 | 1 | 1 | 1 | 1 | 1 | 1 | 1 | 1 | 1 | 1 | 1 | 1 | 1 | 0 | 1 | 1 | 1 | 1 | 1 | 1 | 1 | 1 | 1 | 1 | 1 | 1 | 1 | 1 | 1 | 1 | 1 | 1 | 1 | 1 | 1 | 1 | 1 | 1 | 1 | 1 | 1 | 1 | 1 | 1 | 1 | 1 | 1 | 1 | 1 | 1 | 1 | 1 | 1 | 1 | 1 |
| *Vanda falcata* Beer | 1 | 1 | 1 | 1 | 1 | 1 | 1 | 0 | 1 | 0 | 1 | 1 | 1 | 1 | 1 | 1 | 1 | 1 | 1 | 1 | 1 | 0 | 1 | 1 | 1 | 1 | 1 | 1 | 1 | 1 | 1 | 1 | 1 | 1 | 1 | 1 | 1 | 1 | 1 | 1 | 1 | 1 | 1 | 1 | 1 | 1 | 1 | 1 | 1 | 0 | 1 | 1 | 1 | 1 | 1 | 1 | 1 | 1 | 1 | 1 | 1 | 1 | 1 |
| *Vanilla aphylla* Blume | 0 | 1 | 1 | 1 | 0 | 1 | 1 | 1 | 0 | 0 | 1 | 0 | 1 | 1 | 0 | 1 | 1 | 1 | 1 | 1 | 1 | 0 | 1 | 1 | 1 | 1 | 1 | 1 | 1 | 1 | 0 | 1 | 1 | 1 | 1 | 1 | 1 | 1 | 1 | 0 | 1 | 1 | 1 | 1 | 0 | 1 | 1 | 1 | 0 | 0 | 0 | 1 | 1 | 1 | 1 | 1 | 0 | 1 | 1 | 0 | 1 | 0 | 1 |
| *Vanilla planifolia* Andrews | 0 | 1 | 1 | 1 | 1 | 1 | 1 | 0 | 1 | 0 | 1 | 0 | 1 | 1 | 1 | 1 | 1 | 1 | 0 | 1 | 1 | 0 | 1 | 1 | 1 | 1 | 1 | 1 | 1 | 1 | 1 | 1 | 1 | 1 | 1 | 1 | 1 | 0 | 1 | 1 | 1 | 1 | 1 | 1 | 1 | 1 | 1 | 1 | 1 | 0 | 0 | 1 | 1 | 1 | 0 | 1 | 1 | 0 | 1 | 1 | 1 | 1 | 1 |
| *Vanilla pompona* Schiede | 0 | 1 | 1 | 1 | 1 | 1 | 1 | 1 | 0 | 0 | 1 | 0 | 1 | 1 | 1 | 1 | 1 | 1 | 1 | 1 | 1 | 0 | 1 | 1 | 1 | 1 | 1 | 1 | 1 | 1 | 1 | 1 | 1 | 1 | 1 | 1 | 1 | 1 | 1 | 1 | 1 | 1 | 1 | 1 | 0 | 1 | 1 | 1 | 0 | 1 | 0 | 1 | 1 | 1 | 1 | 1 | 1 | 1 | 1 | 1 | 1 | 1 | 1 |
| *Zootrophion hirtzii* Luer | 1 | 1 | 1 | 1 | 1 | 1 | 1 | 1 | 1 | 0 | 1 | 1 | 1 | 1 | 1 | 1 | 1 | 1 | 1 | 1 | 1 | 0 | 1 | 1 | 1 | 1 | 1 | 1 | 1 | 1 | 1 | 1 | 1 | 1 | 1 | 0 | 1 | 1 | 1 | 1 | 1 | 1 | 1 | 1 | 1 | 1 | 1 | 1 | 1 | 1 | 1 | 1 | 1 | 1 | 1 | 1 | 1 | 1 | 1 | 1 | 1 | 1 | 1 |

**Legend:** 1=Present 0=Absent

Number of genes included in phylogenetic analysis = 63

Number of genes recovered in the complete dataset of 468 samples = 27924

Number of missing genes across the dataset =1560

Percentage of missing data = 5.58%

### Figure S1

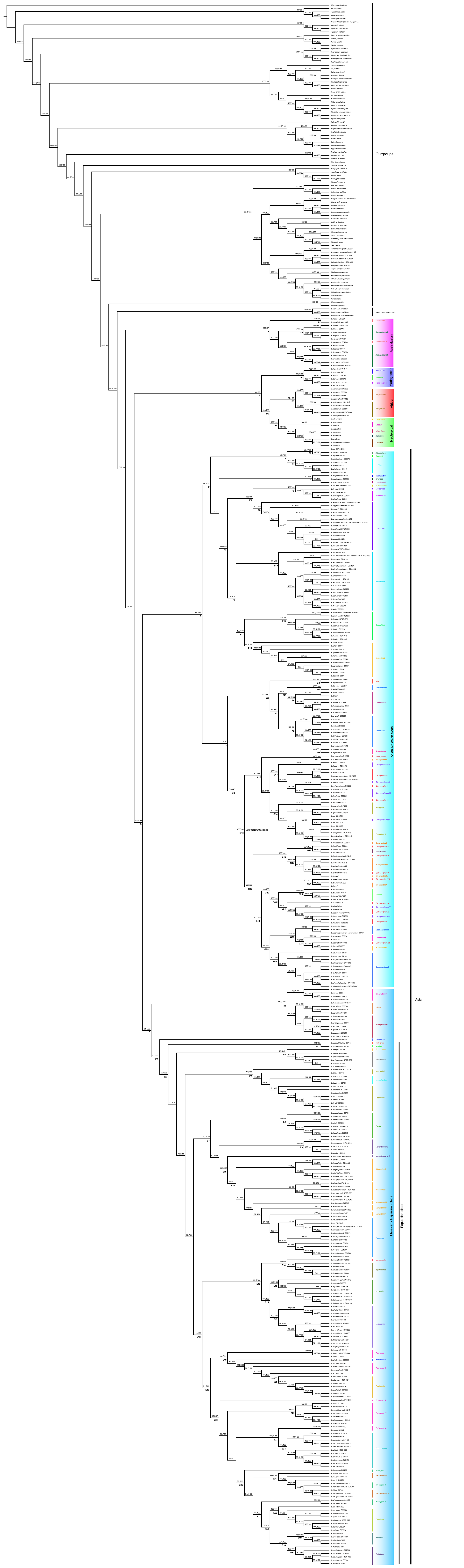

**Figure S1:** Maximum likelihood phylogenetic tree showing the relationships between *Bulbophyllum*, its sister group *Dendrobium*, and other outgroups representing all major lineages of Orchidaceae. The tree also illustrates the relationships among major biogeographic lineages within *Bulbophyllum*. The tree is based on 468 taxa and was constructed using 63 plastid loci. Branch support values are shown at each node, with SH-aLRT (Shimodaira-Hasegawa-like Approximate Likelihood Ratio Test) values presented first, followed by UFBoot (Ultrafast Bootstrap) support values.

#### Figure S2

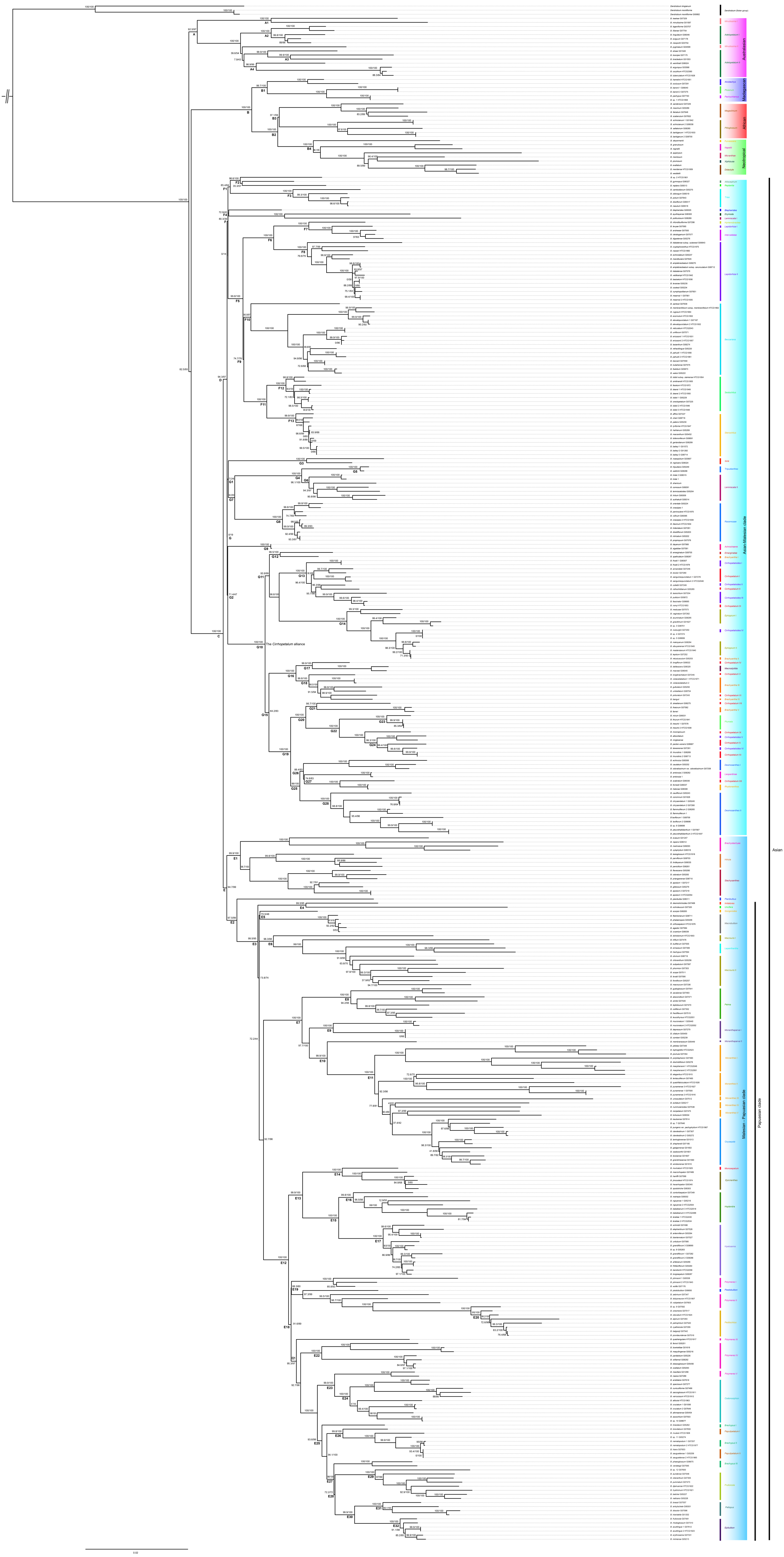

**Figure S2:** Maximum Likelihood phylogenetic tree with original branch lengths showing the relationships between *Bulbophyllum*, its sister group *Dendrobium*, with all other outgroups pruned. The tree also illustrates the relationships among major biogeographic lineages within *Bulbophyllum*.The tree is based on 468 taxa and was constructed using 63 plastid loci. Branch support values are shown at each node, with SH-aLRT (Shimodaira Hasegawa-Like Approximate Likelihood Ratio Test) values presented first, followed by UFBoot (Ultrafast Bootstrap) support values. The scale bar indicates branch length, corresponding to 0.02 substitutions per site.
